# Out of the abyss: Genome and metagenome mining reveals unexpected environmental distribution of abyssomicins

**DOI:** 10.1101/789859

**Authors:** Alba Iglesias, Adriel Latorre-Pérez, James E. M. Stach, Manuel Porcar, Javier Pascual

## Abstract

Natural products have traditionally been discovered through the screening of culturable microbial isolates from all sort of environments. The sequencing revolution allowed the identification of dozens of biosynthetic gene clusters (BGCs) within single bacterial genomes, either from cultured or uncultured strains. However, we are still far from fully exploiting the microbial reservoir, as most of the species are non-model organisms with complex regulatory systems and yet recalcitrant to be engineered. Today, genomic and metagenomic data produced by laboratories worldwide covering the most different natural and artificial environments on Earth, are an invaluable source of raw information from which natural product biosynthesis can be accessed. In the present work, we describe the environmental distribution and evolution of the abyssomicin BGC through the analysis of publicly available genomic and metagenomic data. Our results demonstrate that the selection of a pathway-specific enzyme to direct the genome mining is an excellent strategy that led to the identification of 74 new Diels-Alderase homologs and unveiled a surprising prevalence of the abyssomicin BGC within terrestrial habitats, mainly soil and plant-associated, where we have identified five complete and 12 partial new abyssomicin BGCs and 23 new potential abyssomicin BGCs. Our results strongly support the potential of genome and metagenome mining as a key preliminary tool to inform bioprospecting strategies aiming at the identification of new bioactive compounds such as -but not restricted to-abyssomicins.

## Introduction

Natural products are the main source of pharmaceutically interesting biomolecules. In particular, the search of microbial specialised metabolites has yielded a broad range of chemical structures with bioactivities, from antibiotics or antimycotics to immunosuppressants and anticancer compounds. Among those, compounds featuring tetronate moieties are uniquely attractive for their versatile biological activities. Most of these compounds are produced by bacteria from the phylum *Actinobacteria* and are built of a linear fatty acid or polyketide chain with a characteristic tetronic acid 4-hydroxy-2(5H)-furanone ring system.

Within the growing family of tetronates, compounds are classified taking into account the linearity or macrocyclization of the carbon backbone and the size of the central ring system (Vieweg *et al.*, 2014). Spirotetronates are tetronate compounds in which two rings are linked to each other by a spiroatom, include, amongst many others, the abyssomicins, chlorothricins, tetrocarcins, lobophorines and quartromicins. This class of tetronates shares important biosynthetic and structural features: a conjugated pair of carbon–carbon double bonds at the end of a linear polyketide chain, a characteristic exocyclic double bond on the tetronate ring system and a Diels–Alder reaction that forms the cyclohexene moiety and an additional macrocycle (Vieweg *et al.*, 2014; Weixin *et al.*, 2013).

The abyssomicin is an actively growing family of small spirotetronate natural products with a polyketide backbone and a C_11_ central ring system that has been widely studied due to the unique structural features and bioactivities that some of its members exhibit. Abyssomicin biosynthesis occurs in a variety of hosts isolated from different ecosystems. The first abyssomicins (B-D) were discovered in 2004 during the screening of 930 actinomycetes extracts in a successful attempt to find antibacterial compounds targeting folate biosynthesis. Those abyssomicins were fermentation products of the marine actinomycete *Verrucosispora maris* AB-18-032^T^, later reclassified as *Micromonospora maris* AB-18-032^T^ (Nouioui *et al.*, 2018), isolated from sediments of the Sea of Japan (Riedlinger *et al.*, 2004). Years later, other research groups found new abyssomicins produced by soil isolates of *Streptomyces* sp. HKI0381, *Streptomyces* sp. CHI39 and *Streptomyces* sp. Ank 210, in Senegal, Mexico and Germany, respectively (Abdalla *et al.*, 2011; Igarashi *et al.*, 2010; Niu *et al.*, 2007). After that, the production of abyssomicins was again reported in marine isolates: *Verrucosispora* sp. MS100128 (Wang *et al.*, 2013), *Streptomyces* sp. RLUS1487 (León *et al.*, 2015) and *Verrucosispora* sp. MS100047 (Huang *et al.*, 2016). Finally, the last abyssomicins discovered were synthesised by the soil *Streptomyces* sp. LC-6-2 (Wang *et al.*, 2017a) and the marine *S. koyangensis* SCSIO 5802 (Huang *et al.*, 2018; Song *et al.*, 2017) (Table S1).

The narrow number of bacterial strains identified so far as abyssomycin producers is in contrast with the structural diversity of these molecules. In fact, this family has as many as 37 members classified to date as type I or type II abyssomicins, depending on their structure, where the type I family includes abyssomicins B–E, G, H, J–L and atrop-abyssomicin C, and type II abyssomicins are the enantiomeric counterparts of the type I compounds (Sadaka *et al.*, 2018).

Previous works also elucidated the complete abyssomicin BGC present in the genome of *M. maris* AB-18-032 and proposed a model for the biosynthesis of atrop-abyssomicin C, the atropisomer of abyssomicin C and main product synthesized by *M. maris* AB-18-032 (Gottardi *et al.*, 2011; Keller *et al.*, 2007; Nicolaou & Harrison, 2006, 2007). This abyssomicin biosynthetic gene cluster (*aby*) comprises 25 genes, distributed along 56 kb in *M. maris* AB-18-032 genome. The cluster consists of several transcriptional regulators (*abyR, abyH, abyI* and *abyC*), an ABC exporter system (*abyF1-F4*), a drug resistance transporter (*abyD*), a cytochrome P450 system (*abyV, abyW* and *abyZ*), a cytochrome P450 gene (*abyX*), a monooxygenase (*abyE*), a type II thioesterase (*abyT*), a Diels-Alderase (*abyU*), the PKS I genes (*abyB1, abyB2* and *abyB3*) and five genes (*abyA1-A5*) involved in the assembly of the tetronic acid moiety (Figure 1A and Table S10) (Gottardi *et al*., 2011). The partially sequenced cluster of the isolate *Verrucosispora* sp. MS100047 is 99% similar to *aby* BGC (Figure 1B and Table S13).

**Figure 1.**
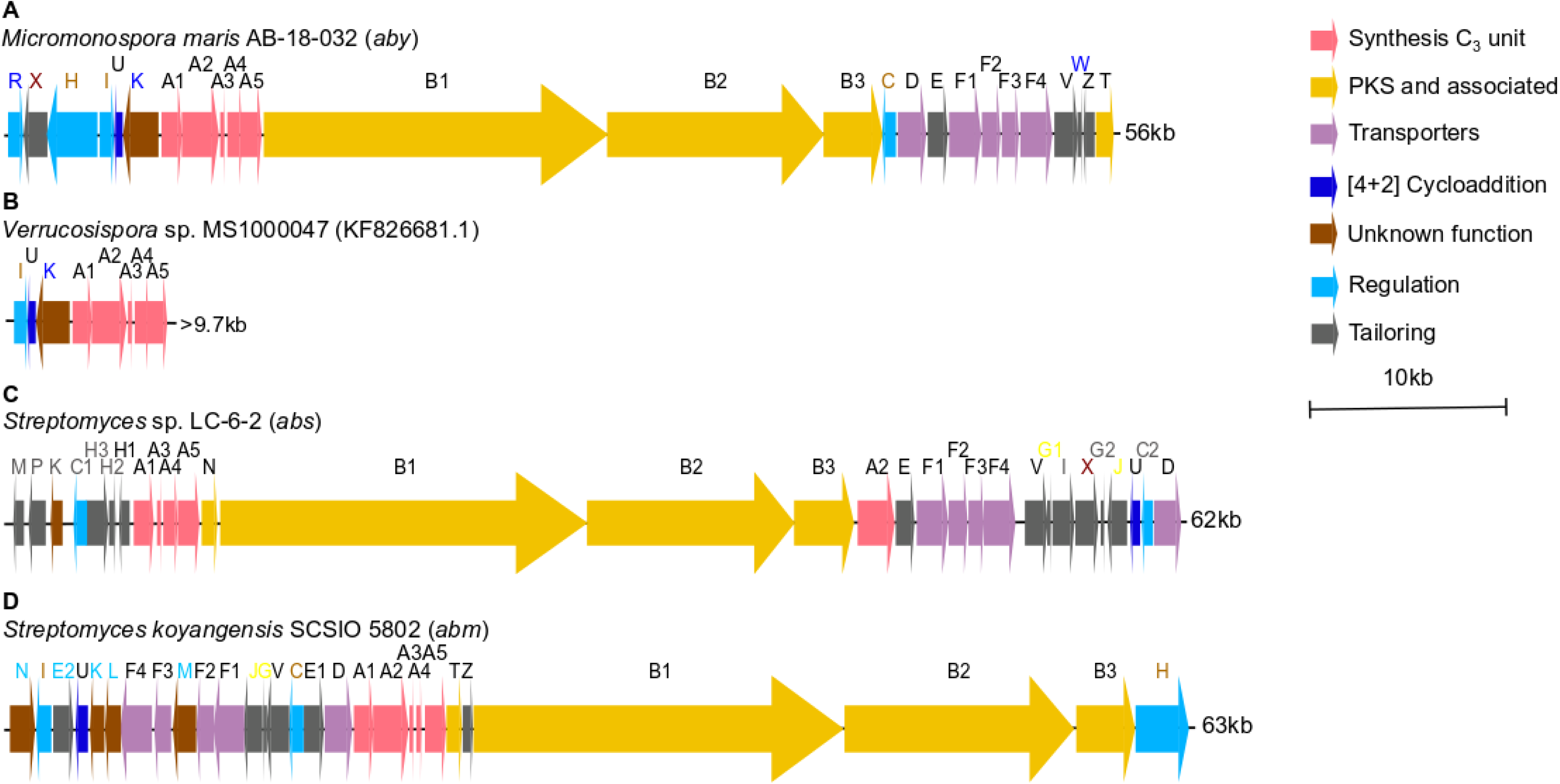
A) Abyssomicin biosynthetic gene cluster (*aby*) of *M. maris* AB-18-032. B) Partial abyssomicin biosynthetic gene cluster of *Verrucosispora* sp. MS1000047. C) Abyssomicin biosynthetic gene cluster (*abs*) of *Streptomyces* sp. LC-6-2. D) Neoabyssomicin/abyssomicin biosynthetic gene cluster (*abm*) of *S. koyangensis* SCSIO 5802. Gene names in black are common to *aby*, *abs* and *abm* BGCs. Blue font represents genes present only in *M. maris* AB-18-032, grey font represents genes present only in *Streptomyces* sp. LC-6-2 and light blue font represent genes unique to *S. koyangensis* SCSIO 5802. In maroon font appear those genes that appear both in *aby* and *abs* BGCs, in light brown those genes that appear both in *aby* and *abm* BGCs and in yellow those genes that appear both in *abs* and *abm* BGCs.

After that, the discovery of abyssomicins M-X as fermentation products of *Streptomyces* sp. LC-6-2 led to the description of a new abyssomicin BGC (*abs*). This cluster presents homologs to most of the genes within *aby* BGC (Table S12) but displays also a new array of tailoring genes. It consists of 30 genes disposed along 62 kb that include a dehydrogenase (*absM*), a hydrolase (*absP*), oxidoreductases (*absH3* and *absH2*), the NADPH-dependent flavin reductase *absH1* homolog of *abyZ*, the five genes involved in the assembly of the tetronic acid moiety (*absA1-A5*), a thioesterase (*absN*) homolg of *abyT*, the PKS genes (*absB1, absB2* and *absB3*), an ABC exporter system (*absF1-F4*), a drug resistance transporter (*absD*), cytochrome P450 gene (*absX*), a monooxygenase (*absE*) and a Diels-Alderase (*absU*). Moreover, this cluster has two unique regulators (*absC1* and *absC2*) and a set of four new tailoring genes (*absG1, absG2, absI* and *absJ*) (Wang *et al.*, 2017b) (Figure 1C).

Finally, a third cluster responsible for neoabyssomicin/abyssomicin biosynthesis (*abm*) was identified in *S. koyangensis* SCSIO 5802. Composed of 28 genes distributed along 63 kb, the *abm* cluster shares with *aby* and *abs* clusters the three PKS genes (*abmB1*–*B3*), the five consecutive genes that code for tetronate biosynthesis (*abmA1*–*A5*), the Diels–Alderase (*abmU*), the oxygenase (*abmV*), as well as the transporters (*abmD*, *abmF1*–*F4*) and regulators (*abmI* and *abmH*). Moreover, there are five genes (a*bmK*, *abmL*, *abmM*, *abmN* and *abmE2*) with no apparent homologous counterparts in the *aby* cluster and two more (*abmJ* and *abmG)* that appear to be in *abs* BGC but not in *aby* BGC (Figure 1D and Table S11)(Tu et al., 2018).

The environmental diversity of the abyssomicin-producing isolates suggests that abyssomicin biosynthesis could be, *a priori*, ubiquitously distributed in nature, and bioprospecting could focus on those environments heavily colonised by *Actinobacteria* of the genus *Micromonospora* and *Streptomyces*. Moreover, very little is known in the field of natural products about the driving forces behind the transmissibility and evolution of BGCs. In the abyssomicin family, the chemical diversity found resulted from having different abyssomicin BGCs distributed across several *Actinobacteria* with different genomic contexts and biosynthetic capabilities, in addition to the natural amenability of these clusters as the enzymes involved in the synthesis of the tetronate (AbyA1-A5) and the spiro-tetronate-forming Diels-Alderase (AbyU) are capable of accepting structurally diverse substrates. This, together with the PKS module composition explains the chemical diversity found so far within this family of natural products and opens up the possibility of expanding this diversity if other different abyssomicin BGCs were found in nature.

In the present work, in order to investigate the environmental colonization of abyssomicin-producing bacteria as well as the structural diversity of abyssomicin BGCs, we have systematically explored the distribution of abyssomicin BGC and its evolution through the analysis of publicly available genomic and metagenomic data, targeting the Diels-Alderase (AbyU/AbsU/AbmU) that catalyses the intramolecular [4 + 2] cycloaddition reaction of the linear abyssomicin polyketide precursor.

## Materials and methods

### Diels-Alderase directed metagenome mining

A total of 3027 metagenomes available in the JGI metagenomes database (IGM; https://img.jgi.doe.gov/; accessed February-April 2019) were analysed for AbyU/AbsU/AbmU homologs presence using the site option to carry out BLASTp (default parameters). The sequences of AbyU (*Micromonospora maris* AB-18-032), AbsU (*Streptomyces* sp. LC-6-2) and AbmU (*Streptomyces koyangensis* SCSIO 5802) used as query can be found in Table S3. Habitats frequently populated by *Micromonospora* and *Streptomyces* species were selected, primarily from soil and aquatic environments but also from other less common environments, including fresh-water, artificial and host-associated environments. The detailed classification of the metagenomic samples from aquatic, terrestrial, engineered and host-associated environments mined for AbyU, AbsU and AbmU can be found in Tables S5-S8. For complete details on the metagenomes analysed and the Diels-Alderase positive metagenomes please refer to Supplementary file 1.

In order to investigate possible taxonomic biases within the bacterial communities of Diels-Alderase positive and negative metagenomes, the taxonomic profiles of ten Diels-Alderase positive metagenomes were analysed compared against ten marine and ten terrestrial Diels-Alderase negative metagenomes, randomly selected from the 3027 pool.

### Diels-Alderase directed genome mining and identification of putative abyssomicin BGCs

BLASTp of AbyU, AbsU and AbmU were carried out against the non-redundant protein sequences database (NCBI; accessed April 2019). The Diels-Alderase containing genomes were then submitted to antiSMASH (Blin *et al.*, 2019) (accessed April 2019; default parameters used) for BGC mining. The location of the Diels-Alderase homolog within the genome was used to verify BGC presence in antiSMASH. When a BGC was found by antiSMASH in the desired location, ORF, protein size and proposed annotation were collated and BLASTp of every protein was carried out against the non-redundant protein sequences database (NCBI) to obtain the closest homolog (Tables S14-S84). BLASTp was used to verify/redefine the BGCs limits stablished by antiSMASH. In cases where antiSMASH did not identify any BGC, reconstruction of the Diels-Alderase homolog nearby genomic region was done manually from the corresponding genome in NCBI. All recovered BGCs were classified based on their completeness and novelty (Table 1).

**Table 1.** Classification of the recovered BGCs found through Diels-Alderase directed genome mining.

| **BGC** | **The Diels-Alderase homolog is** |
| --- | --- |
| Abyssomicin, total | Part of an abyssomicin BGC and it is possible to recover the sequence and structure of the entire BGC. |
| Abyssomicin, partial | Part of an abyssomicin BGC that is likely to be complete but due to the sequencing technology used there are some incomplete genes, frame shifts, gaps or the cluster is on a contig edge. |
| Potential abyssomicin, total | Part of a BGC whose product may potentially be an abyssomicin according to antiSMASH and it is possible to recover the sequence and structure of the entire BGC. |
| Potential abyssomicin, partial | Part of a BGC whose product may potentially be an abyssomicin according to antiSMASH but there are some genes missing or incomplete, frame shifts, gaps or the cluster is on a contig edge. |
| Potential BGC, total | Surrounded by genes that could form a BGC altogether, but it is unclear which could be its product. |
| Potential BGC, partial | Surrounded by genes that could form a BGC altogether, but it is unclear which could be its product and there were some incomplete genes, frame shifts, gaps or the cluster was on a contig edge. |
| Not a BGC | Not likely to be part of any BGC. |
| Not enough data | In a contig whose length makes it not possible to gain any knowledge. |
| Quartromycin, total | In a quartromycin BGC and the sequence of the cluster is complete. |
| Quartromycin, partial | In a quartromycin BGC but the sequence of the corresponding PKS is incomplete. |
| Potential tetronomycin, total | Part of a potential tetronomycin BGC. |
| Potential chlorothricin, partial | In a chlorothricin BGC but the sequence of the corresponding PKS is incomplete. |

### Evolutionary analysis

All the proteins identified through genome mining that produced significative alignments (E-value < 10^−6^) against AbyU, AbsU and AbmU were aligned and the Neighbor-Joining algorithm was used to create a phylogenetic tree using MAFFT (https://mafft.cbrc.jp/alignment/server/phylogeny.html; accessed May 2019) (Katoh *et al.*, 2002). The RefSeq annotated genomes of the microorganisms harbouring those proteins were used to create a phylogenomic tree using UBCG (default parameters) (Na *et al.*, 2018). The produced phylogenies were visualised and annotated with iTOL (Letunic & Bork, 2019).

A manual synteny analysis was carried out for all the newly discovered abyssomicin and potential abyssomicin BGCs (both total and partial), which were classified accordingly as described below (Table 2). The presence of mobile elements within the Diels-Alderase positive mined genomes was studied using Island Viewer 4 (Bertelli *et al.*, 2017).

**Table 2.**
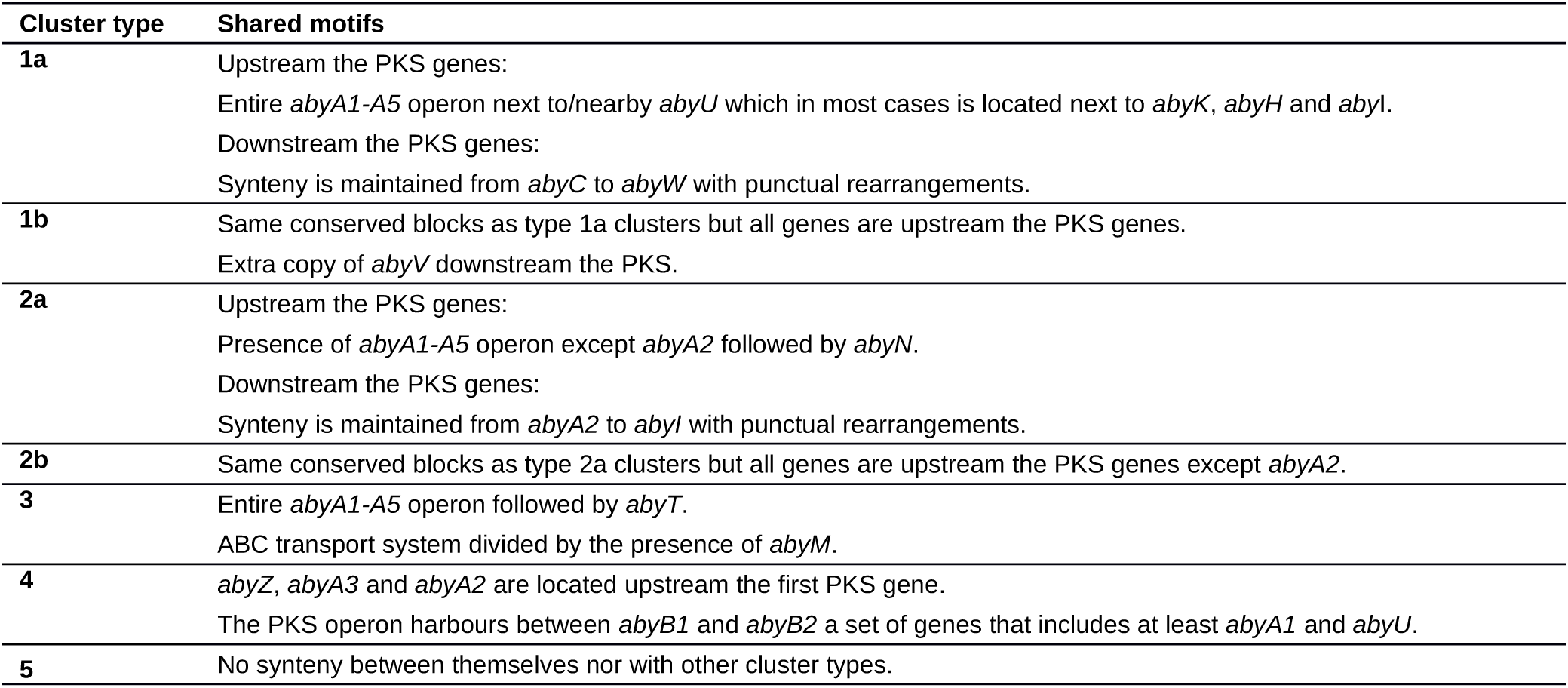
Features shared by the potential abyssomicin BGCs described in this study.

| Cluster type | Shared motifs |
| --- | --- |
| **1a** | Upstream the PKS genes:<br>Entire *abyA1-A5* operon next to/nearby *abyU* which in most cases is located next to *abyK*, *abyH* and *abyI*.<br>Downstream the PKS genes:<br>Synteny is maintained from *abyC* to *abyW* with punctual rearrangements. |
| **1b** | Same conserved blocks as type 1a clusters but all genes are upstream the PKS genes.<br>Extra copy of *abyV* downstream the PKS. |
| **2a** | Upstream the PKS genes:<br>Presence of *abyA1-A5* operon except *abyA2* followed by *abyN*.<br>Downstream the PKS genes:<br>Synteny is maintained from *abyA2* to *abyI* with punctual rearrangements. |
| **2b** | Same conserved blocks as type 2a clusters but all genes are upstream the PKS genes except *abyA2*. |
| **3** | Entire *abyA1-A5* operon followed by *abyT*.<br>ABC transport system divided by the presence of *abyM*. |
| **4** | *abyZ*, *abyA3* and *abyA2* are located upstream the first PKS gene.<br>The PKS operon harbours between *abyB1* and *abyB2* a set of genes that includes at least *abyA1* and *abyU*. |
| **5** | No synteny between themselves nor with other cluster types. |

## Results

### Habitat distribution of the Diels-Alderase positive metagenomes

In order to study the habitat distribution of the bacteria harbouring an abyssomicin BGC, and considering that the Diels-Alderase AbyU could be used as an abyssomicin-biosynthesis specific marker, we mined 3027 publicly available metagenomes for the presence of AbyU and its already known homologs AbsU and AbmU (Table S3-S4). 27% of the analysed metagenomes had aquatic origin, 31% belonged to soil samples, 22% were plant-associated and the remaining 20% covered human-built environments and different host-associated microbiomes (Figure S1). Our results showed that the three Diels-Alderase homologs share a similar habitat distribution, 31% of the AbyU positive metagenomes were from soil, 68% were plant-associated and 1% Arthropoda-associated (Figure 2B); 55% of the AbsU-positive had soil origin and 45% were plant-associated (Figure 2C) and AbmU displayed a similar distribution to AbyU with the only difference of being additionally present in an artificial bioreactor environment (Figure 2D). None of the AbyU, AbsU or AbmU positive metagenomes had aquatic origin.

**Figure 2.**
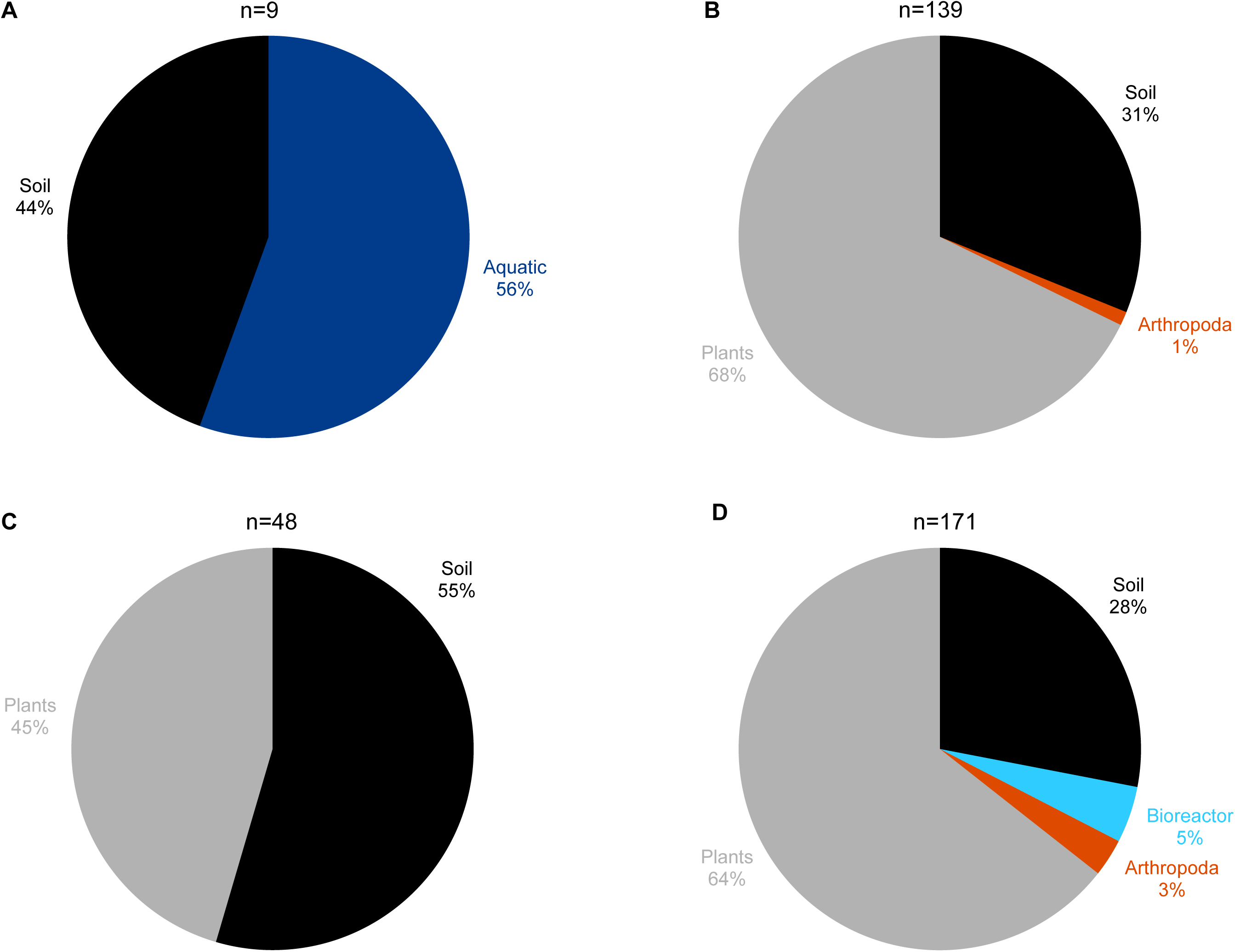
Habitat distribution of **A)** The abyssomicin producing bacteria isolated and reported in the literature until this manuscript was written. **B)** Metagenomes containing AbyU homologs. **C)** Metagenomes containing AbsU homologs. **D)** Metagenomes containing AbmU homologs.

Following these results, ten marine, ten soil and ten Diels-Alderase positive metagenomes (six from soil, three plant-associated, one Arthropoda-associated) were randomly selected from the 3027 pool to confirm that the absence of Diels-Alderase homologs in the aquatic metagenomes was not due to significative differences in *Actinobacteria* abundance nor in sequencing depth. Metagenome size and taxonomic composition were analysed in each of the 30 metagenomes. The sequencing depth of aquatic metagenomes was similar to other habitats and therefore, was discarded as a plausible explanation (Figure S2). The taxonomic assemblage of the 30 metagenomes was similar with a few exceptions. Specifically, *Proteobacteria* was the dominant phylum independently of the type of habitat and *Actinobacteria* was generally found more abundant in soil environments in comparison to aquatic habitats. Moreover, three of the Diels-Alderase positive metagenomes (two from soil and one Arthropoda-associated) displayed an abundance of *Actinobacteria* similar to the aquatic metagenomes (Figure S2). Within the class *Actinobacteria*, all 30 metagenomes displayed a similar taxonomic profile independently of the habitat, being *Streptomycetales* more represented only in three (two from soil and one plant-associated) of the Diels-Alderase positive metagenomes, *Micrococcales* more abundant in one aquatic and one soil metagenome and *Corynebacteriales* in one aquatic metagenome (Figure S3). The taxonomic composition of the other 24 metagenomes was heterogeneous, where at least 5 to 10 taxa were numerically dominant, including the genus *Micromonospora* (Figure S3).

### Diels-Alderase directed genome mining and diversity of abyssomicin BGCs

In order to gain a better overview over how abyssomicin-producing bacteria are environmentally distributed and the structural diversity of abyssomicin BGCs in nature, both partial and complete genomes available in public databases were mined. In a BLASTp of AbyU, AbsU and AbmU against the RefSeq NR database, 74 Diels-Alderase homologs from 66 different genomes were identified (Table S9).

All the 66 Diels-Alderase positive genomes belonged to culturable strains. The habitat distribution of these isolates was, overall, similar to that found by metagenome mining (Figure S4). Specifically, about one third of the strains were recovered from soil, one third from plant-associated environments, and the remaining were recovered from marine environments, mammals, annelids and lichens.

The bacterial genomes were analysed in order to locate those Diels-Alderase homologs and study whether they were part of a potential abyssomicin BGC. This way, it was possible to identify and annotate five total and 12 partial new abyssomicin BGCs and 23 new potential abyssomicin BGCs never described until now and with similar but not identical architectures to *aby*, *abs* and *abm* clusters (Figure 3). 11 of the Diels-Alderase homologs could be located in potential BGCs, three more were found in genomic regions apparently unrelated to any BGC and 11 were located in short contigs from which it was impossible to infer any information. Finally, two Diels-Alderase homologs were found in two different quartromicin BGCs and another two in potential tetronomycin and chlorothricin BGCs.

**Figure 3.**
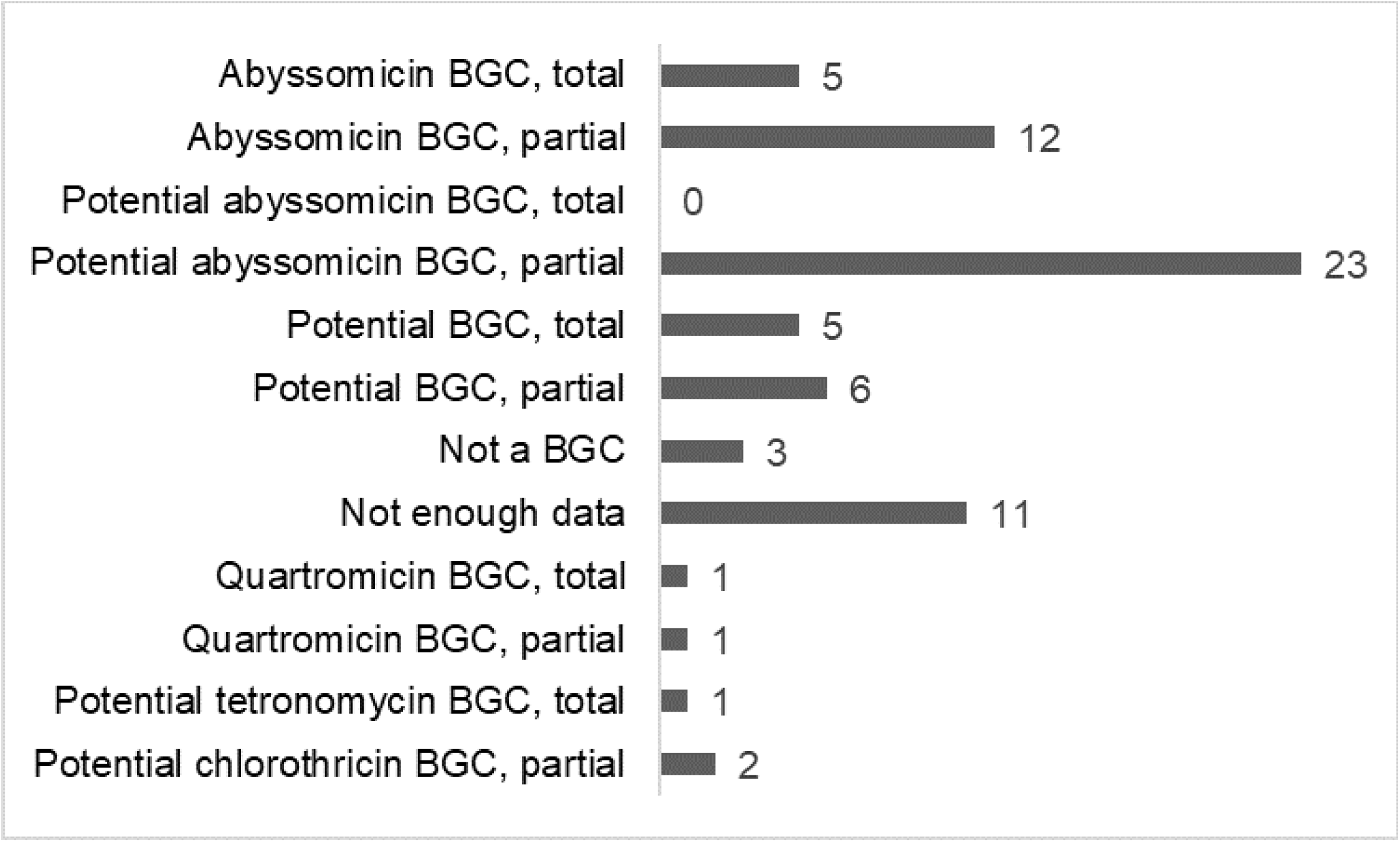
Recovered BGCs found through Diels-Alderase directed genome mining.

From the newly identified Diels-Alderase homologs it was possible to recover 40 total or partial new clusters potentially involved in the biosynthesis of abyssomicins (Figure S7-S10; Table S14-S84). These clusters were further classified according to their synteny in order to analyse their structural diversity. The analysis was carried out manually, as the modular nature of BGCs made the application of general synteny analysis tools impossible. Considering the diversity of biosynthetic genes and their disposition, abyssomicin and potential abyssomicin BGCs were classified into seven cluster types (Table 2). There were four genomes containing type 1a clusters and ten genomes displaying type 1b clusters from the genera *Micromonospora*, *Actinokineospora*, *Frankia*, *Herbidospora* and *Streptomyces* (Figure S7). There were seven clusters classified as type 2a and two clusters classified as type 2b. In this case, type 2a clusters were found in *Streptomyces*, *Actinokineospora* and *Micromonospora* and type 2b only in *Frankia* (Figure S8). Five clusters were classified as type 3, all belonging to *Streptomyces* and three clusters were type 4 clusters found in *Streptomyces* and *Streptacidiphilus* (Figure S9). Finally, there were 13 clusters that did not present enough similarity to any of the cluster types described above. These cluster were found in *Frankia*, *Actinokineospora*, *Lentzea*, *Kutzneria*, *Micromonospora*, *Streptomyces*, *Saccharothrix* and *Actinocrispum* and did not share any outstanding synteny pattern amongst themselves (Figure S10) neither with the five potential tetronomycin, chlorothricin, or quartromycin BGCs that were also found from the Diels-Alderase directed genome mining (Figure S11). The genomes that harboured a Diels-Alderase that was not part of an abyssomicin or potential abyssomicin BGC were not considered for this classification.

### Evolutionary history of abyssomicin BGCs

Most of the Diels-Alderase positive bacteria were taxonomically identified as belonging to the phylum *Actinobacteria* and most of them to the genus *Streptomyces* (37 isolates), followed by seven *Frankia*, three *Herbipospora*, three *Actinomadura* and three *Micromonospora* strains (Figure 4). As was expected, all the genera formed monophyletic clusters, corroborating their correct taxonomic assignment. The abyssomicin BGCs were only identified in several species of some actinobacterial genera but not in all, suggesting that the abyssomicin BGCs may be acquired through horizontal gene transfer (HGT) events, duplications or rearrangements. This hypothesis was reinforced by the fact that the phylogenetic history of the Diels-Alderase (Figure 5) does not follow the same evolutionary history as of the species tree (Figure 4).

**Figure 4.**
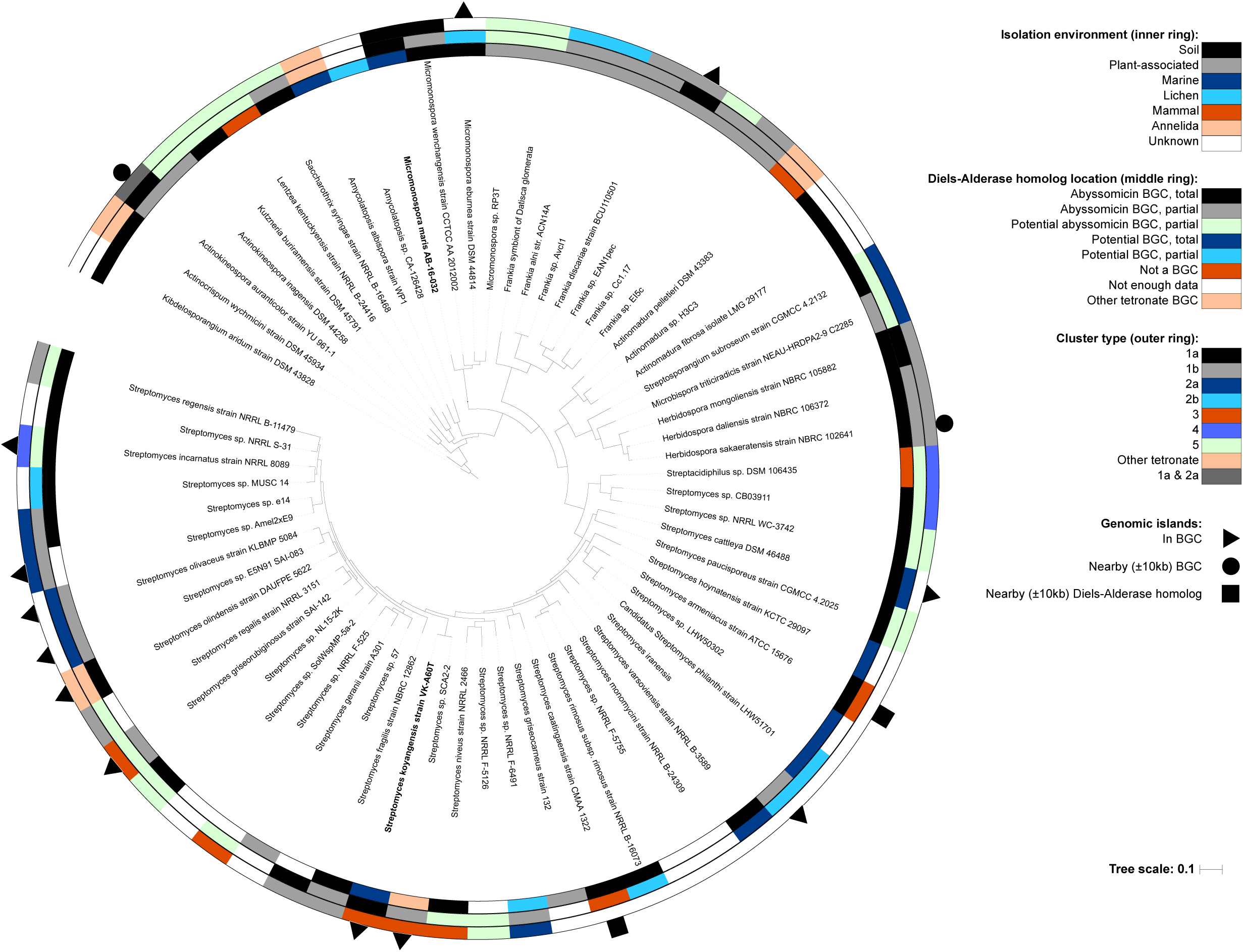
Phylogenomic tree of bacterial genomes containing a Diels-Alderase homolog. The inner ring represents the environment where each strain was isolated, the middle ring depicts the location of the Diels-Alderase homolog and the outer ring shows the cluster type for those isolates found to have abyssomicin and potential abyssomicin BGCs both total and partial. Outer symbols indicate presence of genomic island inside the abyssomicin or potential abyssomicin BGC, nearby it (±10 kb upstream or downstream BGC) or nearby the Diels-Alderase (±10 kb upstream or downstream) when the isolate did not present an abyssomicin BGC.

**Figure 5.**
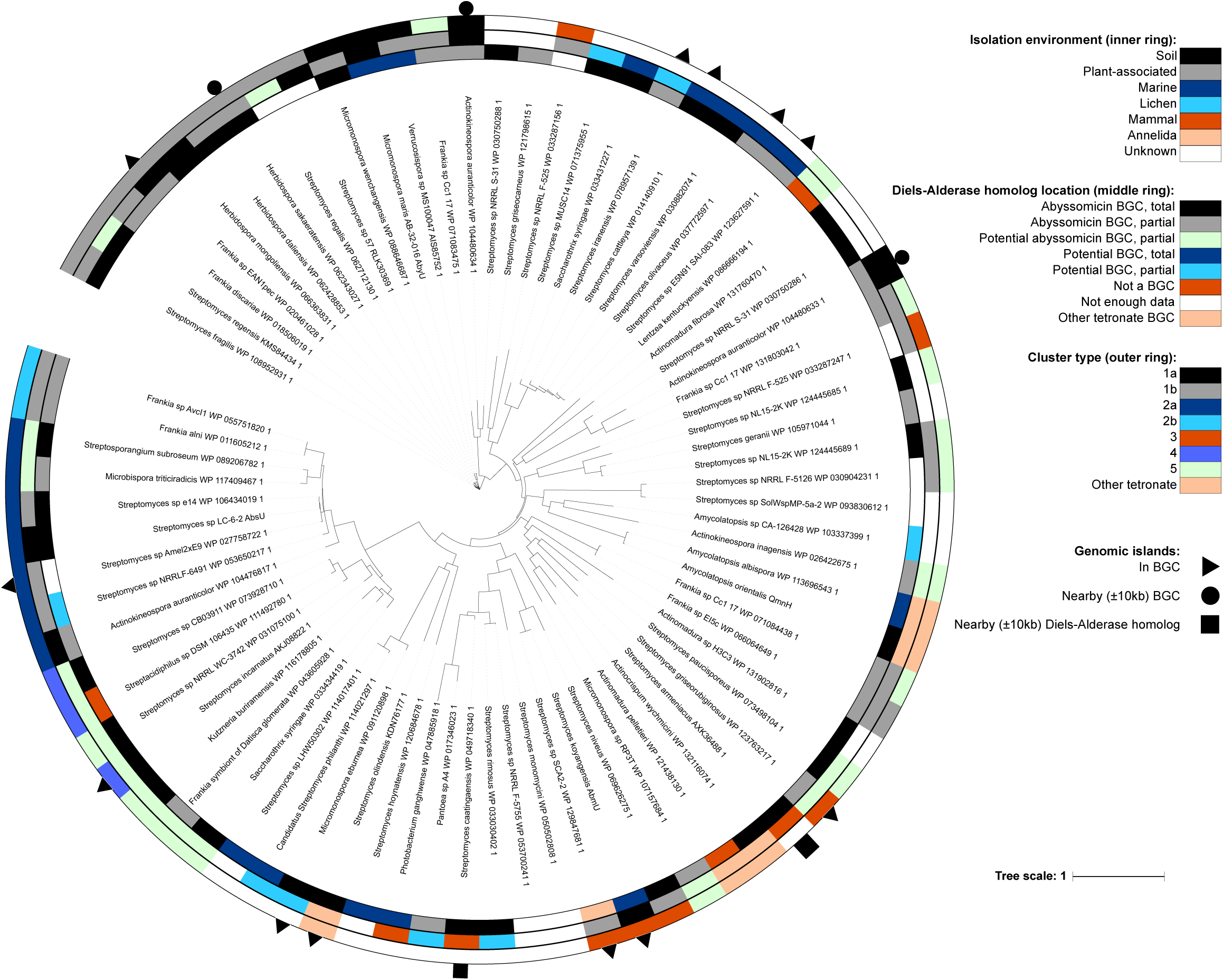
Phylogenetic tree the Diels-Alderase homologs. The inner ring represents the environment where each strain was isolated, the middle ring depicts the location of the Diels-Alderase homolog and the outer ring shows the cluster type for those isolates found to have abyssomicin and potential abyssomicin BGCs both total and partial. Outer symbols indicate presence of genomic island inside the abyssomicin or potential abyssomicin BGC, nearby it (±10 kb upstream or downstream BGC) or nearby the Diels-Alderase (±10 kb upstream or downstream) when the isolate did not present an abyssomicin BGC.

Interestingly, abyssomicin BGCs are usually associated with genomic islands (GI; Figure S7-S10; Table S85-S100) which may allow its HGT among taxa. Specifically, GI were identified in the abyssomicin BGC of some *Streptomyces*, *Frankia*, *Herbidospora*, *Micromonospora* and *Actinokineospora*, nearby it (±10 kb upstream or downstream BGC) or nearby the Diels-Alderase (±10 kb upstream or downstream) (Figure 5). Albeit that the vast majority this HGT events take place among members of the phylum *Actinobacteria*, two *Proteobacteria*, namely *Pantoea* sp. A4 and *Photobacterium ganghwense* JCM 12487, harboured a Diels-Alderase. The Diels-Alderase genes of both *Proteobacteria* strains were very phylogenetically related among them and in turn related to other *Streptomyces* strains (Figure 5). The closest neighbour to both proteobacterias was *S. caatingaesis* WP_049718340. Thereby, these intra- and inter-phyla HGT events may be explained by the presence of mobile elements such as transposases and integrases flanking or within the BGCs (Table S85-S100). Moreover, several abyssomicin paralogs were found in the mined genomes of *A. auranticolor* YU 961-1, *Frankia* sp. Cc1.17, *Streptomyces* sp. NL15-2K, *Streptomyces* sp. NRRL F-525, *Streptomyces* sp. NRRL S-31 and *S. syringae* NRRL B-16468 (Figure 5).

On the other hand, evolutive pressure has shaped the abyssomicin BGCs, widening the functional and structural diversity of this secondary metabolite. In fact, the presence of tailoring genes is variable among species as well as the Diels-Alderase gene location within the BGCs (Figure S7-S10; Table S14-S84). However, the synteny of abyssomicin BGCs lacks phylogenetic signal and hence the abyssomicin BGC classification that we propose in the present study could not be used to trace its evolutive history.

## Discussion

### Habitat distribution of the Diels-Alderase hosts discovered through metagenome and genome mining

Until today, only nine bacterial strains have been reported to produce abyssomicins (Table S1). From these strains, 37 abyssomicins with differences at structural and bioactivity levels have been characterized (Sadaka *et al.*, 2018). With the aim of studying the distribution of those microorganisms capable of producing new abyssomicin molecules, we have analysed *in silico* an extensive diversity of metagenomes and genomes. AbyU is the natural Diels-Alderase present in abyssomicin BGC that catalyses the formation of the heterobicyclic ring system that characterises this family of natural products. Very few enzymes in nature can catalyse this reaction and despite being capable of accepting structurally diverse substrates, sequence conservation with the closest known spirotetronate cyclases is minimal (Byrne *et al.*, 2016). We selected this enzyme to lead the mining as it is essential in abyssomicin biosynthesis.

Here we mined 3027 metagenomes for the presence of AbyU, AbsU and AbmU, and our results showed that Diels-Alderase positive microorganisms have a strikingly diverse environmental distribution, being mainly present in soil and plant-associated microbiomes but totally absent in aquatic habitats (Figure S1). Since the few isolates reported in the literature to produce abyssomicins were equally distributed between aquatic and soil environments (Figure 2A; Table S1), our results were totally unexpected. In order to further study this, we compared the relative abundance of Diels-Alderase positive bacteria with regard to the sequencing depth of each study and the taxonomic composition of the communities, namely the different abundance of *Actinobacteria* (Figure S2-S3). However, none of both factors showed a direct correlation with the absence of Diels-Alderase positive bacteria in aquatic habitats. A possible explanation for the absence of Diels-Alderase positive bacteria in aquatic habitats could be related to the ecological role of the abyssomicins. Specifically, we hypothesize that abyssomicins could have key ecological roles in benthonic regions, since all the abyssomicin producing strains isolated from aquatic environments so far come from marine sediments (Table S2). Moreover, it is tempting to hypothesize that abyssomicin-producing bacteria may be involved in symbioses with higher organisms, which has been seen before for other different antibiotic-producing strains that play an important role as defensive symbionts both in marine and terrestrial ecosystems (Adnani *et al.*, 2017; Gunatilaka, 2006; Seipke *et al.*, 2012). The abyssomicins could also act as signal molecule in plant-bacteria communication or as precursors involved in plant growth and development, as reported before in the *Frankia* and *Micromonospora* genera through, for example, the formation of nitrogen fixing actinonodules (Sellstedt & Richau, 2013; Trujillo et al., 2010). Further investigations will be needed in order to unravel the biased habitat distribution of Diels-Alderase positive bacteria.

In addition, we identified 74 Diels-Alderase homologs present in 66 different genomes (Table S9) from which it was possible to identify and annotate five total and 12 partial new abyssomicin BGCs and 23 new potential abyssomicin BGCs. Indeed, all these 40 abyssomicin and potential abysomicin producers are culturable strains whose habitat distribution follows the same patterns found through the metagenome mining as none of them was recovered from aquatic samples (Figure S4-S5). In our case, 66% of the Diels-Alderase positive genomes displayed an abyssomicin or potential abyssomicin BGC. In the remaining genomes in which the Diels-Alderase was not located in any BGC, we could not predict its metabolic function. Previous studies reported other Diels-Alderases involved in the synthesis of other natural products, with the exception of riboflavin synthases that are involved in primary metabolism (Lichman *et al.*, 2019).

Therefore, based on the genome- and metagenome mining, we can conclude that the potential abyssomicin producers have a cosmopolitan distribution albeit their presence in aquatic habitat is limited. This strongly suggests that abyssomicin bioprospecting efforts should not be focused on aquatic environments but rather on soil and plant-associated ones. Also, two Diels-Alderase homologs were found in two different quartromicin BGCs and another two in potential tetronomycin and chlorothricin BGCs. The presence of those four Diels-Alderase homologs within BGCs belonging to other natural products is well justified, as quartromicin, tetronomycin and chlorothricin share the same tetronate cycloaddition as the abyssomicins (Vieweg *et al.*, 2014).

Moreover, 11 of the Diels-Alderase homologs detected in the mined genomes were in potential non-abyssomicins BGCs, three more were found in genomic regions *a priori* unrelated to any BGC and 11 appeared in short contigs from which it was impossible to infer any information. In this case, only ten of the 66 genomes analysed were completely sequenced and only seven isolates were sequenced with third generation sequencing technologies (Table S9). The identification of the Diels-Alderase homologs location within the genomes and the recovery of potential BGCs was influenced by the quality of the sequencing technology used and the assembly level achieved by each previous individual study. Some factors such as the high G+C content of actinomycete genomes affect the sequencing reactions and the assembly process (Nakamura *et al.*, 2011), however, the biggest challenge appears to recover the highly-conserved and modular sequences of polyketide synthases (PKS) characterised by displaying highly similar intragenic and intergenic tandem repeats at nucleotide level, which in many cases are longer than the read-length of the sequencing technology used (Gomez-Escribano *et al.*, 2016). Moreover, large PKS clusters can often be distributed along several contigs, and it has been demonstrated that sequencing errors can introduce false frameshifts into the large PKS sequences (Blažič et al., 2012). Finally, the presence of Diels-Alderase homologs outside abyssomicin BGCs, could be explained by the presence of transposases flanking Diels-Alderase homologs allowing their genetic recombination along the genome (Table S85-S100). Specifically, the Diels-Alderase homologs of *Streptomyces caatingaensis* CMAA 1322 and *Streptomyces armeniacus* ATCC 15676 were not part of an abyssomicin BGC but showed transposases on both sides (Figure 5; Table S89; Table S91).

### Evolutive history of abyssomicin BGC

It is well-known that *Actinobacteria* are characterised by its ability to produce a wide variety of specialised metabolites and, despite the problem of re-discovering already known molecules, bacteria from the phyla *Actinobacteria* are still one of the most prolific sources of chemical diversity (Genilloud, 2017). Abyssomicin BGCs presence is limited to the phylum *Actinobacteria*, mainly representatives of the genus *Streptomyces* and *Frankia*. The constraint of the abyssomicin BGC to some specific strains suggests that speciation was not the primary driver for dissemination of this cluster (Figure 4). Instead, HGT may have played an important role in the transmission of abyssomicin BGCs, which may have jumped among taxa through mobile elements (Hall *et al.*, 2017; Ziemert *et al.*, 2014). Indeed, many integrases and transposases were found surrounding or inside the abyssomicin BGCs (Table S85-S100).

Many BGCs in *Actinobacteria* evolve through HGT events, but only a few studies have demonstrated it (Choudoir *et al.*, 2018). For example, in a genome mining study on 75 *Salinispora* strains, 124 pathways involved in the synthesis of PKS and non-ribosomal peptide synthetase (NRPS) natural products were identified and showed that HGT events were responsible for the majority of pathways, which occurred in only one or two strains, as acquired pathways were incorporated into genomic islands (Ziemert *et al.*, 2014). In another example, the secondary metabolite clusters on the chromosome of *S. avermitilis* ATCC31267 were found to contain many transposase genes in the regions near both ends of the clusters, suggesting these transposases might have been involved in the transfer of these clusters (Omura *et al.*, 2001). Similarly, it was demonstrated that the rifamycin BGC in *Salinispora arenicola* CNS-205 had been acquired through HGT directly from *Amycolatopsis mediterranei* S699 by genomic island movement (Penn *et al.*, 2009).

Although HGT events are more frequent among phylogenetically close taxa, in this case within the phylum *Actinobacteria*, HGT events can take place among different phyla. In the present study, we could identify a HGT event of Diels-Alderases from a representative of the genus *Streptomyces* to two strains of the phylum *Proteobacteria*, namely *Pantoea* sp. A4 and *Photobacterium ganghwense* JCM 12487 (Figure 5). The transmission of functional BGCs among phyla was also reported by other authors (Zeng *et al.*, 2014). Unfortunately, neither transposases nor integrases were identified nearby the Diels-Alderases of *Pantoea* sp. A4 and *Photobacterium ganghwense* JCM 12487, which could have explained the HGT event.

The acquisition of an abyssomicin BGC by a bacterial strain could increase its evolutionary fitness and therefore enhance its competitiveness against other members of the community. In fact, the biological activity of abyssomicins includes antimicrobial activities against Gram-positive bacteria and *Mycobacteria (Freundlich et al., 2010; Riedlinger et al., 2004)*. Other biological activities discovered so far are antitumor properties, latent human immunodeficiency virus (HIV) reactivator, anti-HIV and HIV replication inducer properties (Sadaka *et al.*, 2018). The wide diversity of abyssomicin BGCs that we have found through genome mining suggests that a plethora of abyssomicin-like molecules remains unknown.

### Conclusions and future perspectives

The aim of this study was to shed some light into the structural diversity, habitat distribution and evolutionary history of abyssomicin BGC. Through metagenome and genome mining, we discovered that the habitat distribution of microorganisms harbouring a Diels-Alderase is restricted to that of the phylum *Actinobacteria*, with mainly representatives of the genus *Streptomyces* and *Frankia*, which are primarily present in soil and plant-associated environments. Surprisingly, we did not find any Diels-Alderase positive bacterium in aquatic environments although five out of nine reported abyssomicin producers were isolated from marine sediments. Therefore, all the strains that present abyssomicin BGCs have been observed to be associated to organic or inorganic solid substrates. Based on the habitat distribution of Diels-Alderase positive bacteria, we hypothesize that microorganisms producing abyssomicin-like molecules could play key ecological roles in the corresponding microbial communities.

Moreover, the vast structural diversity of abyssomicin BGCs that we have found could reflect its horizontal evolutionary history, and we predict that a plethora of abyssomicins remains unknown to date. Additionally, Diels-Alderase enzymes are of great value in synthetic chemistry, as the [4+2] cycloaddition reaction they catalyse could facilitate the development of environmentally friendly synthetic routes to a wide variety of useful compounds. Finally, the discovery of Diels-Alderase homologs, could hold great potential as part of the synthetic biology toolbox to generate libraries of novel non-natural biomolecules. Taken together, the results of the present work reveal the interest of a new bioprospecting strategy to identify biocompounds such as abyssomicins out of their currently assumed environmental distribution.

