## Supporting Information for "Out of the abyss: Genome and metagenome mining reveals unexpected environmental distribution of abyssomicins" (from another version of this preprint: version 1, 2019-10-10)


| Contents | | Page |
|---|---|---|
| **Figure S1.** | Habitat distribution of the metagenomes analysed for the presence of AbyU, AbmU and AbsU. | 2 |
| **Figure S2.** | Taxonomic profile at the phylum level of ten marine (1-10), ten terrestrial (11-20) and ten Diels-Alderase positive (21-30) metagenomes. These metagenomes were randomly selected from the 3027 metagenomes analysed. The ten Diels-Alderase positive were six terrestrial (21-26), one Arthropoda-associated (27) and three plant-associated (28-30). Only the most abundant phyla are shown. Numbers above each bar plot indicate metagenome size (Mb). | 3 |
| **Figure S3.** | Taxonomic profile at the order level of ten marine (1-10), ten terrestrial (11-20) and ten Diels-Alderase positive (21-30) metagenomes. These metagenomes were randomly selected from the 3027 metagenomes analysed. The ten Diels-Alderase positive were six terrestrial (21-26), one Arthropoda-associated (27) and three plant-associated (28-30). | 4 |
| **Figure S4.** | Habitat distribution of the Diels-Alderase positive isolates found by genome mining. | 5 |
| **Figure S5.** | Habitat distribution of the Diels-Alderase positive isolates found by genome mining to have an abyssomicin or potential abyssomicin BGC (both total and partial). | 6 |
| **Figure S6.** | Habitat distribution of the Diels-Alderase positive isolates found by genome mining that do not harbour any abyssomicin nor potential abyssomicin BGC. | 7 |
| **Figure S7.** | Abyssomicin and potential abyssomicin BGCs recovered from genome mining classified as clusters type 1a and 1b. | 8 |
| **Figure S8.** | Abyssomicin and potential abyssomicin BGCs recovered from genome mining classified as clusters type 2a and 2b. | 9 |
| **Figure S9.** | Abyssomicin and potential abyssomicin BGCs recovered from genome mining classified as clusters type 3 and 4. | 10 |
| **Figure S10.** | Abyssomicin and potential abyssomicin BGCs recovered from genome mining classified as clusters type 5. | 11 |
| **Figure S11.** | Potential tetronomycin BGC in *S. olindensis*, potential chlorothricin BGC from *A. wychmicini* and *A. pelletieri* and quartromycin BGCs from *A. albispora* and *A. orientalis*. | 12 |
| **Table S1.** | Abyssomicin producing bacteria as reported in literature. | 13 |
| **Table S2.** | Isolation details of the abyssomicin producers isolated from aquatic environments. | 14 |
| **Table S3.** | Nucleotide and protein sequences of the Diels Alderase proteins present in aby, abs and abm BGCs. | 15 |
| **Table S4.** | Identity percentage at protein (above) and nucleotide (below) level between the Diels Alderase proteins present in aby, abs and abm BGCs. | 16 |
| **Table S5.** | Classification of metagenomic samples from aquatic environments mined for AbyU, AbsU and AbmU. Numbers represent Diels-Alderase positive/mined metagenomes. | 17 |
| **Table S6.** | Classification of metagenomic samples from terrestrial environments mined for AbyU, AbsU and AbmU. Numbers represent Diels-Alderase positive/mined metagenomes. | 18 |
| **Table S7.** | Classification of metagenomic samples from engineered environments mined for AbyU, AbsU and AbmU. Numbers represent Diels-Alderase positive/mined metagenomes. | 19 |
| **Table S8.** | Classification of metagenomic samples from host-associated environments mined for AbyU, AbsU and AbmU. Numbers represent Diels-Alderase positive/mined metagenomes. | 20-21 |
| **Table S9.** | Proteins within the non-redundant sequence database (NCBI) with significative alignments to AbyU, AbmU, AbsU and VASRM7_509. Columns displaying the Diels-Alderase homologs show protein ID, alignment E-value and similarity percentage. | 22-25 |
| **Table S10.** | The abyssomicin biosynthetic gene cluster from *M. maris* AB-16-032 (modified from Gottardi et al., 2011). | 26 |
| **Table S11.** | The abyssomicin biosynthetic gene cluster from *S. koyangensis* SCSIO 5802 (modified from Song *et al*., 2017). | 27-28 |
| **Table S12.** | The abyssomicin biosynthetic gene cluster from *Streptomyces* sp. LC-6-2 (adapted from Wang et al., 2017). | 29-30 |
| **Table S13.** | The abyssomicin biosynthetic gene cluster from *Verrucosispora* sp. MS100047 (KF826681.1). | 31 |
| **Table S14-S84.** | Recovered BGCs and ORFs surrounding AbyU homologs. | 32-140 |
| **Table S85-S100.** | Predicted genomic islands. | 141-171 |
| **References** | | 172 |

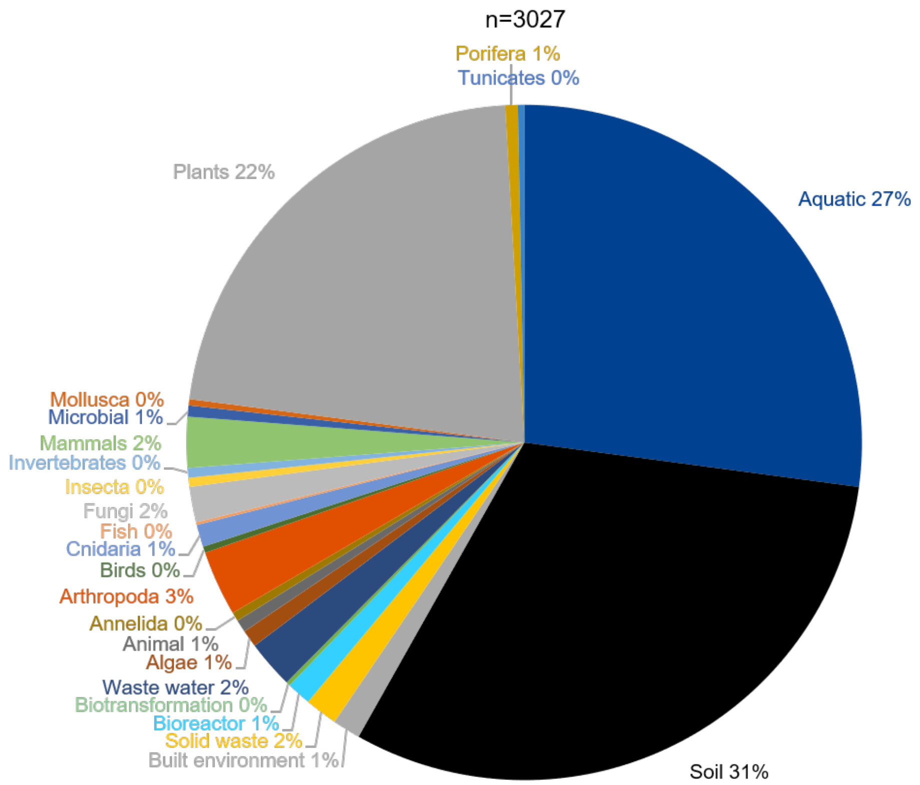


**Figure S1.** Habitat distribution of the metagenomes analysed for the presence of AbyU, AbmU and AbsU.

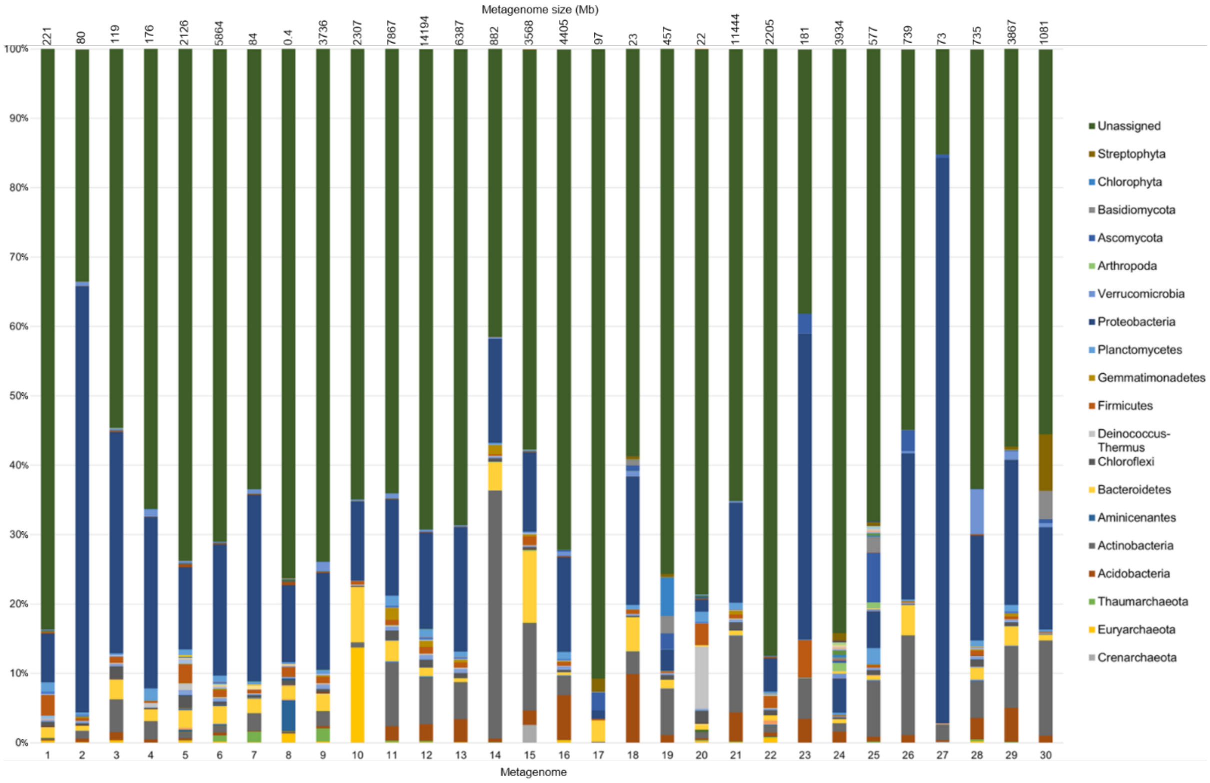


**Figure S2.** Taxonomic profile at the phylum level of ten marine (1-10), ten terrestrial (11-20) and ten Diels-Alderase positive (21-30) metagenomes. These metagenomes were randomly selected from the 3027 metagenomes analysed. The ten Diels-Alderase positive were six terrestrial (21-26), one Arthropoda-associated (27) and three plant-associated (28-30). Only the most abundant phyla are shown. Numbers above each bar plot indicate metagenome size (Mb).

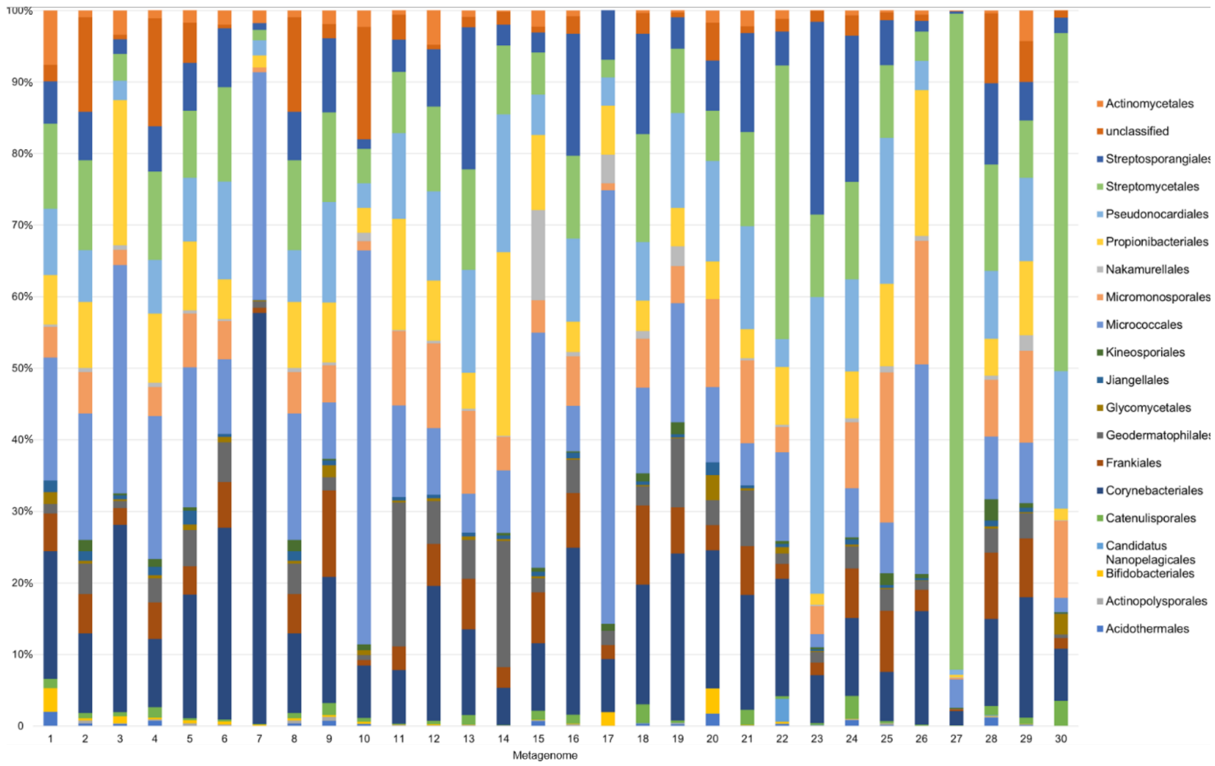


**Figure S3.** Taxonomic profile at the order level of ten marine (1-10), ten terrestrial (11-20) and ten Diels-Alderase positive (21-30) metagenomes. These metagenomes were randomly selected from the 3027 metagenomes analysed. The ten Diels-Alderase positive were six terrestrial (21-26), one Arthropoda-associated (27) and three plant-associated (28-30).

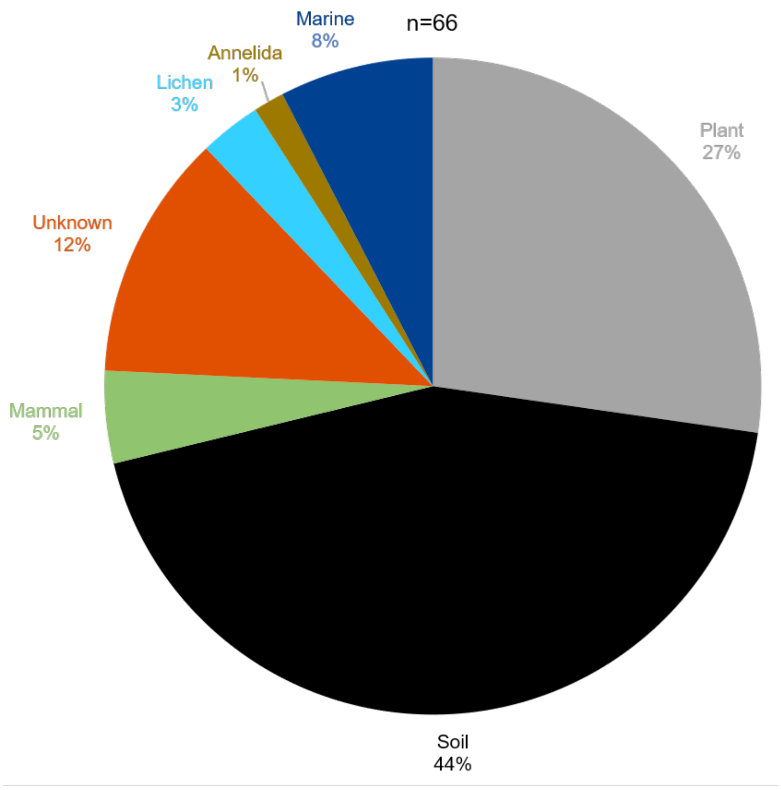


**Figure S4.** Habitat distribution of the Diels-Alderase positive isolates found by genome mining.

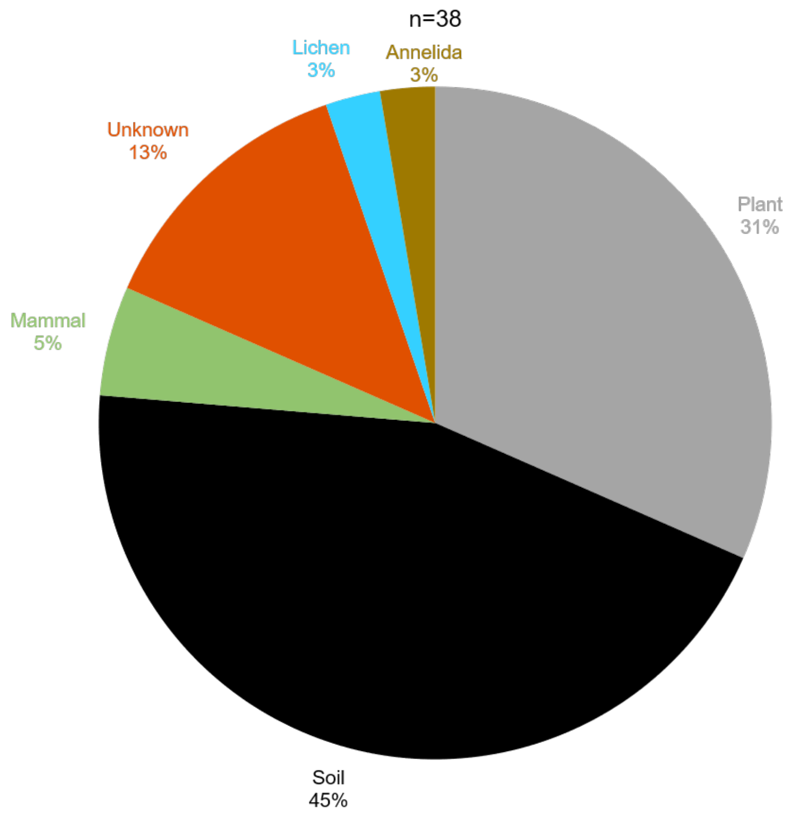


**Figure S5.** Habitat distribution of the Diels-Alderase positive isolates found by genome mining to have an abyssomicin or potential abyssomicin BGC (both total and partial).

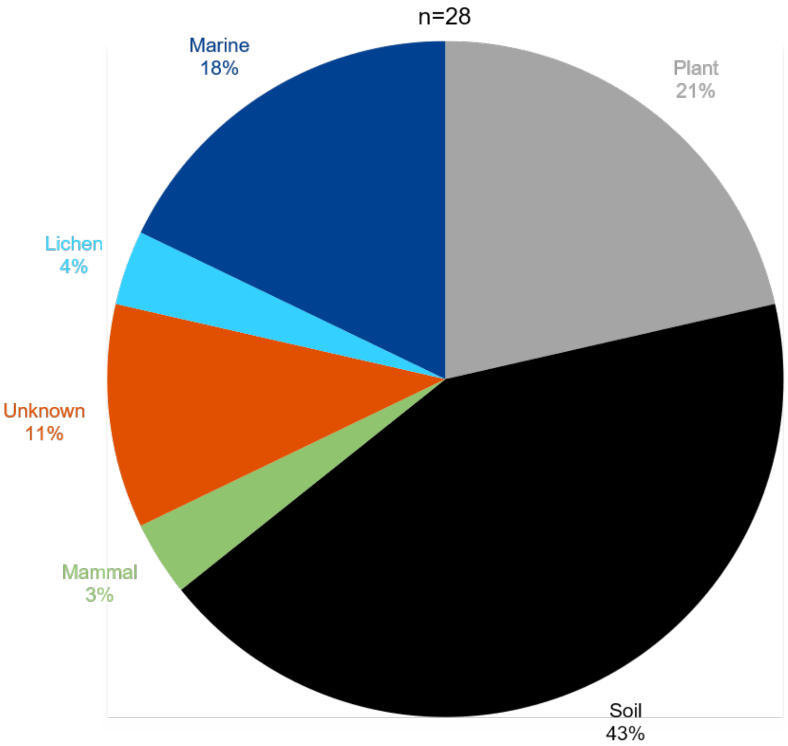


**Figure S6.** Habitat distribution of the Diels-Alderase positive isolates found by genome mining that do not harbour any abyssomicin nor potential abyssomicin BGC.

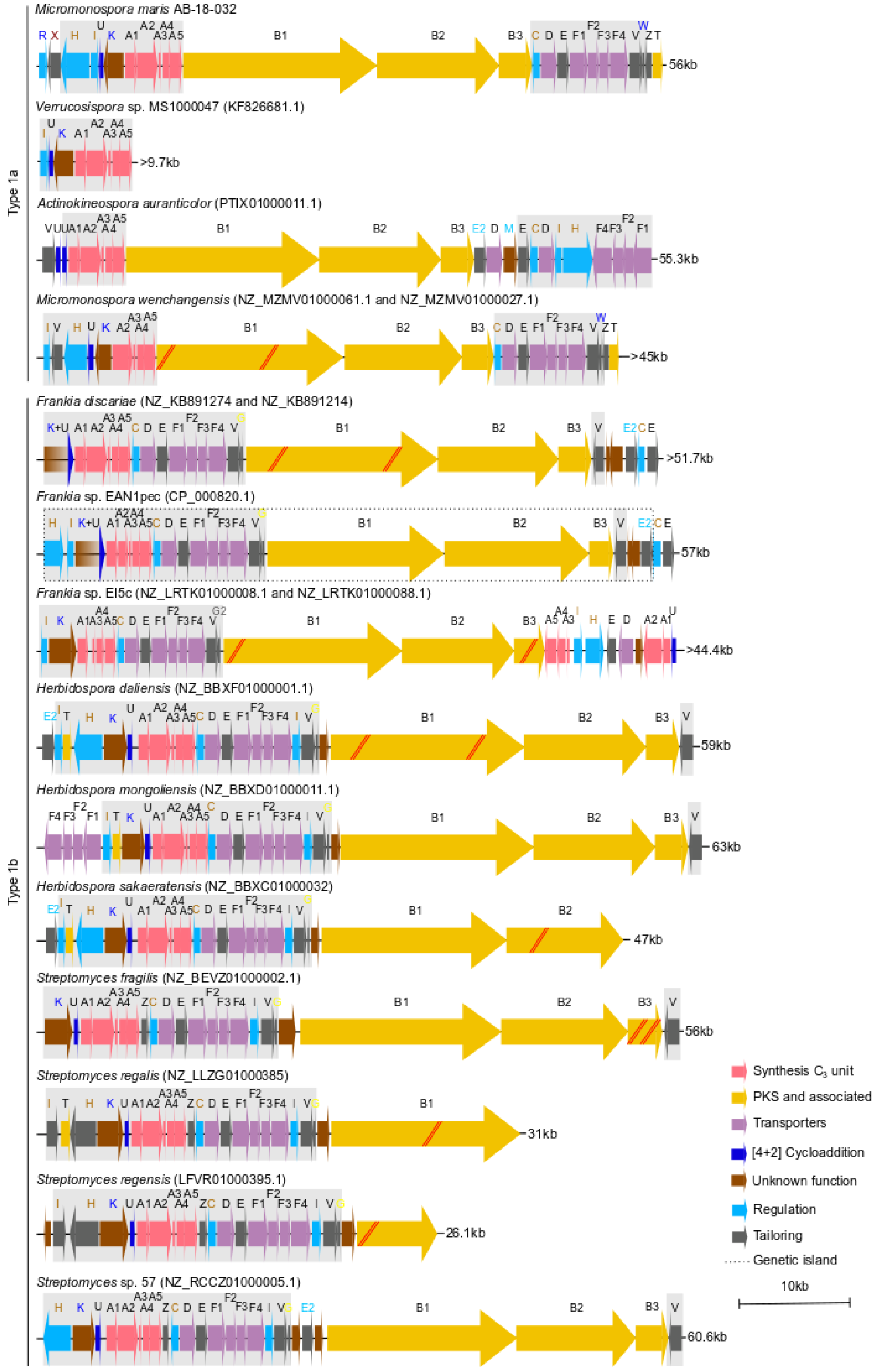


**Figure S7.** Abyssomicin and potential abyssomicin BGCs recovered from genome mining classified as clusters type 1a and 1b. Gene names in black are common to *aby*, *abs* and *abm* BGCs. Blue font represents genes present only in *M. maris* AB-18-032, grey font represents genes present only in *Streptomyces* sp. LC-6-2 and light blue font represent genes unique to *S. koyangensis* SCSIO 5802. In maroon font appear those genes that appear both in *aby* and *abs* BGCs, in light brown those genes that appear both in *aby* and *abm* BGCs and in yellow those genes that appear both in *abs* and *abm* BGCs. Grey boxes indicate the conserved regions shared between the type 1 clusters. Dotted lines indicate genetic islands.

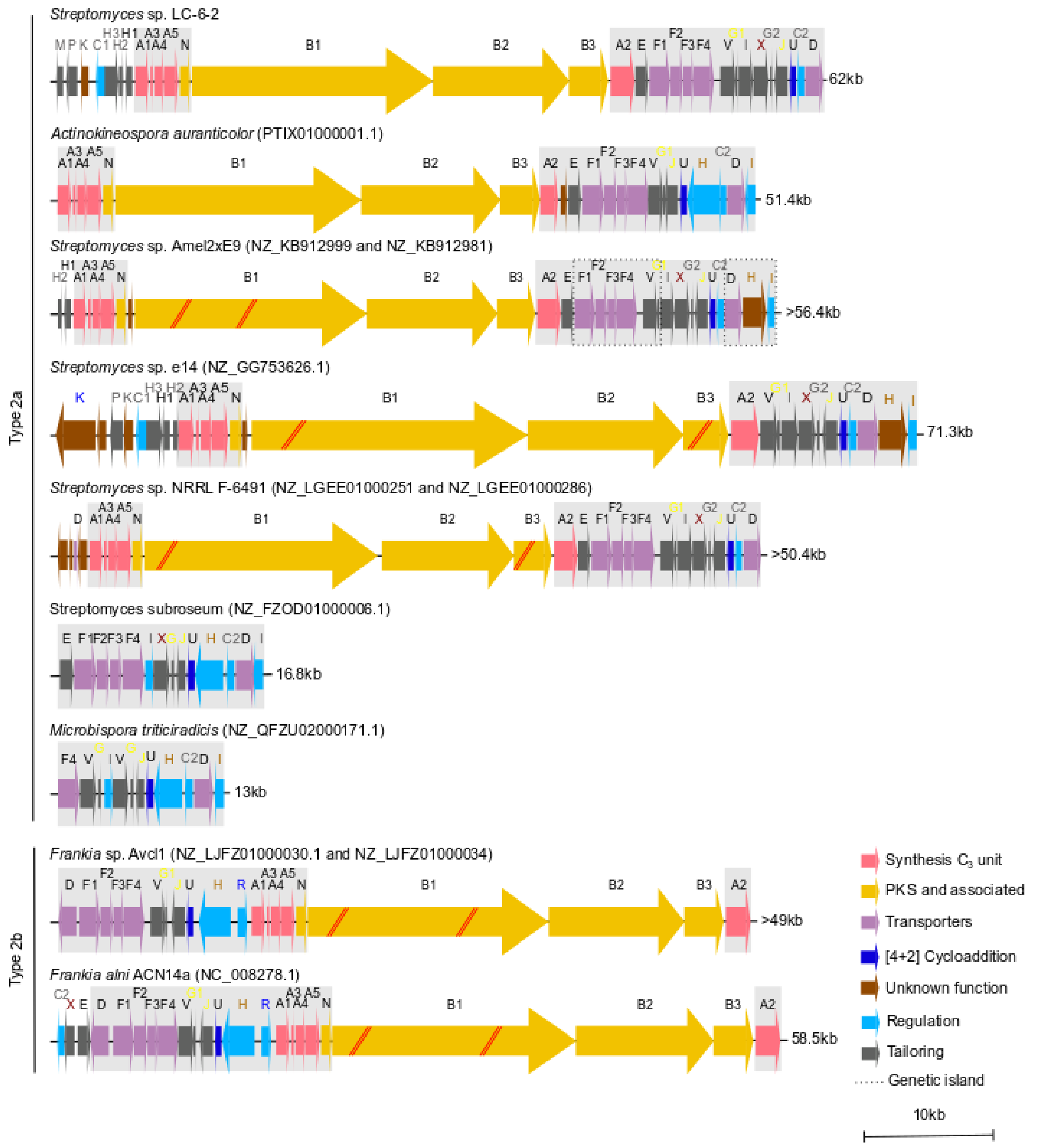


**Figure S8.** Abyssomicin and potential abyssomicin BGCs recovered from genome mining classified as clusters type 2a and 2b. Gene names in black are common to *aby*, *abs* and *abm* BGCs. Blue font represents genes present only in *M. maris* AB-18-032, grey font represents genes present only in *Streptomyces* sp. LC-6-2 and light blue font represent genes unique to *S. koyangensis* SCSIO 5802. In maroon font appear those genes that appear both in *aby* and *abs* BGCs, in light brown those genes that appear both in *aby* and *abm* BGCs and in yellow those genes that appear both in *abs* and *abm* BGCs. Grey boxes indicate the conserved regions shared between the type 2 clusters. Dotted lines indicate genetic islands.

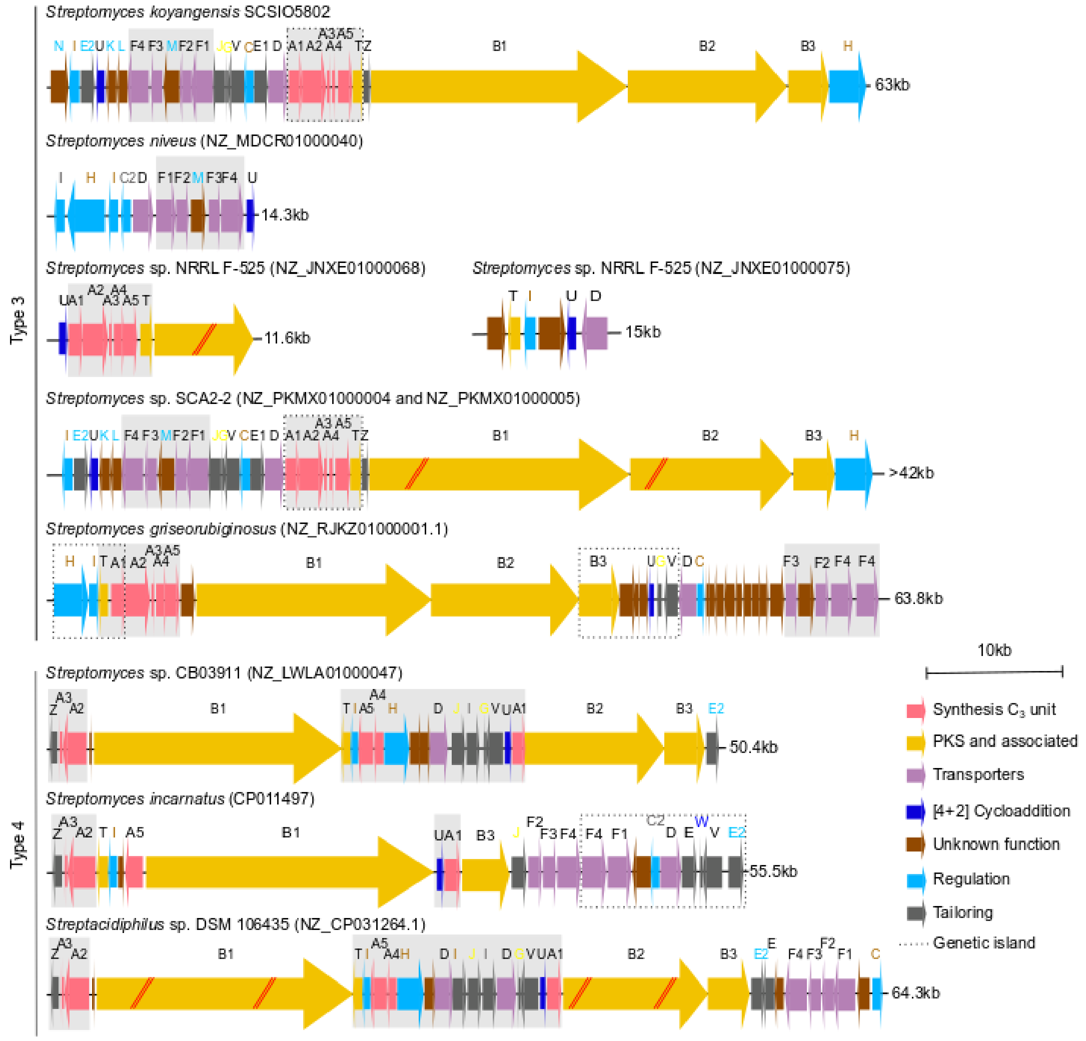


**Figure S9.** Abyssomicin and potential abyssomicin BGCs recovered from genome mining classified as clusters type 3 and 4. Gene names in black are common to aby, abs and abm BGCs. Gene names in black are common to *aby*, *abs* and *abm* BGCs. Blue font represents genes present only in *M. maris* AB-18-032, grey font represents genes present only in *Streptomyces* sp. LC-6-2 and light blue font represent genes unique to *S. koyangensis* SCSIO 5802. In maroon font appear those genes that appear both in *aby* and *abs* BGCs, in light brown those genes that appear both in *aby* and *abm* BGCs and in yellow those genes that appear both in *abs* and *abm* BGCs. Grey boxes indicate the conserved regions shared between the type 3 and type 4 clusters. Dotted lines indicate genetic islands.

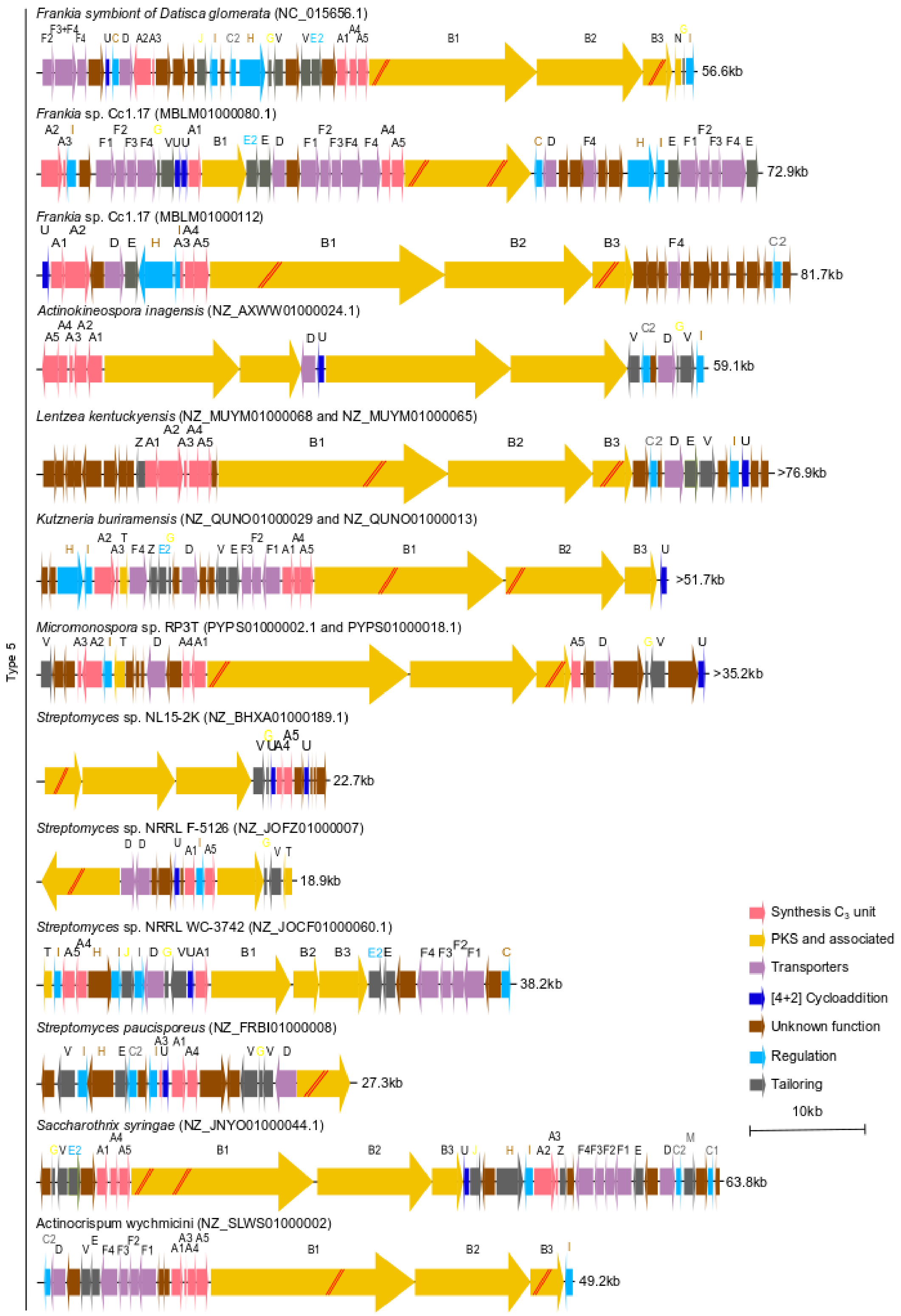


**Figure S10.** Abyssomicin and potential abyssomicin BGCs recovered from genome mining classified as clusters type 5. Gene names in black are common to *aby*, *abs* and *abm* BGCs. Blue font represents genes present only in *M. maris* AB-18-032, grey font represents genes present only in *Streptomyces* sp. LC-6-2 and light blue font represent genes unique to *S. koyangensis* SCSIO 5802. In maroon font appear those genes that appear both in *aby* and *abs* BGCs, in light brown those genes that appear both in *aby* and *abm* BGCs and in yellow those genes that appear both in *abs* and *abm* BGCs.

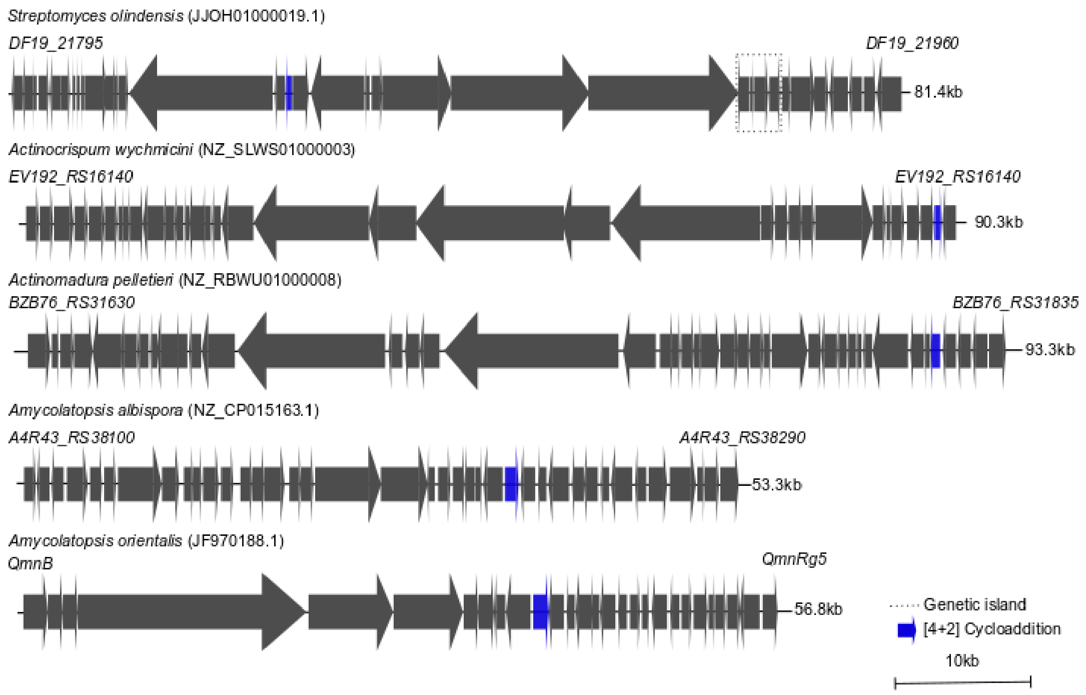


**Figure S11.** Potential tetronomycin BGC in *S. olindensis* DAUFPE 5622, potential chlorothricin BGC from *A. wychmicini* DSM 45934 and *A. pelletieri* DSM 43383 and quartromycin BGCs from *A. albispora* WP1 and *A. orientalis* Q427-8. The Diels-Alderase homologs are displayed in blue. Dotted lines indicate genetic islands.

**Table S1.** Abyssomicin producing bacteria as reported in literature.

| Microorganism | Isolation location | Habitat | Reference | Accession | Abyssomicin structure |
|---|---|---|---|---|---|
| *Micromonospora maris* AB-18-032 | Sediment Sea of Japan (-289 m) | Aquatic | (Riedlinger *et al.*, 2004) | JF752342 | B, C, atrop-C, D, G, H |
| *Streptomyces sp.* HKI0381 | Soil Ile de Paradis (Senegal) | Terrestrial | (Niu *et al.*, 2007) | - | E |
| *Streptomyces sp.* CHI39 | Rock soil Campeche (Mexico) | Terrestrial | (Igarashi *et al.*, 2010) | - | I |
| *Streptomyces sp.* Ank 210 | Forest soil Kaiserslautern (Germany) | Terrestrial | (Abdalla *et al.*, 2011) | - | ent-homoA, ent-homoB |
| *Verrucosispora sp.* MS100128 | South China Sea deep-sea sediment (−2733 m) | Aquatic | (Wang *et al.*, 2013) | - | B, C, atrop-C, D, H, J, K, L |
| *Streptomyces sp.* RLUS1487 | Marine | Aquatic | (León *et al.*, 2015) | - | 2, 3, 4, 5 |
| *Verrucosispora sp.* MS100047 | South China Sea sediment | Aquatic | (Huang *et al.*, 2016) | KF826681 | B |
| *Streptomyces sp.* LC-6-2 | Soil Lotts Creek coal fire (United States) | Terrestrial | (Wang *et al.*, 2017) | KY432814 | M, N, O, P, Q, R, S, T, U, V, W, X |
| *Streptomyces koyangensis* SCSIO 5802 | South China Sea sediment (−3536 m) | Aquatic | (Song *et al.*, 2017) | MG243704 | 2, 4, neo-A, neo-B, neo-C, neo-D, neo-E, neo-F, neo-G, neo-2 |

**Table S2.** Isolation details of the abyssomicin producers isolated from aquatic environments.

| Microorganism | Isolation details | Reference |
|---|---|---|
| *Micromonospora maris* AB-18-032 | Isolated on a colloidal chitin agar plate which had been inoculated with a suspension of a sediment sample collected from the Sea of Japan and incubated at 30°C for 4 weeks. | (Riedlinger *et al.*, 2004) |
| *Verrucosispora sp.* MS100128 | Isolated using oatmeal agar from a sediment sample collected from the South China Sea at 2733 m below sea level. | (Wang *et al.*, 2013) |
| *Streptomyces sp.* RLUS1487 | Isolated on AIS medium from a marine sediment sample collected by SCUBA near American Samoa. | (León *et al.*, 2015) |
| *Verrucosispora sp.* MS100047 | Isolated on a VER01 agar plate from a sediment sample collected in the South China Sea at 28 °C. | (Huang *et al.*, 2016) |
| *Streptomyces koyangensis* SCSIO 5802 | Isolated from a sediment sample collected from the South China Sea at a depth of -3536 m. | (Song *et al.*, 2017) |

**Table S3.** Nucleotide and protein sequences of the Diels Alderase proteins present in *aby*, *abs* and *abm* BGCs.

| **Gene/Protein** | **Sequence** |
|---|---|
| abyU | atgactgagcgactggagacgcgaccgcaggccctgctcatcaaggtgcccaccgagatcgtggtgaaggtggtcgacgacgtggacgtggccgctccggcggtggggcaggtgggcaaattcgacgacgagttgtacgacgaggccggtgcccagatcggcacgtccagcggcaacttccgcatcgagtacgtgcgaccgaccgacggcggactgctcacctactaccaggaggacatcactctctccgatggggtgatccacgcggagggctgggcggacttcaacgacgtgcggacgagtaagtgggtgttctacccggcgaccggggtgagcggccgctacctgggcctcaccggcttccggcagtggcggatgacgggcgtgcgcaagtccgccgaggcgcggatcctgctcggcgagtga |
| abmU | atgaacgaacgcttcaccctgcccgcccacagccccgccctcgcggcgctcgtccccgagttcctcgacctggcgcgagccgcgagcggcgatccggccgccgaggagcgcgacctcgcggtctgggagaacctcacggaacacgtctcgctggactaccggttcgccaacccgcccgtgcacggtcccggcgactgggacacgtacgacagccgcttcgtggaccccgccggcgtggagatcggcaccctccagggcaccggacgcatcctgtacgagcgttcgtcggacgcgcacctgatgatgtactaccgcgagcagctgaccttccccgacgggacggcccagaccgcgggctgggtcgacggcaccgcgatcctcggcggggcctggcagcgcttccccatcctggggtcgggcggccggtacggctccatgatcgggctgcgctccttccagcccacccccgaggcgccgcacagcctctaccgcacccacctggtgctccgggagatccccggcgggcacgggctgaccgaccccgaggagatcgacgcggcactgtcgctgctcggcgccttcgtgggcccctcggtcaacccggcgaccggcaacggccgcctcgaaccccccgtacgcgccgggcgcaccgcctga |
| absU | gtggtgttgcaggtcctgtccgactggctcacgccgctggtcgcgacgcccccgaagaccgtctcgccggaggtcggcgccctcaaggacacgggcaggtcgctcatcctgcgcgacctgagggagaaggtggtcgcctacgagtcgaacaaccccgaccccaccggcaccacccccaccgagaacgacttcgccacggtccggctggagatcttcggccccgacggtacgcagatcgggaccaccgagggcgccgggcggatgctgtaccggcaggagaaggacgagcacttcatcgcctacttcggcgaggagatcacgctcaacgacggcaacgtcatccgcgcgggcgggctcgtggacgacgcgcggctgacggcgggcgaacacgccacgttccccgcggtggtggtcagcgggccgctgcgcggcgcgatcggcttccgccagttccggccgctggtcaaggagtcgcacacgacgtacgagtcctcgatcgtcgtctaccggaggtga |
| AbyU | MTERLETRPQALLIKVPTEIVVKVVDDVDVAAPAVGQVGKFDDELYDEAGAQIGTSSGNFRIEYVRPTDGGLLTYYQEDITLSDGVIHAEGWADFNDVRTSKWVFYPATGVSGRYLGLTGFRQWRMTGVRKSAEARILLGE |
| AbmU | MNERFTLPAHSPALAALVPEFLDLARAASGDPAAEERDLAVWENLTEHVSLDYRFANPPVHGPGDWDTYDSRFVDPAGVEIGTLQGTGRILYERSSDAHLMMYYREQLTFPDGTAQTAGWVDGTAILGGAWQRFPILGSGGRYGSMIGLRSFQPTPEAPHSLYRTHLVLREIPGGHGLTDPEEIDAALSLLGAFVGPSVNPATGNGRLEPPVRAGRTA |
| AbsU | MVLQVLSDWLTPLVATPPKTVSPEVGALKDTGRSLILRDLREKVVAYESNNPDPTGTTPTENDFATVRLEIFGPDGTQIGTTEGAGRMLYRQEKDEHFIAYFGEEITLNDGNVIRAGGLVDDARLTAGEHATFPAVVVSGPLRGAIGFRQFRPLVKESHTTYESSIVVYRR |

**Table S4.** Identity percentage at protein (above) and nucleotide (below) level between the Diels Alderase proteins present in *aby*, *abs* and *abm* BGCs.

<table>
<tr><th></th><th>AbyU</th><th>AbmU</th><th>AbsU</th></tr>
<tr><td rowspan="2">AbyU</td><td rowspan="2">*</td><td>37%</td><td>31%</td></tr>
<tr><td>56%</td><td>55%</td></tr>
<tr><td rowspan="2">AbmU</td><td>37%</td><td rowspan="2">*</td><td>30%</td></tr>
<tr><td>56%</td><td>57%</td></tr>
<tr><td rowspan="2">AbsU</td><td>31%</td><td>30%</td><td rowspan="2">*</td></tr>
<tr><td>55%</td><td>57%</td></tr>
</table>

**Table S5.** Classification of metagenomic samples from aquatic environments mined for AbyU, AbsU and AbmU. Numbers represent Diels-Alderase positive/mined metagenomes.

| | | | | | |
|---|---|---|---|---|---|
| Environmental | Aquatic | Freshwater | Drinking water | Unchlorinated | 0/9 |
| | | | | Unclassified | 0/8 |
| | | | Ground water | Acid Mine Drainage | 0/7 |
| | | | | Cave water | 0/11 |
| | | | | Contaminated | 0/14 |
| | | | | Mine drainage | 0/4 |
| | | | | Unclassified | 0/71 |
| | | | Lentic | Epilimnion | 0/25 |
| | | | | Hypolimnion | 0/8 |
| | | | | Littoral zone | 0/2 |
| | | | | Sediment | 0/15 |
| | | | | Unclassified | 0/60 |
| | | | Lotic | Sediment | 0/9 |
| | | | | Unclassified | 0/12 |
| | | | Pond | Sediment | 0/12 |
| | | | | Unclassified | 0/7 |
| | | | River | Unclassified | 0/39 |
| | | | Sediment | Unclassified | 0/41 |
| | | | Unclassified | Unclassified | 0/18 |
| | | | Ice | Glacial Lake | 0/19 |
| | | | | Glacier | 0/9 |
| | | | | Ice accretions | 0/1 |
| | | | Lake | Sediment | 0/46 |
| | | | Wetlands | Bog | 0/25 |
| | | Marine | Coastal | Sediment | 0/25 |
| | | | Intertidal zone | Estuary | 0/25 |
| | | | | Salt marsh | 0/19 |
| | | | | Sediment | 0/10 |
| | | | | Unclassified | 0/40 |
| | | | Oceanic | Abyssal plane | 0/3 |
| | | | | Aphotic zone | 0/22 |
| | | | | Sediment | 0/57 |
| | | | | Unclassified | 0/94 |
| | | Non-marine Saline and Alkaline | Alkaline | Sediment | 0/10 |
| | | | | Unclassified | 0/7 |
| | | | Hypersaline | Microbial mats | 0/6 |
| | | | Saline | Epilimnion | 0/1 |
| | | | | Sediment | 0/4 |
| | | | | Unclassified | 0/20 |

**Table S6.** Classification of metagenomic samples from terrestrial environments mined for AbyU, AbsU and AbmU. Numbers represent Diels-Alderase positive/mined metagenomes.

| | | | | | |
|---|---|---|---|---|---|
| Environmental | Terrestrial | Soil | Crop | Agricultural land | 0/17 |
| | | | Loam | Agricultural soil | 18/57 |
| | | | | Forest Soil | 0/26 |
| | | | | Grasslands | 2/42 |
| | | | | Unclassified | 0/8 |
| | | | Clay | Oil-contaminated | 0/8 |
| | | | | Unclassified | 1/18 |
| | | | Fossil | Unclassified | 0/1 |
| | | | Hot | Acidic | 0/4 |
| | | | Sand | Desert | 1/36 |
| | | | | Oil contaminated | 0/2 |
| | | | | Unclassified | 1/19 |
| | | | Unclassified | Agricultural | 7/18 |
| | | | | Agricultural land | 0/84 |
| | | | | Desert | 0/20 |
| | | | | Forest Soil | 5/233 |
| | | | | Grasslands | 1/32 |
| | | | | Permafrost | 0/45 |
| | | | | Shrubland | 0/1 |
| | | | | Tropical rainforest | 0/17 |
| | | | Wetlands | Permafrost | 0/12 |
| | | | | Unclassified | 0/50 |
| | | Geologic | Mine | Unclassified | 0/4 |
| | | | Sediment | Unclassified | 0/16 |
| | | Plant litter | Unclassified | Unclassified | 8/130 |
| | | Rock-dwelling (endoliths) | Unclassified | Unclassified | 0/18 |
| | | Rock-dwelling (subaerial biofilms) | Unclassified | Unclassified | 0/8 |
| | | Volcanic | Fumaroles | Unclassified | 0/9 |

**Table S7.** Classification of metagenomic samples from engineered environments mined for AbyU, AbsU and AbmU. Numbers represent Diels-Alderase positive/mined metagenomes.

| | | | | | |
|---|---|---|---|---|---|
| Engineered | Built environment | City | Subway | Unclassified | 0/34 |
| | | Solar panel | Unclassified | Unclassified | 0/3 |
| | | Unclassified | Unclassified | Unclassified | 0/4 |
| | Solid waste | Composting | Grass | Bioreactor | 0/11 |
| | | | | Unclassified | 0/1 |
| | | | Unclassified | Unclassified | 0/23 |
| | | | Wood | Bioreactor | 0/4 |
| | | Landfield | Unclassified | Unclassified | 0/3 |
| | | Solid Animal Waste | Unclassified | Unclassified | 0/6 |
| | Bioreactor | Aerobic | Unclassified | Unclassified | 6/6 |
| | | Continuous culture | Marine intertidal flat sediment inoculum | Unclassified | 0/2 |
| | | | Marine sediment inoculum | Unclassified | 0/11 |
| | | Unclassified | Unclassified | Unclassified | 0/17 |
| | Biotransformation | Microbial solubilization of coal | Unclassified | Unclassified | 0/4 |
| | | Mixed alcohol bioreactor | Unclassified | Unclassified | 0/2 |
| | Waste water | Activated Sludge | Unclassified | Unclassified | 0/21 |
| | | Anaerobic digestor | Unclassified | Unclassified | 0/20 |
| | | Industrial waste water | Mine water | Unclassified | 0/2 |
| | | | Petrochemical | Unclassified | 0/14 |
| | | | Unclassified | Unclassified | 0/13 |

**Table S8.** Classification of metagenomic samples from host-associated environments mined for AbyU, AbsU and AbmU. Numbers represent Diels-Alderase positive/mined metagenomes.

| | | | | | |
|---|---|---|---|---|---|
| Host-associated | Algae | Green algae | Ectosymbionts | Unclassified | 0/11 |
| | | Red algae | Ectosymbionts | Unclassified | 0/15 |
| | Animal | Digestive system | Fecal | Unclassified | 0/18 |
| | | Skin | Unclassified | Unclassified | 0/1 |
| | Annelida | Integument | Cuticle | Epibionts | 0/1 |
| | | | Unclassified | Unclassified | 0/8 |
| | | Intracellular endosymbiont | Trophosome | Unclassified | 0/2 |
| | | Reproductive system | Egg capsule | Unclassified | 0/2 |
| | | Unclassified | Unclassified | Unclassified | 0/1 |
| | Arthropoda | Ant dump | Unclassified | Unclassified | 0/11 |
| | | Digestive system | Foregut | Unclassified | 0/1 |
| | | | Gut | P3 segment | 0/1 |
| | | | | Proctodeal segment | 0/1 |
| | | | | Unclassified | 1/40 |
| | | | Midgut | Unclassified | 0/4 |
| | | Intracellular endosymbiont | Secondary | Unclassified | 0/1 |
| | | | Unclassified | Unclassified | 1/1 |
| | | Symbiotic fungal gardens and galleries | Fungus gallery | Unclassified | 0/1 |
| | | | Fungus garden | Garden dump | 0/1 |
| | | | | Unclassified | 1/9 |
| | | | Unclassified | Unclassified | 0/6 |
| | | Tissue | Unclassified | Unclassified | 0/15 |
| | | Unclassified | Unclassified | Unclassified | 1/3 |
| | Birds | Digestive system | Ceca | Lumen | 0/1 |
| | | | Crop | Lumen | 0/9 |
| | Cnidaria | Unclassified | Unclassified | Unclassified | 0/33 |
| | Fish | Digestive system | Unclassified | Unclassified | 0/1 |
| | | Skin | Epidermal mucus | Unclassified | 0/3 |
| | Fungi | Mycelium | Unclassified | Unclassified | 0/41 |
| | | Unclassified | Unclassified | Unclassified | 0/11 |
| | Insecta | Digestive system | Unclassified | Unclassified | 0/14 |
| | Invertebrates | Cnidaria | Coral | Unclassified | 0/15 |
| | Mammals | Digestive system | Fecal | unclassified | 0/17 |
| | | | Foregut | Rumen | 0/18 |
| | | | | Unclassified | 0/12 |
| | | | Stomach | Rumen | 0/6 |
| | | | | Unclassified | 0/2 |
| | | Nervous system | Brain | Unclassified | 0/11 |
| | | Tissue | Unclassified | Unclassified | 0/8 |
| | Microbial | Bacteria | Unclassified | Unclassified | 0/8 |
| | | Dinoflagellates | Endosymbionts | Unclassified | 0/2 |
| | | | Unclassified | Unclassified | 0/7 |
| | Mollusca | Digestive system | Ceca | Uncharacterized | 0/1 |
| | | Respiratory system | Gills | Extracellular | 0/7 |
| | | Shell | Unclassified | Unclassified | 0/2 |
| | Plants | Endosphere | Unclassified | Unclassified | 1/11 |
| | | Leaf | Unclassified | Unclassified | 0/5 |
| | | Nodule | Unclassified | Unclassified | 0/4 |
| | | Peat moss | Unclassified | Unclassified | 0/14 |
| | | Phylloplane | Epiphytes | Unclassified | 5/32 |
| | | | Unclassified | Unclassified | 0/27 |
| | | Phyllosphere | Unclassified | Unclassified | 4/57 |
| | | Rhizoplane | Endophytes | Unclassified | 0/1 |

| | | | | | |
|---|---|---|---|---|---|
| | | | Epiphytes | Unclassified | 19/106 |
| | | | Soil | Unclassified | 2/2 |
| | | | Unclassified | Unclassified | 1/54 |
| | | Rhizosphere | Epiphytes | Unclassified | 0/2 |
| | | | Soil | Unclassified | 11/116 |
| | | | Unclassified | Unclassified | 41/157 |
| | | Roots | Nodule | Unclassified | 0/2 |
| | | | Unclassified | Unclassified | 12/58 |
| | | Wood | Unclassified | Unclassified | 0/16 |
| | Porifera | Unclassified | Unclassified | Unclassified | 0/19 |
| | Tunicates | Ascidians | Unclassified | Unclassified | 0/9 |
| | | Unclassified | Unclassified | Unclassified | 0/1 |

**Table S9.** Proteins within the non-redundant sequence database (NCBI) with significative alignments to AbyU, AbmU, AbsU and VASRM7_509. Columns displaying the Diels-Alderase homologs show protein ID, alignment E-value and similarity percentage.

| Microorganism | AbyU homolog | AbsU homolog | AbmU homolog | Genome accession, number of contigs, sequencing technology | Protein location |
|---|---|---|---|---|---|
| *Actinokineospora auranticolor* YU 961-1 | WP_104480634.1 (5e-31/44.62%) | WP_104476817.1 (4e-62/70.99%) | WP_104476817.1 (9e-11/34.44%) | NZ_PTIX00000000.1, 61 contigs, Illumina HiSeq | 2 Abyssomicin BGCs, total |
| | WP_104480633.1 (4e-11/33.68%) | | | | |
| | WP_104476817.1 (2e-07/38.67%) | | | | |
| *Actinomadura* sp. H3C3 | WP_131902816.1 (1e-07/30.97%) | WP_131902816.1 (9e-09/27.34%) | WP_131902816.1 (2e-10/31.90%) | SMKU00000000.1, 805 contigs, Illumina HiSeq | Not enough data |
| *Amycolatopsis albispora* WP1 | WP_113696543.1 (1e-13/38.10%) | WP_113696543.1 (2e-06/32.38%) | WP_113696543.1 (8e-11/30.67%) | NZ_CP015163.1, Complete genome, PacBio RSII | Quartromicin BGC, partial |
| *Streptomyces armeniacus* ATCC 15676 | AXK36488.1 (4e-13/34.88%) | AXK36488.1 (4e-08/32.58%) | AXK36488.1 (6e-12/28.23%) | CP031320.1, 21 contigs, Illumina MiSeq | Not a BGC |
| *Amycolatopsis* sp. CA-126428 | WP_103337399.1 (4e-16/36.19%) | No hit | WP_103337399.1 (5e-12/37.37%) | NZ_PPHF00000000.1, 188 contigs, Illumina HiSeq | Not enough data |
| *Streptomyces caatingaensis* CMAA 1322 | WP_049718340.1 (4e-20/40.37%) | WP_049718340.1 (1e-11/36.17%) | WP_049718340.1 (5e-30/38.41%) | NZ_LFXA00000000.1, 18 contigs, PacBio | Not a BGC |
| *Streptomyces cattleya* DSM 46488 | WP_014140910.1 (3e-21/34.55%) | WP_014140910.1 (2e-14/36.90%) | WP_014140910.1 (4e-14/34.31%) | NC_017586.1, Complete genome, no data | Potential BGC |
| *Streptomyces* sp. CB03911 | WP_073928710.1 (7e-08/30.43%) | WP_073928710.1 (2e-45/57.81%) | WP_073928710.1 (1e-06/29.17%) | NZ_LWLA00000000.1, 49 contigs, Illumina MiSeq | Potential abyssomicin BGC, partial |
| *Streptomyces* sp. E5N91 SAI-083 | WP_123627591.1 (6e-22/34.55%) | WP_123627591.1 (9e-15/36.90%) | WP_123627591.1 (8e-13/32.35%) | NZ_RJKF00000000.1, 2 contigs, PacBio | Potential BGC, total |
| *Micromonospora eburnea* DSM 44814 | No hit | No hit | WP_091120898.1 (2e-14/34.23%) | NZ_FMHY00000000.1, 2 contigs, no data | Potential BGC, partial |
| *Frankia* sp. AvcI1 | WP_055751820.1 (1e-08/29.70%) | WP_055751820.1 (1e-64/73.44%) | WP_055751820.1 (1e-09/30.53%) | NZ_LJFZ00000000.1, 77 contigs, Illumina HiSeq | Abyssomicin BGC, partial |
| *Actinomadura fibrosa* LMG 29177 | WP_131760470.1 (2e-12/32.71%) | No hit | WP_131760470.1 (6e-07/30.70%) | CAACUY000000000, 569 contigs, no data | Not enough data |
| *Frankia alni* ACN14A | WP_011605212.1 (2e-08/29.70%) | WP_011605212.1 (1e-65/74.22%) | WP_011605212.1 (1e-10/31.58%) | NC_008278, Complete genome, no data | Abyssomicin BGC, partial |
| *Frankia discariae* BCU110501 | WP_018506019.1 (1e-54/74.05%) | WP_018506019.1 (1e-08/36.14%) | WP_018506019.1 (4e-14/36.96%) | NZ_ARDT00000000.1, 200 contigs, no data | Abyssomicin BGC, partial |
| *Frankia* sp. EAN1pec | WP_020461028.1 (4e-56/75.57%) | WP_020461028.1 (2e-08/36.14%) | WP_020461028.1 (3e-14/35.64%) | NC_009921.1, Complete genome, no data | Abyssomicin BGC, total |
| *Frankia* sp. EI5c | WP_066064649.1 (2e-19/34.31%) | WP_066064649.1 (1e-06/29.29% ) | WP_066064649.1 (1e-17/42.05%) | NZ_LRTK01000000, 159 contigs, Illumina HiSeq | Abyssomicin BGC, partial |
| *Frankia* symbiont of *Datisca glomerata* | WP_043605928.1 (5e-12/35.19%) | WP_043605928.1 (6e-51/62.50%) | WP_043605928.1 (3e-12/33.67%) | NC_015656.1, Complete genome, 454/Illumina | Potential abyssomicin BGC, partial |
| *Frankia* sp. Cc1.17 | WP_071083475.1 (1e- | WP_071084438.1 (2e- | WP_071084438.1 (1e- | NZ_MBLM00000000, | 2 Abyssomicin BGCs, partial |

| | | | | | |
|---|---|---|---|---|---|
| | 35/51.33%)<br>WP_071084438.1 (3e-19/34.31%)<br>WP_131803042.1 (6e-10/28.97%) | 06/28.28%) | 17/42.05%)<br>WP_071083475.1 (8e-06/28.57%) | 195 contigs, Illumina HiSeq | |
| *Photobacterium ganghwense* JCM 12487 | WP_047885918.1 (1e-09/34.26%) | No hit | WP_047885918.1 (4e-12/28.49%) | NZ_PYMI00000000,<br>39 contigs, Illumina MiSeq | Not a BGC |
| *Streptomyces geranii* A301 | WP_105971044.1 (1e-18/36.09%) | WP_105971044.1 (4e-10/34.19%) | WP_105971044.1 (7e-16/36.80%) | NZ_PJME00000000.1,<br>104 contigs, Illumina HiSeq | Not enough data |
| *Streptomyces griseocarneus* 132 | WP_121798615.1 (3e-26/40.00%) | WP_121798615.1 (3e-07/29.41%) | WP_121798615.1 (2e-10/27.21%) | NZ_PENC00000000,<br>227 contigs, Illumina HiSeq | Not enough data |
| *Streptomyces griseorubiginosus* SAI-142 | WP_123763217.1 (1e-15/38.04%) | WP_123763217.1 (1e-07/28.69%) | WP_123763217.1 (3e-14/31.90%) | NZ_RJKZ00000000,<br>4 contigs, PacBio | Potential abyssomicin BGC, partial |
| *Herbidospora daliensis* NBRC 106372 | WP_062428853.1 (4e-63/76.56%) | WP_062428853.1 (5e-10/37.35%) | WP_062428853.1 (2e-15/38.64%) | NZ_BBXF01000001,<br>Complete genome, Illumina MiSeq | Abyssomicin BGC, partial |
| *Herbidospora mongoliensis* NBRC 105882 | WP_066363831.1 (6e-63/76.56%) | WP_066363831.1 (4e-09/32.98%) | WP_066363831.1 (9e-15/37.50%) | NZ_BBXD00000000,<br>47 contigs, Illumina MiSeq | Abyssomicin BGC, total |
| *Herbidospora sakaeratensis* NBRC 102641 | WP_062343027.1 (2e-63/75.38%) | WP_062343027.1 (6e-10/37.35%) | WP_062343027.1 (2e-15/38.64%) | NZ_BBXC00000000,<br>45 contigs, Illumina MiSeq | Abyssomicin BGC, partial |
| *Actinokineospora inagensis* DSM 44258 | WP_084467520.1 (9e-16/37.74%) | No hit | WP_084467520.1 (7e-10/37.00%) | NZ_AXWW00000000,<br>106 contigs,Illumina HiSeq 2000 | Potential abyssomicin BGC, partial |
| *Streptomyces iranensis* DSM 41954 | WP_078957139.1 (8e-23/32.54%) | WP_078957139.1 (9e-15/38.10%) | WP_078957139.1 (8e-13/32.58%) | NZ_LK022848,<br>Complete genome, no data | Potential BGC, partial |
| *Lentzea kentuckyensis* NRRL B-24416 | WP_086666194.1 (3e-13/38.30%) | No hit | WP_086666194.1 (1e-06/28.03%) | NZ_MUYM00000000,<br>317 contigs, Illumina MiSeq | Potential abyssomicin BGC, partial |
| *Kutzneria buriramensis* DSM 45791 | WP_116178805.1 (2e-10/33.94% ) | WP_116178805.1 (3e-50/61.72%) | WP_116178805.1 (8e-10/28.23%) | NZ_QUNO00000000,<br>65 contigs,Illumina HiSeq | Potential abyssomicin BGC, partial |
| *Streptomyces* sp. LHW50302 | WP_114017401.1 (2e-05/27.27%) | No hit | WP_114017401.1 (2e-14/33.83%) | NZ_QOIM00000000,<br>70 contigs,Illumina HiSeq | Not enough data |
| *Microbispora triticiradicis* NEAU-HRDPA2-9 | WP_117409467.1 (2e-07/33.73%) | WP_117409467.1 (1e-76/80.60%) | WP_117409467.1 (2e-11/29.01%) | NZ_QFZU00000000,<br>285 contigs,Illumina HiSeq | Potential abyssomicin BGC, partial |
| *Micromonospora wenchangensis CCTCC AA 2012002* | WP_088646687.1 (2e-72/85.11%) | WP_088646687.1 (7e-11/38.55%) | WP_088646687.1 (8e-13/35.63%) | NZ_MZMV00000000.1,<br>150 contigs,Illumina HiSeq | Abyssomicin BGC, partial |
| *Micromonospora* sp. RP3T | WP_107157684.1 (2e-23/40.19%) | WP_107157684.1 (6e-16/32.85%) | WP_107157684.1 (2e-25/38.40%) | NZ_PYPS00000000,<br>174 contigs,Illumina HiSeq | Potential abyssomicin BGC, partial |
| *Streptomyces monomycini* NRRL B-24309 | WP_050502808.1 (2e-15/36.08%) | WP_050502808.1 (5e-07/33.72% ) | WP_050502808.1 (7e-24/37.59%) | NZ_JNYL00000000,<br>643 contigs,Illumina | Not enough data |
| *Streptomyces* sp. MUSC 14 | WP_071375955.1 (1e-24/41.44%) | WP_071375955.1 (2e-13/35.63%) | WP_071375955.1 (8e-11/30.39%) | NZ_MLYN00000000,<br>174 contigs, Illumina MiSeq | Potential BGC, partial |
| *Streptomyces niveus* NRRL 2466 | WP_069626275.1 (9e-11/29.70%) | WP_069626275.1 (5e-15/28.87%) | WP_069626275.1 (4e-76/53.02%) | NZ_MDCR00000000,<br>608 contigs, Illumina MiSeq | Potential abyssomicin BGC, partial |
| *Streptomyces* sp. NL15-2K | WP_124445689.1 (5e-17/35.83%)<br>WP_124445685.1 (7e-17/39.56%) | WP_124445689.1 (8e-11/34.19%)<br>WP_124445685.1 (6e-08/30.34%) | WP_124445689.1 (2e-13/36.73%)<br>WP_124445685.1 (2e-11/33.70%) | NZ_BHXA00000000,<br>292 contigs, Illumina HiSeq2500 | Potential abyssomicin BGC, partial |

| | | | | | |
|---|---|---|---|---|---|
| *Streptomyces* sp. NRRL F-525 | WP_033287247.1 (1e-20/39.50%) | WP_033287156.1 (6e-09/29.56%) | WP_033287247.1 (4e-12/34.78%) | NZ_JNXE00000000,<br>242 contigs, Illumina | Potential abyssomicin BGC, partial |
| | WP_033287156.1 (6e-09/33.33%) | | WP_033287156.1 (2e-07/29.21% ) | | |
| *Streptomyces* sp. NRRL F-5126 | WP_030904231.1 (3e-12/35.11%) | No hit | No hit | NZ_JOFZ00000000,<br>168 contigs, Illumina | Potential abyssomicin BGC, partial |
| *Streptomyces* sp. NRRL F-5755 | WP_053700241.1 (2e-12/36.70%) | No hit | WP_053700241.1 (2e-22/36.59%) | NZ_LGCW00000000,<br>327 contigs, Illumina HiSeq | Not enough data |
| *Streptomyces* sp. NRRL S-31 | WP_030750288.1 (3e-35/49.22%) | No hit | WP_030750286.1 (4e-08/29.03%) | NZ_JOCB00000000,<br>322 contigs, Illumina | Not enough data |
| | WP_030750286.1 (1e-15/32.71%) | | | | |
| *Streptomyces* sp. NRRL WC-3742 | WP_031075100.1 (3e-09/33.33%) | WP_031075100.1 (6e-45/57.48% ) | No hit | NZ_JOCF01000000,<br>233 contigs, Illumina | Potential abyssomicin BGC, partial |
| *Streptomyces olindensis* DAUFPE 5622 | KDN76177.1 (2e-11/31.15%) | KDN76177.1 (3e-06/28.30%) | No hit | JJOH00000000,<br>233 contigs, Illumina | Potential tetronomycin BGC, total |
| *Amycolatopsis orientalis* Q427-8 | AFI57012.1 (4e-09/36.75%) | No hit | AFI57012.1 (6e-06/30.82%) | JF970188,<br>Quartromicin BGC, no data | Quartromicin BGC, total |
| *Pantoea* sp. A4 | WP_017346023.1 (1e-08/31.18%) | No hit | WP_017346023.1 (9e-11/28.87%) | NZ_ALXE00000000,<br>71 contigs, Illumina HiSeq | Potential BGC, partial |
| *Streptomyces paucisporeus* CGMCC 4.2025 | WP_073498104.1 (8e-13/33.33%) | WP_073498104.1 (6e-11/29.17%) | WP_073498104.1 (4e-08/28.71%) | NZ_FRBI00000000,<br>79 contigs, no data | Potential abyssomicin BGC, partial |
| *Actinomadura pelletieri* DSM 43383 | WP_121438130.1 (2e-16/40.00% ) | WP_121438130.1 (1e-07/25.83%) | WP_121438130.1 (3e-09/29.66%) | NZ_RBWU00000000,<br>15 contigs, Illumina HiSeq | Potential chlorothricin BGC, partial |
| *Candidatus Streptomyces philanthi* LHW51701 | WP_114021297.1 (4e-06/29.09%) | No hit | WP_114021297.1 (6e-14/33.83%) | NZ_QOIN00000000,<br>80 contigs, Illumina MiSeq | Potential BGC, partial |
| *Streptomyces rimosus subsp. rimosus* NRRL B-16073 | WP_033030402.1 (2e-10/38.14%) | No hit | WP_033030402.1 (8e-23/34.76%) | NZ_JNWX00000000,<br>140 contigs, Illumina | Potential BGC, partial |
| *Streptomyces* sp. Amel2xE9 | WP_027758722.1 (1e-11/39.76%) | WP_027758722.1 (6e-91/98.44%) | WP_027758722.1 (3e-15/32.08%) | NZ_ARPE00000000,<br>73 contigs, no data | Abyssomicin BGC, partial |
| *Streptomyces* sp. e14 | WP_106434019.1 (1e-11/39.76% ) | WP_106434019.1 (1e-125/100.00%) | WP_106434019.1 (4e-15/31.34%) | NZ_ACUR01000000,<br>13 scaffolds, ABI | Abyssomicin BGC, partial |
| *Streptomyces fragilis* NBRC 12862 | WP_108952931.1 (5e-65/78.46%) | WP_108952931.1 (2e-10/36.73%) | WP_108952931.1 (2e-16/38.64%) | NZ_BEVZ00000000,<br>19 contigs, Illumina MiSeq | Abyssomicin BGC, partial |
| *Streptomyces incarnatus* NRRL 8089 | AKJ08822.1 (1e-12/33.91%) | AKJ08822.1 (4e-52/60.58%) | AKJ08822.1 (3e-13/33.08%) | CP011497,<br>Complete genome, 454 | Potential abyssomicin BGC, partial |
| *Streptomyces* sp. NRRL F-6491 | WP_053650217.1 (3e-10/36.14%) | WP_053650217.1 (1e-115/91.81%) | WP_053650217.1 (7e-17/32.84%) | NZ_LGEE00000000,<br>287 contigs, Illumina HiSeq | Abyssomicin BGC, partial |
| *Streptomyces olivaceus* KLBMP 5084 | WP_037772597.1 (3e-21/34.55% ) | WP_037772597.1 (1e-14/36.90%) | WP_037772597.1 (3e-12/32.35%) | NZ_CP016795,<br>Complete genome, PacBio | Potential BGC, total |
| *Streptomyces regalis* NRRL 3151 | WP_062712130.1 (8e-66/78.46%) | WP_062712130.1 (9e-09/34.94%) | WP_062712130.1 (7e-16/38.64%) | NZ_LLZG00000000,<br>424 contigs, Illumina HiSeq | Potential abyssomicin BGC, partial |
| *Streptomyces regensis* NRRL B-11479 | KMS84434.1 (1e-63/76.92%) | KMS84434.1 (1e-08/37.00% ) | KMS84434.1 (2e-17/40.91% ) | LFVR00000000,<br>960 contigs, Illumina HiSeq | Potential abyssomicin BGC, partial |
| *Saccharothrix syringae* NRRL B-16468 | WP_033431227.1 (4e-23/34.13%) | WP_033434419.1 (4e-49/59.38%) | WP_033431227.1 (3e-13/33.71%) | NZ_JNYO00000000,<br>190 contigs, Illumina | Abyssomicin BGC, partial |
| | WP_033434419.1 (5e- | WP_033431227.1 (8e- | WP_033434419.1 (2e- | | Potential BGC, total |

| | | | | | |
|---|---|---|---|---|---|
| | 11/32.17%) | 16/39.29%) | 11/33.33%) | | |
| *Streptomyces* sp. SCA2-2 | WP_129847681.1 (2e-15/33.93%) | WP_129847681.1 (4e-15/31.34%) | WP_129847681.1 (9e-158/99.09%) | NZ_PKMX00000000, 48 contigs, Illumina HiSeq | Abyssomicin BGC, partial |
| *Streptomyces* sp. SolWspMP-5a-2 | WP_093830612.1 (2e-13/36.17%) | WP_093830612.1 (8e-06/31.00%) | WP_093830612.1 (4e-06/28.09%) | NZ_FMCI00000000, 400 contigs, no data | Not enough data |
| *Streptacidiphilus* sp. DSM 106435 | WP_111492780.1 (9e-07/32.46%) | WP_111492780.1 (3e-47/60.16% ) | WP_111492780.1 (4e-06/29.17%) | NZ_CP031264.1, Complete genome, PacBio RSII and Illumina MiSeq | Potential abyssomicin BGC, partial |
| *Streptomyces hoynatensis* KCTC 29097 | WP_120684678.1 (1e-08/31.87%) | WP_120684678.1 (5e-11/29.06%) | WP_120684678.1 (9e-12/30.84% ) | NZ_RBAL00000000.1, 53 contigs, Illumina MiSeq | Not enough data |
| *Streptomyces* sp. 57 | RLK30369.1 (2e-52/71.76%) | No hit | RLK30369.1 (7e-11/38.64%) | RCCZ00000000.1, 8 contigs, PacBio | Abyssomicin BGC, total |
| *Streptomyces subroseum* CGMCC 4.2132 | WP_089206782.1 (2e-08/33.73%) | WP_089206782.1 (9e-75/82.81%) | WP_089206782.1 (8e-12/28.23%) | NZ_FZOD00000000.1, 180 scaffolds, no data | Potential abyssomicin BGC, partial |
| *Streptomyces varsoviensis* NRRL B-3589 | WP_030882074.1 (6e-21/35.45%) | WP_030882074.1 (4e-15/36.90%) | WP_030882074.1 (8e-14/33.33%) | NZ_JOFN00000000.1, 155 contigs, Illumina | Potential BGC, total |
| *Actinocrispum wychmicini* DSM 45934 | - | - | - | NZ_SLWS00000000.1, 43 Scaffolds, Illumina | Potential abyssomicin BGC, partial |
| | WP_132116074.1 (9e-17/42.39%) | No hit | WP_132116074.1 (1e-07/37.31%) | | Potential chlorothricin BGC, partial |

**Table S10.** The abyssomicin biosynthetic gene cluster from *M. maris* AB-16-032 (modified from Gottardi *et al.*, 2011).

| ORF | Size (aa) | Proposed function | Closest homolog, host (protein ID); Identity/Similarity (%) | Abs homolog | Abm homolog |
|---|---|---|---|---|---|
| *abyR* | 248 | transcriptional regulator, SARP family | SARP family transcriptional regulator, *Verrucosispora maris* (WP_013733064.1); 99/100 | - | - |
| *abyX* | 396 | cytochrome P450 | cytochrome P450, *Micromonospora wenchangensis* (WP_088646685.1); 82/90 | AbsX | - |
| *abyH* | 889 | LuxR family transcriptional regulator | helix-turn-helix transcriptional regulator, *Verrucosispora* sp. FIM060022 (WP_126713160.1); 99/99 | - | AbmH |
| *abyI* | 252 | transcriptional regulator, SARP family | putative pathway specific activator, *Streptomyces longisporoflavus* (ACR50789.1); 49/63 | - | AbmI |
| *abyU* | 141 | Diels–Alderase | YD repeat-containing protein*, Verrucosispora* sp. MS100047 (AIS85752.1); 100/100 | AbsU | AbmU |
| *abyK* | 619 | YD repeat | RHS repeat protein, *Verrucosispora sp*. FIM060022 WP_126713159.1] 99/99 | - | - |
| *abyA1* | 341 | β-ketoacyl-acyl-carrier protein synthase I | 3-oxoacyl-ACP synthase III family protein, *Verrucosispora* sp. FIM060022 (WP_126713158.1); 99/99 | AbsA1 | AbmA1 |
| *abyA2* | 622 | phosphatase and glyceryl transferase | FkbH like protein, *Verrucosispora* sp. MS100047 (AIS85751.1); 99/99 | AbsA2 | AbmA2 |
| *abyA3* | 78 | discrete ACP | acyl carrier protein, *Verrucosispora sp*. FIM060022 (RUL90284.1); 99/100 | AbsA3 | AbmA3 |
| *abyA4* | 251 | dehydrogenase catalytic domain-containing protein | dehydrogenase catalytic domain-containing protein, *Verrucosispora* sp. MS100047 (AIS85747.1); 99/99 | AbsA4 | AbmA4 |
| *abyA5* | 355 | hydrolase superfamily dihydrolipoamide acyltransferase-like protein | alpha/beta hydrolase, *Verrucosispora* sp. FIM060022 (WP_126713156.1); 99/99 | AbsA5 | AbmA5 |
| *abyB1* | 5781 | PKS I | type I polyketide synthase, *Streptomyces* sp. KhCrAH-43 (WP_018522876.1); 54/63 | AbsB1 | AbmB1 |
| *abyB2* | 3645 | PKS I | type I polyketide synthase, *Streptomyces* sp. 2131.1 (WP_093709984.1); 54/63 | AbsB2 | AbmB2 |
| *abyB3* | 992 | PKS I | acyltransferase domain-containing protein, *Verrucosispora* sp. FIM060022 (WP_126713147.1); 99/99 | AbsB3 | AbmB3 |
| *abyC* | 230 | regulatory protein, TetR | TetR family transcriptional regulator, *Verrucosispora* sp. FIM060022 (WP_126713146.1); 100/100 | - | AbmC |
| *abyD* | 475 | drug resistance transporter EmrB/QacA | DHA2 family efflux MFS transporter, *Micromonospora wenchangensis* (WP_088644887.1); 86/92 | AbsD | AbmD |
| *abyE* | 335 | luciferase; alkanal monooxygenase α-chain | LLM class flavin-dependent oxidoreductase, *Verrucosispora* sp. FIM060022 (RUL90371.1); 100/100 | AbsE | AbmE1 |
| *abyF1* | 538 | ABC transporter periplasmic peptide-binding protein | ABC transporter substrate-binding protein, *Verrucosispora* sp. FIM060022 (RUL90370.1); 100/100 | AbsF1 | AbmF1 |
| *abyF2* | 311 | ABC transporter inner membrane component | ABC transporter permease, *Micromonospora wenchangensis* (WP_088644890.1); 82/89 | AbsF2 | AbmF2 |
| *abyF3* | 283 | ABC transporter inner membrane component | ABC transporter permease, *Verrucosispora* sp. FIM060022 (WP_126713145.1); 99/99 | AvsF3 | AbmF3 |
| *abyF4* | 539 | ABC transporter ATP-binding protein | dipeptide ABC transporter ATP-binding protein, *Verrucosispora* sp. FIM060022 (WP_126713144.1); 99/99 | AbsF4 | AbmF4 |
| *abyV* | 395 | cytochrome P450 | cytochrome P450, *Verrucosispora* sp. FIM060022 (WP_126713143.1); 99/99 | AbsV | AbmV |
| *abyW* | 83 | alcohol dehydrogenase zinc-binding domain | Ferredoxin, *Verrucosispora* sp. FIM060022 (WP_126713142.1); 98/100 | - | - |
| *abyZ* | 192 | NAD(P)H-dependent FMN reductase | FMN reductase (NADPH), *Verrucosispora* sp. FIM060022 (WP_126713141.1); 99/100 | AbsH1 | AbmZ |
| *abyT* | 298 | thioesterase | Thioesterase, *Verrucosispora* sp. FIM060022 (RUL90363.1); 100/100 | AbsN | AbmT |

**Table S11.** The abyssomicin biosynthetic gene cluster from *S. koyangensis* SCSIO 5802 (modified from Song *et al.*, 2017).

| ORF | Size (aa) | Proposed function | Closest homolog, host (protein ID); Identity/Similarity (%) | Aby homolog | Abs homolog |
|---|---|---|---|---|---|
| *abmN* | 445 | rRNA (Uracil-5-)-methyltransferase | RlmD, *Acinetobacter baumannii* AB307-0294 (B7H018.1); 30/48 | - | - |
| *abmI* | 268 | Transcriptional activator, SARP family | DnrI, *Streptomyces peucetius* (P25047.1); 35/50 | AbyI | - |
| *abmE2* | 356 | Luciferase-like monooxygenase, α-subunit | LuxA, *Photorhabdus luminescens* (P23146.1); 24/42 | - | - |
| *abmU* | 218 | Diels–Alderase | YD repeat-containing protein, *Streptomyces regensis* (KMS84434.1); 41/52 | AbyU | AbsU |
| *abmK* | 256 | 4′-Phosphopantetheinyl transferase superfamily (PPTase) | Npt, *Nocardia iowensis* (A1YCA5.1); 42/54 | - | - |
| *abmL* | 281 | Metallophosphoesterase | GsiA, *Salmonella enterica* (Q57RB2.2); 46/60 | - | - |
| *abmF4* | 560 | ABC transporter system ATP-binding protein | OppD, *Lactococcus lactis* (AIS04392.1); 42/73 | AbyF4 | AbsF4 |
| *abmF3* | 298 | ABC transporter system substrate-binding protein dependent permease | OppC, *Lactococcus lactis* (ABA47380.1); 27/63 | AbyF3 | AbsF3 |
| *abmM* | 413 | Amidohydrolase | Mb2939c, *Mycobacterium bovis* AF2122/97 (P68916.1); 28/37 | - | - |
| *abmF2* | 313 | ABC transporter system permease | OppB*, Lactococcus lactis* (ABA47381.1); 27/58 | AbyF2 | AbsF2 |
| *abmF1* | 546 | ABC transport system substrate-binding protein | OppA, *Lactococcus lactis* (AAO63469.1); 20/52 | AbyF1 | AbsF1 |
| *abmJ* | 331 | Aldo/keto reductase | OsI_15387, *Oryza sativaIndica* (A2XRZ0.1); 50/68 | - | AbsJ |
| *abmG* | 77 | Ferredoxin | Fd-1, *Streptomyces griseolus* (P18324.3); 58/75 | - | AbsG1 |
| *abmV* | 405 | Cytochrome P450 | Vitamin D3 dihydroxylase, *Streptomyces griseolus* (P18326.2); 55/70 | AbyV | AbsV |
| *abmC* | 257 | TetR regulatory protein | Mce3R, *Mycobacterium tuberculosis* H37Rv (P95251.2); 33/48 | AbyC | - |
| *abmE1* | 353 | Luciferase-like monooxygenase, β-subunit | LuxB, *Photorhabdus luminescens* (P19840.1); 20/41 | AbyE | AbsE |
| *abmD* | 487 | Major facilitator superfamily of transporter | EmrB, *Mycobacterium tuberculosis* CDC1551 (P9WG88.1); 34/56 | AbyD | AbsD |
| *abmA1* | 343 | Ketoacyl-*S*-ACP synthase | ChlM, *Streptomyces antibioticus* (AAZ77702.1); 61/74 | AbyA1 | AbsA1 |
| *abmA2* | 628 | Glyceryl-*S*-ACP synthase | ChlD, *Streptomyces antibioticus* (AAZ77703.1); 61/70 | AbyA2 | AbsA2 |
| *abmA3* | 75 | Acyl carrier protein | ChlD2, *Streptomyces antibioticus* (AAZ77704.1); 56/73 | AbyA3 | AbsA3 |
| *abmA4* | 280 | 2-Oxoacid dehydrogenase multienzymes acyltransferase E2 component | ChlD3, *Streptomyces antibioticus* (AAZ77705.1); 65/76 | AbyA4 | AbsA4 |
| *abmA5* | 373 | α/β hydrolase fold protein | ChlD4*, Streptomyces antibioticus* (AAZ77706.1); 51/64 | AbmA5 | AbsA5 |
| *abmT* | 274 | Type II thioesterase | PikA5, *Streptomyces venezuelae* (Q9ZGI1.1); 32/45 | AbyT | AbsN |
| *abmZ* | 178 | NADPH-dependent flavin reductase | HsaB, *Rhodococcus jostii* RHA1 (Q0S808.1); 39/57 | AbyZ | AbsH1 |
| *abmB1* | 6540 | PKS I | PikA1, *Streptomyces venezuelae* (Q9ZGI5.1); 54/65 | AbyB1 | AbsB1 |
| *abmB2* | 4054 | PKS I | PikA2, *Streptomyces venezuelae* (Q9ZGI4.1); 49/59 | AbyB2 | AbsB2 |

| | | | | | |
|---|---|---|---|---|---|
| *abmB3* | 1040 | PKS I | PikA1, *Streptomyces venezuelae* (Q9ZGI5.1); 54/64 | AbyB3 | AbsB3 |
| *abmH* | 942 | LuxR family transcriptional regulator | NreC, *Staphylococcus carnosus subsp. carnosus* TM300 (Q7WZY4.1); 48/67 | AbyH | - |

**Table S12.** The abyssomicin biosynthetic gene cluster from *Streptomyces* sp. LC-6-2 (adapted from Wang *et al.*, 2017).

| ORF | Size (aa) | Proposed function | Closest homolog, host (protein ID); Identity/Similarity (%) | Aby homolog | Abm homolog |
|---|---|---|---|---|---|
| *absM* | 187 | short chain dehydrogenase | short-chain dehydrogenase/reductase family oxidoreductase, *Streptomyces* sp. E14 (EFF94110.1); 97/98 | - | - |
| *absP* | 301 | alpha/beta hydrolase | alpha/beta fold hydrolase, *Streptomyces* sp. E14 (WP_009191652.1); 99/99 | - | - |
| *absK* | 209 | histidine phosphatase family protein | histidine phosphatase family protein, *Streptomyces* sp. E14 (WP_009191653.1); 97/98 | - | - |
| *absC1* | 237 | TetR family transcription regulator | TetR family transcriptional regulator, *Streptomyces* sp. E14 (EFF94113.1); 99/100 | - | - |
| *absH3* | 387 | Oxidoreductase | oxidoreductase, *Streptomyces* sp. E14 (WP_009191655.1); 98/98 | - | - |
| *absH2* | 117 | Oxidoreductase | oxidoreductase, *Streptomyces scabrisporus* (WP_026218602.1); 85/90 | - | - |
| *absH1* | 185 | NADPH-dependent flavin reductase | flavin reductase domain-containing protein, *Streptomyces* sp. E14 (EFF94116.1); 99/100 | AbyZ | AbmZ |
| *absA1* | 351 | oxoacyl-ACP synthase III | 3-oxoacyl-ACP synthase III family protein, *Streptomyces* sp. E14 (WP_063821841.1); 100/100 | AbyA1 | AbmA1 |
| *absA3* | 77 | acyl carrier protein | acyl carrier protein, *Streptomyces* sp. E14 (WP_043261562.1); 100/100 | AbyA3 | AbmA3 |
| *absA4* | 251 | 2-oxoacid dehydrogenases acyltransferase | acyltransferase, *Microbispora triticiradicis* (WP_111700702.1); 82/88 | AbyA4 | AbmA4 |
| *absA5* | 381 | hydrolase superfamily dihydrolipoamide acyltransferase-like protein | alpha/beta hydrolase, *Streptomyces* sp. Amel2xE9 (WP_019984789.1); 98/98 | AbmA5 | AbmA5 |
| *absN* | 275 | thioesterase | thioesterase, *Streptomyces* sp. E14 (WP_009191660.1); 99/99 | AbyT | AbmT |
| *absB1* | 6424 | PKS I | type I polyketide synthase, *Streptomyces* sp. Amel2xE9 (WP_078625043.1); 97/97 | AbyB1 | AbmB1 |
| *absB2* | 3644 | PKS I | type I polyketide synthase, *Streptomyces* sp. Amel2xE9 (WP_019985563.1); 95/95 | AbyB2 | AbmB2 |
| *absB3* | 1049 | PKS I | type I polyketide synthase, *Streptomyces* sp. Amel2xE9 (WP_019985562.1); 98/98 | AbyB3 | AbmB3 |
| *absA2* | 649 | FkbH-like protein | HAD-IIIC family phosphatase, *Streptomyces* sp. Amel2xE9 (WP_019985561.1); 99/99 | AbyA2 | AbmA2 |
| *absE* | 328 | FMN-dependent alkanal monooxygenase | LLM class flavin-dependent oxidoreductase, *Streptomyces* sp. Amel2xE9 (WP_106962181.1); 99/99 | AbyE | AbmE |
| *absF1* | 554 | ABC transporter | ABC transporter substrate-binding protein, *Streptomyces* sp. Amel2xE9 (WP_027758725.1); 99/99 | AbyF1 | AbmF1 |
| *absF2* | 333 | ABC transporter | ABC transporter permease, *Streptomyces* sp. Amel2xE9 (WP_019985558.1); 99/99 | AbyF2 | AbmF2 |
| *absF3* | 269 | ABC transporter | ABC transporter permease, *Streptomyces* sp. Amel2xE9 (WP_106962190.1); 99/99 | AbyF3 | AbmF3 |
| *absF4* | 555 | ABC transporter | ABC transporter ATP-binding protein, *Streptomyces* sp. Amel2xE9 (WP_019985556.1); 98/98 | AbyF4 | AbmF4 |
| *absV* | 397 | cytochrome P450 | cytochrome P450, *Streptomyces* sp. Amel2xE9 (WP_027758724.1); 99/100 | AbyV | AbmV |
| *absG1* | 68 | ferredoxin | ferredoxin-1, *Streptomyces* sp. NRRL F-6491 (KOX15570.1); 84/94 | - | AbmG |
| *absI* | 387 | acyltransferase | acyltransferase, *Streptomyces* sp. E14 (WP_009191675.1); 100/100 | - | - |
| *absX* | 403 | cytochrome P450 | cytochrome P450, *Streptomyces* sp. E14 (WP_009191676.1); 100/100 | AbyX | - |
| *absG2* | 64 | Ferredoxin | ferredoxin-1, *Streptomyces* sp. NRRL F-6491 (KOX15554.1); 89/95 | - | - |

| | | | | | |
|---|---|---|---|---|---|
| *absJ* | 344 | aldo/keto reductase | aldo/keto reductase, *Streptomyces* sp. E14 (WP_050790870.1); 100/100 | - | AbmJ |
| *absU* | 171 | Diels-Alderase | hypothetical protein, *Streptomyces* sp. E14 (WP_106434019.1); 100/100 | AbyU | AbmU |
| *absC2* | 196 | TetR familt transcription regulator | TetR/AcrR family transcriptional regulator, *Streptomyces* sp. E14 (WP_009191680.1); 99/99 | - | - |
| *absD* | 476 | MFS transporter | MFS transporter, *Streptomyces* sp. E14 (WP_043261567.1); 100/100 | AbyD | AbmD |

**Table S13.** The abyssomicin biosynthetic gene cluster from *Verrucosispora* sp. MS100047 (KF826681.1).

| **ORF** | **Size (aa)** | **Proposed function** | **Closest homolog, host (protein ID); Identity/Similarity (%)** | **Abs homolog** | **Abm homolog** |
|---|---|---|---|---|---|
| *VASRM7_503* | 355 | hypothetical protein | AbyA5, *Verrucosispora maris* AB-18-032 (AEK75501.1); 99/99 | AbsA5 | AbmA5 |
| *VASRM7_504* | 251 | dehydrogenase catalytic domain-containing protein | acyltransferase, *Verrucosispora maris* (WP_013733055.1); 99/99 | AbsA4 | AbmA4 |
| *VASRM7_505* | 78 | hypothetical protein | acyl-carrier protein, *Verrucosispora maris* AB-18-032 (AEK75499.1); 100/100 | AbsA3 | AbmA3 |
| *VASRM7_508* | 622 | FkbH like protein | FkbH like protein, *Verrucosispora maris* AB-18-032 (AEB44397.1); 99/99 | AbsA2 | AbmA2 |
| *VASRM7_506* | 341 | 3-oxoacyl-[acyl-carrier-protein | Chain A, 3-oxoacyl-acp Synthase III, *Verrucosispora maris* AB-18-032 (5BY7_A); 99/99 | AbsA1 | AbmA1 |
| *VASRM7_507* | 579 | YD repeat-containing protein | RHS repeat protein, *Verrucosispora* sp. FIM060022 (WP_126713159.1); 99/100 | - | - |
| *VASRM7_509* | 141 | YD repeat-containing protein | YD repeat-containing protein, *Verrucosispora maris* AB-18-032 (AEB44400.1); 100/100 | AbsU | AbmU |
| *VASRM7_510* | 241 | SARP family pathway specific transcriptional activator | SARP family pathway specific transcriptional activator, *Verrucosispora maris* AB-18-032 (AEB44401.1); 100/100 | - | AbmI |
| *VASRM7_511* | 288 | hypothetical protein | - | - | - |

**Table S14.** Predicted functions of ORFs in abyssomicin BGC from *Actinokineospora auranticolor* YU 961-1 (PTIX01000001.1).

| ORF | Size (aa) | Proposed function | Closest homolog, host (protein ID); Identity/Similarity (%) | Aby homolog | Abs homolog | Abm homolog |
|---|---|---|---|---|---|---|
| *CLV40_RS02920* | 314 | 3-oxoacyl-[acyl-carrier-protein] synthase-3 | 3-oxoacyl-ACP synthase III family protein, *Microbispora triticiradicis* (WP_111700704.1); 76/84 | AbyA1 | AbsA1 | AbmA1 |
| *CLV40_RS02925* | 74 | phosphopantetheine binding protein | acyl carrier protein, *Actinocrispum wychmicini* (WP_132114012.1); 58/75 | AbyA3 | AbsA3 | AbmA3 |
| *CLV40_RS02930* | 247 | 2-oxoacid dehydrogenase/acyltransferase | acyltransferase, *Microbispora triticiradicis* (WP_111700702.1); 70/77 | AbyA4 | AbsA4 | AbmA4 |
| *CLV40_RS02935* | 357 | alpha/beta hydrolase family protein DUF1100 | alpha/beta hydrolase, *Microbispora triticiradicis* (WP_111700701.1); 66/78 | AbyA5 | AbsA5 | AbmA5 |
| *CLV40_RS02940* | 237 | surfactin synthase thioesterase subunit | thioesterase, *Microbispora triticiradicis* (WP_111700742.1); 55/64 | AbyT | AbsN | AbmT |
| *CLV40_RS02945* | 5456 | PKS I | putative type I polyketide synthase, *Frankia alni* ACN14a (CAJ62714.1); 53/62 | AbyB1 | AbsB1 | AbmB1 |
| *CLV40_RS02950* | 3217 | PKS I | type I polyketide synthase, *Frankia* sp. AvcI1 (WP_055752030.1); 58/67 | AbyB2 | AbsB2 | AbmB2 |
| *CLV40_RS02955* | 989 | PKS I | acyltransferase domain-containing protein, *Frankia alni* (WP_011605200.1); 58/67 | AbyB3 | AbsB3 | AbmB3 |
| *CLV40_RS02960* | 629 | HAD superfamily phosphatase (TIGR01681 family)/FkbH-like protein | HAD-IIIC family phosphatase, *Frankia alni* (WP_011605199.1); 63/72 | AbyA2 | AbsA2 | AbmA2 |
| *CLV40_RS02965* | 161 | DDE superfamily endonuclease | transposase IS4 family protein, *Saccharomonospora azurea* SZMC 14600 (EHK83939.1); 51/67 | - | - | - |
| *CLV40_RS02970* | 332 | luciferase family oxidoreductase group 1 | LLM class flavin-dependent oxidoreductase, *Streptomyces* sp. Amel2xE9 (WP_106962181.1);68/76 | AbyE | AbsE1 | AbmE1 |
| *CLV40_RS02975* | 542 | peptide/nickel transport system substrate-binding protein | ABC transporter substrate-binding protein, *Microbispora triticiradicis* (WP_117408852.1); 62/73 | AbyF1 | AbsF1 | AbmF1 |
| *CLV40_RS02980* | 278 | peptide/nickel transport system permease protein | ABC transporter permease, *Streptosporangium subroseum* (WP_089206641.1); 68/80 | AbyF2 | AbsF2 | AbmF2 |
| *CLV40_RS02985* | 270 | peptide/nickel transport system permease protein | ABC transporter permease, *Actinocorallia herbida* (WP_123665082.1); 61/72 | AbyF3 | AbsF3 | AbmF3 |
| *CLV40_RS02990* | 535 | peptide/nickel transport system ATP-binding protein | ABC transporter ATP-binding protein, *Streptosporangium subroseum* (WP_089206779.1); 66/74 | AbyF4 | AbsF4 | AbmF4 |
| *CLV40_RS02995* | 396 | pentalenic acid synthase | cytochrome P450, *Microbispora triticiradicis* (WP_117409456.1); 74/85 | AbyV | AbsV | AbmV |
| *CLV40_RS03000* | 63 | ferredoxin | ferredoxin, *Streptosporangium subroseum* (WP_089206642.1); 61/76 | - | AbsG1 | AbmG |
| *CLV40_RS03005* | 333 | aryl-alcohol dehydrogenase-like predicted oxidoreductase | aldo/keto reductase, *Microbispora triticiradicis* (WP_117409459.1); 69/78 | - | AbsJ | AbmJ |
| *CLV40_RS03010* | 131 | Diels-Alderase | hypothetical protein, *Streptomyces* sp. E14 (WP_106434019.1); 68/77 | AbyU | AbsU | AbmU |
| *CLV40_RS03015* | 915 | regulatory LuxR family protein | LuxR family transcriptional regulator, *Microbispora triticiradicis* (WP_133306130.1); 55/67 | AbyH | - | AbmH |
| *CLV40_RS03020* | 213 | AcrR family transcriptional regulator | TetR/AcrR family transcriptional regulator, *Microbispora* sp. GKU 823 (WP_079317081.10; 73/84 | - | AbsC2 | - |
| *CLV40_RS03025* | 480 | EmrB/QacA drug resistance transporter | MFS transporter, *Saccharothrix syringae* (WP_033434362.1);62/75 | AbyD | AbsD | AbmD |

| | | | | | | |
|---|---|---|---|---|---|---|
| *CLV40_RS03030* | 277 | SARP family transcriptional regulator | SARP family transcriptional regulator, *Micromonospora wenchangensis* (WP_088646684.1); 74/82 | AbyI | - | AbmI |

**Table S15.** Predicted functions of ORFs in abyssomicin BGC from *Actinokineospora auranticolor* YU 961-1 (PTIX01000011.1).

| ORF | Size (aa) | Proposed function | Closest homolog, host (protein ID); Identity/Similarity (%) | Aby homolog | Abs homolog | Abm homolog |
|---|---|---|---|---|---|---|
| *CLV40_RS20825* | 402 | cytochrome P450 | cytochrome P450, *Frankia* sp. Cc1.17 (WP_071083429.1); 65/77 | AbyV | AbsV | AbmV |
| *CLV40_RS20830* | 171 | Diels-Alderase | hypothetical protein, *Streptomyces* sp. NRRL S-31 (WP_030750286.1); 53/64 | AbyU | AbsU | AbmU |
| *CLV40_RS20835* | 155 | Diels-Alderase | hypothetical protein, *Streptomyces* sp. NRRL S-31 (WP_030750288.1); 60/76 | AbyU | AbsU | AbmU |
| *CLV40_RS20840* | 343 | 3-oxoacyl-ACP synthase III family protein | 3-oxoacyl-ACP synthase III family protein, *Streptomyces* sp. NRRL S-31 (WP_030750290.1); 70/83 | AbyA1 | AbsA1 | AbmA1 |
| *CLV40_RS20845* | 631 | HAD-IIIC family phosphatase | HAD-IIIC family phosphatase, *Kutzneria buriramensis* (WP_116181645.1); 61/73 | AbyA2 | AbsA2 | AbmA2 |
| *CLV40_RS20850* | 75 | acyl carrier protein | acyl carrier protein, *Streptomyces* sp. NRRL F-5123 (WP_031525362.10; 54/79 | AbyA3 | AbsA3 | AbmA3 |
| *CLV40_RS20855* | 230 | acyltransferase | acyltransferase, *Actinomadura pelletieri* (WP_121438112.1); 63/73 | AbyA4 | AbsA4 | AbmA4 |
| *CLV40_RS20860* | 358 | alpha/beta hydrolase | alpha/beta hydrolase, *Streptomyces* sp. 2131.1 (WP_093709996.1); 59/70 | AbyA5 | AbsA5 | AbmA5 |
| *CLV40_RS20865* | 6037 | PKS I | type I polyketide synthase, *Streptomyces fragilis* (WP_108952947.1); 54/62 | AbyB1 | AbsB1 | AbmB1 |
| *CLV40_RS20870* | 3369 | PKS I | type I polyketide synthase, *Actinomadura macra* (WP_067456430.1); 49/58 | AbyB2 | AbsB2 | AbmB2 |
| *CLV40_RS20875* | 1036 | PKS I | type I polyketide synthase, *Streptomyces regalis* (WP_062710596.1); 53/64 | AbyB3 | AbsB3 | AbmB3 |
| *CLV40_RS20880* | 347 | LLM class flavin-dependent oxidoreductase | LLM class flavin-dependent oxidoreductase, *Frankia* sp. Cc1.17 (WP_071083425.1); 68/80 | - | - | AbmE2 |
| *CLV40_RS20885* | 532 | MFS transporter | MFS transporter, *Streptomyces regalis* (KUL37079.10); 48/63 | AbyD | AbsD | AbmD |
| *CLV40_RS20890* | 407 | amidohydrolase family protein | amidohydrolase family protein, *Streptomyces* sp. SCA2-2 (WP_129847676.1); 53/64 | - | - | AbmM |
| *CLV40_RS20895* | 350 | LLM class flavin-dependent oxidoreductase | LLM class flavin-dependent oxidoreductase, *Frankia* sp. Cc1.17 (WP_071083424.1);60/71 | AbyE | AbsE1 | AbmE1 |
| *CLV40_RS20900* | 225 | TetR/AcrR family transcriptional regulator | TetR/AcrR family transcriptional regulator, *Streptomyces* sp. CB02414 (WP_073730525.1); 56/70 | AbyC | - | AbmC |
| *CLV40_RS20905* | 483 | DHA2 family efflux MFS transporter permease subunit | DHA2 family efflux MFS transporter permease subunit, *Frankia* sp. Cc1.17 (WP_071083423.1); 67/78 | AbyD | AbsD | AbmD |
| *CLV40_RS20910* | 280 | AfsR/SARP family transcriptional regulator | AfsR/SARP family transcriptional regulator, *Streptomyces* sp. SA15 (WP_095750847.10; 52/69 | AbyI | - | AbmI |
| *CLV40_RS20915* | 885 | helix-turn-helix transcriptional regulator | hypothetical protein AMK11_25530, *Streptomyces* sp. CB02414 (OKI81332.1); 33/42 | AbyH | - | AbmH |
| *CLV40_RS20920* | 526 | ABC transporter ATP-binding protein | dipeptide ABC transporter ATP-binding protein, *Actinocrispum wychmicini* (WP_132114000.1); 61/71 | AbyF4 | AbsF4 | AbmF4 |
| *CLV40_RS20925* | 282 | ABC transporter permease | ABC transporter permease, *Actinomadura macra* (WP_067456459.1); 68/79 | AbyF3 | AbsF3 | AbmF3 |
| *CLV40_RS20930* | 288 | ABC transporter permease | ABC transporter permease subunit, *Actinocrispum wychmicini* (WP_132114004.1); 64/81 | AbyF2 | AbsF2 | AbmF2 |
| *CLV40_RS20935* | 531 | ABC transporter substrate-binding protein | AbmF1, *Streptomyces koyangensis* (AVI57419.1); 54/67 | AbyF1 | AbsF1 | AbmF1 |

**Table S16.** Predicted functions of ORFs surrounding AbyU homolog from *Actinomadura sp.* H3C3 (NZ_SMKU01000406.1).

| **ORF** | **Size (aa)** | **Proposed function** | **Closest homolog, host (protein ID); Identity/Similarity (%)** | **Aby homolog** | **Abs homolog** | **Abm homolog** |
|---|---|---|---|---|---|---|
| *E1298_RS41175* | - | PKS I | type I polyketide synthase, *Streptomyces iranensis* (WP_044580929.1); 53/61 | - | - | - |
| *E1298_RS41180* | 209 | TetR family transcriptional regulator | TetR family transcriptional regulator, *Streptomyces* sp. NRRL F-525 (WP_078652913.1); 45/67 | - | - | - |
| *E1298_RS41185* | 722 | MMPL family transporter | MMPL family transporter, *Streptomyces* sp. NRRL F-525 (WP_051801844.1); 60/70 | - | - | - |
| *E1298_RS41190* | 181 | Diels-Alderase | hypothetical protein, *Actinomadura* sp. 6K520 (WP_131984851.1); 31/51 | AbyU | AbsU | AbmU |

**Table S17.** Predicted functions of ORFs in quartromicin BGC from *Amycolatopsis albispora* WP1 (NZ_CP015163.1).

| ORF | Size (aa) | Proposed function | Closest homolog, host (protein ID); Identity/Similarity (%) | Aby homolog | Abs homolog | Abm homolog |
|---|---|---|---|---|---|---|
| *A4R43_RS38100* | 399 | cytochrome P450 | cytochrome p450, *Amycolatopsis orientalis* (AFI57027.1); 95/97 | - | - | - |
| *A4R43_RS38105* | 267 | acyltransferase | 2-oxoacid dehydrogenase, acyltransferase, *Amycolatopsis orientalis* (AFI57026.1); 86/90 | - | - | - |
| *A4R43_RS38110* | 74 | acyl carrier protein | ACP, *Amycolatopsis orientalis* (AFI57025.1); 94/97 | - | - | - |
| *A4R43_RS38115* | 609 | HAD-IIIC family phosphatase | glyceryltransferase/phosphatase, *Amycolatopsis orientalis* (AFI57024.1); 91/94 | - | - | - |
| *A4R43_RS38120* | 343 | 3-oxoacyl-ACP synthase III family protein | 3-oxoacyl-ACP synthase III (KS), *Amycolatopsis orientalis* (AFI57023.1); 95/98 | - | - | - |
| *A4R43_RS38125* | 221 | response regulator transcription factor | QmnRg3, *Amycolatopsis orientalis* (AFI57015.1); 99/99 | - | - | - |
| *A4R43_RS38130* | 446 | HAMP domain-containing protein | QmnRg2, *Amycolatopsis orientalis* (AFI57016.1); 90/94 | - | - | - |
| *A4R43_RS38135* | 150 | hypothetical protein | QmnL, *Amycolatopsis orientalis* (AFI57019.1); 88/90 | - | - | - |
| *A4R43_RS38140* | 352 | HlyD family efflux transporter periplasmic adaptor subunit | QmnK, *Amycolatopsis orientalis* (AFI57020.1); 96/97 | - | - | - |
| *A4R43_RS38145* | 227 | ABC transporter ATP-binding protein | QmnRs2, *Amycolatopsis orientalis* (AFI57021.1); 96/99 | - | - | - |
| *A4R43_RS38150* | 406 | ABC transporter permease | QmnRs1, *Amycolatopsis orientalis* (AFI57022.1); 97/98 | - | - | - |
| *A4R43_RS38155* | 160 | hypothetical protein | QmnJ, *Amycolatopsis orientalis* (AFI57014.1); 88/93 | - | - | - |
| *A4R43_RS38160* | 348 | HlyD family efflux transporter periplasmic adaptor subunit | QmnI, *Amycolatopsis orientalis* (AFI57013.1); 88/93 | - | - | - |
| *A4R43_RS38165* | 376 | Diels-Alderase | QmnH, *Amycolatopsis orientalis* (AFI57012.1); 94/97 | AbyU | AbsU | AbmU |
| *A4R43_RS38170* | 533 | hypothetical protein | PQQ-dependent dehydrogenase, *Amycolatopsis orientalis* (AFI57011.1); 93/95 | - | - | - |
| *A4R43_RS38175* | 72 | hypothetical protein | hypothetical protein, *Streptomyces sparsogenes* (WP_065968263.1); 57/63 | - | - | - |
| *A4R43_RS38180* | 255 | AfsR/SARP family transcriptional regulator | regulator, *Amycolatopsis orientalis* (AFI57010.1); 98/98 | - | - | - |
| *A4R43_RS38185* | 396 | cysteine desulfurase-like protein | QmnF, *Amycolatopsis orientalis* (AFI57009.1); 89/92 | - | - | - |
| *A4R43_RS38190* | 383 | hypothetical protein | QmnE, *Amycolatopsis orientalis* (AFI57008.1); 90/93 | - | - | - |
| *A4R43_RS38195* | 153 | hypothetical protein | - | - | - | - |
| *A4R43_RS38200* | 1285 | PKS I | QmnA3, *Amycolatopsis orientalis* (AFI57007.1); 86/90 | - | - | - |
| *A4R43_RS38205* | 1767 | PKS I | QmnA2, *Amycolatopsis orientalis* (AFI57006.1); 93/96 | - | - | - |
| *A4R43_RS38210* | - | PKS I | QmnA1, *Amycolatopsis orientalis* (AFI57005.1); 92/94 | - | - | - |
| *A4R43_RS38215* | - | PKS I | QmnA1, *Amycolatopsis orientalis* (AFI57005.1); 89/94 | - | - | - |

| | | | | | | |
|---|---|---|---|---|---|---|
| *A4R43_RS38220* | - | PKS I | QmnA1, *Amycolatopsis orientalis* (AFI57005.1); 89/93 | - | - | - |
| *A4R43_RS38225* | - | PKS I | QmnA1, *Amycolatopsis orientalis* (AFI57005.1); 92/94 | - | - | - |
| *A4R43_RS38230* | - | PKS I | QmnA1, *Amycolatopsis orientalis* (AFI57005.1); 86/92 | - | - | - |
| *A4R43_RS38235* | - | PKS I | QmnA1, *Amycolatopsis orientalis* (AFI57005.1); 91/95 | - | - | - |
| *A4R43_RS38240* | - | PKS I | QmnA1, *Amycolatopsis orientalis* (AFI57005.1); 94/96 | - | - | - |
| *A4R43_RS38245* | - | PKS I | QmnA1, *Amycolatopsis orientalis* (AFI57005.1); 91/93 | - | - | - |
| *A4R43_RS38250* | - | PKS I | QmnA1, *Amycolatopsis orientalis* (AFI57005.1); 86/90 | - | - | - |
| *A4R43_RS38255* | - | PKS I | QmnA1, *Amycolatopsis orientalis* (AFI57005.1); 86/91 | - | - | - |
| *A4R43_RS38260* | - | PKS I | QmnA1, *Amycolatopsis orientalis* (AFI57005.1); 90/93 | - | - | - |
| *A4R43_RS38265* | 322 | alpha/beta hydrolase | 2-oxoacid dehydrogenase, acyltransferase, *Amycolatopsis orientalis* (AFI57004.1); 90/93 | - | - | - |
| *A4R43_RS38270* | 251 | thioesterase | thioesterase, *Amycolatopsis orientalis* (AFI57003.1); 95/98 | - | - | - |
| *A4R43_RS38275* | 471 | acyl-CoA carboxylase subunit beta | propionyl-CoA carboxylase, *Amycolatopsis orientalis* (AFI57002.1); 97/98 | - | - | - |
| *A4R43_RS38280* | 304 | PAC2 family protein | hypothetical protein, *Amycolatopsis orientalis* (AFI57001.1); 97/97 | - | - | - |
| *A4R43_RS38285* | 247 | SDR family NAD(P)-dependent oxidoreductase | short chain dehydrogenase, *Amycolatopsis orientalis* (AFI56999.1); 97/98 | - | - | - |
| *A4R43_RS38290* | 243 | FadR family transcriptional regulator | regulatory protein, *Amycolatopsis orientalis* (AFI56998.1); 89/93 | - | - | - |

**Table S18.** Predicted functions of ORFs surrounding AbyU homolog from *Streptomyces armeniacus* ATCC 15676 (CP031320.1)

| ORF | Size (aa) | Proposed function | Closest homolog, host (protein ID); Identity/Similarity (%) | Aby homolog | Abs homolog | Abm homolog |
|---|---|---|---|---|---|---|
| *DVA86_31825* | 380 | LacI family transcriptional regulator | LacI family transcriptional regulator, *Streptomyces* sp. CNH287 (WP_037760459.1); 81/84 | - | - | - |
| *DVA86_31830* | 326 | sugar ABC transporter permease | sugar ABC transporter permease, *Streptomyces* sp. CNH287 (WP_027750470.1); 89/92 | - | - | - |
| *DVA86_31835* | 300 | carbohydrate ABC transporter permease | carbohydrate ABC transporter permease, *Streptomyces* sp. CNH287 (WP_051262793.1); 88/93 | - | - | - |
| *DVA86_31840* | 444 | extracellular solute-binding protein | extracellular solute-binding protein, *Streptomyces* sp. Z26 (WP_121516363.1); 82/89 | - | - | - |
| *DVA86_31845* | 334 | ADP-ribosylglycohydrolase family protein | crystallin, *Streptomyces rimosus* (WP_033027683.1); 87/91 | - | - | - |
| *DVA86_31855* | 1954 | polyketide synthase | ChlA1, *Streptomyces antibioticus* (AAZ77693.1); 61/71 | - | - | - |
| *DVA86_31860* | 182 | Diels-Alderase | hypothetical protein, *Actinocrispum wychmicini* (WP_132116074.1); 41/52 | AbyU | AbsU | AbmU |
| *DVA86_31865* | 262 | SDR family NAD(P)-dependent oxidoreductase | SDR family oxidoreductase, *Pseudonocardia acaciae* (WP_028921087.1); 66/76 | - | - | - |
| *DVA86_31870* | 433 | class I SAM-dependent methyltransferase | class I SAM-dependent methyltransferase, *Streptomyces aureocirculatus* (WP_030566749.1); 73/83 | - | - | - |
| *DVA86_31875* | 258 | NAD-dependent epimerase/dehydratase family protein | ChlC5, *Streptomyces antibioticus* (AAZ77680.1); 53/68 | - | - | - |
| *DVA86_31880* | 404 | DUF1205 domain-containing protein | DUF1205 domain-containing protein, *Micromonospora* sp. HK10 (WP_046564311.1); 55/66 | - | - | - |
| *DVA86_31885* | 415 | DUF1205 domain-containing protein | DUF1205 domain-containing protein, *Micromonospora* sp. HK10 (WP_046564311.1); 49/64 | - | - | - |

**Table S19.** Predicted functions of ORFs surrounding AbyU homolog from *Amycolatopsis sp.* CA-126428 (NZ_PPHF01000036).

| **ORF** | **Size (aa)** | **Proposed function** | **Closest homolog, host (protein ID); Identity/Similarity (%)** | **Aby homolog** | **Abs homolog** | **Abm homolog** |
|---|---|---|---|---|---|---|
| *C2L58_RS10660* | 1398 | PKS I | type I polyketide synthase, *Actinokineospora inagensis* (WP_084467521.1); 65/73 | - | - | - |
| *C2L58_RS10665* | 171 | Diels-Alderase | DHA2 family efflux MFS transporter permease subunit, *Actinokineospora inagensis* (WP_084467520.1); 76/85 | AbyU | AbsU | AbmU |
| *C2L58_RS10670* | 468 | MFS transporter | DHA2 family efflux MFS transporter permease subunit, *Actinokineospora inagensis* (WP_084467520.1); 85/91 | - | - | - |
| *C2L58_RS10675* | 173 | hypothetical protein | hypothetical protein, *Actinokineospora inagensis* (WP_026422675.1); 68/78 | - | - | - |
| *C2L58_RS10680* | - | hypothetical protein | type I polyketide synthase, *Actinokineospora inagensis* (WP_051385696.1); 74/82 | - | - | - |

**Table S20.** Predicted functions of ORFs surrounding AbyU homolog from *Streptomyces caatingaensis* CMAA 1322 (NZ_LFXA01000017).

| ORF | Size (aa) | Proposed function | Closest homolog, host (protein ID); Identity/Similarity (%) | Aby homolog | Abs homolog | Abm homolog |
|---|---|---|---|---|---|---|
| *AC230_RS23505* | 161 | MarR family transcriptional regulator | MarR family transcriptional regulator, *Streptomyces griseocarneus* (WP_121798617.1); 70/79 | - | - | - |
| *AC230_RS23510* | 228 | respiratory nitrate reductase subunit gamma | respiratory nitrate reductase subunit gamma, *Streptomyces* sp. NRRL B-1347 (WP_078868664.1); 73/83 | - | - | - |
| *AC230_RS23515* | 192 | nitrate reductase molybdenum cofactor assembly chaperone | nitrate reductase molybdenum cofactor assembly chaperone, *Streptomyces orinoci* (WP_109280517.1); 71/79 | - | - | - |
| *AC230_RS23520* | 527 | nitrate reductase subunit beta | nitrate reductase subunit beta, *Streptomyces orinoci* (WP_109280530.1); 84/89 | - | - | - |
| *AC230_RS23525* | - | nitrate reductase subunit alpha | nitrate reductase subunit alpha, *Streptomyces orinoci* (WP_109280516.1); 84/89 | - | - | - |
| *AC230_RS23530* | 871 | M4 family peptidase | M4 family peptidase, *Streptomyces olivoreticuli* (WP_116215423.1); 58/69 | - | - | - |
| *AC230_RS23535* | 426 | DegT/DnrJ/EryC1/StrS family aminotransferase | DegT/DnrJ/EryC1/StrS aminotransferase, *Streptomyces rimosus subsp. rimosus* ATCC 10970 (ELQ83841.1); 82/88 | - | - | - |
| *AC230_RS30460* | 335 | hypothetical protein | hypothetical protein, *Streptomyces rimosus* (WP_050503688.1); 68/75 | - | - | - |
| *AC230_RS23545* | 77 | hypothetical protein | acyl carrier protein, *Streptomyces rimosus* (WP_078897713.1); 80/88 | - | - | - |
| *AC230_RS23550* | 551 | (2,3-dihydroxybenzoyl)adenylate synthase | (2,3-dihydroxybenzoyl)adenylate synthase, *Streptomyces rimosus* (WP_053801678.1); 74/79 | - | - | - |
| *AC230_RS23555* | 263 | thioesterase | thioesterase, *Streptomyces rimosus* (WP_030371682.1); 62/71 | - | - | - |
| *AC230_RS23560* | 397 | FAD-dependent oxidoreductase | FAD-dependent oxidoreductase, *Streptomyces rimosus* (WP_030670919.1); 81/88 | - | - | - |
| *AC230_RS23565* | 966 | type I polyketide synthase | type I polyketide synthase, *Streptomyces rimosus* (WP_050508728.1); 74/80 | - | - | - |
| *AC230_RS23570* | 171 | CGNR zinc finger domain-containing protein | CGNR zinc finger domain-containing protein, *Streptomyces* sp. NRRL F-5755 (WP_053700243.1); 72/83 | - | - | - |
| *AC230_RS23575* | 480 | MFS transporter | MFS transporter, *Streptomyces sp*. AM-2504 (WP_131121744.1); 74/83 | - | - | - |
| *AC230_RS23580* | 348 | pyridoxal-phosphate dependent enzyme | pyridoxal-phosphate dependent enzyme, *Streptomyces rimosus* (WP_079027570.1); 60/74 | - | - | - |
| *AC230_RS23585* | 217 | serine acetyltransferase | serine acetyltransferase, *Streptomyces albus* (WP_060729303.1); 68/79 | - | - | - |
| *AC230_RS23590* | 175 | Diels-Alderase | hypothetical protein, *Streptomyces rimosus* (WP_033030402.1); 71/79 | AbyU | AbsU | AbmU |
| *AC230_RS23595* | 435 | tetratricopeptide | hypothetical protein, *Streptomyces rimosus* (WP_003980475.1); 64/73 | - | - | - |
| *AC230_RS23600* | 320 | iron ABC transporter | iron ABC transporter, *Streptomyces rimosus subsp. pseudoverticillatus* (KOT92518.1); 73/81 | - | - | - |
| *AC230_RS23605* | 344 | iron ABC transporter permease | iron ABC transporter permease, *Kribbella* sp. YM53 (WP_131512197.1); 60/76 | - | - | - |
| *AC230_RS23610* | 330 | ABC transporter substrate-binding protein | hypothetical protein, *Streptomyces rimosus* (WP_053801671.1); 60/70 | - | - | - |
| *AC230_RS23620* | 455 | hypothetical protein | hypothetical protein, *Streptomyces mobaraensis* (WP_004952143.1); 70/75 | - | - | - |
| *AC230_RS23625* | 69 | hypothetical protein | - | - | - | - |

| | | | | | | |
|---|---|---|---|---|---|---|
| *AC230_RS23630* | 397 | argininosuccinate synthase | argininosuccinate synthase, *Streptomyces mobaraensis* (WP_004952138.1); 94/97 | - | - | - |
| *AC230_RS23635* | 477 | argininosuccinate | argininosuccinate lyase, *Streptomyces mobaraensis* (WP_004952137.1); 94/95 | - | - | - |
| *AC230_RS23640* | 182 | TetR/AcrR family transcriptional regulator | TetR/AcrR family transcriptional regulator, *Streptomyces orinoci* (WP_109278895.1); 88/91 | - | - | - |
| *AC230_RS23645* | 519 | MFS transporter | MFS transporter, *Streptomyces cinnamoneus* (WP_099197714.1); 85/91 | - | - | - |
| *AC230_RS23650* | 276 | alpha/beta fold hydrolase | alpha/beta fold hydrolase, *Streptomyces luteoverticillatus* (WP_126913376.1); 88/93 | - | - | - |
| *AC230_RS23655* | 225 | 1-acyl-sn-glycerol-3-phosphate acyltransferase | 1-acyl-sn-glycerol-3-phosphate acyltransferase, *Streptomyces mobaraensis* (WP_004955746.1); 93/96 | - | - | - |
| *AC230_RS23660* | 394 | glycerophosphodiester phosphodiesterase | glycerophosphodiester phosphodiesterase, *Streptomyces mobaraensis* (WP_040892776.1); 87/91 | - | - | - |

**Table S21.** Predicted functions of ORFs in potential BGC from *Streptomyces cattleya* DSM 46488 (NC_017586.1).

| ORF | Size (aa) | Proposed function | Closest homolog, host (protein ID); Identity/Similarity (%) | Aby homolog | Abs homolog | Abm homolog |
|---|---|---|---|---|---|---|
| *SCATT_RS00545* | 164 | DUF1360 domain-containing protein | DUF1360 domain-containing protein, *Actinomadura* sp. WAC 06369 (RSN60148.1); 59/69 | - | - | - |
| *SCATT_RS00550* | 466 | dihydrolipoyl dehydrogenase | dihydrolipoyl dehydrogenase, *Streptomyces populi* (WP_103553595.1); 85/93 | - | - | - |
| *SCATT_RS00555* | 90 | hypothetical protein | hypothetical protein, *Streptomyces olivaceus* (WP_031047094.1); 76/82 | - | - | - |
| *SCATT_RS00560* | 393 | LLM class flavin-dependent oxidoreductase | monooxygenase, *Streptomyces olivaceus* (AOW90810.1); 95/98 | - | - | - |
| *SCATT_RS00565* | 277 | SDR family oxidoreductase | SDR family NAD(P)-dependent oxidoreductase, *Streptomyces varsoviensis* (WP_030882095.1); 95/97 | - | - | - |
| *SCATT_RS35820* | 101 | hypothetical protein | hypothetical protein BC342_34615, *Streptomyces olivaceus* (AOW90812.1); 79/78 | - | - | - |
| *SCATT_RS00570* | 454 | crotonyl-CoA carboxylase/reductase | crotonyl-CoA carboxylase/reductas, *Streptomyces olivaceus* (WP_070390081.1); 97/98 | - | - | - |
| *SCATT_RS00575* | 109 | integrase | IS3 family transposase, *Streptomyces* sp. AmelKG-D3 (WP_099217416.1); 86/89 | - | - | - |
| *SCATT_RS00580* | 925 | helix-turn-helix transcriptional regulator | LuxR family transcriptional regulator, *Streptomyces varsoviensis* (WP_030882089.1); 89/93 | - | - | - |
| *SCATT_RS00585* | 75 | hypothetical protein | - | - | - | - |
| *SCATT_RS00590* | 408 | cytochrome P450 | cytochrome P450, *Streptomyces varsoviensis* (WP_078643428.1); 91/95 | - | - | - |
| *SCATT_RS00595* | 339 | thioesterase | thioesterase, *Streptomyces varsoviensis* (WP_078643426.1); 90/94 | - | - | - |
| *SCATT_RS00600* | 158 | nuclear transport factor 2 family protein | nuclear transport factor 2 family protein, *Streptomyces varsoviensis* (WP_030882078.1); 90/94 | - | - | - |
| *SCATT_RS00605* | 280 | NAD(P)-dependent oxidoreductase | NAD(P)-dependent oxidoreductase, *Streptomyces varsoviensis* (WP_030882077.1); 93/97 | - | - | - |
| *SCATT_RS00610* | 140 | Diels-Alderase | hypothetical protein, *Streptomyces varsoviensis* (WP_030882074.1); 96/98 | AbyU | AbsU | AbmU |
| *SCATT_RS00615* | 133 | nuclear transport factor 2 family protein | nuclear transport factor 2 family protein, *Streptomyces olivaceus* (WP_070390078.1); 91/96 | - | - | - |
| *SCATT_RS00620* | 416 | cytochrome P450 | cytochrome P450, *Streptomyces* sp. E5N91 SAI-083 (WP_123627589.1); 93/96 | - | - | - |
| *SCATT_RS00625* | 6125 | PKS I | SDR family NAD(P)-dependent oxidoreductase, *Streptomyces* sp. E5N91 SAI-083 (WP_123627588.1); 92/94 | - | - | - |
| *SCATT_RS00630* | 4349 | PKS I | SDR family NAD(P)-dependent oxidoreductase, *Streptomyces* sp. E5N91 SAI-083 (WP_123627587.1); 90/93 | - | - | - |
| *SCATT_RS00635* | 172 | pyridoxamine 5'-phosphate oxidase family protein | pyridoxamine 5'-phosphate oxidase family protein, *Streptomyces* sp. E5N91 SAI-083 (WP_123627586.1); 90/94 | - | - | - |
| *SCATT_RS00640* | 3931 | PKS I | type I polyketide synthase, *Streptomyces iranensis* (WP_044580009.1); 92/95 | - | - | - |
| *SCATT_RS00645* | 1345 | PKS I | type I polyketide synthase, *Streptomyces iranensis* (WP_044580010.1); 91/94 | - | - | - |

| | | | | | | |
|---|---|---|---|---|---|---|
| *SCATT_RS37820* | 21 | gamma-butyrolactone receptor protein | - | - | - | - |
| *SCATT_RS37825* | 123 | hypothetical protein | - | - | - | - |
| *SCATT_RS00655* | 161 | MBL fold metallo-hydrolase | MBL fold metallo-hydrolase, *Streptomyces viridosporus* (WP_081235511.1); 96/98 | - | - | - |
| *SCATT_RS00660* | 377 | galactose mutarotase | galactose mutarotase, *Streptomyces* sp. Amel2xE9 (WP_037724950.1); 75/82 | - | - | - |
| *SCATT_RS00665* | 202 | TetR/AcrR family transcriptional regulator | TetR/AcrR family transcriptional regulator, *Streptomyces* sp. ADI95-16 (WP_123083105.1); 46/68 | - | - | - |
| *SCATT_RS00675* | 366 | oxidoreductase | oxidoreductase, *Streptomyces hygroscopicus* (WP_030822548.1); 93/95 | - | - | - |
| *SCATT_RS00680* | 59 | hypothetical protein | - | - | - | - |
| *SCATT_RS00685* | 256 | IS5/IS1182 family transposase | IS5/IS1182 family transposase, *Streptomyces* sp. WAC 01438 (AZM62328.1); 96/98 | - | - | - |
| *SCATT_RS00690* | 79 | hypothetical protein | - | - | - | - |
| *SCATT_RS00695* | 416 | hypothetical protein | hypothetical protein, *Kitasatospora mediocidica* (WP_035804847.1); 79/85 | - | - | - |
| *SCATT_RS00700* | 109 | hypothetical protein | TnsA-like heteromeric transposase endonuclease subunit, *Streptomyces* sp. 4121.5 (WP_100841043.1); 77/79 | - | - | - |
| *SCATT_RS00710* | 228 | dipeptidase | dipeptidase, *Streptomyces* sp. CNH287 (WP_027749012.1); 87/92 | - | - | - |
| *SCATT_RS00715* | 317 | NADP-dependent oxidoreductase | NADP-dependent oxidoreductase, *Streptomyces* sp. NRRL S-340 (WP_037861743.1); 86/91 | - | - | - |
| *SCATT_RS00720* | 136 | SDR family NAD(P)-dependent oxidoreductase | SDR family oxidoreductase, *Streptomyces* sp. 840.1 (WP_123531887.1); 86/93 | - | - | - |
| *SCATT_RS00725* | 236 | SDR family oxidoreductase | SDR family oxidoreductase, *Streptomyces* sp. 840.1 (WP_123531928.1); 92/95 | - | - | - |
| *SCATT_RS00730* | 307 | LysR family transcriptional regulator | LysR family transcriptional regulator, *Streptomyces* sp. 840.1 (WP_123531885.1); 92/94 | - | - | - |
| *SCATT_RS00735* | 56 | hypothetical protein | NAD-dependent epimerase/dehydratase family protein, *Streptomyces* sp. H23 (WP_134653989.1); 96/100 | - | - | - |
| *SCATT_RS00740* | 83 | hypothetical protein | NAD-dependent epimerase/dehydratase family protein, *Streptomyces natalensis* (WP_044366192.1); 53/65 | - | - | - |
| *SCATT_RS00745* | 503 | NAD(P)/FAD-dependent oxidoreductase | NAD(P)/FAD-dependent oxidoreductase, *Streptomyces scabrisporus* (WP_020548648.1); 52/65 | - | - | - |
| *SCATT_RS00750* | 194 | CGNR zinc finger domain-containing protein | zf-CGNR multi-domain protein, *Streptomyces* sp. MspMP-M5 (WP_026247789.1); 95/96 | - | - | - |
| *SCATT_RS00755* | 481 | MFS transporter | major facilitator superfamily MFS_1, *Streptomyces iranensis* (CDR01235.1); 96/97 | - | - | - |

**Table S22.** Predicted functions of ORFs in potential abyssomicin BGC from *Streptomyces sp.* CB03911 (NZ_LWLA01000047).

| ORF | Size (aa) | Proposed function | Closest homolog, host (protein ID); Identity/Similarity (%) | Aby homolog | Abs homolog | Abm homolog |
|---|---|---|---|---|---|---|
| *A6A07_RS37465* | 354 | LLM class flavin-dependent oxidoreductase | LLM class flavin-dependent oxidoreductase, *Streptacidiphilus* sp. DSM 106435 (WP_111490410.1); 95/97 | - | - | AbmE2 |
| *A6A07_RS37470* | 1094 | PKS I | acyltransferase domain-containing protein, *Streptacidiphilus* sp. DSM 106435 (WP_111490411.1); 94/95 | AbyB3 | AbsB3 | AbmB3 |
| *A6A07_RS37475* | 2188 | PKS I | SDR family NAD(P)-dependent oxidoreductase, *Streptacidiphilus* sp. DSM 106435 (WP_114914558.1); 91/93 | AbyB2 | AbsB2 | AbmB2 |
| *A6A07_RS37480* | 344 | 3-oxoacyl-ACP synthase III family protein | 3-oxoacyl-ACP synthase III family protein, *Streptacidiphilus* sp. DSM 106435 (WP_111492779.1); 97/97 | AbyA1 | AbsA1 | AbmA1 |
| *A6A07_RS37485* | 126 | Diels-Alderase | hypothetical protein, *Streptacidiphilus* sp. DSM 106435 (WP_111492780.1); 92/96 | AbyU | AbsU | AbmU |
| *A6A07_RS37490* | 437 | cytochrome P450 | cytochrome P450, *Streptacidiphilus* sp. DSM 106435 (WP_111492778.1); 98/99 | AbyX/AbyV | AbsV/AbsX | AbmV |
| *A6A07_RS37495* | 70 | ferredoxin | ferredoxin, *Streptacidiphilus* sp. DSM 106435 (WP_111492777.1); 88/94 | - | AbsG2/AbsG1 | AbmG |
| *A6A07_RS37500* | 395 | acyltransferase | acyltransferase, *Streptacidiphilus* sp. DSM 106435 (WP_111492775.1); 92/94 | - | AbsI | - |
| *A6A07_RS37505* | 331 | aldo/keto reductase | aldo/keto reductase, *Streptacidiphilus* sp. DSM 106435 (WP_111492774.1); 98/98 | - | AbsJ | AbmJ |
| *A6A07_RS37510* | 480 | MFS transporter | MFS transporter, *Streptacidiphilus* sp. DSM 106435 (WP_114914554.1); 97/97 | AbyD | AbsD | AbmD |
| *A6A07_RS37515* | 422 | LLM class flavin-dependent oxidoreductase | LLM class flavin-dependent oxidoreductase, *Kutzneria buriramensis* (WP_116181636.1); 56/67 | - | - | - |
| *A6A07_RS37520* | 353 | LLM class flavin-dependent oxidoreductase | LLM class flavin-dependent oxidoreductase, *Streptomyces* sp. Ru71 (WP_103783148.1); 49/57 | - | - | - |
| *A6A07_RS37525* | 601 | LuxR family transcriptional regulator | LuxR family transcriptional regulator, *Streptacidiphilus* sp. DSM 106435 (WP_114914553.1); 91/93 | AbyH | - | AbmH |
| *A6A07_RS37530* | 94 | acyltransferase | acyltransferase, *Streptacidiphilus* sp. DSM 106435 (WP_111494374.1); 95/97 | AbyA4 | AbsA4 | AbmA4 |
| *A6A07_RS37535* | 370 | alpha/beta hydrolase | alpha/beta hydrolase, *Streptacidiphilus* sp. DSM 106435 (WP_111494372.1); 96/97 | AbyA5 | AbsA5 | AbmA5 |
| *A6A07_RS37540* | 296 | AfsR/SARP family transcriptional regulator | AfsR/SARP family transcriptional regulator, *Streptacidiphilus* sp. DSM 106435 (WP_111494370.1); 96/97 | AbyI | - | AbmI |
| *A6A07_RS37545* | 279 | thioesterase | thioesterase, *Streptacidiphilus* sp. DSM 106435 (WP_111494376.1); 89/91 | AbyT | AbsN | AbmT |
| *A6A07_RS37550* | 6420 | PKS I | SDR family NAD(P)-dependent oxidoreductase, *Streptacidiphilus* sp. DSM 106435 (WP_114914725.1); 90/92 | AbyB1 | AbsB1 | AbmB1 |
| *A6A07_RS37555* | 76 | hypothetical protein | hypothetical protein, *Streptacidiphilus* sp. DSM 106435 (WP_114914550.1); 68/73 | - | - | - |
| *A6A07_RS37560* | 639 | HAD-IIIC family phosphatase | HAD-IIIC family phosphatase, *Streptacidiphilus* sp. DSM 106435 (WP_111488988.1); 94/96 | AbyA2 | AbsA2 | AbmA2 |
| *A6A07_RS37565* | 82 | acyl carrier protein | acyl carrier protein, *Streptacidiphilus* sp. DSM 106435 (WP_111488990.1); 93/95 | AbyA3 | AbsA3 | AbmA3 |
| *A6A07_RS37570* | 180 | NADPH-dependent FMN reductase | NADPH-dependent FMN reductase, *Streptacidiphilus* sp. DSM 106435 (WP_111488992.1); 94/96 | AbyZ | AbsH1 | AbmZ |

**Table S23.** Predicted functions of ORFs in potential BGC from *Streptomyces sp.* E5N91 SAI-083 (NZ_RJKF01000001.1).

| ORF | Size (aa) | Proposed function | Closest homolog, host (protein ID); Identity/Similarity (%) | Aby homolog | Abs homolog | Abm homolog |
|---|---|---|---|---|---|---|
| *EDC84_RS34320* | 235 | SDR family NAD(P)-dependent oxidoreductase | SDR family NAD(P)-dependent oxidoreductase, *Streptomyces* sp. E5N298 (WP_121703707.1); 99/98 | - | - | - |
| *EDC84_RS34325* | 67 | hypothetical protein | conserved hypothetical protein, *Streptomyces lividans* TK24 (EFD71066.1); 99/98 | - | - | - |
| *EDC84_RS34330* | 301 | LysR family transcriptional regulator | LysR family transcriptional regulator, *Streptomyces* sp. M1013 (WP_076972954.1); 99/99 | - | - | - |
| *EDC84_RS34335* | 187 | NAD(P)H-dependent oxidoreductase | NAD(P)H-dependent oxidoreductase, *Streptomyces* sp. CS113 (WP_087805165.1); 98/99 | - | - | - |
| *EDC84_RS34340* | 234 | helix-turn-helix transcriptional regulator | TetR/AcrR family transcriptional regulator, *Streptomyces* sp. M1013 (WP_076972906.1); 97/97 | - | - | - |
| *EDC84_RS34345* | 344 | LacI family transcriptional regulator | LacI family transcriptional regulator, *Streptomyces* sp. M1013 (OMI91275.1); 99/100 | - | - | - |
| *EDC84_RS34350* | 131 | thioredoxin | thioredoxin, *Streptomyces canus* (WP_059296951.1); 99/100 | - | - | - |
| *EDC84_RS34355* | 477 | NAD(P)/FAD-dependent oxidoreductase | NAD(P)/FAD-dependent oxidoreductase, *Streptomyces* sp. E5N298 (WP_121703708.1); 99/99 | - | - | - |
| *EDC84_RS34360* | 218 | peptide deformylase | peptide deformylase, *Streptomyces* sp. E5N298 (WP_121703724.1); 96/98 | - | - | - |
| *EDC84_RS34365* | 115 | tetratricopeptide repeat protein | tetratricopeptide repeat protein, *Streptomyces* sp. S10(2018) (WP_127893298.1); 97/100 | - | - | - |
| *EDC84_RS34370* | 321 | pirin family protein | pirin family protein, *Streptomyces canus* (WP_059296947.1); 98/98 | - | - | - |
| *EDC84_RS34375* | 107 | hypothetical protein | transposase, *Streptomyces viridosporus* ATCC 14672 (EFE71945.1); 64/68 | - | - | - |
| *EDC84_RS34380* | 511 | alpha/beta hydrolase | alpha/beta hydrolase, *Streptomyces parvulus* (WP_114532779.1); 98/98 | - | - | - |
| *EDC84_RS34385* | - | MerR family transcriptional regulator | MerR family transcriptional regulator, *Streptomyces parvulus* (WP_114532776.1); 97/98 | - | - | - |
| *EDC84_RS34390* | - | IS701 family transposase | SRSO17 transposase, *Streptomyces* sp. 75 (REE37585.1); 84/92 | - | - | - |
| *EDC84_RS34395* | 281 | methyltransferase domain-containing protein | methyltransferase domain-containing protein, *Streptomyces olivaceus* (WP_070390074.1); 97/98 | - | - | - |
| *EDC84_RS34400* | 1365 | acyltransferase domain-containing protein | type I polyketide synthase, *Streptomyces olivaceus* (WP_070390075.1); 95/96 | - | - | - |
| *EDC84_RS34405* | 3938 | SDR family NAD(P)-dependent oxidoreductase | type I polyketide synthase, *Streptomyces olivaceus* (WP_031033161.1); 97/97 | - | - | - |
| *EDC84_RS34410* | 117 | pyridoxamine 5'-phosphate oxidase family protein | pyridoxamine 5'-phosphate oxidase family protein, *Streptomyces olivaceus* (WP_031033163.1); 99/99 | - | - | - |
| *EDC84_RS34415* | 4377 | SDR family NAD(P)-dependent oxidoreductase | type I polyketide synthase, *Streptomyces cattleya* (WP_014140914.1); 90/93 | - | - | - |
| *EDC84_RS34420* | 6126 | SDR family NAD(P)-dependent oxidoreductase | type I polyketide synthase, *Streptomyces olivaceus* (WP_070390077.1); 96/97 | - | - | - |
| *EDC84_RS34425* | 420 | cytochrome P450 | cytochrome P450, *Streptomyces olivaceus* (WP_031047118.1); 97/98 | - | - | - |
| *EDC84_RS34430* | 134 | nuclear transport factor 2 family protein | nuclear transport factor 2 family protein, *Streptomyces olivaceus* (WP_070390078.1); 95/97 | - | - | - |

| | | | | | | |
|---|---|---|---|---|---|---|
| *EDC84_RS34435* | 141 | Diels-Alderase | hypothetical protein BC342_34580, *Streptomyces olivaceus* (AOW90806.1); 98/99 | AbyU | AbsU | AbmU |
| *EDC84_RS34440* | 281 | NAD(P)-dependent oxidoreductase | NAD(P)-dependent oxidoreductase, *Streptomyces olivaceus* (WP_031047111.1); 98/98 | - | - | - |
| *EDC84_RS34445* | 158 | nuclear transport factor 2 family protein | nuclear transport factor 2 family protein, *Streptomyces olivaceus* (WP_031047108.1); 97/98 | - | - | - |
| *EDC84_RS34450* | 332 | thioesterase | thioesterase, *Streptomyces varsoviensis* (WP_078643426.1); 85/90 | - | - | - |
| *EDC84_RS34455* | 409 | cytochrome P450 | cytochrome P450, *Streptomyces varsoviensis* (WP_078643428.1); 90/94 | - | - | - |
| *EDC84_RS34460* | 394 | LLM class flavin-dependent oxidoreductase | LLM class flavin-dependent oxidoreductase, *Streptomyces cattleya* (WP_014140896.1); 96/98 | - | - | - |
| *EDC84_RS34465* | 278 | SDR family oxidoreductase | SDR family oxidoreductase, *Streptomyces cattleya* (WP_014140897.1); 94/96 | - | - | - |
| *EDC84_RS34470* | 145 | hypothetical protein | hypothetical protein BC342_34615, *Streptomyces olivaceus* (AOW90812.1); 97/97 | - | - | - |
| *EDC84_RS34475* | 455 | crotonyl-CoA carboxylase/reductase | crotonyl-CoA carboxylase/reductase, *Streptomyces olivaceus* (WP_070390081.1); 99/98 | - | - | - |
| *EDC84_RS34480* | 926 | helix-turn-helix transcriptional regulator | helix-turn-helix transcriptional regulator, *Streptomyces olivaceus* (WP_070387930.1); 93/96 | - | - | - |

**Table S24.** Predicted functions of ORFs in potential BGC from *Micromonospora eburnean* DSM 44814 (NZ_FMHY01000002.1).

| ORF | Size (aa) | Proposed function | Closest homolog, host (protein ID); Identity/Similarity (%) | Aby homolog | Abs homolog | Abm homolog |
|---|---|---|---|---|---|---|
| *GA0070604_RS20550* | 1339 | SDR family NAD(P)-dependent oxidoreductase | type I polyketide synthase, *Streptomyces* sp. RTd22 (WP_079152145.1); 45/57 | - | - | - |
| *GA0070604_RS20555* | 77 | hypothetical protein | acetylhydrolase, *Micromonospora* sp. CB01531 (WP_073835454.1); 71/80 | - | - | - |
| *GA0070604_RS20560* | 253 | hypothetical protein | methyltransferase, *Candidatus Streptomyces philanthi* (WP_114021299.1); 67/81 | - | - | - |
| *GA0070604_RS20565* | 437 | cytochrome P450 | cytochrome P450, *Streptomyces oceani* (WP_070196845.1); 47/62 | - | - | - |
| *GA0070604_RS20570* | 578 | ABC transporter ATP-binding protein | ABC transporter ATP-binding protein, *Streptomyces varsoviensis* (WP_030881385.1); 68/83 | - | - | - |
| *GA0070604_RS20575* | 652 | ABC transporter ATP-binding protein | ABC transporter ATP-binding protein, *Streptomyces griseoplanus* (WP_055589414.1); 70/80 | - | - | - |
| *GA0070604_RS20580* | 412 | DUF1205 domain-containing protein | DUF1205 domain-containing protein, *Streptomyces* sp. LHW50302 (WP_114017411.1); 55/67 | - | - | - |
| *GA0070604_RS20585* | 506 | hypothetical protein | FAD-dependent oxidoreductase, *Streptomyces* sp. LHW50302 (WP_114019176.1); 60/71 | - | - | - |
| *GA0070604_RS20590* | 1759 | type I polyketide synthase | type I polyketide synthase, *Micromonospora haikouensis* (WP_091284722.1); 86/90 | - | - | - |
| *GA0070604_RS20595* | 348 | 3-oxoacyl-ACP synthase | 3-oxoacyl-ACP synthase, *Micromonospora haikouensis* (WP_091284724.1); 83/93 | - | - | - |
| *GA0070604_RS20600* | 403 | cytochrome P450 | cytochrome P450, *Candidatus Streptomyces philanthi* (WP_114021298.1); 77/85 | - | - | - |
| *GA0070604_RS20605* | 200 | Diels-Alderase | hypothetical protein, *Streptomyces* sp. LHW50302 (WP_114017401.1); 73/80 | AbyU | AbsU | AbmU |
| *GA0070604_RS20610* | 1866 | type I polyketide synthase | acyl transferase domain-containing protein, *Actinomadura pelletieri* DSM 43383 (RKS68196.1); 54/65 | - | - | - |
| *GA0070604_RS20615* | 3995 | type I polyketide synthase | ChlA5, *Streptomyces antibioticus* (AAZ77698.1); 57/68 | - | - | - |
| *GA0070604_RS20625* | 923 | hypothetical protein | acyltransferase domain-containing protein, *Candidatus Streptomyces philanthi* (WP_114025062.1); 66/77 | - | - | - |
| *GA0070604_RS20630* | 498 | hypothetical protein | acyltransferase domain-containing protein, *Actinomadura pelletieri* (WP_121438117.1); 46/56 | - | - | - |
| *GA0070604_RS20635* | 160 | hypothetical protein | acyltransferase domain-containing protein, *Streptomyces* LHW50302 (WP_114019184.1); 50/62 | - | - | - |
| *GA0070604_RS20640* | 492 | hypothetical protein | hypothetical protein, *Actinomadura pelletieri* (WP_121438116.1); 60/71 | - | - | - |
| *GA0070604_RS20645* | 344 | 3-oxoacyl-ACP synthase III family protein | 3-oxoacyl-ACP synthase III family protein, *Candidatus Streptomyces philanthi* (WP_114025060.1); 70/82 | - | - | - |
| *GA0070604_RS20650* | 330 | alpha/beta hydrolase | alpha/beta hydrolase, *Streptomyces armeniacus* (AXK32423.1); 64/76 | - | - | - |
| *GA0070604_RS20655* | 258 | AfsR/SARP family transcriptional regulator | AfsR/SARP family transcriptional regulator, *Candidatus Streptomyces philanthi* (WP_114025055.1); 69/82 | - | - | - |
| *GA0070604_RS20660* | 263 | thioesterase | thioesterase, *Actinomadura pelletieri* (WP_121438108.1); 57/72 | - | - | - |

**Table S25.** Predicted functions of ORFs in abyssomicin BGC from *Frankia sp.* AvcI1 (NZ_LJFZ01000030.1 and NZ_LJFZ01000034).

| ORF | Size (aa) | Proposed function | Closest homolog, host (protein ID); Identity/Similarity (%) | Aby homolog | Abs homolog | Abm homolog |
|---|---|---|---|---|---|---|
| UK94_RS25170 | 496 | MFS transporter | MFS transporter, *Frankia alni* (WP_011605223.1); 99/99 | AbyD | AbsD | AbmD |
| UK94_RS25175 | 552 | ABC transporter substrate-binding protein | ABC transporter substrate-binding protein, *Frankia alni* (WP_011605219.1); 98/98 | AbyF1 | AbsF1 | AbmF1 |
| UK94_RS25180 | 306 | ABC transporter permease | ABC transporter permease, *Frankia alni* ACN14a (CAJ62730.1); 99/99 | AbyF2 | AbsF2 | AbmF2 |
| UK94_RS25185 | 262 | ABC transporter permease | ABC transporter permease, *Frankia alni* (WP_011605217.1); 98/98 | AbyF3 | AbsF3 | AbmF3 |
| UK94_RS25190 | 582 | ABC transporter ATP-binding protein | ABC transporter ATP-binding protein, *Frankia alni* (WP_050997164.1); 99/99 | AbyF4 | AbsF4 | AbmF4 |
| UK94_RS25195 | 396 | cytochrome P450 | cytochrome P450, *Frankia alni* (WP_011605215.1); 98/98 | AbyX/AbyV | AbsV | AbmV |
| UK94_RS25200 | 73 | ferredoxin | ferredoxin, *Frankia alni* (WP_011605214.1); 99/98 | - | AbsG1 | AbmG |
| UK94_RS25205 | 314 | aldo/keto reductase | aldo/keto reductase, *Frankia alni* (WP_011605213.1); 99/99 | - | AbsJ | AbmJ |
| UK94_RS25210 | 129 | hypothetical protein | hypothetical protein, *Frankia alni* (WP_011605212.1); 99/99 | AbyU | AbsU | AbmU |
| UK94_RS25215 | 951 | LuxR family transcriptional regulator | LuxR family transcriptional regulator, *Frankia alni* (WP_011605211.1); 99/99 | AbyH | - | AbmH |
| UK94_RS25220 | 256 | AfsR/SARP family transcriptional regulator | AfsR/SARP family transcriptional regulator, *Frankia alni* (WP_041939444.1); 99/99 | AbyR | - | - |
| UK94_RS25225 | 344 | 3-oxoacyl-ACP synthase III family protein | 3-oxoacyl-ACP synthase III family protein, *Frankia alni* (WP_011605207.1); 99/99 | AbyA1 | AbsA1 | AbmA1 |
| UK94_RS25230 | 74 | acyl carrier protein | hypothetical protein FRAAL4076, *Frankia alni* ACN14a (CAJ62718.1); 98/100 | AbyA3 | AbsA3 | AbmA3 |
| UK94_RS25235 | 252 | acyltransferase | acyltransferase, *Frankia alni* (WP_011605205.1); 99/99 | AbyA4 | AbsA4 | AbmA4 |
| UK94_RS25240 | 360 | alpha/beta hydrolase | alpha/beta hydrolase, *Frankia alni* (WP_011605204.1); 99/99 | AbyA5 | AbsA5 | AbmA5 |
| UK94_RS25245 | 281 | thioesterase | thioesterase, *Frankia alni* (WP_011605203.1); 99/99 | AbyT | AbsN | AbmT |
| UK94_RS26870 | 4436 | PKS I | type I polyketide synthase, *Frankia alni* ACN14a (CAJ62714.1); 99/98 | AbyB1 | AbsB1 | AbmB1 |
| //////////////////////////////////////////////////////////////////////////////////////////////////// | | | | | | |
| UK94_RS26865 | 3490 | PKS I | type I polyketide synthase, *Frankia alni* (WP_011605201.1); 99/99 | AbyB2 | AbsB2 | AbmB2 |
| UK94_RS26860 | 1042 | PKS I | acyltransferase domain-containing protein, *Frankia alni* (WP_011605200.1); 99/99 | AbyA3 | AbsB3 | AbmB3 |
| UK94_RS26855 | 629 | HAD-IIIC family phosphatase | HAD-IIIC family phosphatase, *Frankia alni* (WP_011605199.1); 99/99 | AbyA2 | AbsA2 | AbmA2 |

**Table S26.** Predicted functions of ORFs surrounding AbyU homolog from *Actinomadura fibrosa* LMG 29177 (CAACUY010000117.1).

| ORF | Size (aa) | Proposed function | Closest homolog, host (protein ID); Identity/Similarity (%) | Aby homolog | Abs homolog | Abm homolog |
|---|---|---|---|---|---|---|
| *E1300_RS30030* | 1651 | acyltransferase domain-containing protein | Erythronolide synthase, modules 1 and 2, *Streptomyces* sp. M56 (AUA16058.1); 50/61 | - | - | - |
| *E1300_RS30035* | 520 | multicopper oxidase family protein | copper oxidase, *Streptomyces* sp. NRRL S-813 (WP_030177947.1); 69/76 | - | - | - |
| *E1300_RS30040* | 158 | hypothetical protein | hypothetical protein, *Amycolatopsis taiwanensis* (WP_052372290.1); 36/55 | - | - | - |
| *E1300_RS30045* | 94 | hypothetical protein | hypothetical protein, *Streptomyces* sp. DvalAA-14 (WP_093739540.1); 43/55 | - | - | - |
| *E1300_RS30050* | 523 | acyl-CoA carboxylase subunit beta | acyl-CoA carboxylase subunit beta, *Streptomyces* sp. DvalAA-14 (WP_093739542.1); 82/88 | - | - | - |
| *E1300_RS30055* | 160 | Diels-Alderase | hypothetical protein, *Streptomyces* sp. NRRL F-525 (WP_033287247.1); 32/54 | AbyU | AbsU | AbmU |
| *E1300_RS30060* | 466 | DHA2 family efflux MFS transporter permease subunit | DHA2 family efflux MFS transporter permease subunit, *Rhodococcus* sp. SMB37 (WP_132471207.1); 51/66 | - | - | - |
| *E1300_RS30065* | 192 | MarR family transcriptional regulator | MarR family transcriptional regulator, *Streptomyces formicae* (WP_098245770.1); 43/63 | - | - | - |
| *E1300_RS30070* | 496 | hypothetical protein | hypothetical protein, *Streptomyces agglomeratus* (WP_069929832.1); 53/68 | - | - | - |
| *E1300_RS30075* | 323 | hypothetical protein | - | - | - | - |
| *E1300_RS30080* | 279 | hypothetical protein | Phage integrase family protein, *Actinomadura echinospora* (SEF92328.1); 69/79 | - | - | - |
| *E1300_RS30085* | 160 | NUDIX domain-containing protein | hypothetical protein, *Actinokineospora inagensis* (WP_035306982.1); 50/66 | - | - | - |
| *E1300_RS30090* | 579 | DUF2326 domain-containing protein | NUDIX domain-containing protein, *Nonomuraea wenchangensis* (WP_091081257.1); 62/76 | - | - | - |
| *E1300_RS30095* | 84 | hypothetical protein | - | - | - | - |

**Table S27.** Predicted functions of ORFs in abyssomicin BGC from *Frankia alni* ACN14a (NC_008278.1).

| ORF | Size (aa) | Proposed function | Closest homolog, host (protein ID); Identity/Similarity (%) | Aby homolog | Abs homolog | Abm homolog |
|---|---|---|---|---|---|---|
| *FRAAL_RS17820* | 629 | HAD-IIIC family phosphatase | HAD-IIIC family phosphatase, *Frankia* sp. AvcI1 (WP_055752028.1); 99/99 | AbyA2 | AbsA2 | AbmA2 |
| *FRAAL_RS17825* | 1042 | acyltransferase domain-containing protein | acyltransferase domain-containing protein, *Frankia* sp. AvcI1 (WP_055752029.1); 99/99 | AbyB3 | AbsB3 | AbmB3 |
| *FRAAL_RS17830* | 3486 | PKS I | type I polyketide synthase, *Frankia* sp. AvcI1 (WP_055752030.1); 99/99 | AbyB2 | AbsB2 | AbmB2 |
| *FRAAL_RS32455* | - | hypothetical protein | type I polyketide synthase, *Frankia* sp. AvcI1 (WP_095213022.1); 99/99 | AbyB1 | AbsB1 | AbmB1 |
| *FRAAL_RS32460* | | polyketide synthase | type I polyketide synthase, *Frankia* sp. AvcI1 (WP_095213022.1); 99/99 | | | |
| *FRAAL_RS32465* | | hypothetical protein | type I polyketide synthase, *Frankia* sp. AvcI1 (WP_095213022.1); 99/98 | | | |
| *FRAAL_RS31810* | | hypothetical protein | type I polyketide synthase, *Frankia* sp. AvcI1 (WP_095213022.1); 100/100 | | | |
| *FRAAL_RS31815* | | polyketide synthase | type I polyketide synthase, *Frankia* sp. AvcI1 (WP_095213022.1); 99/99 | | | |
| *FRAAL_RS31820* | | hypothetical protein | type I polyketide synthase, *Frankia* sp. AvcI1 (WP_095213022.1); 97/96 | | | |
| *FRAAL_RS31825* | | hypothetical protein | type I polyketide synthase, *Frankia* sp. AvcI1 (WP_095213022.1); 99/99 | | | |
| *FRAAL_RS31830* | | polyketide synthase | AbsB1, *Streptomyces* sp. LC-6-2 (ARE67853.1); 75/83 | | | |
| *FRAAL_RS32470* | | 3-ketoacyl-ACP synthase | AbsB1, *Streptomyces* sp. LC-6-2 (ARE67853.1); 60/69 | | | |
| *FRAAL_RS31835* | | modular polyketide synthase BFAS4 | AbsB1, *Streptomyces* sp. LC-6-2 (ARE67853.1); 67/73 | | | |
| *FRAAL_RS32475* | | hypothetical protein | AbsB1, *Streptomyces* sp. LC-6-2 (ARE67853.1); 68;76 | | | |
| *FRAAL_RS31850* | | polyketide synthase | AbsB1, *Streptomyces* sp. LC-6-2 (ARE67853.1); 65/75 | | | |
| *FRAAL_RS31855* | | hypothetical protein | polyketide synthase, *Microbispora* sp. GKU 823 (WP_079314734.1); 57/70 | | | |
| *FRAAL_RS31860* | | hypothetical protein | polyketide synthase type I, *Streptomyces* sp. E14 (EFF94120.1); 78/83 | | | |
| *FRAAL_RS17840* | 281 | thioesterase | thioesterase, *Frankia* sp. AvcI1 (WP_055751806.1); 99/99 | AbyT | AbsN | AbmT |
| *FRAAL_RS17845* | 360 | alpha/beta hydrolase | alpha/beta hydrolase, *Frankia* sp. AvcI1 (WP_055751805.1); 99/99 | AbyA5 | AbsA5 | AbmA5 |
| *FRAAL_RS17850* | 252 | acyltransferase | acyltransferase, *Frankia* sp. AvcI1 (WP_055751804.1); 99/99 | AbyA4 | AbsA4 | AbmA4 |
| *FRAAL_RS17855* | 74 | acyl carrier protein | acyl carrier protein, *Streptomyces* sp. NRRL F-5126 (WP_030912403.1); 53/73 | AbyA3 | AbsA3 | AbmA3 |
| *FRAAL_RS17860* | 344 | 3-oxoacyl-ACP synthase III family protein | 3-oxoacyl-ACP synthase III family protein, *Frankia* sp. AvcI1 (WP_055751803.1); 99/99 | AbyA1 | AbsA1 | AbmA1 |
| *FRAAL_RS17865* | 256 | AfsR/SARP family transcriptional regulator | AfsR/SARP family transcriptional regulator, *Frankia* sp. AvcI1 (WP_055751802.1); 99/98 | AbyR | - | - |
| *FRAAL_RS17870* | 951 | LuxR family transcriptional regulator | LuxR family transcriptional regulator, *Frankia* sp. AvcI1 (WP_055751801.1); 99/99 | AbyH | - | AbmH |
| *FRAAL_RS17875* | 129 | Diels-Alderase | hypothetical protein, *Frankia* sp. AvcI1 (WP_055751820.1); 99/99 | AbyU | AbsU | AbmU |

| | | | | | | |
|---|---|---|---|---|---|---|
| *FRAAL_RS17880* | 341 | aldo/keto reductase | aldo/keto reductase, *Frankia* sp. AvcI1 (WP_055751800.1); 99/99 | - | AbsJ | AbmJ |
| *FRAAL_RS17885* | 73 | ferredoxin | ferredoxin, *Frankia* sp. AvcI1 (WP_055751799.1); 99/98 | - | AbsG1 | AbmG |
| *FRAAL_RS17890* | 396 | cytochrome P450 | cytochrome P450, *Frankia* sp. AvcI1 (WP_055751798.1); 98/98 | AbyX | AbsV | AbmV |
| *FRAAL_RS17895* | 598 | ABC transporter ATP-binding protein | ABC transporter ATP-binding protein, *Frankia* sp. AvcI1 (WP_095212991.1); 99/99 | AbyF4 | AbsF4 | AbmF4 |
| *FRAAL_RS17900* | 262 | ABC transporter permease | ABC transporter permease, *Frankia* sp. AvcI1 (WP_055751819.1); 98/98 | AbyF3 | AbsF3 | AbmF3 |
| *FRAAL_RS17905* | 337 | ABC transporter permease | ABC transporter permease, *Frankia* sp. AvcI1 (WP_063845354.1); 99/99 | AbyF2 | AbsF2 | AbmF2 |
| *FRAAL_RS17910* | 552 | ABC transporter substrate-binding protein | ABC transporter substrate-binding protein, *Frankia* sp. AvcI1 (WP_055751795.1); 98/98 | AbyF1 | AbsF1 | AbmF1 |
| *FRAAL_RS17915* | 497 | MFS transporter | MFS transporter, *Frankia* sp. AvcI1 (WP_055751794.1); 99/99 | AbyD | AbsD | AbmD |
| *FRAAL_RS17920* | 331 | LLM class flavin-dependent oxidoreductase | LLM class flavin-dependent oxidoreductase, *Frankia* sp. AvcI1 (WP_055751793.1); 99/99 | AbyE | AbsE | AbmE1 |
| *FRAAL_RS17925* | 373 | cytochrome P450 | cytochrome P450, *Frankia* sp. AvcI1 (WP_055751792.1); 99/99 | AbyX/AbyV | AbsX | AbmV |
| *FRAAL_RS17930* | 203 | TetR/AcrR family transcriptional regulator | TetR/AcrR family transcriptional regulator, *Frankia* sp. AvcI1 (WP_055751791.1); 98/98 | - | AbsC2 | - |

**Table S28.** Predicted functions of ORFs in abyssomicin BGC from *Frankia discariae* BCU110501 (NZ_KB891214 and NZ_KB891274).

| ORF | Size (aa) | Proposed function | Closest homolog, host (protein ID); Identity/Similarity (%) | Aby homolog | Abs homolog | Abm homolog |
|---|---|---|---|---|---|---|
| *B056_RS0115515* | 392 | LLM class flavin-dependent oxidoreductase | LLM class flavin-dependent oxidoreductase, *Frankia* sp. EAN1pec (WP_020461007.1); 94/95 | AbyE | AbsE | AbmE1 |
| *B056_RS0115520* | 234 | TetR/AcrR family transcriptional regulator | TetR/AcrR family transcriptional regulator, *Frankia* sp. EAN1pec (WP_020461008.1); 94/96 | AbyC | - | AbmC |
| *B056_RS0115525* | 360 | LLM class flavin-dependent oxidoreductase | LLM class flavin-dependent oxidoreductase, *Frankia* sp. EAN1pec (WP_020461009.1); 96/96 | - | - | AbmE2 |
| *B056_RS0115530* | 499 | hypothetical protein | propionyl-CoA carboxylase subunit beta, *Frankia* sp. EI5c (WP_066072904.1); 81/85 | - | - | - |
| *B056_RS0115535* | 70 | hypothetical protein | hypothetical protein, *Frankia* sp. EAN1pec (WP_041254286.1); 88/89 | - | - | - |
| *B056_RS0115540* | 405 | cytochrome P450 | cytochrome P450, *Frankia* sp. EAN1pec (WP_020461012.1); 98/99 | AbyX/AbyV | AbsV/AbsX | AbmV |
| *B056_RS0115545* | 1093 | PKS I | type I polyketide synthase, *Frankia* sp. EI5c (WP_083987208.1); 77/82 | AbyB3 | AbsB3 | AbmB3 |
| *B056_RS0115550* | 2276 | PKS I | type I polyketide synthase, *Frankia* sp. EAN1pec (WP_020461014.1); 90/92 | AbyB2 | AbsB2 | AbmB2 |
| *B056_RS36660* | - | PKS I | type I polyketide synthase, *Streptomyces* sp. KhCrAH-43 (WP_018522876.1); 60/67 | AbyB1 | AbsB1 | AbmB1 |
| //////////////////////////////////////////////////////////////////////////////////////////////////////////////////////////////// | | | | | | |
| *B056_RS42215* | - | PKS I | type I polyketide synthase, *Streptomyces* sp. KhCrAH-43 (WP_018522876.1); 60/67 | AbyB1 | AbsB1 | AbmB1 |
| *B056_RS0132560* | 65 | ferredoxin | ferredoxin, *Frankia* sp. EI5c (WP_066073232.1); 92/96 | AbyW | AbsG2/AbsG1 | AbmG |
| *B056_RS0132555* | 399 | cytochrome P450 | cytochrome P450, *Frankia* sp. EAN1pec (WP_020461016.1); 98/99 | AbyV/AbyX | AbsV/AbsX | AbmV |
| *B056_RS0132550* | 564 | ABC transporter ATP-binding protein | ABC transporter ATP-binding protein, *Frankia* sp. EAN1pec (WP_049795952.1); 94/95 | AbyF4 | AbsF4 | AbmF4 |
| *B056_RS0132545* | 288 | ABC transporter permease | ABC transporter permease, *Frankia* sp. EAN1pec (WP_020461018.1); 93/94 | AbyF3 | AbsF3 | AbmF3 |
| *B056_RS0132540* | 317 | ABC transporter permease | ABC transporter permease, *Frankia* sp. EAN1pec (WP_020461019.1); 95/97 | AbyF2 | AbsF2 | AbmF2 |
| *B056_RS0132535* | 547 | ABC transporter substrate-binding protein | ABC transporter substrate-binding protein, *Frankia* sp. EAN1pec (WP_020461020.1); 96/97 | AbyF1 | AbsF1 | AbmF1 |
| *B056_RS0132530* | 347 | LLM class flavin-dependent oxidoreductase | LLM class flavin-dependent oxidoreductase, *Frankia* sp. EAN1pec (WP_020461021.1); 99/99 | AbyE | AbsE | AbmE1 |
| *B056_RS0132525* | 486 | DHA2 family efflux MFS transporter permease subunit | DHA2 family efflux MFS transporter permease subunit, *Frankia* sp. EAN1pec (WP_020461022.1); 99/99 | AbyD | AbsD | AbmD |
| *B056_RS0132520* | 250 | TetR/AcrR family transcriptional regulator | TetR/AcrR family transcriptional regulator, *Frankia* sp. EAN1pec (WP_020461023.1); 99/99 | AbyC | - | AbmC |
| *B056_RS0132515* | 374 | alpha/beta hydrolase | alpha/beta hydrolase, *Frankia* sp. EAN1pec (WP_020461024.1); 95/95 | AbyA5 | AbsA5 | AbmA5 |
| *B056_RS0132510* | 259 | acyltransferase | acyltransferase, *Frankia* sp. EAN1pec (WP_049795953.1); 98/100 | AbyA4 | AbsA4 | AbmA4 |
| *B056_RS0132505* | 78 | acyl carrier protein | acyl carrier protein, *Frankia* sp. EI5c (WP_066073198.1); 92/94 | AbyA3 | AbsA3 | AbmA3 |

| | | | | | | |
|---|---|---|---|---|---|---|
| *B056_RS0132500* | 658 | HAD-IIIC family phosphatase | HAD-IIIC family phosphatase, *Streptomyces* sp. KhCrAH-43 (WP_018522892.1); 70/78 | AbyA2 | AbsA2 | AbmA2 |
| *B056_RS0132495* | 348 | 3-oxoacyl-ACP synthase III family protein | 3-oxoacyl-ACP synthase III family protein, *Frankia* sp. EAN1pec (WP_020461027.1); 99/100 | AbyA1 | AbsA1 | AbmA1 |
| *B056_RS0132490* | 1024 | RHS repeat protein + Diels-Alderase | RHS repeat protein, *Frankia* sp. EAN1pec (WP_020461028.1); 95/95 | AbyK+AbyU | AbsU | AbmU |

**Table S29.** Predicted functions of ORFs in abyssomicin BGC from *Frankia sp.* EAN1pec (CP000820.1).

| ORF | Size (aa) | Proposed function | Closest homolog, host (protein ID); Identity/Similarity (%) | Aby homolog | Abs homolog | Abm homolog |
|---|---|---|---|---|---|---|
| *Franean1_3465* | 386 | luciferase family protein | LLM class flavin-dependent oxidoreductase, *Frankia discariae* (WP_026239744.1);94/95 | AbyE | AbsE | AbmE1 |
| *Franean1_3466* | 245 | transcriptional regulator, TetR family | TetR/AcrR family transcriptional regulator, *Frankia discariae* (WP_020572516.1); 94/96 | AbyC | - | AbmC |
| *Franean1_3467* | 354 | luciferase family protein | LLM class flavin-dependent oxidoreductase, *Frankia discariae* (WP_018502788.1); 96/96 | - | - | AbmE2 |
| *Franean1_3468* | 141 | putative acetyl/propionyl CoA carboxylase beta subunit | hypothetical protein, *Frankia discariae* (WP_018502789.1); 80/80 | - | - | - |
| *Franean1_3469* | 100 | hypothetical protein | hypothetical protein, *Frankia discariae* (WP_018502790.1); 86/88 | - | - | - |
| *Franean1_3470* | 404 | cytochrome P450 | cytochrome P450, *Frankia discariae* (WP_026239745.1); 98/99 | AbyX/AbyV | AbsV/AbsX | AbmV |
| *Franean1_3471* | 1071 | PKS I | type I polyketide synthase, *Frankia discariae* (WP_026239746.1); 93/94 | AbyB3 | AbsB3 | AbmB3 |
| *Franean1_3472* | 4111 | PKS I | type I polyketide synthase, *Streptomyces* sp. 2131.1 (WP_093709984.1); 59/67 | AbyB2 | AbsB2 | AbmB2 |
| *Franean1_3473* | 4840 | PKS I | type I polyketide synthase, *Streptomyces* sp. 2131.1 (WP_093709985.1); 53/62 | AbyB1 | AbsB1 | AbmB1 |
| *Franean1_3474* | 64 | protein of unknown function DUF1271 | Ferredoxin*, Streptomyces fragilis* (WP_108952945.1); 59/78 | AbyW | AbsG2/AbsG1 | AbmG |
| *Franean1_3475* | 398 | cytochrome P450 | cytochrome P450, *Frankia discariae* (WP_018506032.1); 98/99 | AbyV/AbyX | AbsV/AbsX | AbmV |
| *Franean1_3476* | 594 | ABC transporter related | ABC transporter ATP-binding protein, *Frankia discariae* (WP_051105801.1); 94/95 | AbyF4 | AbsF4 | AbmF4 |
| *Franean1_3477* | 287 | binding-protein-dependent transport systems inner membrane component | ABC transporter permease, *Frankia discariae* (WP_026240411.1); 93/94 | AbyF3 | AbsF3 | AbmF3 |
| *Franean1_3478* | 316 | binding-protein-dependent transport systems inner membrane component | ABC transporter permease, *Frankia discariae* (WP_018506029.1); 95/97 | AbyF2 | AbsF2 | AbmF2 |
| *Franean1_3479* | 546 | extracellular solute-binding protein family 5 | ABC transporter substrate-binding protein, *Frankia discariae* (WP_018506028.1); 96/97 | AbyF1 | AbsF1 | AbmF1 |
| *Franean1_3480* | 346 | luciferase family protein | LLM class flavin-dependent oxidoreductase, *Frankia discariae* (WP_018506027.1); 99/99 | AbyE | AbsE | AbmE1 |
| *Franean1_3481* | 845 | drug resistance transporter, EmrB/QacA subfamily | DHA2 family efflux MFS transporter permease subunit, *Frankia discariae* (WP_018506026.1); 99/99 | AbyD | AbsD | AbmD |
| *Franean1_3482* | 248 | transcriptional regulator, TetR family | etR/AcrR family transcriptional regulator, *Frankia discariae* (WP_018506025.1); 94/94 | AbyC | - | AbmC |
| *Franean1_3483* | 373 | conserved hypothetical protein | alpha/beta hydrolase, *Frankia discariae* (WP_018506024.1); 95/95 | AbyA5 | AbsA5 | AbmA5 |
| *Franean1_3484* | 294 | catalytic domain of components of various dehydrogenase complexes | Acyltransferase*, Frankia discariae* (WP_026240410.1); 98/100 | AbyA4 | AbsA4 | AbmA4 |
| *Franean1_3485* | 77 | hypothetical protein | acyl carrier protein, *Streptomyces* sp. NRRL WC-3725 (WP_031029037.1); 80/88 | AbyA3 | AbsA3 | AbmA3 |

| | | | | | | |
|---|---|---|---|---|---|---|
| *Franean1_3486* | 142 | conserved hypothetical protein | HAD-IIIC family phosphatase, *Frankia discariae* (WP_018506021.1); 84/87 | AbyA2 | AbsA2 | AbmA2 |
| *Franean1_3487* | 348 | Beta-ketoacyl-acyl-carrier-protein synthase I | 3-oxoacyl-ACP synthase III family protein, *Frankia discariae* (WP_018506020.1); 99/100 | AbyA1 | AbsA1 | AbmA1 |
| *Franean1_3488* | 1023 | YD repeat protein + Diels-Alderase | RHS repeat protein, *Frankia discariae* (WP_018506019.1); 94/95 | AbyK+AbyU | AbsU | AbmU |
| *Franean1_3489* | 257 | transcriptional regulator, SARP family | SARP family transcriptional regulator, *Frankia discariae* (WP_018506018.1); 98/99 | AbyI | - | AbmI |
| *Franean1_3490* | 950 | LuxR family transcriptional regulator | helix-turn-helix transcriptional regulator, *Frankia discariae* (WP_018506017.1); 97/97 | AbyH | - | AbmH |

**Table S30.** Predicted functions of ORFs in abyssomicin BGC from *Frankia sp.* EI5c (NZ_LRTK01000008.1 and NZ_LRTK01000088.1).

| ORF | Size (aa) | Proposed function | Closest homolog, host (protein ID); Identity/Similarity (%) | Aby homolog | Abs homolog | Abm homolog |
|---|---|---|---|---|---|---|
| *UG55_RS07380* | 142 | Diels-Alderase | hypothetical protein, *Frankia sp*. Cc1.17 (WP_071084438.1); 97/99 | AbyU | AbsU | AbmU |
| *UG55_RS07385* | 349 | 3-oxoacyl-ACP synthase III family protein | 3-oxoacyl-ACP synthase III family protein, *Frankia* sp. Cc1.17 (WP_071084440.1); 94/96 | AbyA1 | AbsA1 | AbmA1 |
| *UG55_RS07390* | 672 | HAD-IIIC family phosphatase | HAD-IIIC family phosphatase, *Frankia* sp. Cc1.17 (WP_116287792.1); 87/89 | AbyA2 | AbsA2 | AbmA2 |
| *UG55_RS07395* | 188 | TetR family transcriptional regulator | TetR/AcrR family transcriptional regulator, *Frankia* sp. Cc1.17 (WP_084132131.1); 94/96 | - | - | - |
| *UG55_RS07400* | 472 | DHA2 family efflux MFS transporter permease subunit | DHA2 family efflux MFS transporter permease subunit, *Frankia* sp. Cc1.17 (WP_116287793.1); 95/97 | AbyD | AbsD | AbmD |
| *UG55_RS07405* | 336 | LLM class flavin-dependent oxidoreductase | LLM class flavin-dependent oxidoreductase, *Frankia* sp. Cc1.17 (WP_071084502.1); 93/96 | AbyE | AbsE | AbmE1 |
| *UG55_RS07410* | 1049 | helix-turn-helix transcriptional regulator | LuxR family transcriptional regulator, *Frankia* sp. Cc1.17 (WP_071084446.1); 88/91 | AbyH | - | AbmH |
| *UG55_RS07415* | 257 | AfsR/SARP family transcriptional regulator | hypothetical protein, *Frankia* sp. Cc1.17 (WP_071084449.1); 89/94 | AbyI | - | AbmI |
| *UG55_RS07420* | 76 | acyl carrier protein | acyl carrier protein, *Frankia* sp. Cc1.17 (WP_071084451.1); 97/97 | AbyA3 | AbsA3 | AbmA3 |
| *UG55_RS07425* | 304 | acyltransferase | acyltransferase, *Frankia* sp. Cc1.17 (WP_084132123.1); 86/88 | AbyA4 | AbsA4 | AbmA4 |
| *UG55_RS07430* | 359 | alpha/beta hydrolase | alpha/beta hydrolase, *Frankia* sp. Cc1.17 (WP_071084457.1) | AbyA5 | AbsA5 | AbmA5 |
| *UG55_RS07435* | - | PKS I | Contig edge | PKS I | PKS I | PKS I |
| ////////////////////////////////////////////////////////////////////////////////////////////////////////////////// | | | | | | |
| *UG55_RS23610* | - | PKS I | Contig edge | PKS I | PKS I | PKS I |
| *UG55_RS23605* | 65 | ferredoxin | ferredoxin, *Streptomyces regensis* (KMS84448.1); 59/81 | - | AbsG2 | - |
| *UG55_RS23600* | 399 | cytochrome P450 | cytochrome P450, *Frankia* sp. EAN1pec (WP_020461016.1); 90/93 | AbyV/AbyX | AbsV/AbsX | AbmV |
| *UG55_RS23595* | 577 | ABC transporter ATP-binding protein | ABC transporter ATP-binding protein, *Frankia discariae* (WP_051105801.1); 80/86 | AbyF4 | AbsF4 | AbmF4 |
| *UG55_RS23590* | 287 | ABC transporter permease | ABC transporter permease, *Frankia* sp. Cc1.17 (WP_084131831.1); 81/88 | AbyF3 | AbsF3 | AbmF3 |
| *UG55_RS23585* | 197 | ABC transporter permease | ABC transporter permease, *Frankia discariae* (WP_018506029.1); 79/88 | AbyF2 | AbsF2 | AbmF2 |
| *UG55_RS23580* | 547 | ABC transporter substrate-binding protein | ABC transporter substrate-binding protein, *Frankia discariae* (WP_018506028.1); 87/91 | AbyF1 | AbsF1 | AbmF1 |
| *UG55_RS23575* | 348 | LLM class flavin-dependent oxidoreductase | LLM class flavin-dependent oxidoreductase, *Frankia* sp. BMG5.11 (TCJ32075.1); 82/87 | AbyE | AbsE | AbmE1 |
| *UG55_RS2357* | 491 | DHA2 family efflux MFS transporter | DHA2 family efflux MFS transporter permease subunit, *Frankia* sp. EAN1pec | AbyD | AbsD | AbmD |

| | | | | | | |
|---|---|---|---|---|---|---|
| *0* | | permease subunit | (WP_020461022.1); 90/94 | | | |
| *UG55_RS23565* | 274 | TetR/AcrR family transcriptional regulator | TetR/AcrR family transcriptional regulator, *Frankia* sp. Ea1.12 (WP_112105117.1); 83/87 | AbyC | - | AbmC |
| *UG55_RS23560* | 362 | alpha/beta hydrolase | alpha/beta hydrolase, *Frankia* sp. EAN1pec (WP_020461024.1); 84/85 | AbyA5 | AbsA5 | AbmA5 |
| *UG55_RS23555* | 246 | acyltransferase | acyltransferase, *Frankia discariae* (WP_026240410.1); 85/92 | AbyA4 | AbsA4 | AbmA4 |
| *UG55_RS23550* | 78 | acyl carrier protein | acyl carrier protein, *Streptomyces* sp. NRRL WC-3725 (WP_031029037.1); 75/85 | AbyA3 | AbsA3 | AbmA3 |
| *UG55_RS23545* | 342 | 3-oxoacyl-ACP synthase III family protein | 3-oxoacyl-ACP synthase III family protein, *Frankia* sp. EAN1pec (WP_020461027.1); 87/91 | AbyA1 | AbsA1 | AbmA1 |
| *UG55_RS23535* | 1080 | RHS repeat protein | RHS repeat protein, *Frankia discariae* (WP_018506019.1); 67/73 | AbyK | - | - |
| *UG55_RS23530* | 258 | activator protein | SARP family transcriptional regulator, *Frankia discariae* (WP_018506018.1); 90/93 | AbyI | - | AbmI |

**Table S31.** Predicted functions of ORFs in potential abyssomicin BGC from *Frankia* symbiont of *Datisca glomerata* (NC_015656.1).

| ORF | Size (aa) | Proposed function | Closest homolog, host (protein ID); Identity/Similarity (%) | Aby homolog | Abs homolog | Abm homolog |
|---|---|---|---|---|---|---|
| *FSYMDG_RS09425* | 528 | hypothetical protein | acyltransferase, *Amycolatopsis circi* (WP_116201221.1); 55/68 | - | AbsI | - |
| *FSYMDG_RS09430* | 68 | ferredoxin | ferredoxin, *Frankia coriariae* (KLL10399.1); 93/97 | - | AbsG1 | AbmG |
| *FSYMDG_RS09435* | 274 | thioesterase | thioesterase, *Frankia* sp. AvcI1 (WP_055749039.1); 66/76 | AbyT | AbsN | AbmT |
| *FSYMDG_RS24310* | - | PKS I | acyltransferase domain-containing protein, partial, *Frankia symbiont of Coriaria nepalensis* (WP_131772417.1); 99/100 | PKS I | PKS I | PKS I |
| *FSYMDG_RS25265* | - | PKS I | type I polyketide synthase, *Frankia* sp. BMG5.30 (WP_076843523.1); 70/77 | PKS I | PKS I | PKS I |
| *FSYMDG_RS25270* | - | PKS I | type I polyketide synthase, *Streptomyces alfalfae* (WP_076682132.1); 55/62 | PKS I | PKS I | PKS I |
| *FSYMDG_RS24325* | - | PKS I | acyltransferase domain-containing protein, partial, *Frankia symbiont of Coriaria nepalensis* (WP_131772419.1); 100/100 | PKS I | PKS I | PKS I |
| *FSYMDG_RS24330* | - | PKS I | type I polyketide synthase, *Streptomyces alboviridis* (WP_032759890.1); 69/76 | PKS I | PKS I | PKS I |
| *FSYMDG_RS24340* | - | PKS I | SDR family NAD(P)-dependent oxidoreductase, partial, *Frankia symbiont of Coriaria nepalensis* (WP_131772420.1); 99/99 | PKS I | PKS I | PKS I |
| *FSYMDG_RS24345* | - | PKS I | type I polyketide synthase, *Streptomyces exfoliatus* (WP_078626965.1); 57/67 | PKS I | PKS I | PKS I |
| *FSYMDG_RS25275* | - | PKS I | acyltransferase domain-containing protein, *Frankia symbiont of Coriaria nepalensis* (WP_131772422.1); 99/100 | PKS I | PKS I | PKS I |
| *FSYMDG_RS24360* | - | PKS I | acyltransferase domain-containing protein, partial, *Frankia symbiont of Coriaria myrtifolia* (WP_131773939.1); 99/100 | PKS I | PKS I | PKS I |
| *FSYMDG_RS09445* | 345 | alpha/beta hydrolase | alpha/beta hydrolase, *Frankia* sp. BMG5.30 (ONH34857.1); 96/96 | AbyA5 | AbsA5 | AbmA5 |
| *FSYMDG_RS09450* | 310 | acyltransferase | acyltransferase, *Frankia* sp. BMG5.30 (WP_076843553.1); 92/93 | AbyA4 | AbsA4 | AbmA4 |
| *FSYMDG_RS09455* | 343 | 3-oxoacyl-ACP synthase III family protein | 3-oxoacyl-ACP synthase, *Frankia coriariae* (KLL11317.1); 98/98 | AbyA1 | AbsA1 | AbmA1 |
| *FSYMDG_RS09460* | 569 | monooxygenase FAD-binding protein | hypothetical protein, *Frankia* sp. BMG5.30 (WP_076843525.1); 89/91 | - | - | - |
| *FSYMDG_RS09465* | 356 | LLM class flavin-dependent oxidoreductase | LLM class flavin-dependent oxidoreductase, *Frankia* sp. BMG5.30 (WP_076843526.1); 96/98 | - | - | AbmE2 |
| *FSYMDG_RS09470* | 230 | cytochrome P450 | cytochrome P450, *Frankia coriariae* (KLL11355.1); 99/100 | AbyX/AbyV | AbsV/AbsX | AbmV |
| *FSYMDG_RS09475* | 478 | IS66 family transposase | IS66 family transposase, *Frankia* sp. ACN1ag (WP_055409628.1); 82/85 | - | - | - |
| *FSYMDG_RS09480* | 155 | cytochrome P450 | cytochrome P450, *Frankia coriariae* (KLL11355.1); 99/99 | AbyX/AbyV | AbsX/AbsV | AbmV |
| *FSYMDG_RS09485* | 83 | ferredoxin | ferredoxin, *Frankia coriariae* (KLL11356.1); 95/96 | - | AbsG1/AbsG2 | AbmG |
| *FSYMDG_RS09490* | 292 | response regulator transcription factor | LuxR family transcriptional regulator, *Frankia* sp. BMG5.30 (WP_083731095.1); 92/92 | AbyH | - | AbmH |
| *FSYMDG_RS09495* | 202 | TetR/AcrR family | TetR/AcrR family transcriptional regulator, *Frankia coriariae* | - | AbsC2 | - |

| | | | | | | |
|---|---|---|---|---|---|---|
| | | transcriptional regulator | (WP_052914596.1); 97/97 | | | |
| *FSYMDG_RS09500* | 114 | hypothetical protein | LuxR family transcriptional regulator, *Frankia* sp. BMG5.30 (WP_083731095.1); 77/80 | - | - | - |
| *FSYMDG_RS09505* | 246 | AfsR/SARP family transcriptional regulator | activator protein, *Frankia* sp. BMG5.30 (WP_076843528.1); 99//99 | AbyI | - | AbmI |
| *FSYMDG_RS09510* | 328 | aldo/keto reductase | aldo/keto reductase, *Frankia* sp. BMG5.30 (WP_076843529.1); 98/98 | - | AbsJ | AbmJ |
| *FSYMDG_RS09515* | 147 | hypothetical protein | hypothetical protein, *Frankia* sp. BMG5.30 (WP_076843530.1); 95/96 | - | - | - |
| *FSYMDG_RS09530* | 496 | FMN-binding glutamate synthase family protein | FMN-binding glutamate synthase family protein, *Frankia* sp. BMG5.30 (WP_076843531.1); 97/97 | - | - | - |
| *FSYMDG_RS09535* | 530 | propionyl-CoA carboxylase subunit beta | propionyl-CoA carboxylase subunit beta, *Frankia coriariae* (KLL11331.1); 97/97 | - | - | - |
| *FSYMDG_RS09545* | 75 | acyl carrier protein | acyl carrier protein, *Frankia* sp. EI5c (WP_066064666.1); 65/76 | AbyA3 | AbsA3 | AbmA3 |
| *FSYMDG_RS09550* | 654 | HAD-IIIC family phosphatase | HAD-IIIC family phosphatase, *Frankia* sp. BMG5.30 (WP_076843558.1); 98/98 | AbyA2 | AbsA2 | AbmA2 |
| *FSYMDG_RS09555* | 498 | MFS transporter | MFS transporter, *Microbispora triticiradicis* (WP_117409462.1); 50/67 | AbyD | AbsD | AbmD |
| *FSYMDG_RS09560* | 238 | TetR/AcrR family transcriptional regulator | TetR/AcrR family transcriptional regulator, *Streptacidiphilus* sp. DSM 106435 (WP_111490402.1); 62/74 | AbyC | - | AmbC |
| *FSYMDG_RS09565* | 125 | Diels-Alderase | hypothetical protein FrCorBMG51_12000, *Frankia coriariae* (KLL11361.1); 96/100 | AbyU | AbsU | AbmU |
| *FSYMDG_RS09570* | 448 | NtaA/DmoA family FMN-dependent monooxygenase | LLM class flavin-dependent oxidoreductase, *Paenibacillus* sp. JDR-2 (WP_015846774.1); 60/76 | - | - | - |
| *FSYMDG_RS09575* | 298 | ABC transporter ATP-binding protein | ABC transporter ATP-binding protein, *Frankia coriariae* (WP_086055414.1); 92/93 | AbyF4 | AbsF4 | AbmF4 |
| *FSYMDG_RS24370* | 674 | ATP-binding cassette domain-containing protein | dipeptide/oligopeptide/nickel ABC transporter permease/ATP-binding protein, *Frankia coriariae* (WP_047224248.1); 53/64 | AbyF3+AbyF4 | AbsF3+AbsF4 | AbmF3+AbmF4 |
| *FSYMDG_RS09590* | 337 | ABC transporter permease | ABC transporter permease, *Frankia coriariae* (KLL11332.1); 97/98 | AbmF2 | AbsF2 | AbmF2 |

**Table S32.** Predicted functions of ORFs in abyssomicin BGC from *Frankia* sp. Cc1.17 (MBLM01000080.1).

| ORF | Size (aa) | Proposed function | Closest homolog, host (protein ID); Identity/Similarity (%) | Aby homolog | Abs homolog | Abm homolog |
|---|---|---|---|---|---|---|
| *CC117_RS09725* | 371 | LLM class flavin-dependent oxidoreductase | luciferase, *Frankia* sp. Cc1.17 (OHV40281.1); 99/100 | AbyE | AbsE | AbmE1 |
| *CC117_RS09735* | 611 | ABC transporter ATP-binding protein | ABC transporter related, *Frankia* sp. EAN1pec (ABW12878.1); 84/87 | AbyF4 | AbsF4 | AbmF4 |
| *CC117_RS09740* | 287 | ABC transporter permease | ABC transporter permease, *Frankia* sp. EAN1pec (WP_020461018.1); 85/91 | AbyF3 | AbsF3 | AbmF3 |
| *CC117_RS09745* | 316 | ABC transporter permease | ABC transporter permease, *Frankia discariae* (WP_018506029.1); 92/96 | AbyF2 | AbsF2 | AbmF2 |
| *CC117_RS09750* | 546 | ABC transporter substrate-binding protein | ABC transporter substrate-binding protein, *Frankia* sp. EAN1pec (WP_020461020.1); 88/92 | AbyF1 | AbsF1 | AbmF1 |
| *CC117_RS09755* | 339 | LLM class flavin-dependent oxidoreductase | LLM class flavin-dependent oxidoreductase, *Frankia* sp. EAN1pec (WP_020461021.1); 85/91 | AbyE | AbsE | AbmE1 |
| *CC117_RS09760* | 285 | AfsR/SARP family transcriptional regulator | activator protein, *Frankia* sp. EI5c (WP_066073189.1); 83/89 | AbyI | - | AbmI |
| *CC117_RS09765* | 921 | LuxR family transcriptional regulator | helix-turn-helix transcriptional regulator, *Frankia* discariae (WP_018506017.1); 76/82 | AbyH | - | AbmH |
| *CC117_RS09770* | 277 | MetQ/NlpA family ABC transporter substrate-binding protein | ABC transporter substrate-binding protein, *Frankia* sp. EUN1f (WP_006542932.1); 85/91 | - | - | - |
| *CC117_RS09775* | 199 | ABC transporter permease | ABC transporter permease, *Frankia* sp. EUN1f (WP_006542931.1); 91/93 | - | - | - |
| *CC117_RS09780* | 360 | ATP-binding cassette domain-containing protein | ATP-binding cassette domain-containing protein, *Frankia* sp. EUN1f (WP_006542930.1); 86/90 | AbyF4 | AbsF4 | AbmF4 |
| *CC117_RS09785* | 334 | FadR family transcriptional regulator | FadR family transcriptional regulator, *Frankia* sp. EI5c (WP_083986722.1); 68/77 | - | - | - |
| *CC117_RS09795* | 207 | TetR/AcrR family transcriptional regulator | TetR/AcrR family transcriptional regulator, *Frankia* sp. BMG5.36 (WP_071055116.1); 77/85 | - | - | - |
| *CC117_RS09800* | 476 | MFS transporter | MFS transporter, *Frankia* sp. EUN1h (OHV31341.1); 81/87 | AbyD | AbsD | AbmD |
| *CC117_RS09805* | 225 | TetR/AcrR family transcriptional regulator | TetR/AcrR family transcriptional regulator, *Frankia* sp. EI5c (WP_066072908.1); 74/81 | AbyC | - | AbmC |
| *CC117_RS34465* | - | PKS I | type I polyketide synthase, *Umezawaea tangerina* (WP_106189546.1); 48/61 | PKS I | PKS I | PKS I |
| *CC117_RS36135* | - | PKS I | SDR family NAD(P)-dependent oxidoreductase, partial, *Streptomyces coelicolor* (WP_134115609.1); 61/72 | PKS I | PKS I | PKS I |
| *CC117_RS36140* | - | PKS I | type I polyketide synthase, *Streptomyces* sp. NBRC 109436 (WP_064455271.1); 59/70 | PKS I | PKS I | PKS I |
| *CC117_RS34485* | - | PKS I | SDR family NAD(P)-dependent oxidoreductase, *Saccharopolyspora* sp. 16K404 (WP_132624326.1); 56/67 | PKS I | PKS I | PKS I |
| *CC117_RS36145* | - | PKS I | SDR family NAD(P)-dependent oxidoreductase, partial, *Frankia symbiont of Coriaria nepalensis* (WP_131772418.1); 68/71 | PKS I | PKS I | PKS I |
| *CC117_RS36150* | - | PKS I | SDR family NAD(P)-dependent oxidoreductase, *Streptomyces* sp. MK-45 (WP_126395712.1); 54/63 | PKS I | PKS I | PKS I |
| *CC117_RS09815* | 363 | alpha/beta hydrolase | alpha/beta hydrolase, *Frankia* sp. BMG5.30 (ONH34857.1); 66/73 | AbyA5 | AbsA5 | AbmA5 |

| | | | | | | |
|---|---|---|---|---|---|---|
| *CC117_RS09820* | 229 | acyltransferase | acyltransferase, *Frankia coriariae* (WP_086055410.1); 74/83 | AbyA4 | AbsA4 | AbmA4 |
| *CC117_RS09825* | 337 | ABC transporter ATP-binding protein | ATP-binding cassette domain-containing protein, *Frankia* sp. EI5c (WP_066072919.1); 74/80 | AbyF4 | AbsF4 | AbmF4 |
| *CC117_RS09830* | 356 | ABC transporter ATP-binding protein | ABC transporter ATP-binding protein, *Frankia* sp. EAN1pec (WP_020461003.1); 72/79 | AbyF4 | AbsF4 | AbmF4 |
| *CC117_RS09835* | 278 | ABC transporter permease | ABC transporter permease, *Frankia* sp. EAN1pec (WP_020461004.1); 81/90 | AbyF3 | AbsF3 | AbmF3 |
| *CC117_RS09840* | 324 | ABC transporter permease | ABC transporter permease, *Frankia* sp. EAN1pec (WP_020461005.1); 82/89 | AbyF2 | AbsF2 | AbmF2 |
| *CC117_RS09845* | 523 | ABC transporter substrate-binding protein | ABC transporter substrate-binding protein, *Frankia* sp. EAN1pec (WP_020461006.1); 80/90 | AbyF1 | AbsF1 | AbmF1 |
| *CC117_RS09850* | 484 | LLM class flavin-dependent oxidoreductase | LLM class flavin-dependent oxidoreductase, *Millisia brevis* (WP_066905197.1); 53/66 | - | - | - |
| *CC117_RS09855* | 481 | DHA2 family efflux MFS transporter permease subunit | DHA2 family efflux MFS transporter permease subunit, *Frankia symbiont of Datisca glomerata* (WP_013873862.1); 80/86 | AbyD | AbsD | AbmD |
| *CC117_RS09860* | 354 | LLM class flavin-dependent oxidoreductase | LLM class flavin-dependent oxidoreductase, *Actinokineospora auranticolor* (WP_104480644.1); 60/71 | AbyE | AbsE | AbmE1 |
| *CC117_RS09865* | 348 | LLM class flavin-dependent oxidoreductase | LLM class flavin-dependent oxidoreductase, *Frankia* sp. EAN1pec (WP_020461009.1); 80/86 | - | - | AbmE2 |
| *CC117_RS09870* | 1148 | PKS I | type I polyketide synthase, *Actinomadura macra* (WP_067456435.1); 58/67 | AbyB1 | AbsB1 | AbmB1 |
| *CC117_RS09875* | 388 | 3-oxoacyl-ACP synthase III family protein | 3-oxoacyl-ACP synthase III family protein, *Streptomyces* sp. NRRL S-31 (WP_030750290.1); 77/86 | AbyA1 | AbsA1 | AbmA1 |
| *CC117_RS09880* | 131 | Diels-Alderase | hypothetical protein CLV40_111123, *Actinokineospora auranticolor* (PPK66159.1); 61/78 | AbyU | AbsU | AbmU |
| *CC117_RS09885* | 184 | Diels-Alderase | hypothetical protein, *Streptomyces* sp. NRRL S-31 (WP_030750286.1); 57/68 | AbyU | AbsU | AbmU |
| *CC117_RS09890* | 417 | cytochrome P450 | cytochrome P450, *Streptomyces* sp. NRRL S-31 (WP_030750284.1); 73/81 | AbyV/AvyX | AbsV/AbsX | AbmV |
| *CC117_RS09895* | 64 | ferredoxin | ferredoxin-1, *Streptomyces* sp. CC71 (KYK09758.1); 60/75 | - | AbsG1 | AbmG |
| *CC117_RS09900* | 588 | ABC transporter ATP-binding protein | dipeptide ABC transporter ATP-binding protein, *Frankia symbiont of Datisca glomerata* (WP_131768082.1); 77/82 | AbyF4 | AbsF4 | AbmF4 |
| *CC117_RS09905* | 316 | ABC transporter permease | ABC transporter permease subunit, *Frankia symbiont of Datisca glomerata* (WP_131768081.1); 80/86 | AbyF3 | AbsF3 | AbmF3 |
| *CC117_RS09910* | 322 | ABC transporter permease | ABC transporter permease subunit, *Frankia symbiont of Datisca glomerata* (WP_131768080.1); 84/92 | AbyF2 | AbsF2 | AbmF2 |
| *CC117_RS09915* | 528 | ABC transporter substrate-binding protein | ABC transporter substrate-binding protein, *Frankia symbiont of Datisca glomerata* (WP_131768079.1); 79/86 | AbyF1 | AbsF1 | AbmF1 |
| *CC117_RS09920* | 307 | taurine dioxygenase | TauD/TfdA family dioxygenase, *Frankia symbiont of Datisca glomerata* (WP_131768078.1); 79/90 | - | - | - |
| *CC117_RS09925* | 260 | AfsR/SARP family transcriptional regulator | AfsR/SARP family transcriptional regulator, *Candidatus Streptomyces philanthi* (WP_114025055.1); 52/68 | AbyI | - | AbmI |
| *CC117_RS09930* | 78 | acyl carrier protein | acyl carrier protein, *Streptomyces olindensis* (KDN76173.1); 61/74 | AbyA3 | AbsA3 | AbmA3 |
| *CC117_RS09935* | 641 | HAD-IIIC family phosphatase | HAD-IIIC family phosphatase, *Kutzneria buriramensis* (WP_116181645.1); 64/75 | AbyA2 | AbsA2 | AbmA2 |

**Table S33.** Predicted functions of ORFs in abyssomicin BGC from *Frankia* sp. Cc1.17 (NZ_MBLM01000112).

| ORF | Size (aa) | Proposed function | Closest homolog, host (protein ID); Identity/Similarity (%) | Aby homolog | Abs homolog | Abm homolog |
|---|---|---|---|---|---|---|
| *CC117_RS14595* | 142 | Diels-Alderase | hypothetical protein, *Frankia* sp. EI5c (WP_066064649.1); 97/99 | AbyU | AbsU | AbmU |
| *CC117_RS14600* | 349 | 3-oxoacyl-ACP synthase III family protein | 3-oxoacyl-ACP synthase III family protein, *Frankia* sp. EI5c (WP_066064652.1); 94/96 | AbyA1 | AbsA1 | AbmA1 |
| *CC117_RS14605* | 666 | HAD-IIIC family phosphatase | HAD-IIIC family phosphatase, *Frankia* sp. EI5c (WP_083986533.1); 88/90 | AbyA2 | AbsA2 | AbmA2 |
| *CC117_RS14610* | 209 | TetR/AcrR family transcriptional regulator | transcriptional regulator, *Frankia* sp. EI5c (OAA27545.1); 94/96 | - | - | - |
| *CC117_RS14615* | 519 | MFS transporter | drug resistance transporter, EmrB/QacA subfamily, *Frankia* sp. EI5c (OAA27546.1); 88/90 | AbyD | AbsD | AbmD |
| *CC117_RS14620* | 336 | LLM class flavin-dependent oxidoreductase | luciferase family oxidoreductase, group 1, *Frankia* sp. EI5c (OAA27547.1); 93/96 | AbyE | AbsE | AbmE1 |
| *CC117_RS14625* | 1031 | LuxR family transcriptional regulator | helix-turn-helix transcriptional regulator, *Frankia* sp. EI5c (WP_066064661.1); 88/91 | AbyH | - | AbmH |
| *CC117_RS14630* | 257 | AfsR/SARP family transcriptional regulator | AfsR/SARP family transcriptional regulator, *Frankia* sp. EI5c (WP_066064664.1); 89/94 | AbyI | - | AbmI |
| *CC117_RS14635* | 76 | acyl carrier protein | acyl carrier protein, *Frankia* sp. EI5c (WP_066064666.1); 97/97 | AbyA3 | AbsA3 | AbmA3 |
| *CC117_RS14640* | 293 | acyltransferase | acyltransferase, *Frankia* sp. EI5c (WP_066064669.1); 86/88 | AbyA4 | AbsA4 | AbmA4 |
| *CC117_RS14645* | 359 | alpha/beta hydrolase | alpha/beta hydrolase, *Frankia* sp. EI5c (WP_066064673.1); 94/95 | AbyA5 | AbsA5 | AbmA5 |
| *CC117_RS36320* | - | PKS I | type I polyketide synthase, partial, *Frankia* sp. EI5c (WP_066064676.1); 83/85 | PKS I | PKS I | PKS I |
| *CC117_RS34795* | - | PKS I | SDR family NAD(P)-dependent oxidoreductase, partial, *Frankia* sp. EI5c (WP_128423201.1); 85/88 | PKS I | PKS I | PKS I |
| *CC117_RS34800* | - | PKS I | SDR family NAD(P)-dependent oxidoreductase, partial, *Frankia* sp. EI5c (WP_128423201.1); 73/77 | PKS I | PKS I | PKS I |
| *CC117_RS34805* | - | PKS I | SDR family NAD(P)-dependent oxidoreductase, partial, *Frankia* sp. EI5c (WP_128423201.1); 94/96 | PKS I | PKS I | PKS I |
| *CC117_RS34810* | - | PKS I | SDR family NAD(P)-dependent oxidoreductase, partial, *Frankia* sp. EI5c (WP_128423201.1); 64/68 | PKS I | PKS I | PKS I |
| *CC117_RS34820* | - | PKS I | SDR family NAD(P)-dependent oxidoreductase, partial, *Frankia* sp. EI5c (WP_128423201.1); 79/82 | PKS I | PKS I | PKS I |
| *CC117_RS34825* | - | PKS I | SDR family NAD(P)-dependent oxidoreductase, partial, *Frankia* sp. EI5c (WP_128423201.1); 77/81 | PKS I | PKS I | PKS I |
| *CC117_RS36325* | - | PKS I | SDR family NAD(P)-dependent oxidoreductase, partial, *Frankia* sp. EI5c (WP_128423201.1 ); 66/74 | PKS I | PKS I | PKS I |
| *CC117_RS34840* | - | PKS I | SDR family NAD(P)-dependent oxidoreductase, partial, *Frankia* sp. EI5c (WP_128423214.1); 75/78 | PKS I | PKS I | PKS I |
| *CC117_RS34845* | - | PKS I | SDR family NAD(P)-dependent oxidoreductase, partial, *Frankia* sp. EI5c (WP_128423214.1); 78/81 | PKS I | PKS I | PKS I |
| *CC117_RS36330* | - | PKS I | KR domain-containing protein, partial, *Frankia* sp. EI5c (WP_128423221.1); 87/88 | PKS I | PKS I | PKS I |

| | | | | | | |
|---|---|---|---|---|---|---|
| *CC117_RS14660* | - | PKS I | SDR family NAD(P)-dependent oxidoreductase, partial, *Frankia* sp. EI5c (WP_066069977.1); 80/84 | PKS I | PKS I | PKS I |
| *CC117_RS36335* | - | PKS I | type I polyketide synthase, *Frankia* sp. EI5c (WP_066069974.1); 90/92 | PKS I | PKS I | PKS I |
| *CC117_RS36340* | - | PKS I | type I polyketide synthase, *Frankia* sp. EI5c (WP_066069974.1); 78/80 | PKS I | PKS I | PKS I |
| *CC117_RS36345* | - | PKS I | type I polyketide synthase, *Frankia* sp. EI5c (WP_066069974.1); 94/96 | PKS I | PKS I | PKS I |
| *CC117_RS36350* | - | PKS I | type I polyketide synthase, *Frankia* sp. EI5c (WP_066069974.1); 83/85 | PKS I | PKS I | PKS I |
| *CC117_RS36355* | - | PKS I | type I polyketide synthase, *Frankia* sp. EI5c (WP_066069974.1); 77/79 | PKS I | PKS I | PKS I |
| *CC117_RS36360* | - | PKS I | type I polyketide synthase, *Frankia* sp. EI5c (WP_066069974.1); 85/88 | PKS I | PKS I | PKS I |
| *CC117_RS14670* | - | PKS I | type I polyketide synthase, *Frankia* sp. EI5c (WP_066069971.1); 83/86 | PKS I | PKS I | PKS I |
| *CC117_RS14675* | 501 | hypothetical protein | hypothetical protein, *Frankia* sp. EI5c (WP_066069968.1); 90/93 | - | - | - |
| *CC117_RS14680* | 344 | hypothetical protein | hypothetical protein, *Frankia* sp. EI5c (WP_066069965.1); 93/96 | - | - | - |
| *CC117_RS14685* | 255 | ABC transporter permease | ABC transporter permease, *Frankia* sp. EI5c (WP_066069963.1); 97/98 | - | - | - |
| *CC117_RS14690* | 228 | ABC transporter ATP-binding protein | ABC transporter ATP-binding protein, *Frankia* sp. EI5c (WP_066069960.1); 95/97 | AbyF4 | AbsF4 | AbmF4 |
| *CC117_RS14695* | 212 | nuclear transport factor 2 family protein | nuclear transport factor 2 family protein, *Frankia* sp. EI5c (WP_066069958.1); 87/87 | - | - | - |
| *CC117_RS14700* | 474 | FAD-binding protein | FAD-dependent oxidoreductase, *Frankia* sp. EI5c (WP_066069956.1); 93/95 | - | - | - |
| *CC117_RS14705* | 214 | TetR/AcrR family transcriptional regulator | TetR/AcrR family transcriptional regulator, *Streptomyces mirabilis* (WP_075032561.1); 98/99 | - | - | - |
| *CC117_RS14710* | 245 | SDR family oxidoreductase | SDR family oxidoreductase, *Streptosporangium* sp. 'caverna' (WP_110706182.1); 99/99 | - | - | - |
| *CC117_RS14715* | 208 | NADP oxidoreductase | NADP oxidoreductase, *Streptomyces mirabilis* (WP_075032563.1); 98/99 | - | - | - |
| *CC117_RS14720* | 301 | TIGR03619 family F420-dependent LLM class oxidoreductase | TIGR03619 family F420-dependent LLM class oxidoreductase, *Streptomyces violaceoruber* (WP_030946214.1); 91/92 | - | - | - |
| *CC117_RS34855* | 277 | hypothetical protein | hypothetical protein, *Frankia* sp. EI5c (WP_066069953.1); 92/94 | - | - | - |
| *CC117_RS14730* | 206 | TetR/AcrR family transcriptional regulator | TetR/AcrR family transcriptional regulator, *Frankia* sp. EI5c (WP_083986911.1); 79/82 | - | AbsC2 | - |
| *CC117_RS14735* | 209 | hypothetical protein | hypothetical protein, *Frankia elaeagni* (WP_018637321.1); 94/97 | - | - | - |

**Table S34.** Predicted functions of ORFs surrounding AbyU homolog from *Photobacterium ganghwense* JCM 12487 (NZ_PYMI01000004.1).

| ORF | Size (aa) | Proposed function | Closest homolog, host (protein ID); Identity/Similarity (%) | Aby homolog | Abs homolog | Abm homolog |
|---|---|---|---|---|---|---|
| *C9I92_RS15395* | - | IS481 family transposase | ISSod13, transposase, *Vibrio cholerae* (SYZ80217.1); 85/91 | - | - | - |
| *C9I92_RS15400* | 157 | TetR family transcriptional regulator | TetR/AcrR family transcriptional regulator, *Vibrio proteolyticus* (WP_081693121.1); 87/94 | - | - | - |
| *C9I92_RS15405* | 439 | LLM class flavin-dependent oxidoreductase | LLM class flavin-dependent oxidoreductase, *Oceanimonas baumannii* (WP_094278553.1); 83/89 | - | - | - |
| *C9I92_RS15410* | 167 | ABC transporter ATP-binding protein | ABC transporter ATP-binding protein, *Oceanimonas baumannii* (WP_094278554.1); 58/73 | - | - | - |
| *C9I92_RS15415* | 683 | TonB-dependent siderophore receptor | TonB-dependent siderophore receptor, *Oceanimonas baumannii* (WP_094278589.1); 79/89 | - | - | - |
| *C9I92_RS15420* | 518 | MFS transporter | DHA2 family efflux MFS transporter permease subunit, *Salinivibrio* sp. YCSC6 (WP_096632410.1); 89/93 | - | - | - |
| *C9I92_RS15425* | 210 | TetR/AcrR family transcriptional regulator | TetR/AcrR family transcriptional regulator, *Vibrio gazogenes* (WP_088133275.1); 88/92 | - | - | - |
| *C9I92_RS15430* | 263 | ABC transporter ATP-binding protein | ABC transporter ATP-binding protein, *Janthinobacterium lividum* (WP_128140012.1); 61/77 | - | - | - |
| *C9I92_RS15435* | 376 | iron ABC transporter permease | iron-siderophore ABC transporter permease, *Burkholderia* sp. BDU5 (KVE40442.1); 63/80 | - | - | - |
| *C9I92_RS15440* | 164 | TonB-dependent receptor | TonB-dependent siderophore receptor, *Janthinobacterium lividum* (WP_128140009.1); 36/44 | - | - | - |
| *C9I92_RS15445* | 91 | hypothetical protein | hypothetical protein BW21_4870, *Burkholderia* sp. 2002721687 (AJY38756.1); 61/71 | - | - | - |
| *C9I92_RS15450* | 536 | siderophore biosynthesis | siderophore biosynthesis protein, *Xenorhabdus thuongxuanensis* (WP_074020647.1); 64/77 | - | - | - |
| *C9I92_RS15455* | 260 | alpha/beta fold hydrolase | thioesterase, *Xenorhabdus thuongxuanensis* (WP_074020648.1); 58/72 | - | - | - |
| *C9I92_RS15460* | 348 | pyridoxal-phosphate dependent enzyme | pyridoxal-phosphate dependent enzyme family protein, *Burkholderia* sp. ABCPW 111 (KGS01917.1); 74/84 | - | - | - |
| *C9I92_RS15465* | 191 | Diels-Alderase | hypothetical protein, *Xenorhabdus beddingii* (WP_086111861.1); 71/86 | AbyU | AbsU | AbmU |
| *C9I92_RS15470* | 965 | acyltransferase domain-containing protein | acyltransferase domain-containing protein, *Xenorhabdus beddingii* (WP_086111860.1); 63/77 | - | - | - |
| *C9I92_RS15475* | 335 | leucine dehydrogenase | leucine dehydrogenase, *Xenorhabdus beddingii* (WP_086111859.1); 68/82 | - | - | - |
| *C9I92_RS15480* | 380 | ABC transporter substrate-binding protein | ABC transporter substrate-binding protein, *Xenorhabdus beddingii* (WP_086111858.1); 67/78 | - | - | - |
| *C9I92_RS15485* | 424 | FAD-dependent oxidoreductase | FAD-dependent oxidoreductase, *Xenorhabdus thuongxuanensis* (WP_074020653.1); 74/85 | - | - | - |
| *C9I92_RS15490* | 266 | phosphonate ABC transporter, permease protein PhnE | phosphonate ABC transporter, permease protein PhnE, *Vibrio parahaemolyticus* (WP_069543805.1); 81/90 | - | - | - |
| *C9I92_RS15495* | 271 | phosphonate ABC transporter ATP-binding protein | phosphonate ABC transporter ATP-binding protein, *Vibrio campbellii* (WP_122020405.1); 83/91 | - | - | - |
| *C9I92_RS15500* | 283 | phosphonate ABC transporter substrate-binding protein | phosphate/phosphite/phosphonate ABC transporter substrate-binding protein, *Vibrio maritimus* (WP_081941364.1); 81/89 | - | - | - |

| | | | | | | |
|---|---|---|---|---|---|---|
| *C9I92_RS15505* | 363 | phosphonoacetaldehyde reductase | iron-containing alcohol dehydrogenase, *Vibrio maritimus* (WP_042495142.1); 62/76 | - | - | - |
| *C9I92_RS15510* | 299 | LysR family transcriptional regulator | LysR family transcriptional regulator, *Vibrio campbellii* (WP_045456615.1); 65/84 | - | - | - |
| *C9I92_RS15515* | 295 | DMT family transporter | DMT family transporter, *Photobacterium marinum* (WP_007465237.1); 71/82 | - | - | - |
| *C9I92_RS15520* | 675 | elongation factor G | elongation factor G, *Photobacterium sanctipauli* (WP_036815829.1); 78/89 | - | - | - |

**Table S35.** Predicted functions of ORFs surrounding AbyU homolog from *Streptomyces geranii* A301 (NZ_PJME01000012.1).

| ORF | Size (aa) | Proposed function | Closest homolog, host (protein ID); Identity/Similarity (%) | Aby homolog | Abs homolog | Abm homolog |
|---|---|---|---|---|---|---|
| *CW359_RS18380* | 1119 | peptidase | 1,4-dihydropyridine esterase, *Streptomyces* sp. L-9-10 (WP_129768931.1); 52/65 | - | - | - |
| *CW359_RS18385* | 69 | hypothetical protein | hypothetical protein, *Streptomyces* sp. NL15-2K (WP_124445691.1); 93/98 | - | - | - |
| *CW359_RS18390* | 75 | hypothetical protein | hypothetical protein, *Streptomyces* sp. NL15-2K (WP_124445690.1); 80/87 | - | - | - |
| *CW359_RS18395* | 145 | Diels-Alderase | hypothetical protein, *Streptomyces* sp. NL15-2K (WP_124445689.1); 92/97 | AbyU | AbsU | AbmU |
| *CW359_RS18400* | 354 | hypothetical protein | hypothetical protein, *Streptomyces* sp. NL15-2K (WP_124445688.1); 92/95 | - | - | - |
| *CW359_RS18405* | 355 | alpha/beta hydrolase | alpha/beta fold hydrolase, *Streptomyces* sp. NL15-2K (WP_124445687.1); 76/80 | - | - | - |
| *CW359_RS18410* | 197 | hypothetical protein | acyltransferase, *Streptomyces* sp. NL15-2K (WP_124445686.1); 79/86 | - | - | - |
| *CW359_RS18415* | 1543 | PKS I | SDR family NAD(P)-dependent oxidoreductase, *Streptomyces* sp. NL15-2K (WP_124445724.1); 82/87 | - | - | - |
| *CW359_RS18420* | 287 | thioesterase | thioesterase, *Streptomyces* sp. NL15-2K (WP_124445723.1); 79/84 | - | - | - |
| *CW359_RS18425* | 481 | multidrug efflux MFS transporter | DHA2 family efflux MFS transporter permease subunit, *Streptomyces* sp. NL15-2K (WP_124445722.1); 87/92 | - | - | - |
| *CW359_RS18430* | 1029 | non-ribosomal peptide synthetase | amino acid adenylation domain-containing protein, *Streptomyces* sp. NL15-2K (WP_124445721.1); 83/87 | - | - | - |
| *CW359_RS18435* | 273 | hypothetical protein | hypothetical protein, *Streptomyces* sp. NL15-2K (WP_124445720.1); 81/86 | - | - | - |
| *CW359_RS18440* | 248 | thioesterase | alpha/beta fold hydrolase, *Streptomyces* sp. NL15-2K (WP_124445719.1); 82/89 | - | - | - |
| *CW359_RS18445* | 347 | pyridoxal-phosphate dependent enzyme | pyridoxal-phosphate dependent enzyme, *Streptomyces* sp. NL15-2K (WP_124445718.1); 83/87 | - | - | - |
| *CW359_RS18450* | 312 | helix-turn-helix domain containing protein | IS630 family transposase, *Streptomyces* sp. NL15-2K (WP_124445717.1); 59/67 | - | - | - |
| *CW359_RS18455* | 257 | AfsR/SARP family transcriptional regulator | regulatory protein, *Streptomyces* sp. NL15-2K (GCB53297.1); 80/90 | - | - | - |
| *CW359_RS18460* | 935 | helix-turn-helix transcriptional regulator | helix-turn-helix transcriptional regulator, *Streptomyces* sp. NL15-2K (WP_124445716.1); 70/79 | - | - | - |
| *CW359_RS18465* | 105 | hypothetical protein | hypothetical protein, *Streptomyces* sp. NL15-2K (WP_124445715.1); 69/82 | - | - | - |
| *CW359_RS18470* | 196 | TetR/AcrR family transcriptional regulator | TetR/AcrR family transcriptional regulator, *Streptomyces* sp. NL15-2K (WP_124445714.1); 88/94 | - | - | - |

**Table S36.** Predicted functions of ORFs surrounding AbyU homolog from *Streptomyces griseocarneus* 132 (NZ_PENC01000003).

| ORF | Size (aa) | Proposed function | Closest homolog, host (protein ID); Identity/Similarity (%) | Aby homolog | Abs homolog | Abm homolog |
|---|---|---|---|---|---|---|
| CTZ27_RS09805 | 1351 | PKS I | SDR family NAD(P)-dependent oxidoreductase, *Streptomyces orinoci* (WP_109280288.1); 46/56 | - | - | - |
| CTZ27_RS09810 | 2970 | PKS I | type I polyketide synthase, *Saccharothrix* sp. CB00851 (WP_073887745.1); 53/63 | - | - | - |
| CTZ27_RS09815 | 413 | cytochrome P450 | cytochrome P450, *Streptomyces* sp. NRRL F-6491 (KOX15956.1); 60/72 | - | - | - |
| CTZ27_RS09820 | 149 | nuclear transport factor 2 family protein | nuclear transport factor 2 family protein, *Micromonosporaceae bacterium* CPCC 204380 (WP_117208645.1); 45/60 | - | - | - |
| CTZ27_RS09825 | 133 | Diels-Alderase | hypothetical protein, *Streptomyces* sp. MUSC 14 (WP_071375955.1); 40/55 | AbyU | AbsU | AbmU |
| CTZ27_RS09830 | 418 | crotonyl-CoA carboxylase/reductase | crotonyl-CoA carboxylase/reductase, *Saccharothrix texasensis* (ROP35577.1); 72/82 | - | - | - |
| CTZ27_RS09835 | 306 | LuxR family transcriptional regulator | hypothetical protein ADL06_14810, *Streptomyces* sp. NRRL F-6491 (KOX27187.1); 45/59 | - | - | - |
| CTZ27_RS09840 | 178 | MarR family transcriptional regulator | MarR family transcriptional regulator, *Streptomyces orinoci* (WP_109280519.1); 82/88 | - | - | - |
| CTZ27_RS09845 | 103 | hypothetical protein | hypothetical protein, *Streptomyces canus* (WP_059204811.1); 58/64 | - | - | - |
| CTZ27_RS09850 | 412 | sensor histidine kinase | sensor histidine kinase, *Streptacidiphilus rugosus* (WP_037608268.1); 76/81 | - | - | - |
| CTZ27_RS09855 | 216 | response regulator transcription factor | response regulator, *Streptomyces* sp. BK308 (WP_132857399.1); 88/93 | - | - | - |
| CTZ27_RS09860 | - | uroporphyrinogen-III synthase | uroporphyrinogen-III synthase, *Kitasatospora mediocidica* (WP_051966293.1); 80/87 | - | - | - |
| CTZ27_RS09865 | 712 | nitrite reductase | nitrite reductase, Streptomyces cattleya (WP_014151313.1); 84/87 | - | - | - |
| CTZ27_RS09870 | 405 | NAD(P)/FAD-dependent oxidoreductase | nitrite reductase, *Streptomyces malaysiense* (OIK25113.1); 71/76 | - | - | - |
| CTZ27_RS09875 | 855 | nitrite reductase large subunit | nitrite reductase large subunit, *Streptomyces cattleya* (WP_014151311.1); 83/89 | - | - | - |
| CTZ27_RS09880 | 145 | nitrite reductase small subunit NirD | nitrite reductase (NADH) small subunit, *Streptomyces misionensis* (SED96007.1); 71/84 | - | - | - |
| CTZ27_RS09885 | 458 | NarK/NasA family nitrate transporter | nitrite reductase small subunit NirD, *Streptomyces* sp. MBT76 (WP_079110603.1); 75/84 | - | - | - |
| CTZ27_RS09890 | 176 | GNAT family N-acetyltransferase | GNAT family N-acetyltransferase, *Streptomyces olivoreticuli* (WP_116209751.1); 85/92 | - | - | - |

**Table S37.** Predicted functions of ORFs in potential abyssomicin BGC from *Streptomyces griseorubiginosus* SAI-142 (NZ_RJKZ01000001.1).

| ORF | Size (aa) | Proposed function | Closest homolog, host (protein ID); Identity/Similarity (%) | Aby homolog | Abs homolog | Abm homolog |
|---|---|---|---|---|---|---|
| *EDC83_RS30485* | 912 | LuxR family transcriptional regulator | LuxR family transcriptional regulator, *Streptomyces regalis* (WP_062712128.1); 38/51 | AbyH | - | AbmH |
| *EDC83_RS30490* | 257 | AfsR/SARP family transcriptional regulator | activator protein, *Streptomyces* sp. BK438 (WP_132903690.1); 70/79 | AbyI | - | AbmI |
| *EDC83_RS30495* | 278 | thioesterase | thioesterase, *Streptomyces hoynatensis* (WP_120684679.1); 48/58 | AbyT | AbsN | AbmT |
| *EDC83_RS30500* | 343 | 3-oxoacyl-ACP synthase III family protein | 3-oxoacyl-ACP synthase III family protein, *Streptomyces paucisporeus* (WP_073498468.1); 64/77 | AbyA1 | AbsA1 | AbmA1 |
| *EDC83_RS30505* | 257 | HAD-IIIC family phosphatase | HAD-IIIC family phosphatase, *Streptomyces armeniacus* (AXK32418.1); 59/71 | AbyA2 | AbsA2 | AbmA2 |
| *EDC83_RS30510* | 75 | acyl carrier protein | acyl carrier protein, *Actinomadura* sp. 7K507 (WP_132147234.1); 57/82 | AbyA3 | AbsA3 | AbmA3 |
| *EDC83_RS30515* | 268 | acyltransferase | acyltransferase, *Streptomyces kanamyceticus* (WP_055549000.1); 60/77 | AbyA4 | AbsA4 | AbmA4 |
| *EDC83_RS30520* | 372 | alpha/beta hydrolase | alpha/beta hydrolase, *Streptomyces* sp. NRRL F-525 (WP_033287161.1); 52/64 | AbyA5 | AbsA5 | AbmA5 |
| *EDC83_RS30525* | 248 | hypothetical protein | S-adenosyl methyltransferase*, Actinomadura umbrina* (REE95466.1); 51/69 | - | - | - |
| *EDC83_RS30530* | 6378 | PKS I | type I polyketide synthase, *Actinomadura macra* (WP_067456430.1); 55/66 | AbyB1 | AbsB1 | AbmB1 |
| *EDC83_RS30535* | 2213 | PKS I | type I polyketide synthase, *Streptomyces formicae* (WP_098241246.1); 56/67 | AbyB2 | AbsB2 | AbmB2 |
| *EDC83_RS30540* | 1554 | PKS I | Erythronolide synthase, *Streptomyces malaysiensis* (PNG90733.1); 49/61 | AbyB3 | AbsB3 | AbmB3 |
| *EDC83_RS30545* | 581 | hypothetical protein | 2-polyprenyl-6-methoxyphenol hydroxylase, *Kibdelosporangium aridum* (WP_037262367.1); 47/59 | - | - | - |
| *EDC83_RS30550* | 61 | hypothetical protein | hypothetical protein, *Streptomyces* sp. 57 (WP_121408886.1); 56/60 | - | - | - |
| *EDC83_RS30555* | 348 | methyltransferase | methyltransferase, *Nonomuraea wenchangensis* (WP_091076886.1); 48/59 | - | - | - |
| *EDC83_RS30560* | 137 | Diels-Alderase | hypothetical protein, *Streptomyces paucisporeus* (WP_073498104.1); 44/58 | AbyU | AbsU | AbmU |
| *EDC83_RS30565* | 78 | ferredoxin | ferredoxin, *Streptomyces* sp. GSSD-12 (WP_114664483.1); 59/70 | - | AbsG1 | AbmG |
| *EDC83_RS30570* | 369 | cytochrome P450 | cytochrome P450, *Streptomyces paucisporeus* (WP_073498464.1); 56/71 | AbyX/AbyV | AbsV/AbsX | AbmV |
| *EDC83_RS30575* | 501 | MFS transporter | DHA2 family efflux MFS transporter permease subunit, *Streptomyces formicae* (WP_098241238.1); 52/68 | AbyD | AbsD | AbmD |
| *EDC83_RS30580* | 174 | TetR/AcrR family transcriptional regulator | TetR family transcriptional regulator, *Nonomuraea* sp. CH32 (WP_132623500.1); 53/67 | AbyC | - | AbmC |
| *EDC83_RS30585* | 45 | nucleotidyl transferase AbiEii/AbiGii toxin family protein | nucleotidyl transferase AbiEii/AbiGii toxin family protein, *Streptomyces* sp. NRRL F-525 (WP_033282780.1); 82/86 | - | - | - |
| *EDC83_RS30590* | 174 | hypothetical protein | - | - | - | - |
| *EDC83_RS30595* | 455 | L-serine ammonia-lyase | L-serine ammonia-lyase, *Streptomyces* sp. 351MFTsu5.1 (WP_020139905.1); 98/99 | - | - | - |
| *EDC83_RS30600* | 421 | serine hydroxymethyltransferase | serine hydroxymethyltransferase, *Streptomyces* sp. NRRL B-24085 (WP_053846967.1); 98/98 | - | - | - |

| | | | | | | |
|---|---|---|---|---|---|---|
| *EDC83_RS30605* | 125 | glycine cleavage system protein GcvH | glycine cleavage system H protein, *Streptomyces sviceus* ATCC 29083 (EDY54000.1); 100/100 | - | - | - |
| *EDC83_RS30610* | 371 | glycine cleavage system aminomethyltransferase GcvT | glycine cleavage system aminomethyltransferase GcvT, *Streptomyces* sp. NRRL B-24085 (WP_053846965.1); 95/97 | - | - | - |
| *EDC83_RS30615* | 222 | ATP-binding protein | AAA domain-containing protein, *Streptomyces* sp. BK205 (TCR18702.1); 95/96 | - | - | - |
| *EDC83_RS30620* | 263 | enhanced serine sensitivity protein SseB | enhanced serine sensitivity protein SseB, *Streptomyces* sp. BK205 (WP_132837573.1); 99/100 | - | - | - |
| *EDC83_RS30625* | 264 | enhanced serine sensitivity protein SseB | type III secretion system (T3SS) SseB-like protein, *Streptomyces* sp. BK205 (TCR18704.1); 97/98 | - | - | - |
| *EDC83_RS30630* | 332 | ABC transporter permease | ABC transporter permease, *Streptomyces* sp. W SAI-097 (WP_123991755.1); 99/99 | AbyF3 | AbsF3 | AbmF3 |
| *EDC83_RS30635* | 582 | ABC transporter substrate-binding protein | ABC transporter substrate-binding protein, *Streptomyces mirabilis* (WP_037711494.1); 98/98 | - | - | - |
| *EDC83_RS30640* | 335 | ABC transporter permease | ABC transporter permease, *Streptomyces* sp. W SAI-097 (WP_123991757.1); 99/100 | AbyF2 | AbsF2 | AbmF2 |
| *EDC83_RS30645* | 363 | ABC transporter ATP-binding protein | ABC transporter ATP-binding protein, *Streptomyces* sp. NRRL B-24085 (WP_053846961.1); 99/99 | AbyF4 | AbsF4 | AbmF4 |
| *EDC83_RS30650* | 446 | dipeptide ABC transporter ATP-binding protein | dipeptide ABC transporter ATP-binding protein, *Streptomyces* sp. NRRL B-3229 (WP_037819546.1); 86/88 | AbyF4 | AbsF4 | AbmF4 |

**Table S38.** Predicted functions of ORFs in abyssomicin BGC from *Herbidospora daliensis* NBRC 106372 (NZ_BBXF01000001.1).

| **ORF** | **Size (aa)** | **Proposed function** | **Closest homolog, host (protein ID); Identity/Similarity (%)** | **Aby homolog** | **Abs homolog** | **Abm homolog** |
|---|---|---|---|---|---|---|
| *AW274_RS02325* | 285 | LLM class F420-dependent oxidoreductase | LLM class F420-dependent oxidoreductase, *Herbidospora cretacea* (WP_030454059.1); 93/95 | - | - | AbmE2 |
| *AW274_RS02330* | 257 | AfsR/SARP family transcriptional regulator | activator protein, *Herbidospora sakaeratensis* (WP_062343034.1); 96/98 | AbyI | - | AbmI |
| *AW274_RS02335* | 248 | thioesterase | thioesterase, *Herbidospora sakaeratensis* (WP_062343032.1); 90/92 | AbyT | AbsN | AbmT |
| *AW274_RS02340* | 897 | LuxR family transcriptional regulator | helix-turn-helix transcriptional regulator, *Herbidospora sakaeratensis* (WP_062343030.1); 92/93 | AbyH | - | AbmH |
| *AW274_RS02345* | 836 | RHS repeat protein | RHS repeat protein, *Herbidospora sakaeratensis* (WP_062343029.1); 91/94 | AbyK | - | - |
| *AW274_RS02350* | 134 | Diels-Alderase | hypothetical protein, *Herbidospora sakaeratensis* (WP_062343027.1); 99/100 | AbyU | AbsU | AbmU |
| *AW274_RS02355* | 340 | 3-oxoacyl-ACP synthase III family protein | 3-oxoacyl-ACP synthase III family protein, *Herbidospora sakaeratensis* (WP_062343025.1); 98/98 | AbyA1 | AbsA1 | AbmA1 |
| *AW274_RS02360* | 614 | HAD-IIIC family phosphatase | HAD-IIIC family phosphatase, *Herbidospora sakaeratensis* (WP_062343023.1); 94/96 | AbyA2 | AbsA2 | AbmA2 |
| *AW274_RS02370* | 322 | acyltransferase | acyltransferase, *Herbidospora sakaeratensis* (WP_062343019.1); 93/95 | AbyA4 | AbsA4 | AbmA4 |
| *AW274_RS02375* | 347 | alpha/beta hydrolase | alpha/beta hydrolase, *Herbidospora sakaeratensis* (WP_062343017.1); 95/97 | AbyA5 | AbsA5 | AbmA5 |
| *AW274_RS02380* | 213 | TetR/AcrR family transcriptional regulator | TetR/AcrR family transcriptional regulator, *Herbidospora yilanensis* (WP_062349591.1); 98/99 | AbyC | - | AbmC |
| *AW274_RS02385* | 476 | DHA2 family efflux MFS transporter permease subunit | DHA2 family efflux MFS transporter permease subunit, *Herbidospora sakaeratensis* (WP_062343013.1); 97/98 | AbyD | AbsD | AbmD |
| *AW274_RS02390* | 335 | LLM class flavin-dependent oxidoreductase | LLM class flavin-dependent oxidoreductase, *Herbidospora sakaeratensis* (WP_062343011.1); 96/98 | AbyE | AbsE | AbmE1 |
| *AW274_RS02395* | 543 | ABC transporter substrate-binding protein | ABC transporter substrate-binding protein, *Herbidospora sakaeratensis* (WP_062343009.1); 97/98 | AbyF1 | AbsF1 | AbmF1 |
| *AW274_RS02400* | 317 | ABC transporter permease | ABC transporter permease, *Herbidospora sakaeratensis* (WP_083977791.1); 98/99 | AbyF2 | AbsF2 | AbmF2 |
| *AW274_RS02405* | 268 | ABC transporter permease | ABC transporter permease, *Herbidospora cretacea* (WP_034385023.1); 95/98 | AbyF3 | AbsF3 | AbmF3 |
| *AW274_RS02410* | 529 | ABC transporter ATP-binding protein | ABC transporter ATP-binding protein, *Herbidospora sakaeratensis* (WP_062343005.1); 95/96 | AbyF4 | AbsF4 | AbmF4 |
| *AW274_RS02415* | 376 | acyltransferase | acyltransferase, *Herbidospora sakaeratensis* (WP_062343003.1); 93/94 | - | AbsI | - |
| *AW274_RS02420* | 393 | cytochrome P450 | cytochrome P450, *Herbidospora sakaeratensis* (WP_062343001.1); 97/98 | AbyV/AbyX | AbsV/AbsX | AbmV |
| *AW274_RS02425* | 63 | ferredoxin | ferredoxin, *Herbidospora sakaeratensis* (WP_062343000.1); 97/98 | - | AbsG2/AbsG1 | AbmG |
| *AW274_RS02430* | 286 | alpha/beta hydrolase | alpha/beta hydrolase, *Herbidospora yilanensis* (WP_062349582.1); 91/95 | - | - | - |
| *AW274_RS38765* | - | PKS I | type I polyketide synthase, *Herbidospora mongoliensis* (WP_066363856.1); 94/95 | AbyB1 | AbsB1 | AbmB1 |

| | | | | | | |
|---|---|---|---|---|---|---|
| *AW274_RS38770* | | PKS I | type I polyketide synthase, *Herbidospora mongoliensis* (WP_066363856.1); 78/79 | | | |
| *AW274_RS38775* | | PKS I | type I polyketide synthase, *Herbidospora mongoliensis* (WP_066363856.1); 86/89 | | | |
| *AW274_RS38780* | | PKS I | type I polyketide synthase, *Herbidospora mongoliensis* (WP_066363856.1); 86/90 | | | |
| *AW274_RS38785* | | PKS I | type I polyketide synthase, *Herbidospora mongoliensis* (WP_066363856.1); 73/82 | | | |
| *AW274_RS38790* | | PKS I | - | | | |
| *AW274_RS38795* | | PKS I | type I polyketide synthase, *Herbidospora mongoliensis* (WP_066363856.1); 75/82 | | | |
| *AW274_RS38800* | | PKS I | type I polyketide synthase, *Herbidospora mongoliensis* (WP_066363856.1); 85/90 | | | |
| *AW274_RS38805* | | PKS I | type I polyketide synthase, *Herbidospora mongoliensis* (WP_066363856.1); 82/87 | | | |
| *AW274_RS38810* | | PKS I | type I polyketide synthase, *Herbidospora mongoliensis* (WP_066363856.1); 78/86 | | | |
| *AW274_RS38815* | | PKS I | type I polyketide synthase, *Herbidospora mongoliensis* (WP_066363856.1); 74/81 | | | |
| *AW274_RS38820* | | PKS I | type I polyketide synthase, *Herbidospora sakaeratensis* (WP_062342995.1); 88/90 | | | |
| *AW274_RS02440* | 3774 | PKS I | SDR family NAD(P)-dependent oxidoreductase, *Streptomyces* sp. BK438 (WP_132903672.1); 63/71 | AbyB2 | AbsB2 | AbmB2 |
| *AW274_RS02445* | 1019 | PKS I | type I polyketide synthase, *Herbidospora sakaeratensis* (WP_062333186.1); 90/91 | AbyB3 | AbsB3 | AbmB3 |
| *AW274_RS02450* | 386 | cytochrome P450 | cytochrome P450, *Herbidospora sakaeratensis* (WP_062333189.1); 94/97 | AbyX/AbyV | AbsV/AbsX | AbmV |

**Table S39.** Predicted functions of ORFs in abyssomicin BGC from *Herbidospora mongoliensis* NBRC 105882 (NZ_BBXD01000011.1).

| ORF | Size (aa) | Proposed function | Closest homolog, host (protein ID); Identity/Similarity (%) | Aby homolog | Abs homolog | Abm homolog |
|---|---|---|---|---|---|---|
| *AW272_RS12035* | 501 | ABC transporter ATP-binding protein | ABC transporter ATP-binding protein, *Herbidospora yilanensis* (WP_062349605.1); 79/84 | AbyF4 | AbsF4 | AbmF4 |
| *AW272_RS12040* | 278 | ABC transporter permease | ABC transporter permease, *Herbidospora yilanensis* (WP_063910013.1); 94/95 | AbyF3 | AbsF3 | AbmF3 |
| *AW272_RS12045* | 318 | ABC transporter permease | ABC transporter permease, *Herbidospora yilanensis* (WP_062349604.1); 94/96 | AbyF2 | AbsF2 | AbmF2 |
| *AW272_RS12050* | 530 | hypothetical protein | ABC transporter substrate-binding protein, *Herbidospora yilanensis* (WP_062349603.1); 89/92 | AbyF1 | AbsF1 | AbmF1 |
| *AW272_RS12055* | 252 | AfsR/SARP family transcriptional regulator | activator protein, *Herbidospora yilanensis* (WP_062349599.1); 88/91 | AbyI | - | AbmI |
| *AW272_RS12060* | 249 | thioesterase | thioesterase, *Herbidospora yilanensis* (WP_083949863.1); 88/89 | AbyT | AbsN | AbmT |
| *AW272_RS40525* | 1708 | hypothetical protein | RHS repeat protein, *Herbidospora yilanensis* (WP_062349597.1); 86/90 | AbyK | - | - |
| *AW272_RS12075* | 134 | Diels-Alderase | hypothetical protein, *Herbidospora* (WP_030454064.1); 97/99 | AbyU | AbsU | AbmU |
| *AW272_RS12080* | 340 | 3-oxoacyl-ACP synthase III family protein | 3-oxoacyl-ACP synthase III family protein, *Herbidospora daliensis* (WP_062428856.1); 94/96 | AbyA1 | AbsA1 | AbmA1 |
| *AW272_RS12085* | 630 | HAD-IIIC family phosphatase | HAD-IIIC family phosphatase, *Herbidospora sakaeratensis* (WP_062343023.1); 89/91 | AbyA2 | AbsA2 | AbmA2 |
| *AW272_RS12090* | 75 | acyl carrier protein | hypothetical protein, *Herbidospora yilanensis* (WP_062349594.1); 92/97 | AbyA3 | AbsA3 | AbmA3 |
| *AW272_RS12095* | 247 | acyltransferase | Acyltransferase, *Herbidospora yilanensis* (WP_062349593.1); 94/96 | AbyA4 | AbsA4 | AbmA4 |
| *AW272_RS12100* | 347 | alpha/beta hydrolase | alpha/beta hydrolase, *Herbidospora yilanensis* (WP_062349592.1); 92/95 | AbyA5 | AbsA5 | AbmA5 |
| *AW272_RS12105* | 217 | TetR/AcrR family transcriptional regulator | TetR/AcrR family transcriptional regulator, *Herbidospora sakaeratensis* (WP_062343015.1); 94/96 | AbyC | - | AbmC |
| *AW272_RS12110* | 475 | DHA2 family efflux MFS transporter permease subunit | DHA2 family efflux MFS transporter permease subunit, *Herbidospora yilanensis* (WP_062349590.1); 92/95 | AbyD | AbsD | AbmD |
| *AW272_RS12115* | 335 | LLM class flavin-dependent oxidoreductase | LLM class flavin-dependent oxidoreductase, *Herbidospora sakaeratensis* (WP_062343011.1); 92/95 | AbyE | AbsE | AbmE1 |
| *AW272_RS12120* | 544 | ABC transporter substrate-binding protein | ABC transporter substrate-binding protein, *Herbidospora daliensis* (WP_062428872.1); 90/94 | AbyF1 | AbsF1 | AbmF1 |
| *AW272_RS12125* | 317 | ABC transporter permease | ABC transporter permease, *Herbidospora yilanensis* (WP_062349773.1); 93/96 | AbyF2 | AbsF2 | AbmF2 |
| *AW272_RS12130* | 268 | ABC transporter permease | ABC transporter permease, *Herbidospora cretacea* (WP_034385023.1); 92/95 | AbyF3 | AbsF3 | AbmF3 |
| *AW272_RS12135* | 529 | ABC transporter ATP-binding protein | ABC transporter ATP-binding protein, *Herbidospora cretacea* (WP_030454076.1); 90/94 | AbyF4 | AbsF4 | AbmF4 |
| *AW272_RS12140* | 388 | acyltransferase | Acyltransferase, *Herbidospora sakaeratensis* (WP_062343003.1); 90/93 | - | AbsI | - |
| *AW272_RS12145* | 393 | cytochrome P450 | cytochrome P450, *Herbidospora sakaeratensis* (WP_062343001.1); 95/97 | AbyV/AbyX | AbsV/AbsX | AbmV |
| *AW272_RS12150* | 63 | ferredoxin | Ferredoxin, *Herbidospora sakaeratensis* (WP_062343000.1); 95/98 | - | AbsG2/AbsG1 | AbmG |

| | | | | | | |
|---|---|---|---|---|---|---|
| *AW272_RS12155* | 288 | alpha/beta hydrolase | alpha/beta hydrolase, *Herbidospora yilanensis* (WP_062349582.1); 91/93 | - | - | - |
| *AW272_RS12160* | 6103 | PKS I | type I polyketide synthase, *Streptomyces fragilis* (WP_108952947.1); 59/67 | AbyB1 | AbsB1 | AbmB1 |
| *AW272_RS12165* | 3854 | PKS I | type I polyketide synthase, *Herbidospora daliensis* (WP_062428883.1); 80/84 | AbyB2 | AbsB2 | AbmB2 |
| *AW272_RS12170* | 1070 | PKS I | type I polyketide synthase, *Herbidospora daliensis* (WP_062428886.1); 81/86 | AbyB3 | AbsB3 | AbmB3 |
| *AW272_RS12175* | 386 | cytochrome P450 | cytochrome P450, *Herbidospora sakaeratensis* (WP_062333189.1); 91/95 | AbyX/AbyV | AbsV/AbsX | AbmV |

**Table S40.** Predicted functions of ORFs in abyssomicin BGC from *Herbidospora sakaeratensis* NBRC 102641 (NZ_BBXC01000032).

| **ORF** | **Size (aa)** | **Proposed function** | **Closest homolog, host (protein ID); Identity/Similarity (%)** | **Aby homolog** | **Abs homolog** | **Abm homolog** |
|---|---|---|---|---|---|---|
| *AW271_RS37390* | - | PKS I | type I polyketide synthase, *Herbidospora daliensis* (WP_062428883.1); 89/91 | PKS I | PKS I | PKS I |
| *AW271_RS37395* | 5982 | PKS I | SDR family NAD(P)-dependent oxidoreductase, *Streptomyces* sp. 57 (WP_121408907.1); 57/65 | AbyB1 | AbsB1 | AbmB1 |
| *AW271_RS37400* | 286 | alpha/beta hydrolase | alpha/beta hydrolase, *Herbidospora yilanensis* (WP_062349582.1);94/96 | - | - | - |
| *AW271_RS37405* | 63 | ferredoxin | ferredoxin, *Herbidospora yilanensis* (WP_062349583.1); 98/100 | - | AbsG2/AbsG1 | AbmG |
| *AW271_RS37410* | 393 | cytochrome P450 | cytochrome P450, *Herbidospora daliensis* (WP_062428876.1); 97/98 | AbyV/AbyX | AbsV/AbsX | AbmV |
| *AW271_RS37415* | 380 | acyltransferase | acyltransferase, *Herbidospora mongoliensis* (WP_066363851.1); 90/93 | - | AbsI | - |
| *AW271_RS37420* | 534 | ABC transporter ATP-binding protein | ABC transporter ATP-binding protein, *Herbidospora daliensis* (WP_062428874.1); 96/98 | AbyF4 | AbsF4 | AbmF4 |
| *AW271_RS37425* | 268 | ABC transporter permease | ABC transporter permease, *Herbidospora cretacea* (WP_034385023.1); 96/98 | AbyF3 | AbsF3 | AbmF3 |
| *AW271_RS37430* | 317 | ABC transporter permease | ABC transporter permeasem, *Herbidospora daliensis* (WP_062430092.1); 98/99 | AbyF2 | AbsF2 | AbmF2 |
| *AW271_RS37435* | 543 | ABC transporter substrate-binding protein | ABC transporter substrate-binding protein, *Herbidospora daliensis* (WP_062428872.1); 97/98 | AbyF1 | AbsF1 | AbmF1 |
| *AW271_RS37440* | 335 | LLM class flavin-dependent oxidoreductase | LLM class flavin-dependent oxidoreductase, *Herbidospora cretacea* (WP_030454072.1); 98/98 | AbyE | AbsE | AbmE1 |
| *AW271_RS37445* | 476 | DHA2 family efflux MFS transporter permease subunit | DHA2 family efflux MFS transporter permease subunit, *Herbidospora daliensis* (WP_062428868.1); 97/98 | AbyD | AbsD | AbmD |
| *AW271_RS37450* | 217 | TetR/AcrR family transcriptional regulator | TetR/AcrR family transcriptional regulator, *Herbidospora yilanensis* (WP_062349591.1); 98/99 | AbyC | - | AbmC |
| *AW271_RS37455* | 347 | alpha/beta hydrolase | alpha/beta hydrolase, *Herbidospora daliensis* (WP_062428864.1); 95/97 | AbyA5 | AbsA5 | AbmA5 |
| *AW271_RS37460* | 244 | acyltransferase | acyltransferase, *Herbidospora yilanensis* (WP_062349593.1); 95/96 | AbyA4 | AbsA4 | AbmA4 |
| *AW271_RS37465* | 75 | hypothetical protein | acyl carrier protein, *Herbidospora mongoliensis* (WP_066363836.1); 92/97 | AbyA3 | AbsA3 | AbmA3 |
| *AW271_RS37470* | 630 | HAD-IIIC family phosphatase | HAD-IIIC family phosphatase, *Herbidospora yilanensis* (WP_062349595.1); 93/96 | AbyA2 | AbsA2 | AbmA2 |
| *AW271_RS37475* | 340 | 3-oxoacyl-ACP synthase III family protein | 3-oxoacyl-ACP synthase III family protein, *Herbidospora daliensis* (WP_062428856.1); 98/98 | AbyA1 | AbsA1 | AbmA1 |
| *AW271_RS37480* | 134 | Diels-Alderase | hypothetical protein, *Herbidospora daliensis* (WP_062428853.1); 99/100 | AbyU | AbsU | AbmU |
| *AW271_RS37485* | 836 | RHS repeat protein | RHS repeat protein, *Herbidospora daliensis* (WP_062428851.1); 91/94 | AbyK | - | - |
| *AW271_RS37490* | 898 | helix-turn-helix transcriptional regulator | LuxR family transcriptional regulator, *Herbidospora daliensis* (WP_062428849.1); 92/93 | AbyH | - | AbmH |
| *AW271_RS37495* | 251 | thioesterase | thioesterase, *Herbidospora yilanensis* (WP_083949863.1); 91/92 | AbyT | AbsN | AbmT |
| *AW271_RS37500* | 257 | AfsR/SARP family transcriptional regulator | activator protein, *Herbidospora yilanensis* (WP_062349599.1); 98/98 | AbyI | - | AbmI |

| | | | | | | |
|---|---|---|---|---|---|---|
| *AW271_RS37505* | 285 | LLM class F420-dependent oxidoreductase | LLM class F420-dependent oxidoreductase, *Herbidospora cretacea* (WP_030454059.1); 93/95 | - | - | AbmE2 |

**Table S41.** Predicted functions of ORFs in potential abyssomicin BGC from *Actinokineospora inagensis* DSM 44258 (NZ_AXWW01000024.1).

| ORF | Size (aa) | Proposed function | Closest homolog, host (protein ID); Identity/Similarity (%) | Aby homolog | Abs homolog | Abm homolog |
|---|---|---|---|---|---|---|
| *H504_RS0110020* | 257 | AfsR/SARP family transcriptional regulator | activator protein, *Amycolatopsis* sp. CA-126428 (WP_103341808.1); 76/86 | AbyI | - | AbmI |
| *H504_RS0110025* | 409 | cytochrome P450 | cytochrome P450, *Amycolatopsis* sp. CA-126428 (WP_103341807.1); 83/87 | AbyX/AbyV | AbsV/AbsX | AbmV |
| *H504_RS0110030* | 64 | ferredoxin | ferredoxin-1, *Amycolatopsis* sp. CA-126428 (WP_103341806.1); 66/76 | - | AbsG1/AbsG2 | AbmG |
| *H504_RS34265* | 498 | multidrug efflux MFS transporter | DHA2 family efflux MFS transporter permease subunit, *Amycolatopsis* sp. CA-126428 (WP_103341805.1); 70/79 | AbyD | AbsD | AbmD |
| *H504_RS34270* | 155 | nuclear transport factor 2 family protein | nuclear transport factor 2 family protein, *Amycolatopsis* sp. CA-126428 (WP_103341804.1); 69/81 | - | - | - |
| *H504_RS34275* | 267 | helix-turn-helix transcriptional regulator | transcriptional regulator, TetR family, *Frankia* sp. CcI6 (ETA00366.1); 56/70 | - | AbsC2 | - |
| *H504_RS0110050* | 412 | cytochrome P450 | cytochrome P450, *Amycolatopsis* sp. CA-126428 (WP_103341801.1); 65/80 | AbyX/AbyV | AbsV/AbsX | AbmV |
| *H504_RS0110055* | 3768 | PKS I | type I polyketide synthase, *Streptomyces* sp. JV178 (WP_099966065.1); 54/64 | PKS I | PKS I | PKS I |
| *H504_RS0110060* | 6097 | PKS I | type I polyketide synthase, *Actinomadura macra* (WP_067456430.1); 53/64 | PKS I | PKS I | PKS I |
| *H504_RS0110065* | 167 | Diels-Alderase | hypothetical protein, *Amycolatopsis* sp. CA-126428 (WP_103337400.1); 68/78 | AbyU | AbsU | AbmU |
| *H504_RS34280* | 663 | MFS transporter | DHA2 family efflux MFS transporter permease subunit, *Amycolatopsis* sp. CA-126428 (WP_103337401.1); 86/92 | AbyD | AbsD | AbmD |
| *H504_RS0110075* | 1381 | PKS I | type I polyketide synthase, *Amycolatopsis* sp. CA-126428 (WP_103337398.1); 65/74 | PKS I | PKS I | PKS I |
| *H504_RS34285* | 3465 | PKS I | SDR family NAD(P)-dependent oxidoreductase, *Streptomyces olivoreticuli* (WP_116210514.1); 54/66 | PKS I | PKS I | PKS I |
| *H504_RS0110090* | 344 | 3-oxoacyl-ACP synthase III family protein | 3-oxoacyl-ACP synthase III family protein, *Amycolatopsis* sp. CA-126428 (WP_103342534.1); 82/90 | AbyA1 | AbsA1 | AbmA1 |
| *H504_RS34290* | 634 | HAD-IIIC family phosphatase | HAD-IIIC family phosphatase, *Amycolatopsis* sp. CA-126428 (WP_103342533.1); 75/83 | AbyA2 | AbsA2 | AbmA2 |
| *H504_RS0110100* | 74 | hypothetical protein | acyl carrier protein, *Amycolatopsis* sp. CA-126428 (WP_103342532.1); 64/79 | AbyA3 | AbsA3 | AbmA3 |
| *H504_RS34295* | 259 | acyltransferase | acyltransferase, *Amycolatopsis* sp. CA-126428 (WP_103342540.1); 81/88 | AbyA4 | AbsA4 | AbmA4 |
| *H504_RS34300* | 372 | alpha/beta hydrolase | alpha/beta hydrolase, *Amycolatopsis* sp. CA-126428 (WP_103342531.1); 68/76 | AbyA5 | AbsA5 | AbmA5 |

**Table S42.** Predicted functions of ORFs in potential BGC from *Streptomyces iranensis* DSM 41954 (NZ_LK022848).

| **ORF** | **Size (aa)** | **Proposed function** | **Closest homolog, host (protein ID); Identity/Similarity (%)** | **Aby homolog** | **Abs homolog** | **Abm homolog** |
|---|---|---|---|---|---|---|
| *SIRAN_RS44090* | 227 | TetR/AcrR family transcriptional regulator | TetR/AcrR family transcriptional regulator, *Streptomyces* sp. 11-1-2 (WP_119984664.1); 89/91 | - | - | - |
| *SIRAN_RS44095* | 319 | NAD(P)-dependent alcohol dehydrogenase | NAD(P)-dependent alcohol dehydrogenase, *Streptomyces rhizosphaericus* (WP_086879937.1); 94/96 | - | - | - |
| *SIRAN_RS44100* | 770 | helicase | helicase, *Streptomyces rhizosphaericus* (WP_086879936.1); 97/98 | - | - | - |
| *SIRAN_RS44105* | 505 | glycoside hydrolase | glycoside hydrolase, *Streptomyces rapamycinicus* (WP_020874169.1); 97/98 | - | - | - |
| *SIRAN_RS44110* | 144 | hypothetical protein | SRPBCC family protein, *Streptomyces hygroscopicus* (WP_030824776.1); 92/96 | - | - | - |
| *SIRAN_RS44115* | 649 | beta-N-acetylglucosaminidase | hyaluronidase, *Streptomyces rapamycinicus* (WP_020874171.1); 95/96 | - | - | - |
| *SIRAN_RS44120* | 440 | FAD-dependent oxidoreductase | FAD-binding monooxygenase, *Streptomyces* sp. WAC05858 (WP_125755201.1); 90/94 | - | - | - |
| *SIRAN_RS44125* | 894 | glucan biosynthesis protein | glucan biosynthesis protein, *Streptomyces rhizosphaericus* (WP_086879933.1); 97/97 | - | - | - |
| *SIRAN_RS51845* | 154 | nuclear transport factor 2 family protein | nuclear transport factor 2 family protein, *Saccharothrix syringae* (WP_051765870.1); 87/92 | - | - | - |
| *SIRAN_RS51850* | 277 | NAD(P)-dependent oxidoreductase | NAD(P)-dependent oxidoreductase, *Saccharothrix syringae* (WP_051765824.1); 87/93 | - | - | - |
| *SIRAN_RS51855* | 140 | Diels-Alderase | hypothetical protein, *Streptomyces cattleya* (WP_014140910.1); 91/95 | AbyU | AbsU | AbmU |
| *SIRAN_RS44130* | 119 | nuclear transport factor 2 family protein | nuclear transport factor 2 family protein, *Streptomyces cattleya* (WP_014627233.1); 93/97 | - | - | - |
| *SIRAN_RS44135* | 417 | cytochrome P450 | cytochrome P450, *Saccharothrix syringae* (WP_033431229.1); 93/97 | - | - | - |
| *SIRAN_RS44140* | 6154 | PKS I | type I polyketide synthase, *Streptomyces cattleya* (WP_014140913.1); 89/93 | - | - | - |
| *SIRAN_RS44145* | 159 | PKS I | type I polyketide synthase, *Saccharothrix syringae* (WP_084716421.1); 89/90 | - | - | - |
| *SIRAN_RS51860* | - | PKS I | type I polyketide synthase, *Saccharothrix syringae* (WP_084716421.1); 89/92 | - | - | - |
| *SIRAN_RS44165* | - | pyridoxamine 5'-phosphate oxidase family protein | hypothetical protein, *Saccharothrix syringae* (WP_033431232.1); 88/94 | - | - | - |
| *SIRAN_RS44170* | - | PKS I | type I polyketide synthase, *Streptomyces cattleya* (WP_014140917.1); 92/95 | - | - | - |
| *SIRAN_RS44175* | - | PKS I | acyltransferase domain-containing protein, *Streptomyces cattleya* (WP_014140918.1); 91/93 | - | - | - |
| *SIRAN_RS51865* | - | hypothetical protein | transcriptional regulator, partial, *Streptomyces milbemycinicus* (WP_086861070.1); 60/67 | - | - | - |
| *SIRAN_RS44180* | 277 | SDR family oxidoreductase | SDR family oxidoreductase, *Streptomyces cattleya* (WP_014140897.1); 94/97 | - | - | - |
| *SIRAN_RS44185* | 393 | LLM class flavin-dependent oxidoreductase | LLM class flavin-dependent oxidoreductase, *Streptomyces cattleya* (WP_014140896.1); 95/97 | - | - | - |
| *SIRAN_RS44190* | 68 | thioesterase | thioesterase, *Streptomyces cattleya* (WP_014140907.1); 96/98 | - | - | - |

| | | | | | | |
|---|---|---|---|---|---|---|
| *SIRAN_RS44195* | 408 | cytochrome P450 | cytochrome P450, *Streptomyces cattleya* (WP_014627229.1); 88/93 | - | - | - |
| *SIRAN_RS51870* | 125 | hypothetical protein | RNA-directed DNA polymerase, *Streptomyces* sp. SYSU K10008 (WP_128381944.1); 87/89 | - | - | - |
| *SIRAN_RS44205* | 933 | helix-turn-helix transcriptional regulator | LuxR family transcriptional regulator, *Saccharothrix syringae* (WP_051765876.1); 88/91 | - | - | - |
| *SIRAN_RS53370* | 83 | hypothetical protein | hypothetical protein, *Streptomyces hygroscopicus* (WP_078640395.1); 71/81 | - | - | - |

**Table S43.** Predicted functions of ORFs in potential abyssomicin BGC from *Lentzea kentuckyensis* NRRL B-24416 (NZ_MUYM01000068 and NZ_MUYM01000065).

| ORF | Size (aa) | Proposed function | Closest homolog, host (protein ID); Identity/Similarity (%) | Aby homolog | Abs homolog | Abm homolog |
|---|---|---|---|---|---|---|
| *B0F77_RS29650* | 306 | DUF418 domain-containing protein | DUF1624 domain-containing protein, *Lechevalieria aerocolonigenes* (WP_035909001.1); 78/88 | - | - | - |
| *B0F77_RS29655* | 171 | hypothetical protein | hypothetical protein, *Lentzea terrae* (WP_112264025.1); 89/92 | - | - | - |
| *B0F77_RS29660* | 303 | helix-turn-helix domain-containing protein | XRE family transcriptional regulator, *Streptomyces iranensis* (WP_044572014.1); 90/93 | - | - | - |
| *B0F77_RS29665* | 267 | SDR family oxidoreductase | 3-oxoacyl-ACP reductase, *Streptomyces violaceusniger* (KUL62983.1); 94/97 | - | - | - |
| *B0F77_RS29670* | 599 | ABC transporter ATP-binding protein | ABC transporter ATP-binding protein, *Lechevalieria deserti* (WP_109636588.1); 94/96 | - | - | - |
| *B0F77_RS29675* | 462 | hypothetical protein | hypothetical protein, *Lentzea waywayandensis* (WP_093606216.1); 92/94 | - | - | - |
| *B0F77_RS29680* | 254 | alpha/beta hydrolase | alpha/beta fold hydrolase, *Lentzea waywayandensis* (WP_093592315.1); 79/91 | - | - | - |
| *B0F77_RS29685* | 244 | enoyl-CoA hydratase/isomerase family protein | enoyl-CoA hydratase/isomerase family protein, *Nocardia aobensis* (WP_051025421.1); 57/74 | - | - | - |
| *B0F77_RS29690* | 167 | flavin reductase family protein | flavin reductase, *Actinomadura* sp. WAC 06369 (WP_125618547.1); 66/73 | AbyZ | AbsH1 | AbmZ |
| *B0F77_RS29695* | 343 | 3-oxoacyl-ACP synthase III family protein | 3-oxoacyl-ACP synthase III family protein, *Streptomyces* sp. NRRL F-525 (WP_033287157.1); 66/81 | AbyA1 | AbsA1 | AbmA1 |
| *B0F77_RS29700* | 606 | HAD-IIIC family phosphatase | HAD-IIIC family phosphatase, *Kutzneria buriramensis* (WP_116181645.1); 58/71 | AbyA2 | AbsA2 | AbmA2 |
| *B0F77_RS29705* | 75 | acyl carrier protein | acyl carrier protein, *Frankia* sp. ACN1ag (KQC35070.1); 64/85 | AbyA3 | AbsA3 | AbmA3 |
| *B0F77_RS29710* | 231 | hypothetical protein | acyltransferase, *Streptomyces olindensis* (KDN76174.1); 65/78 | AbyA4 | AbsA4 | AbmA4 |
| *B0F77_RS29715* | 357 | alpha/beta hydrolase | alpha/beta hydrolase, *Actinomadura macra* (WP_067456402.1); 55/68 | AbyA5 | AbsA5 | AbmA5 |
| *B0F77_RS29720* | 117 | nuclear transport factor 2 family protein | hypothetical protein, *Planobispora rosea* (WP_084780980.1); 38/67 | - | - | - |
| *B0F77_RS29725* | 4307 | PKS I | type I polyketide synthase, *Streptomyces* sp. 2112.2 (WP_093485656.1); 52/64 | AbyB1 | AbsB1 | AbmB1 |
| //////////////////////////////////////////////////////////////////////////////// | | | | | | |
| *B0F77_RS29020* | 4693 | PKS I | SDR family NAD(P)-dependent oxidoreductase, *Streptomyces olivoreticuli* (WP_116210514.1); 54/65 | PKS I | PKS I | PKS I |
| *B0F77_RS29025* | - | PKS I | type I polyketide synthase, *Streptomyces odonnellii* (WP_046496891.1); 52/63 | PKS I | PKS I | PKS I |
| *B0F77_RS29030* | - | PKS I | modular polyketide synthase, *Streptomyces neyagawaensis* (BAW35659.1); 63/74 | PKS I | PKS I | PKS I |
| *B0F77_RS29035* | - | PKS I | type I modular polyketide synthase, *Streptomyces griseochromogenes* (ABV91286.1); 61/70 | PKS I | PKS I | PKS I |
| *B0F77_RS29040* | - | PKS I | hypothetical protein N566_17825, partial, *Streptomycetaceae bacterium* MP113-05 (EST35181.1); 66/74 | PKS I | PKS I | PKS I |
| *B0F77_RS29045* | - | PKS I | polyketide synthase, partial, *Streptomyces* sp. WM6391 (KKD10026.1); 65/78 | PKS I | PKS I | PKS I |

| | | | | | | |
|---|---|---|---|---|---|---|
| *B0F77_RS29050* | - | PKS I | polyketide synthase, partial, *Streptomyces platensis* (BAH67341.1); 67/75 | PKS I | PKS I | PKS I |
| *B0F77_RS29055* | - | PKS I | SDR family NAD(P)-dependent oxidoreductase, *Streptomyces* sp. NL15-2K (WP_124445724.1); 53/63 | PKS I | PKS I | PKS I |
| *B0F77_RS29060* | - | PKS I | Polyketide synthase dehydratase, partial, *Streptomyces* sp. MnatMP-M27 (SCG04427.1); 65/74 | PKS I | PKS I | PKS I |
| *B0F77_RS29065* | - | PKS I | type I polyketide synthase, *Streptacidiphilus neutrinimicus* (WP_052442691.1); 53/64 | PKS I | PKS I | PKS I |
| *B0F77_RS29070* | 500 | hypothetical protein | hypothetical protein, *Actinomadura* sp. H3C3 (WP_131898125.1); 57/67 | - | - | - |
| *B0F77_RS29075* | 183 | TetR/AcrR family transcriptional regulator | TetR/AcrR family transcriptional regulator, *Frankia symbiont of Coriaria ruscifolia* (WP_131786661.1); 76/87 | - | AbsC2 | - |
| *B0F77_RS29080* | 138 | hypothetical protein | nuclear transport factor 2 family protein, *Frankia symbiont of Coriaria ruscifolia* (WP_131786662.1); 85/90 | - | - | - |
| *B0F77_RS29085* | 479 | multidrug efflux MFS transporter | drug resistance transporter, EmrB/QacA subfamily, *Candidatus Frankia californiensis* (SBW22286.1); 76/87 | AbyD | AbsD | AbmD |
| *B0F77_RS29090* | 340 | LLM class flavin-dependent oxidoreductase | MsnO8 family LLM class oxidoreductase, *Actinocrispum wychmicini* (WP_132113998.1); 53/71 | AbyE | AbsE | AbmE1 |
| *B0F77_RS29095* | 398 | cytochrome P450 | cytochrome P450, *Streptomyces* sp. SCA2-2 (WP_129847673.1); 57/72 | AbyX/AbyV | AbsV/AbsX | AbmV |
| *B0F77_RS29100* | 256 | maleylpyruvate isomerase family protein | maleylpyruvate isomerase family mycothiol-dependent enzyme, Actinomadura *fibrosa* (WP_131760704.1); 40/56 | - | - | - |
| *B0F77_RS29105* | 282 | AfsR/SARP family transcriptional regulator | activator protein, *Actinomadura* sp. 6K520 (WP_131977283.1); 53/71 | AbyI | - | AbmI |
| *B0F77_RS29110* | 139 | Diels-Alderase | hypothetical protein, *Streptomyces* sp. FXJ7.023 (WP_037772721.1); 55/66 | AbyU | AbsU | AbmU |
| *B0F77_RS29115* | 268 | 3-hydroxybutyryl-CoA dehydrogenase | 3-hydroxybutyryl-CoA dehydrogenase, *Lechevalieria aerocolonigenes* (WP_030470217.1); 59/74 | - | - | - |
| *B0F77_RS29120* | 448 | ketoacyl-ACP synthase III | ketoacyl-ACP synthase III, *Saccharothrix* sp. NRRL B-16314 (WP_081915703.1); 72/81 | - | - | - |

**Table S44.** Predicted functions of ORFs in potential abyssomicin BGC from *Kutzneria buriramensis* DSM 45791 (NZ_QUNO01000013 and NZ_QUNO01000029).

| ORF | Size (aa) | Proposed function | Closest homolog, host (protein ID); Identity/Similarity (%) | Aby homolog | Abs homolog | Abm homolog |
|---|---|---|---|---|---|---|
| *BCF44_RS50005* | 291 | hypothetical protein | hypothetical protein, *Actinoplanes regularis* (WP_089291877.1); 74/82 | - | - | - |
| *BCF44_RS50000* | 193 | hypothetical protein | transposase, *Kutzneria* sp. 744 (EWM10014.1); 81/87 | - | - | - |
| *BCF44_RS49995* | 1027 | hypothetical protein | LuxR family transcriptional regulator, *Frankia* sp. Cc1.17 (WP_071084446.1); 41/53 | AbyH | - | AbmH |
| *BCF44_RS49990* | 253 | AfsR/SARP family transcriptional regulator | activator protein, *Rhodococcus yunnanensis* (WP_072806089.1); 65/78 | AbyI | - | AbmI |
| *BCF44_RS49985* | 625 | HAD-IIIC family phosphatase | HAD-IIIC family phosphatase, *Micromonospora* sp. RP3T (WP_107154962.1); 65/75 | AbyA2 | AbsA2 | AbmA2 |
| *BCF44_RS49980* | 75 | acyl carrier protein | acyl carrier protein, *Streptomyces* sp. NRRL F-525 (WP_033287159.1); 65/77 | AbyA3 | AbsA3 | AbmA3 |
| *BCF44_RS49975* | 251 | thioesterase | thioesterase, *Streptomyces* sp. 4R-3d (TFI25382.1); 55/65 | AbyT | AbsN | AbmT |
| *BCF44_RS49970* | 519 | ABC transporter ATP-binding protein | ABC transporter ATP-binding protein, *Frankia coriariae* (WP_047222767.1); 67/76 | AbyF4 | AbsF4 | AbmF4 |
| *BCF44_RS49965* | 186 | NADPH-dependent FMN reductase | NADPH-dependent FMN reductase, *Kutzneria albida* (WP_025355017.1); 61/74 | AbyZ | AbsH1 | AbmZ |
| *BCF44_RS49960* | 352 | LLM class flavin-dependent oxidoreductase | LLM class flavin-dependent oxidoreductase, *Saccharothrix syringae* (WP_033434377.1); 70/81 | - | - | AbmE2 |
| *BCF44_RS49955* | 65 | ferredoxin | ferredoxin, *Streptomyces yokosukanensis* (WP_067118731.1); 61/75 | - | AbsG2/AbsG1 | AbmG |
| *BCF44_RS49950* | 201 | TetR/AcrR family transcriptional regulator | regulatory protein TetR, *Frankia symbiont of Datisca glomerata* (AEH09834.1); 72/81 | - | - | - |
| *BCF44_RS49945* | 477 | MFS transporter | DHA2 family efflux MFS transporter permease subunit, *Frankia coriariae* (WP_047222768.1); 69/80 | AbyD | AbsD | AbmD |
| *BCF44_RS49940* | 452 | NtaA/DmoA family FMN-dependent monooxygenase | FMN-dependent oxidoreductase, nitrilotriacetate monooxygenase family, *Frankia symbiont of Datisca glomerata* (AEH09835.1); 71/81 | - | - | - |
| *BCF44_RS49935* | 395 | LLM class flavin-dependent oxidoreductase | LLM class flavin-dependent oxidoreductase, *Streptomyces* sp. RTd22 (WP_063731414.1); 57/66 | - | - | - |
| *BCF44_RS49930* | 406 | cytochrome P450 | cytochrome P450, *Saccharothrix syringae* (WP_033434378.1); 62/77 | AbyX/AbyV | AbsV/AbsX | AbmV |
| *BCF44_RS49925* | 336 | MsnO8 family LLM class oxidoreductase | LLM class flavin-dependent oxidoreductase, *Frankia alni* (WP_011601492.1); 55/65 | AbyE | AbsE | AbmE1 |
| *BCF44_RS49920* | 286 | ABC transporter permease | peptide ABC transporter permease, *Frankia coriariae* (KLL11667.1); 70/79 | AbyF3 | AbsF3 | AbmF3 |
| *BCF44_RS49915* | 308 | ABC transporter permease | ABC transporter permease, *Frankia coriariae* (KLL11700.1); 77/88 | AbyF2 | AbsF2 | AbmF2 |
| *BCF44_RS49910* | 550 | ABC transporter substrate-binding protein | ABC-type transporter, periplasmic subunit, *Frankia symbiont of Datisca glomerata* (AEH09839.1); 65/76 | AbyF1 | AbsF1 | AbmF1 |
| *BCF44_RS49905* | 345 | 3-oxoacyl-ACP synthase III family protein | 3-oxoacyl-ACP synthase III family protein, *Saccharothrix syringae* (WP_033434375.1); 79/85 | AbyA1 | AbsA1 | AbmA1 |
| *BCF44_RS49900* | 261 | acyltransferase | acyltransferas, *Frankia* sp. BMG5.30 (WP_076843553.1); 69/78 | AbyA4 | AbsA4 | AbmA4 |

| | | | | | | |
|---|---|---|---|---|---|---|
| *BCF44_RS49895* | 340 | alpha/beta hydrolase | alpha/beta hydrolase, *Frankia* sp. BMG5.30 (ONH34857.1); 65/75 | AbyA5 | AbsA5 | AbmA5 |
| *BCF44_RS49890* | 3837 | PKS I | SDR family NAD(P)-dependent oxidoreductase, *Streptomyces* sp. MK-45 (WP_126395712.1); 55/65 | PKS I | PKS I | PKS I |
| ////////////////////////////////////////////////////////////////////////////////////////////////////////////////////////////////////////////////////////////////////////////////////////////////////////////////////////////////////////////////////////////////////////////////////// | | | | | | |
| *BCF44_RS33785* | 105 | PKS I | polyketide synthase, *Actinoplanes* sp. N902-109 (AGL15968.1); 45/55 | PKS I | PKS I | PKS I |
| *BCF44_RS33790* | 3424 | PKS I | type I polyketide synthase, *Saccharothrix syringae* (WP_033434373.1); 59/69 | AbyB2 | AbsB2 | AbmB2 |
| *BCF44_RS33795* | 1332 | PKS I | type I polyketide synthase, *Saccharothrix syringae* (WP_051766715.1); 68/75 | AbyB3 | AbsB3 | AbmB3 |
| *BCF44_RS33800* | 125 | Diels-Alderase | hypothetical protein FrCorBMG51_12000, *Frankia coriariae* (KLL11361.1); 77/86 | AbyU | AbsU | AbmU |

**Table S45.** Predicted functions of ORFs surrounding AbyU homolog from *Streptomyces* sp. LHW50302 (NZ_QOIM01000040).

| ORF | Size (aa) | Proposed function | Closest homolog, host (protein ID); Identity/Similarity (%) | Aby homolog | Abs homolog | Abm homolog |
|---|---|---|---|---|---|---|
| *DQ392_RS21910* | 950 | PKS I | polyketide synthase, partial, *Candidatus Streptomyces philanthi* (RCG25677.1); 90/93 | - | - | - |
| *DQ392_RS21915* | 201 | Diels-Alderase | hypothetical protein, *Candidatus Streptomyces philanthi* (WP_114021297.1); 98/99 | AbyU | AbsU | AbmU |
| *DQ392_RS21920* | 403 | cytochrome P450 | cytochrome P450, *Candidatus Streptomyces philanthi* (WP_114021298.1); 99/99 | - | - | - |
| *DQ392_RS21925* | 253 | methyltransferase | methyltransferase, *Candidatus Streptomyces philanthi* (WP_114021299.1); 98/99 | - | - | - |
| *DQ392_RS21930* | 347 | 3-oxoacyl-ACP synthase | 3-oxoacyl-ACP synthase, *Candidatus Streptomyces philanthi* (WP_114021300.1); 99/99 | - | - | - |
| *DQ392_RS21935* | 1825 | PKS I | type I polyketide synthase, *Candidatus Streptomyces philanthi* (WP_114021301.1); 93/94 | - | - | - |
| *DQ392_RS21940* | 323 | dTDP-glucose 4,6-dehydratase | dTDP-glucose 4,6-dehydratase, *Candidatus Streptomyces philanthi* (WP_114021302.1); 98/99 | - | - | - |
| *DQ392_RS21945* | 972 | helix-turn-helix transcriptional regulator | helix-turn-helix transcriptional regulator, *Candidatus Streptomyces philanthi* (WP_114021303.1); 95/96 | - | - | - |
| *DQ392_RS21950* | 776 | DUF2075 domain-containing protein | DUF2075 domain-containing protein, *Candidatus Streptomyces philanthi* (WP_114021304.1); 97/98 | - | - | - |

**Table S46.** Predicted functions of ORFs in potential abyssomicin BGC from *Microbispora triticiradicis* NEAU-HRDPA2-9 (NZ_QFZU02000171.1).

| ORF | Size (aa) | Proposed function | Closest homolog, host (protein ID); Identity/Similarity (%) | Aby homolog | Abs homolog | Abm homolog |
|---|---|---|---|---|---|---|
| *DI270_RS29465* | 298 | ABC transporter ATP-binding protein | ABC transporter ATP-binding protein, *Streptosporangium subroseum* (WP_089206779.1); 76/81 | AbyF4 | AbsF4 | AbmF4 |
| *DI270_RS29470* | 395 | cytochrome P450 | cytochrome P450, *Streptomyces* sp. Amel2xE9 (WP_027758724.1); 86/92 | AbyX/AbyV | AbsV/AbsX | AbmV |
| *DI270_RS29475* | 68 | ferredoxin | ferredoxin-1, *Streptomyces* sp. NRRL F-6491 (KOX15570.1); 66/78 | - | AbsG1/AbsG2 | AbmG |
| *DI270_RS29480* | 385 | acyltransferase | acyltransferase, *Streptomyces* sp. E14 (WP_009191675.1); 71/78 | - | AbsI | - |
| *DI270_RS29485* | 401 | cytochrome P450 | cytochrome P450, *Streptosporangium subroseum* (WP_089206781.1); 85/91 | AbyX/AbyV | AbsV/AbsX | AbmV |
| *DI270_RS29490* | 77 | ferredoxin | ferredoxin, *Streptosporangium subroseum* (WP_089206642.1); 87/91 | - | AbsG2/AbsG1 | AbmG |
| *DI270_RS29495* | 332 | aldo/keto reductase | aldo/keto reductase, *Streptosporangium subroseum* (WP_089206643.1); 82/89 | - | AbsJ | AbmJ |
| *DI270_RS29500* | 134 | Diels-Alderase | hypothetical protein SAMN05216276_1006109, *Streptosporangium subroseum* (SNS24121.1); 91/96 | AbyU | AbsU | AbmU |
| *DI270_RS29505* | 933 | LuxR family transcriptional regulator | LuxR family transcriptional regulator, *Frankia* sp. QA3 (WP_009738951.1); 54/66 | AbyH | - | AbmH |
| *DI270_RS29510* | 195 | TetR/AcrR family transcriptional regulator | TetR/AcrR family transcriptional regulator, *Microbispora rosea* (WP_076442332.1); 91/94 | - | AbsC2 | - |
| *DI270_RS29515* | 475 | MFS transporter | MFS transporter, *Microbispora* sp. GKU 823 (WP_079317079.1); 88/92 | AbyD | AbsD | AbmD |
| *DI270_RS29520* | 252 | AfsR/SARP family transcriptional regulator | AfsR/SARP family transcriptional regulator, *Microbispora* sp. GKU 823 (WP_079317077.1); 88/94 | AbyR/AbyI | - | AbmI |

**Table S47.** Predicted functions of ORFs in abyssomicin BGC from *Micromonospora wenchangensis* CCTCC AA 2012002 (NZ_MZMV01000061.1 and NZ_MZMV01000027.1).

| ORF | Size (aa) | Proposed function | Closest homolog, host (protein ID); Identity/Similarity (%) | Aby homolog | Abs homolog | Abm homolog |
|---|---|---|---|---|---|---|
| *B5D80_RS26560* | 251 | SARP family transcriptional regulator | DNA-binding SARP family transcriptional activator, *Actinokineospora auranticolor* (PPK71426.1); 74/82 | AbyR/AbyI | - | AbmI |
| *B5D80_RS26565* | 397 | cytochrome P450 | cytochrome P450, *Verrucosispora* (WP_013733063.1); 82/90 | AbyX/AbyV | AbsV/AbsX | AbmV |
| *B5D80_RS26570* | 898 | LuxR family transcriptional regulator | LuxR family transcriptional regulator, *Verrucosispora maris* AB-18-032 (AEK75494.1); 68/77 | AbyH | - | AbmH |
| *B5D80_RS26575* | 141 | Diels-Alderase | Chain A, Abyu – Wildtype, *Verrucosispora maris* AB-16-032 (5DYV_A); 85/91 | AbyU | AbsU | AbmU |
| *B5D80_RS26580* | 617 | RHS repeat protein | YD repeat protein, *Verrucosispora maris* AB-18-032 (AEK75496.1); 75/82 | AbyK | - | - |
| *B5D80_RS26585* | 1040 | HAD-IIIC family phosphatase | methoxymalonyl-ACP biosynthesis protein FkbH, *Micromonospora wenchangensis* (OWV01453.1); 99/100 | AbyA2 | AbsA2 | AbmA2 |
| *B5D80_RS26590* | 75 | acyl carrier protein | acyl carrier protein, *Verrucosispora* (WP_043723886.1); 75/82 | AbyA3 | AbsA3 | AbmA3 |
| *B5D80_RS26595* | 250 | acyltransferase | Acyltransferase, *Verrucosispora maris* (WP_013733055.1); 83/91 | AbyA4 | AbsA4 | AbmA4 |
| *B5D80_RS26600* | 355 | alpha/beta hydrolase | alpha/beta hydrolase, *Verrucosispora maris* (WP_013733054.1); 79/87 | AbyA5 | AbsA5 | AbmA5 |
| //////////////////////////////////////////////////////////////////////////////// | | | | | | |
| *B5D80_RS17105* | 2437 | PKS I | modular polyketide synthase, *Verrucosispora maris* AB-18-032 (AEB44393.1); 71/77 | AbyB1 | AbsB1 | AbmB1 |
| *B5D80_RS17110* | 2053 | PKS I | type I polyketide synthase, *Verrucosispora maris* (WP_013733052.1); 73/80 | AbyB2 | AbsB2 | AbmB2 |
| *B5D80_RS17115* | 998 | PKS I | acyltransferase domain-containing protein, *Verrucosispora maris* (WP_013733051.1); 77/84 | AbyB3 | AbsB3 | AbmB3 |
| *B5D80_RS17120* | 230 | TetR/AcrR family transcriptional regulator | TetR/AcrR family transcriptional regulator, *Verrucosispora maris* (WP_013733050.1); 88/92 | AbyC | - | AbmC |
| *B5D80_RS17125* | 474 | DHA2 family efflux MFS transporter permease subunit | DHA2 family efflux MFS transporter permease subunit, *Verrucosispora* (WP_013733049.1); 86/92 | AbyD | AbsD | AbmD |
| *B5D80_RS17130* | 336 | LLM class flavin-dependent oxidoreductase | LLM class flavin-dependent oxidoreductase, *Verrucosispora* (WP_013733048.1): 82/89 | AbyE | AbsE | AbmE1 |
| *B5D80_RS17135* | 566 | ABC transporter substrate-binding protein | ABC transporter substrate-binding protein, *Verrucosispora* (WP_013733047.1); 75/86 | AbyF1 | AbsF1 | AbmF1 |
| *B5D80_RS17140* | 311 | ABC transporter permease | ABC transporter permease, *Verrucosispora* (WP_013733046.1); 82/88 | AbyF2 | AbsF2 | AbmF2 |
| *B5D80_RS17145* | 283 | ABC transporter permease | ABC transporter permease, *Verrucosispora* sp. FIM060022 (WP_126713145.1); 83/87 | AbyF3 | AbsF3 | AbmF3 |
| *B5D80_RS17150* | 539 | ABC transporter ATP-binding protein | ABC transporter ATP-binding protein, *Verrucosispora maris* (WP_013733044.1); 80/87 | AbyF4 | AbsF4 | AbmF4 |
| *B5D80_RS17155* | 396 | cytochrome P450 | cytochrome P450, *Verrucosispora maris* (WP_013733043.1); 83/86 | AbyV/AbyX | AbsV/AbsX | AbmV |
| *B5D80_RS17160* | 79 | ferredoxin-1 | Ferredoxin, *Verrucosispora* sp. FIM060022 (WP_126713142.1); 68/77 | AbyW | AbsG2/AbsG1 | AbmG |

| | | | | | | |
|---|---|---|---|---|---|---|
| *B5D80_RS17165* | 203 | FMN reductase (NADPH) | FMN reductase (NADPH), *Streptomyces* sp. A244 (WP_107460035.1); 81/86 | AbyZ | AbsH1 | AbmZ |
| *B5D80_RS17170* | 303 | thioesterase | Thioesterase, *Verrucosispora* (P_081476081.1); 70/76 | AbyT | AbsN | AbmT |

**Table S48.** Predicted functions of ORFs in potential abyssomicin BGC from *Micromonospora* sp. RP3T (PYPS01000018 and PYPS01000002.1).

| ORF | Size (aa) | Proposed function | Closest homolog, host (protein ID); Identity/Similarity (%) | Aby homolog | Abs homolog | Abm homolog |
|---|---|---|---|---|---|---|
| *C8054_RS16510* | 134 | Diels-Alderase | hypothetical protein, *Streptomyces rimosus* (WP_033030402.1); 37/58 | AbyU | AbsU | AbmU |
| *C8054_RS16515* | 947 | hypothetical protein | hypothetical protein, *Sphaerisporangium* sp. LHW63015 (WP_113983256.1); 50/59 | - | - | - |
| *C8054_RS16520* | 413 | cytochrome P450 | cytochrome P450, *Streptomyces* sp. ICBB 8177 (WP_109446503.1); 66/79 | AbyX/AbyV | AbsV/AbsX | AbmV |
| *C8054_RS16525* | 68 | ferredoxin | ferredoxin, *Nocardia* sp. BMG111209 (WP_026343320.1); 52/64 | - | AbsG1 | AbmG |
| *C8054_RS16530* | 940 | LuxR family transcriptional regulator | regulatory LuxR family protein, *Herbihabitans rhizosphaerae* (RZS36569.1); 39/55 | - | - | - |
| *C8054_RS16535* | 494 | MFS transporter | DHA2 family efflux MFS transporter permease subunit, *Frankia* sp. Cc1.17 (WP_071090374.1); 49/65 | AbyD | AbsD | AbmD |
| *C8054_RS16540* | 316 | TIGR03564 family F420-dependent LLM class oxidoreductase | TIGR03564 family F420-dependent LLM class oxidoreductase, *Amycolatopsis tolypomycina* (WP_091310088.1); 65/78 | - | - | - |
| *C8054_RS16545* | 358 | alpha/beta hydrolase | alpha/beta hydrolase, *Actinocrispum wychmicini* (WP_132114014.1); 59/72 | AbyA5 | AbsA5 | AbmA5 |
| //////////////////////////////////////////////////////////////////////////////////////////////////////////////////////////////////////////////////////////////////////////////////////////////////////// | | | | | | |
| *C8054_01980* | - | PKS I | acyl transferase domain-containing protein, *Streptomyces* sp. 70 (PIG77230.1); 60/70 | PKS I | PKS I | PKS I |
| *C8054_01985* | 344 | 3-oxoacyl-ACP synthase | 3-oxoacyl-ACP synthase III family protein, *Amycolatopsis* sp. CA-126428 (WP_103342534.1); 59/74 | AbyA1 | AbsA1 | AbmA1 |
| *C8054_01990* | 269 | acyltransferase | acyltransferase, *Actinomadura pelletieri* (WP_121438112.1); 62/71 | AbyA4 | AbsA4 | AbmA4 |
| *C8054_01995* | 487 | hypothetical protein | hypothetical protein, *Micromonospora wenchangensis* (WP_088642273.1); 50/64 | - | - | - |
| *C8054_02000* | 488 | hypothetical protein | drug resistance transporter, EmrB/QacA subfamily, partial, *Streptomyces* sp. SolWspMP-5a-2 (SCD36213.1); 43/68 | AbyD | AbsD | AbmD |
| *C8054_02005* | 184 | TetR/AcrR family transcriptional regulator | TetR/AcrR family transcriptional regulator, *Frankia* sp. QA3 (WP_009742630.1); 51/65 | - | - | - |
| *C8054_02010* | 140 | hypothetical protein | hypothetical protein, *Micromonospora auratinigra* (WP_091660051.1); 67/83 | - | - | - |
| *C8054_02015* | 343 | methyltransferase | methyltransferase, *Streptomyces griseorubiginosus* (WP_123763216.1); 42/59 | - | - | - |
| *C8054_02020* | 255 | thioesterase | thioesterase, *Actinomadura pelletieri* (WP_121438108.1); 58/70 | AbyT | AbsN | AbmT |
| *C8054_02025* | 258 | activator protein | activator protein, *Micromonospora endolithica* (WP_120723474.1); 61/74 | AbyI/AbyR | - | AbmI |
| *C8054_02030* | 630 | methoxymalonyl-ACP biosynthesis protein FkbH | HAD-IIIC family phosphatase, *Actinomadura pelletieri* (WP_121438114.1); 65/76 | AbyA2 | AbsA2 | AbmA2 |
| *C8054_02035* | 80 | acyl carrier protein | acyl carrier protein, *Actinomadura* sp. 6K520 (WP_131977265.1); 55/76 | AbyA3 | AbsA3 | AbmA3 |
| *C8054_02040* | 395 | methionine adenosyltransferase | methionine adenosyltransferase, *Actinoplanes* sp. N902-109 (WP_015623988.1); 89/95 | - | - | - |
| *C8054_02045* | 385 | FAD-dependent oxidoreductase | ferredoxin reductase, *Actinoplanes* sp. TFC3 (WP_067499520.1); 75/82 | - | - | - |

| | | | | | | |
|---|---|---|---|---|---|---|
| *C8054_02050* | 407 | cytochrome P450 | cytochrome P450, *Amycolatopsis kentuckyensis* (WP_086841038.1); 61/73 | AbyX/AbyV | AbsX/AbsV | AbmV |

**Table S49.** Predicted functions of ORFs surrounding AbyU homolog from *Streptomyces monomycini* NRRL B-24309 (NZ_KL571104.1).

| ORF | Size (aa) | Proposed function | Closest homolog, host (protein ID); Identity/Similarity (%) | Aby homolog | Abs homolog | Abm homolog |
|---|---|---|---|---|---|---|
| *HY87_RS0129240* | 889 | helix-turn-helix transcriptional regulator | helix-turn-helix transcriptional regulator, *Streptomyces lavendulae* (WP_030241329.1); 53/61 | - | - | - |
| *HY87_RS0129245* | 1131 | hypothetical protein | helix-turn-helix transcriptional regulator, *Streptomyces lavendulae* (WP_030241327.1); 57/68 | - | - | - |
| *HY87_RS0129250* | 245 | AfsR/SARP family transcriptional regulator | activator protein, *Kutzneria buriramensis* (WP_116181646.1); 50/66 | - | - | - |
| *HY87_RS1000000143920* | 169 | Diels-Alderase | hypothetical protein, *Streptomyces caatingaensis* (WP_049718340.1); 44/65 | AbyU | AbsU | AbmU |

**Table S50.** Predicted functions of ORFs in potential BGC from *Streptomyces* sp. MUSC 14 (NZ_MLYN01000052).

| ORF | Size (aa) | Proposed function | Closest homolog, host (protein ID); Identity/Similarity (%) | Aby homolog | Abs homolog | Abm homolog |
|---|---|---|---|---|---|---|
| *BIV25_RS31795* | 320 | LLM class flavin-dependent oxidoreductase | LLM class flavin-dependent oxidoreductase, *Frankia* sp. QA3 (WP_009742601.1); 63/76 | - | - | - |
| *BIV25_RS31800* | 185 | DinB family protein | uncharacterized protein DUF664, *Streptomyces* sp. 67 (RED71639.1); 86/90 | - | - | - |
| *BIV25_RS47940* | 41 | GNAT family N-acetyltransferase | GNAT family N-acetyltransferase, *Streptomyces* sp. BK161 (WP_133927666.1); 78/87 | - | - | - |
| *BIV25_RS31805* | 508 | cytochrome P450 | cytochrome P450, *Streptomyces* sp. MUSC 1 (WP_079173711.1); 92/94 | - | - | - |
| *BIV25_RS31810* | 406 | DUF2029 domain-containing protein | DUF2029 domain-containing protein, *Streptomyces* sp. MUSC 1 (WP_107471405.1); 95/96 | - | - | - |
| *BIV25_RS31815* | 473 | 3-deoxy-7-phosphoheptulonate synthase class II | 3-deoxy-7-phosphoheptulonate synthase class II, *Streptomyces tateyamensis* (WP_110665468.1); 75/84 | - | - | - |
| *BIV25_RS31820* | 285 | 3-hydroxybutyryl-CoA dehydrogenase | 3-hydroxybutyryl-CoA dehydrogenase, *Frankia* sp. BMG5.36 (OHV43725.1); 58/77 | - | - | - |
| *BIV25_RS31825* | 344 | ketoacyl-ACP synthase III | beta-ketoacyl-ACP synthase III, *Amycolatopsis* sp. 8-3EHSu (WP_130478882.1); 70/81 | - | - | - |
| *BIV25_RS31830* | 450 | crotonyl-CoA carboxylase/reductase | crotonyl-CoA carboxylase/reductase, *Actinomadura macra* (WP_067467652.1); 77/86 | - | - | - |
| *BIV25_RS31835* | 255 | thioesterase | thioesterase, *Streptomyces* sp. MA5143a (WP_107466330.1); 62/73 | - | - | - |
| *BIV25_RS31840* | 173 | pyridoxamine 5'-phosphate oxidase family protein | pyridoxamine 5'-phosphate oxidase family protein, *Streptomyces iranensis* (WP_044580008.1); 77/84 | - | - | - |
| *BIV25_RS31845* | 3932 | PKS I | type I polyketide synthase, *Streptomyces iranensis* (WP_044580009.1); 80/87 | - | - | - |
| *BIV25_RS31850* | 1360 | PKS I | type I polyketide synthase, *Streptomyces iranensis* (WP_044580010.1); 82/88 | - | - | - |
| *BIV25_RS31855* | 411 | cytochrome P450 | cytochrome P450, *Saccharothrix syringae* (WP_033431229.1); 66/78 | - | - | - |
| *BIV25_RS31860* | 119 | nuclear transport factor 2 family protein | nuclear transport factor 2 family protein, *Streptomyces cattleya* (WP_014627233.1); 56/76 | - | - | - |
| *BIV25_RS31865* | 139 | Diels-Alderase | hypothetical protein, *Streptomyces* sp. E5N91 SAI-083 (WP_123627591.1); 61/78 | AbyU | AbsU | AbmU |
| *BIV25_RS31870* | 412 | monooxygenase | FAD-dependent monooxygenase, *Saccharopolyspora* sp. 16K309 (WP_132674765.1); 55/65 | - | - | - |
| *BIV25_RS31875* | 445 | salicylate synthase | salicylate synthetase, *Amycolatopsis xylanica* (SDW43103.1); 68/79 | - | - | - |
| *BIV25_RS31880* | 317 | hypothetical protein | malonyl transferase, *Streptomyces uncialis* (WP_073785742.1); 39/57 | - | - | - |
| *BIV25_RS31885* | 355 | arylcarboxylate reductase | arylcarboxylate reductase, *Streptomyces* sp. PRh5 (EXU64039.1); 62/73 | - | - | - |
| *BIV25_RS31890* | 385 | cytochrome P450 | cytochrome P450, *Actinobacteria bacterium* (PZM89993.1); 43/56 | - | - | - |
| *BIV25_RS31895* | 203 | flavin reductase family protein | flavin reductase family protein, *Rhodoplanes* sp. Z2-YC6860 (WP_068017711.1); 51/66 | - | - | - |
| *BIV25_RS31900* | 377 | helix-turn-helix domain-containing protein | helix-turn-helix domain-containing protein, *Streptomyces phaeochromogenes (WP_079053449.1); 68/74* | - | - | - |

| | | | | | | |
|---|---|---|---|---|---|---|
| *BIV25_RS31905* | 918 | helix-turn-helix transcriptional regulator | helix-turn-helix transcriptional regulator, *Streptomyces iranensis* (WP_044580016.1); 64/75 | - | - | - |
| *BIV25_RS31910* | 198 | TetR/AcrR family transcriptional regulator | TetR/AcrR family transcriptional regulator, *Pseudonocardia acaciae* (WP_051579483.1); 55/69 | - | - | - |
| *BIV25_RS31915* | 564 | MFS transporter | DHA2 family efflux MFS transporter permease subunit, *Kutzneria buriramensis* (WP_116174721.1); 54/73 | - | - | - |

**Table S51.** Predicted functions of ORFs in potential abyssomicin BGC from *Streptomyces niveus* NRRL 2466 (NZ_MDCR01000040).

| **ORF** | **Size (aa)** | **Proposed function** | **Closest homolog, host (protein ID); Identity/Similarity (%)** | **Aby homolog** | **Abs homolog** | **Abm homolog** |
|---|---|---|---|---|---|---|
| *BHU15_RS06405* | 388 | acyltransferase | acyltransferase, *Streptomyces luteocolor* (WP_069885825.1); 49/65 | - | AbsI | - |
| *BHU15_RS06410* | 923 | helix-turn-helix transcriptional regulator | helix-turn-helix transcriptional regulator, *Streptomyces* sp. 4R-3d (TFI22095.1); 99/99 | AbyH | - | AbmH |
| *BHU15_RS06415* | 253 | AfsR/SARP family transcriptional regulator | activator protein, *Streptomyces* sp. SCA2-2 (WP_129847683.1); 63/74 | AbyI | - | AbmI |
| *BHU15_RS06420* | 195 | TetR/AcrR family transcriptional regulator | TetR/AcrR family transcriptional regulator, *Sinosporangium album* (WP_093167074.1); 74/84 | - | AbsC2 | - |
| *BHU15_RS06425* | 499 | MFS transporter | MFS transporter, *Sinosporangium album* (WP_093167076.1); 81/88 | AbyD | AbsD | AbyD |
| *BHU15_RS06430* | 321 | hypothetical protein | ABC transporter substrate-binding protein, *Streptomyces* sp. 4R-3d (TFI25403.1); 99/99 | AbyF1 | AbsF1 | AbmF1 |
| *BHU15_RS06435* | 282 | ABC transporter permease | ABC transporter permease, *Streptomyces* sp. SCA2-2 (WP_129847675.1); 69/82 | AbyF2 | AbsF2 | AbsmF2 |
| *BHU15_RS06440* | 410 | amidohydrolase family protein | amidohydrolase family protein, *Streptomyces* sp. SCA2-2 (WP_129847676.1); 62/71 | - | - | AbmM |
| *BHU15_RS06445* | 304 | ABC transporter permease | ABC transporter permease, *Streptomyces* sp. 4R-3d (TFI25375.1); 98/98 | AbyF3 | AbsF3 | AbmF3 |
| *BHU15_RS06450* | 620 | ABC transporter ATP-binding protein | ABC transporter ATP-binding protein, *Streptomyces* sp. 4R-3d (TFI25374.1); 99/99 | AbyF4 | AbsF4 | AbmF4 |
| *BHU15_RS06455* | 217 | Diels-Alderase | AbmU, *Streptomyces koyangensis* (AVI57412.1); 53/68 | AbyU | AbsU | AbmU |

**Table S52.** Predicted functions of ORFs in potential abyssomicin BGC from *Streptomyces* sp. NL15-2K (NZ_BHXA01000189.1).

| ORF | Size (aa) | Proposed function | Closest homolog, host (protein ID); Identity/Similarity (%) | Aby homolog | Abs homolog | Abm homolog |
|---|---|---|---|---|---|---|
| *SNL152K_RS52860* | 1358 | PKS I | SDR family NAD(P)-dependent oxidoreductase, *Streptomyces griseocarneus* (WP_121797418.1); 50/61 | PKS I | PKS I | PKS I |
| *SNL152K_RS52865* | 2156 | PKS I | type I polyketide synthase, *Streptomyces formicae* (WP_098241246.1); 54/64 | PKS I | PKS I | PKS I |
| *SNL152K_RS52870* | 1727 | PKS I | modular polyketide synthase, *Streptomyces* sp. RK95-74 (BAW35608.1); 49/61 | PKS I | PKS I | PKS I |
| *SNL152K_RS52875* | 409 | cytochrome P450 | cytochrome P450, *Streptomyces* sp. WMMB 322 (WP_055484149.1); 55/68 | AbyX/AbyV | AbsV/AbsX | AbmV |
| *SNL152K_RS52880* | 63 | ferredoxin | ferredoxin, *Thermostaphylospora chromogena* (WP_093260901.1); 61/75 | - | AbsG1/AbsG2 | AbmG |
| *SNL152K_RS52885* | 145 | Diels-Alderase | hypothetical protein, *Streptomyces geranii* (WP_105971044.1); 58/72 | AbyU | AbsU | AbmU |
| *SNL152K_RS52890* | 267 | acyltransferase | acyltransferase, *Streptomyces olindensis* (KDN76174.1); 60/72 | AbyA4 | AbsA4 | AbmA4 |
| *SNL152K_RS52895* | 390 | alpha/beta fold hydrolase | alpha/beta hydrolase, *Streptomyces geranii* (WP_107503113.1); 75/80 | AbyA5 | AbsA5 | AbmA5 |
| *SNL152K_RS52900* | 354 | hypothetical protein | hypothetical protein, *Streptomyces geranii* (WP_105971045.1); 92/95 | - | - | - |
| *SNL152K_RS52905* | 151 | Diels-Alderase | hypothetical protein, *Streptomyces geranii* (WP_105971044.1); 91/95 | AbyU | AbsU | AbmU |
| *SNL152K_RS52910* | 72 | hypothetical protein | hypothetical protein, *Streptomyces geranii* (WP_105971043.1); 80/87 | - | - | - |
| *SNL152K_RS52915* | 69 | hypothetical protein | hypothetical protein, *Streptomyces geranii* (WP_105971042.1); 93/98 | - | - | - |
| *SNL152K_RS52920* | 520 | acyl-CoA carboxylase subunit beta | acyl-CoA carboxylase subunit beta, *Streptomyces* sp. SYSU K10008 (WP_128380899.1); 81/89 | - | - | - |

**Table S53.** Predicted functions of ORFs in potential abyssomicin BGC from *Streptomyces* sp. NRRL F-525 (NZ_JNXE01000068).

| ORF | Size (aa) | Proposed function | Closest homolog, host (protein ID); Identity/Similarity (%) | Aby homolog | Abs homolog | Abm homolog |
|---|---|---|---|---|---|---|
| *OO69_RS46840* | 174 | Diels-Alderase | hypothetical protein, *Streptomyces caatingaensis* (WP_049718340.1); 34/47 | AbyU | AbsU | AbmU |
| *OO69_RS46845* | 343 | 3-oxoacyl-ACP synthase III family protein | 3-oxoacyl-ACP synthase III family protein, *Lentzea kentuckyensis* (WP_086666305.1); 66/81 | AbyA1 | AbsA1 | AbmA1 |
| *OO69_RS46850* | 634 | HAD-IIIC family phosphatase | HAD-IIIC family phosphatase, *Actinocrispum wychmicini* (WP_132116038.1); 62/73 | AbyA2 | AbsA2 | AbmA2 |
| *OO69_RS46855* | 76 | acyl carrier protein | acyl carrier protein, *Streptomyces* sp. NRRL F-5123 (WP_031525362.1); 72/89 | AbyA3 | AbsA3 | AbmA3 |
| *OO69_RS46860* | 228 | acyltransferase | acyltransferase, *Streptomyces kanamyceticus* (WP_055549000.1); 61/77 | AbyA4 | AbsA4 | AbmA4 |
| *OO69_RS46865* | 364 | alpha/beta hydrolase | alpha/beta hydrolase, *Actinocrispum wychmicini* (WP_132114014.1); 62/74 | AbyA5 | AbsA5 | AbmA5 |
| *OO69_RS46870* | 268 | thioesterase | thioesterase, *Actinoplanes* sp. N902-109 (WP_051167423.1); 54/62 | AbyT | AbsN | AbmT |
| *OO69_RS46875* | 1637 | PKS I | type I polyketide synthase, *Streptomyces* sp. MBT76 (WP_058042044.1); 62/71 | PKS I | PKS I | PKS I |

**Table S54.** Predicted functions of ORFs in potential abyssomicin BGC from *Streptomyces* sp. NRRL F-525 (NZ_JNXE01000075).

| ORF | Size (aa) | Proposed function | Closest homolog, host (protein ID); Identity/Similarity (%) | Aby homolog | Abs homolog | Abm homolog |
|---|---|---|---|---|---|---|
| *OO69_RS47325* | 497 | hypothetical protein | hypothetical protein, *Actinomadura macra* (WP_067456441.1); 51/62 | - | - | - |
| *OO69_RS47330* | 251 | thioesterase | thioesterase, *Streptomyces* sp. NEAU-S7GS2 (AWN25442.1); 65/79 | AbyT | AbsN | AbmT |
| *OO69_RS47335* | 255 | AfsR/SARP family transcriptional regulator | activator protein, *Actinomadura chibensis* (WP_067904593.1); 60/73 | AbyI | - | AbmI |
| *OO69_RS47345* | 1011 | LuxR family transcriptional regulator | regulatory LuxR family protein, *Herbihabitans rhizosphaerae* (RZS36569.1); 42/57 | - | - | - |
| *OO69_RS47350* | 184 | Diels-Alderase | hypothetical protein, *Streptomyces* sp. NL15-2K (WP_124445685.1); 40/57 | AbyU | AbsU | AbmU |
| *OO69_RS47355* | 518 | MFS transporter | DHA2 family efflux MFS transporter permease subunit, *Streptomyces* sp. FXJ1.172 (WP_107304187.1); 57/75 | AbyD | AbsD | AbyD |

**Table S55.** Predicted functions of ORFs in potential abyssomicin BGC from *Streptomyces* sp. NRRL F-5126 (NZ_JOFZ01000007).

| ORF | Size (aa) | Proposed function | Closest homolog, host (protein ID); Identity/Similarity (%) | Aby homolog | Abs homolog | Abm homolog |
|---|---|---|---|---|---|---|
| *IH48_RS33370* | 555 | PKS I | type I polyketide synthase PikAI, *Streptomyces* sp. SolWspMP-5a-2 (SCD36199.1); 87/92 | PKS I | PKS I | PKS I |
| *IH48_RS0116245* | 537 | MFS transporter | drug resistance transporter, EmrB/QacA subfamily, partial, *Streptomyces* sp. SolWspMP-5a-2 (SCD36213.1); 92/94 | AbyD | AbsD | AbyD |
| *IH48_RS0116250* | 510 | MFS transporter | DHA2 family efflux MFS transporter permease subunit, *Streptomyces* sp. SolWspMP-5a-2 (WP_093830605.1); 91/95 | AbyD | AbsD | AbyD |
| *IH48_RS0116255* | 214 | TetR/AcrR family transcriptional regulator | TetR/AcrR family transcriptional regulator, *Streptomyces* sp. SolWspMP-5a-2 (WP_093830608.1); 90/95 | - | - | - |
| *IH48_RS0116260* | 494 | hypothetical protein | hypothetical protein, *Streptomyces* sp. SolWspMP-5a-2 (WP_093830610.1); 90/93 | - | - | - |
| *IH48_RS0116265* | 133 | Diels-Alderase | hypothetical protein GA0115242_12174, *Streptomyces* sp. SolWspMP-5a-2 (SCE08777.1); 90/94 | AbyU | AbsU | AbmU |
| *IH48_RS0116270* | 76 | hypothetical protein | hypothetical protein, *Streptomyces* sp. SolWspMP-5a-2 (WP_093830614.1); 76/81 | - | - | - |
| *IH48_RS0116275* | 343 | 3-oxoacyl-ACP synthase III family protein | 3-oxoacyl-ACP synthase III family protein, *Streptomyces* sp. SolWspMP-5a-2 (WP_093830729.1); 97/99 | AbyA1 | AbsA1 | AbmA1 |
| *IH48_RS0116280* | 265 | AfsR/SARP family transcriptional regulator | activator protein, *Nonomuraea polychroma* (WP_127931360.1); 54/69 | AbyI | - | AbmI |
| *IH48_RS0116285* | 373 | alpha/beta hydrolase | alpha/beta hydrolase, *Streptomyces* sp. SolWspMP-5a-2 (WP_093830618.1); 90/94 | AbyA5 | AbsA5 | AbmA5 |
| *IH48_RS0116290* | 1519 | PKS I | type I polyketide synthase, *Streptomyces* sp. SolWspMP-5a-2 (WP_093830620.1); 91/94 | - | - | - |
| *IH48_RS0116295* | 65 | ferredoxin | ferredoxin, *Streptomyces* sp. SolWspMP-5a-2 (WP_093830624.1); 88/95 | - | AbsG1 | AbmG |
| *IH48_RS0116300* | 410 | cytochrome P450 | cytochrome P450, *Streptomyces* sp. SolWspMP-5a-2 (WP_093830626.1); 92/95 | AbyX/AbyV | AbsV/AbsX | AbmV |
| *IH48_RS0116305* | 273 | thioesterase | thioesterase, *Streptomyces* sp. SolWspMP-5a-2 (WP_093830628.1); 90/93 | AbyT | AbsN | AbmT |

**Table S56.** Predicted functions of ORFs surrounding AbyU homolog from *Streptomyces* sp. NRRL F-5755 (NZ_LGCW01000306).

| ORF | Size (aa) | Proposed function | Closest homolog, host (protein ID); Identity/Similarity (%) | Aby homolog | Abs homolog | Abm homolog |
|---|---|---|---|---|---|---|
| *ADK86_RS35070* | 435 | hypothetical protein | hypothetical protein, *Streptomyces albus* (WP_060732989.1); 99/99 | - | - | - |
| *ADK86_RS35075* | 433 | MFS transporter | MFS transporter, *Streptomyces albus* (WP_060732990.1); 99/99 | - | - | - |
| *ADK86_RS35080* | 174 | Diels-Alderase | hypothetical protein ADL21_37760, *Streptomyces albus subsp. albus* (KWT56726.1); 98/98 | AbyU | AbsU | AbmU |
| *ADK86_RS35085* | 448 | MFS transporter | MFS transporter, *Streptomyces albus* (WP_060732991.1); 99/99 | - | - | - |
| *ADK86_RS35090* | 173 | CGNR zinc finger domain-containing protein | CGNR zinc finger domain-containing protein, *Streptomyces* sp. WAC 06725 (RSO35445.1); 99/100 | - | - | - |
| *ADK86_RS35095* | 981 | type I polyketide synthase | type I polyketide synthase, *Streptomyces albus* (WP_060732992.1); 96/97 | - | - | - |
| *ADK86_RS35100* | 402 | FAD-dependent oxidoreductase | FAD-dependent oxidoreductase, *Streptomyces rimosus* (WP_030643530.1); 99/99 | - | - | - |
| *ADK86_RS35105* | 255 | thioesterase | thioesteras, *Streptomyces albus* (WP_060732994.1); 98/99 | - | - | - |
| *ADK86_RS35110* | 572 | (2,3-dihydroxybenzoyl)adenylate synthase | 2,3-dihydroxybenzoate--AMP ligase, *Streptomyces griseoflavus* (KOG53532.1); 97/97 | - | - | - |
| *ADK86_RS35115* | 77 | acyl carrier protein | acyl carrier protein, *Streptomyces rimosus* (WP_030643536.1); 99/100 | - | - | - |
| *ADK86_RS35120* | 358 | hypothetical protein | hypothetical protein, *Streptomyces albus* (WP_060732996.1); 98/98 | - | - | - |

**Table S57.** Predicted functions of ORFs surrounding AbyU homolog from *Streptomyces* sp. NRRL S-31 (NZ_JOCB01000102.1).

| ORF | Size (aa) | Proposed function | Closest homolog, host (protein ID); Identity/Similarity (%) | Aby homolog | Abs homolog | Abm homolog |
|---|---|---|---|---|---|---|
| *IF37_RS0131380* | 403 | cytochrome P450 | cytochrome P450, *Frankia* sp. Cc1.17 (WP_071083429.1); 73/81 | - | - | - |
| *IF37_RS0131385* | 181 | Diels-Alderase | hypothetical protein, *Frankia* sp. Cc1.17 (WP_131803042.1); 62/72 | AbyU | AbsU | AbmU |
| *IF37_RS0131390* | 129 | Diels-Alderase | hypothetical protein CLV40_111123, *Actinokineospora auranticolor* (PPK66159.1); 60/76 | AbyU | AbsU | AbmU |
| *IF37_RS0131395* | 342 | 3-oxoacyl-ACP synthase III family protein | 3-oxoacyl-ACP synthase, *Frankia* sp. Cc1.17 (OHV40305.1); 76/85 | - | - | - |
| *IF37_RS0131400* | 136 | hypothetical protein | - | - | - | - |
| *IF37_RS0131405* | - | ABC transporter ATP-binding protein | - | - | - | - |
| *IF37_RS0131410* | 792 | PAS domain S-box protein | PAS domain-containing protein, *Kitasatospora* sp. OK780 (WP_100891745.1); 57/68 | - | - | - |

**Table S58.** Predicted functions of ORFs in potential abyssomicin BGC from *Streptomyces* sp. NRRL WC-3742 (NZ_JOCF01000060.1).

| ORF | Size (aa) | Proposed function | Closest homolog, host (protein ID); Identity/Similarity (%) | Aby homolog | Abs homolog | Abm homolog |
|---|---|---|---|---|---|---|
| *IH61_RS0128745* | 213 | TetR/AcrR family transcriptional regulator | TetR/AcrR family transcriptional regulator, *Streptomyces* sp. CB03911 (WP_073928463.1); 87/95 | AbyC | - | AbmC |
| *IH61_RS0128750* | 320 | alpha/beta hydrolase | alpha/beta hydrolase, *Streptomyces* sp. CB03911 (WP_079198449.1); 78/86 | - | - | - |
| *IH61_RS0128755* | 577 | ABC transporter substrate-binding protein | ABC transporter substrate-binding protein, *Streptacidiphilus* sp. DSM 106435 (WP_111490404.1); 76/84 | AbyF1 | AbsF1 | AbmF1 |
| *IH61_RS0128760* | 311 | ABC transporter permease | ABC transporter permease, *Streptomyces* sp. CB03911 (WP_073928465.1); 90/95 | AbyF2 | AbsF2 | AbmF2 |
| *IH61_RS0128765* | 269 | ABC transporter permease | ABC transporter permease, *Streptomyces* sp. CB03911 (WP_073928702.1); 87/93 | AbyF3 | AbsF3 | AbmF3 |
| *IH61_RS0128770* | 561 | ABC transporter ATP-binding protein | ABC transporter ATP-binding protein, *Streptacidiphilus* sp. DSM 106435 (WP_111490407.1); 79/86 | AbyF4 | AbsF4 | AbmF4 |
| *IH61_RS0128775* | 373 | NtaA/DmoA family FMN-dependent monooxygenase | LLM class flavin-dependent oxidoreductase, *Streptomyces* sp. CB03911 (WP_073928466.1); 90/93 | - | - | - |
| *IH61_RS0128780* | 340 | LLM class flavin-dependent oxidoreductase | LLM class flavin-dependent oxidoreductase, *Streptacidiphilus* sp. DSM 106435 (WP_111490409.1); 86/89 | AbyE | AbsE | AbmE1 |
| *IH61_RS0128785* | 347 | LLM class flavin-dependent oxidoreductase | LLM class flavin-dependent oxidoreductase, *Streptacidiphilus* sp. DSM 106435 (WP_111490410.1); 85/93 | - | - | AbmE2 |
| *IH61_RS0128790* | 1061 | PKS I | acyltransferase domain-containing protein, *Streptacidiphilus* sp. DSM 106435 (WP_111490411.1); 81/86 | AbyB3 | AbsB3 | AbmB3 |
| *IH61_RS46315* | 555 | PKS I | SDR family NAD(P)-dependent oxidoreductase, *Streptacidiphilus* sp. DSM 106435 (WP_114914558.1); 77/82 | AbyB2 | AbyB2 | AbmB2 |
| *IH61_RS44985* | 2041 | PKS I | hypothetical protein, *Streptacidiphilus* sp. DSM 106435 (WP_114914555.1); 81/88 | AbyB1 | AbyB1 | AbmB1 |
| *IH61_RS0128805* | 343 | 3-oxoacyl-ACP synthase III family protein | 3-oxoacyl-ACP synthase III family protein, *Streptacidiphilus* sp. DSM 106435 (WP_111492779.1); 85/90 | AbyA1 | AbsA1 | AbmA1 |
| *IH61_RS0128810* | 125 | Diels-Alderase | hypothetical protein, *Streptomyces* sp. CB03911 (WP_073928710.1); 86/92 | AbyU | AbsU | AbmU |
| *IH61_RS0128815* | 399 | cytochrome P450 | cytochrome, *Streptomyces* sp. CB03911 (OKI12689.1); 88/93 | AbyX/AbyV | AbsV/AbsX | AbmV |
| *IH61_RS0128820* | 79 | ferredoxin | ferredoxin, *Streptomyces* sp. CB03911 (WP_073928505.1); 74/83 | - | AbsG2/AbsG1 | AbmG |
| *IH61_RS0128825* | 480 | DHA2 family efflux MFS transporter permease subunit | DHA2 family efflux MFS transporter permease subunit, *Streptacidiphilus* sp. DSM 106435 (WP_111492776.1); 87/93 | AbyD | AbsD | AbyD |
| *IH61_RS0128830* | 384 | acyltransferase | acyltransferase, *Streptomyces* sp. CB03911 (WP_073928506.1); 77/86 | - | AbsI | - |
| *IH61_RS0128835* | 332 | aldo/keto reductase | aldo/keto reductase, *Streptomyces* sp. CB03911 (WP_073928507.1); 90/93 | - | AbsJ | AbmJ |
| *IH61_RS0128840* | 272 | AfsR/SARP family transcriptional regulator | AfsR/SARP family transcriptional regulator, *Streptacidiphilus* sp. DSM 106435 (WP_111492773.1); 89/93 | AbyI/AbyR | - | AbmI |
| *IH61_RS0128845* | 618 | LuxR family transcriptional regulator | LuxR family transcriptional regulator, *Streptacidiphilus* sp. DSM 106435 (WP_114914553.1); 66/75 | AbyH | - | AbmH |
| *IH61_RS0128850* | 288 | acyltransferase | acyltransferase, *Streptomyces* sp. CB03911 (WP_073928512.1); 80/85 | AbyA4 | AbsA4 | AbmA4 |

| | | | | | | |
|---|---|---|---|---|---|---|
| *IH61_RS0128855* | 362 | alpha/beta hydrolase | alpha/beta hydrolase, *Streptomyces* sp. CB03911 (WP_073928513.1); 84/91 | AbyA5 | AbsA5 | AbmA5 |
| *IH61_RS0128860* | 298 | AfsR/SARP family transcriptional regulator | AfsR/SARP family transcriptional regulator, *Streptomyces* sp. CB03911 (WP_073928514.1); 76/84 | AbyI | - | AbmI |
| *IH61_RS0128865* | 254 | thioesterase | thioesterase, *Streptomyces* sp. CB03911 (WP_079198461.1); 72/78 | AbyT | AbsN | AbmT |

**Table S59.** Predicted functions of ORFs in potential tetronomycin BGC from *Streptomyces olindensis* DAUFPE 5622 (JJOH01000019.1).

| **ORF** | **Size (aa)** | **Proposed function** | **Closest homolog, host (protein ID); Identity/Similarity (%)** | **Aby homolog** | **Abs homolog** | **Abm homolog** |
|---|---|---|---|---|---|---|
| *DF19_21795* | 406 | glucose-1-phosphate adenylyltransferase | glucose-1-phosphate adenylyltransferase, *Streptomyces* sp. Go-475 (WP_114256683.1); 99/100 | - | - | - |
| *DF19_21800* | 383 | glycosyl transferase family 1 | glycogen synthase, *Streptomyces* sp. Go-475 (WP_114256682.1); 99/99 | - | - | - |
| *DF19_21805* | 245 | hypothetical protein | (2Fe-2S)-binding protein, *Streptomyces* sp. Go-475 (WP_114257751.1); 85/86 | - | - | - |
| *DF19_21810* | 414 | membrane protein | hypothetical protein, *Streptomyces* sp. Go-475 (WP_114256681.1); 86/88 | - | - | - |
| *DF19_21815* | 424 | peptidase | peptidase, *Streptomyces* sp. Go-475 (WP_114256680.1); 93/96 | - | - | - |
| *DF19_21820* | 479 | 6-phosphogluconate dehydrogenase | NADP-dependent phosphogluconate dehydrogenase, *Streptomyces* sp. Go-475 (WP_114256679.1); 99/99 | - | - | - |
| *DF19_21825* | 114 | acetyltransferase | N-acetyltransferase, *Streptomyces* sp. Go-475 (WP_114256678.1); 92/95 | - | - | - |
| *DF19_21830* | 139 | aspartate 1-decarboxylase subunit alpha | aspartate 1-decarboxylase, *Streptomyces* sp. Go-475 (WP_114256677.1); 99/99 | - | - | - |
| *DF19_21835* | 75 | hypothetical protein | hypothetical protein, *Streptomyces* sp. 57 (WP_121408886.1); 47/62 | - | - | - |
| *DF19_21840* | 842 | hypothetical protein | helix-turn-helix transcriptional regulator, *Streptomyces* sp. CNH099 (WP_078627912.1); 62/75 | - | - | - |
| *DF19_21845* | 263 | oleoyl-ACP hydrolase | thioesterase, *Streptomyces* sp. CNZ306 (WP_100303313.1); 79/84 | - | - | - |
| *DF19_21850* | 75 | acyl carrier protein | acyl carrier protein, *Streptomyces* sp. CNH099 (WP_027756637.1); 89/96 | - | - | - |
| *DF19_21855* | 278 | acyltransferase | acyltransferase, *Streptomyces* sp. WAC 06738 (WP_125933459.1); 82/90 | - | - | - |
| *DF19_21860* | 4968 | PKS I | type I polyketide synthase, *Streptomyces* sp. CMB-StM0423 (WP_101426746.1); 73/79 | - | - | - |
| *DF19_21865* | 223 | PKS I | DedA family protein, *Streptomyces* sp. CNS335 (WP_018844985.1); 77/88 | - | - | - |
| *DF19_21870* | 173 | Diels-Alderase | hypothetical protein AA958_00640, *Streptomyces* sp. CNQ-509 (AKH80919.1); 67/77 | AbyU | AbsU | AbmU |
| *DF19_21875* | 500 | PKS I | hypothetical protein, *Streptomyces* sp. WAC 06738 (WP_125933460.1); 83/89 | - | - | - |
| *DF19_21880* | 1582 | PKS I | SDR family NAD(P)-dependent oxidoreductase, *Streptomyces* sp. WAC 06738 (WP_125933461.1); 73/80 | - | - | - |
| *DF19_21885* | 146 | PKS I | hypothetical protein, *Streptomyces* sp. CNQ329 (WP_027774440.1); 76/84 | - | - | - |
| *DF19_21890* | 304 | PKS I | erythromycin 3''-O-methyltransferase, *Streptomyces* sp. CNZ306 (PJJ38656.1); 85/93 | - | - | - |
| *DF19_21895* | 1664 | PKS I | type I polyketide synthase, *Streptomyces* sp. CNT371 (WP_027746214.1); 77/83 | - | - | - |

| | | | | | | |
|---|---|---|---|---|---|---|
| *DF19_21900* | 3666 | PKS I | type I polyketide synthase, *Streptomyces* sp. CNQ-509 (WP_047014289.1); 76/81 | - | - | - |
| *DF19_21905* | 3828 | PKS I | type I polyketide synthase, *Streptomyces* sp. CNQ-509 (WP_047014291.1); 70/79 | - | - | - |
| *DF19_21910* | 463 | enterotoxin | FAD-dependent oxidoreductase,*Streptomyces* sp. CNQ329 (WP_027770074.1); 85/89 | - | - | - |
| *DF19_21915* | 73 | hypothetical protein | ferredoxin, *Streptomyces* sp. CNH099 (WP_027757646.1); 73/83 | - | - | - |
| *DF19_21920* | 400 | cytochrome P450 | cytochrome P450, *Streptomyces* sp. CNZ306 (PJJ38667.1); 92/94 | - | - | - |
| *DF19_21925* | 342 | 3-oxoacyl-ACP synthase | 3-oxoacyl-ACP synthase III family protein, *Streptomyces* sp. CNQ-509 (WP_047014295.1); 87/95 | - | - | - |
| *DF19_21930* | 173 | MarR family transcriptional regulator | MarR family transcriptional regulator, *Actinopolyspora mzabensis* (WP_092625240.1); 55/71 | - | - | - |
| *DF19_21935* | 632 | methoxymalonyl-ACP biosynthesis protein FkbH | HAD-IIIC family phosphatase, *Streptomyces* sp. CNH099 (WP_027757643.1); 85/91 | - | - | - |
| *DF19_21940* | 345 | hypothetical protein | alpha/beta fold hydrolase, *Streptomyces* sp. CNQ-509 (WP_047014297.1); 75/84 | - | - | - |
| *DF19_21945* | 543 | hypothetical protein | anibiotic ABC transporter, *Plantactinospora* sp. CNZ321 (WP_130464162.1); 50/67 | - | - | - |
| *DF19_21950* | 306 | ABC transporter ATP-binding protein | ABC transporter ATP-binding protein, *Rhodococcus* sp. S2-17 (WP_109331707.1); 70/80 | - | - | - |
| *DF19_21955* | 256 | activator protein | activator protein, *Streptomyces* sp. CMB-StM0423 (AUH44525.1); 89/93 | - | - | - |
| *DF19_21960* | 776 | hypothetical protein | type I polyketide synthase-related protein, *Streptomyces* sp. NRRL 11266 (BAE93739.1); 66/74 | - | - | - |

**Table S60.** Predicted functions of ORFs in quartromicin BGC from *Amycolatopsis orientalis* Q427-8 (JF970188.1).

| ORF | Size (aa) | Proposed function | Closest homolog, host (protein ID); Identity/Similarity (%) | Aby homolog | Abs homolog | Abm homolog |
|---|---|---|---|---|---|---|
| *QmnB* | 471 | propionyl-CoA carboxylase | acyl-CoA carboxylase subunit beta, *Amycolatopsis albispora* (WP_113696554.1); 97/98# | - | - | - |
| *QmnC* | 251 | thioesterase | thioesterase, *Amycolatopsis albispora* (WP_113696553.1); 95/98 | - | - | - |
| *QmnD4* | 322 | 2-oxoacid dehydrogenase, acyltransferase | alpha/beta hydrolase, *Amycolatopsis albispora* (WP_113696552.1); 90/93 | - | - | - |
| *QmnA1* | 5924 | PKS I | type I polyketide synthase, *Streptomyces cellostaticus* (WP_067002540.1); 54/64 | - | - | - |
| *QmnA2* | 1771 | PKS I | SDR family NAD(P)-dependent oxidoreductase, *Amycolatopsis albispora* (WP_113696551.1); 93/96 | - | - | - |
| *QmnA3* | 1292 | PKS I | acyltransferase domain-containing protein, *Amycolatopsis albispora* (WP_113696550.1); 86/90 | - | - | - |
| *QmnE* | 412 | - | serine hydrolase, *Sorangium cellulosum* (KYF55935.1); 58/71 | - | - | - |
| *QmnF* | 396 | - | cysteine desulfurase-like protein, *Amycolatopsis albispora* (WP_113696547.1); 89/92 | - | - | - |
| *QmnRg 1* | 255 | - | activator protein, *Amycolatopsis albispora* (WP_113696546.1); 98/98 | - | - | - |
| *QmnG* | 533 | PQQ-dependent dehydrogenase | hypothetical protein, *Amycolatopsis albispora* (WP_113696544.1); 93/95 | - | - | - |
| *QmnH* | 376 | Diels-Alderase | hypothetical protein, *Amycolatopsis albispora* (WP_113696543.1); 94/97 | AbyU | AbsU | AbmU |
| *QmnI* | 348 | - | HlyD family efflux transporter periplasmic adaptor subunit, *Amycolatopsis albispora* (WP_113696542.1); 88/93 | - | - | - |
| *QmnJ* | 161 | - | hypothetical protein, *Amycolatopsis albispora* (WP_113696541.1); 88/93 | - | - | - |
| *QmnRs1* | 392 | transporter | ABC transporter permease, *Amycolatopsis albispora* (WP_113696540.1); 97/98 | - | - | - |
| *QmnRs2* | 227 | transporter | ABC transporter ATP-binding protein, *Amycolatopsis albispora* (WP_113696539.1); 96/99 | - | - | - |
| *QmnK* | 352 | - | HlyD family efflux transporter periplasmic adaptor subunit, *Amycolatopsis albispora* (WP_113696538.1); 96/97 | - | - | - |
| *QmnL* | 152 | - | hypothetical protein, *Amycolatopsis albispora* (WP_113696537.1); 88/90 | - | - | - |
| *QmnM* | 219 | - | hypothetical protein, *Streptomyces rimosus* (WP_030595123.1); 58/69 | - | - | - |
| *QmnN* | 130 | | hypothetical protein, *Streptomyces rimosus* (WP_030674267.1); 65/74 | - | - | - |
| *QmnRg 2* | 449 | regulator | HAMP domain-containing protein, *Amycolatopsis albispora* (WP_113696536.1); 90/94 | - | - | - |
| *QmnRg 3* | 222 | regulator | response regulator transcription factor, *Amycolatopsis albispora* (WP_113698159.1); 99/99 | - | - | - |
| *QmnD5* | 343 | 3-oxoacyl-ACP synthase III (KS) | 3-oxoacyl-ACP synthase III family protein, *Amycolatopsis albispora* (WP_113696535.1); 95/98 | - | - | - |
| *QmnD1* | 613 | glyceryltransferase/phosphatase | HAD-IIIC family phosphatase, *Amycolatopsis albispora* (WP_113696534.1); 91/94 | - | - | - |
| *QmnD2* | 71 | ACP | acyl carrier protein, *Amycolatopsis albispora* (WP_113696533.1); 94/97 | - | - | - |

| | | | | | | |
|---|---|---|---|---|---|---|
| *QmnD3* | 281 | 2-oxoacid dehydrogenase, acyltransferase | acyltransferase, *Amycolatopsis albispora* (WP_113696532.1); 83/87 | - | - | - |
| *QmnO* | 403 | cytochrome p450 | cytochrome P450, *Amycolatopsis albispora* (WP_113696531.1); 95/97 | - | - | - |
| *QmnRg 4* | 935 | regulator | LuxR family transcriptional regulator, *Amycolatopsis albispora* (WP_113698158.1); 95/96 | - | - | - |
| *QmnRg 5* | 821 | regulator | transcriptional regulator, *Amycolatopsis albispora* (WP_113696530.1); 90/92 | - | - | - |

**Table S61.** Predicted functions of ORFs in potential BGC from *Pantoea* sp. A4 (NZ_ALXE01000017).

| ORF | Size (aa) | Proposed function | Closest homolog, host (protein ID); Identity/Similarity (%) | Aby homolog | Abs homolog | Abm homolog |
|---|---|---|---|---|---|---|
| *B880_RS0103640* | 163 | DMT family transporter | DMT family transporter, *Enterobacteriaceae bacterium* strain FGI 57 (WP_015965015.1); 94/97 | - | - | - |
| *B880_RS0103645* | 458 | hypothetical protein | MmgE/PrpD family protei, *Serratia* sp. P2ACOL2 (WP_122079351.1); 74/83 | - | - | - |
| *B880_RS0103650* | 503 | DHA2 family efflux MFS transporter permease subunit | Multidrug resistance protein stp, *Serratia marcescens* (SAY43176.1); 72/83 | - | - | - |
| *B880_RS0103655* | 212 | TetR/AcrR family transcriptional regulator | TetR family transcriptional regulator, *Yersinia intermedia* (CQD48424.1); 71/81 | - | - | - |
| *B880_RS0103660* | 385 | ABC transporter substrate-binding protein | ABC transporter substrate-binding protein, *Serratia* sp. P2ACOL2 (WP_122079353.1); 64/79 | - | - | - |
| *B880_RS0103665* | 715 | TonB-dependent siderophore receptor | TonB-dependent siderophore receptor, *Serratia* sp. P2ACOL2 (WP_122079354.1); 69/84 | - | - | - |
| *B880_RS0103670* | 73 | hypothetical protein | hypothetical protein, *Yersinia intermedia* (WP_050881852.1); 59/77 | - | - | - |
| *B880_RS0103675* | 535 | hypothetical protein | RosA, *Erwinia rhapontici* (AMB18979.1); 70/80 | - | - | - |
| *B880_RS0103680* | 242 | thioesterase | thioesterase, *Erwinia persicina* (WP_118665708.1); 58/72 | - | - | - |
| *B880_RS0103685* | 342 | pyridoxal-phosphate dependent enzyme | pyridoxal-phosphate dependent enzyme, *Erwinia rhapontici* (WP_133843159.1); 85/91 | - | - | - |
| *B880_RS0103690* | 184 | Diels-Alderase | RosD, *Erwinia rhapontici* (AMB18976.1); 78/89 | AbyU | AbsU | AbmU |
| *B880_RS0103695* | 391 | FAD-dependent oxidoreductase | - | - | - | - |
| *B880_RS0103700* | 961 | PKS I | acyltransferase domain-containing protein, *Erwinia persicina* (WP_062742714.1); 69/81 | - | - | - |
| *B880_RS0103705* | 344 | Glu/Leu/Phe/Val dehydrogenase | RosG, *Erwinia rhapontici* (AMB18973.1); 78/87 | - | - | - |
| *B880_RS0103710* | 366 | hypothetical protein | hypothetical protein, *Serratia* sp. P2ACOL2 (WP_122079360.1); 59/71 | - | - | - |
| *B880_RS0103715* | 131 | hypothetical protein | DNA-binding protein, *Pantoea rwandensis* (WP_084932292.1); 61/75 | - | - | - |
| *B880_RS0103720* | 289 | aldo/keto reductase | aldo/keto reductase, *Escherichia marmotae* (PGF73638.1); 85/93 | - | - | - |
| *B880_RS0103725* | 439 | FAD-dependent oxidoreductase | FAD-binding oxidoreductase, *Pantoea wallisii* (WP_128601728.1); 81/93 | - | - | - |
| *B880_RS0103730* | 443 | MFS transporter | MFS transporter, *Pseudomonas reidholzensis* (WP_119143972.1); 77/89 | - | - | - |
| *B880_RS0103735* | 277 | transporter substrate-binding domain-containing protein | transporter substrate-binding domain-containing protein, *Pantoea* sp. YU22 (WP_126689428.1); 81/88 | - | - | - |

**Table S62.** Predicted functions of ORFs in potential abyssomicin BGC from *Streptomyces paucisporeus* CGMCC 4.2025 (NZ_FRBI01000008).

| ORF | Size (aa) | Proposed function | Closest homolog, host (protein ID); Identity/Similarity (%) | Aby homolog | Abs homolog | Abm homolog |
|---|---|---|---|---|---|---|
| *BUE44_RS14365* | - | PKS I | type I polyketide synthase, *Micromonospora* sp. GMKU326 (BAQ25511.1); 58/67 | PKS I | PKS I | PKS I |
| *BUE44_RS14370* | 505 | MFS transporter | MFS transporter, *Streptomyces griseorubiginosus* (WP_123763219.1); 58/74 | AbyD | AbsD | AbmD |
| *BUE44_RS14375* | 390 | cytochrome P450 | cytochrome P450, *Amycolatopsis* sp. CA-126428 (WP_103341807.1); 54/71 | AbyX/AbyV | AbsV/AbsX | AbmV |
| *BUE44_RS14380* | 102 | ferredoxin | ferredoxin, *Lentzea kentuckyensis* (WP_086665861.1); 63/77 | - | AbsG1/AbsG2 | AbmG |
| *BUE44_RS14385* | 388 | cytochrome P450 | cytochrome P450, *Amycolatopsis* sp. CA-126428 (WP_103341807.1); 54/69 | AbyX/AbyV | AbsV/AbsX | AbmV |
| *BUE44_RS14390* | 239 | SDR family NAD(P)-dependent oxidoreductase | SDR family oxidoreductase, *Streptoalloteichus hindustanus* (WP_073489819.1); 53/66 | - | - | - |
| *BUE44_RS14395* | 680 | FAD/NAD(P)-binding protein | FAD/NAD(P)-binding protein, *Streptomyces cattleya* (WP_014141705.1); 49/59 | - | - | - |
| *BUE44_RS14400* | 238 | hypothetical protein | acyltransferase, *Streptomyces* sp. WAC 06738 (WP_125933459.1); 67/78 | AbyA4 | AbsA4 | AbmA4 |
| *BUE44_RS14405* | 353 | 3-oxoacyl-ACP synthase III family protein | 3-oxoacyl-ACP synthase III family protein, *Streptomyces* sp. 2131.1 (WP_093710000.1); 66/77 | AbyA1 | AbsA1 | AbmA1 |
| *BUE44_RS14410* | 188 | Diels-Alderase | hypothetical protein, *Streptomyces griseorubiginosus* (WP_123763217.1); 44/58 | AbyU | AbsU | AbmU |
| *BUE44_RS14415* | 75 | acyl carrier protein | acyl carrier protein, *Umezawaea tangerina* (WP_106194434.1); 68/80 | AbyA3 | AbsA3 | AbmA3 |
| *BUE44_RS14420* | 258 | AfsR/SARP family transcriptional regulator | SARP family transcriptional regulator, *Streptomyces* sp. E14 (WP_009191683.1); 51/64 | AbyI/AbyR | - | AbmI |
| *BUE44_RS14425* | 273 | SAM-dependent methyltransferase | SAM-dependent methyltransferase, *Nonomuraea* sp. KC201 (WP_132330820.1); 53/65 | - | - | - |
| *BUE44_RS14430* | 202 | TetR/AcrR family transcriptional regulator | TetR/AcrR family transcriptional regulator, *Streptomyces hoynatensis* (WP_120678650.1); 50/60 | - | AbsC2 | - |
| *BUE44_RS14435* | 356 | LLM class flavin-dependent oxidoreductase | LLM class flavin-dependent oxidoreductase, *Streptomyces* f*ormicae* (WP_098241239.1); 59/73 | AbyE | AbsE | AbmE1 |
| *BUE44_RS14440* | 950 | LuxR family transcriptional regulator | LuxR family transcriptional regulator, *Streptomyces* sp. 57 (WP_121408891.1); 41/53 | AbyH | - | AbmH |
| *BUE44_RS14445* | 263 | AfsR/SARP family transcriptional regulator | SARP family transcriptional regulator, *Streptomyces* sp. E14 (WP_009191683.1); 62/77 | AbyI/AbyR | - | AbmI |
| *BUE44_RS14450* | 403 | cytochrome P450 | cytochrome P450, *Streptomyces griseoruber* (WP_055634261.1); 55/71 | AbyX/AbyV | AbsV/AbsX | AbmV |
| *BUE44_RS14455* | 277 | alpha/beta fold hydrolase | alpha/beta fold hydrolase, *Streptomyces* sp. MUSC 1 (WP_071384385.1); 65/77 | - | - | - |

**Table S63.** Predicted functions of ORFs in potential chlorothricin BGC from *Actinomadura pelletieri* DSM 43383 (NZ_RBWU01000008).

| ORF | Size (aa) | Proposed function | Closest homolog, host (protein ID); Identity/Similarity (%) | Aby homolog | Abs homolog | Abm homolog |
|---|---|---|---|---|---|---|
| *BZB76_RS31630* | 476 | NDP-hexose 2,3-dehydratase | ChlC3, *Streptomyces antibioticus* (AAZ77682.1); 65/73 | - | - | - |
| *BZB76_RS31635* | 79 | hypothetical protein | hypothetical protein, *Actinomadura* sp. LMG 30035 (WP_131741591.1); 48/62 | - | - | - |
| *BZB76_RS31640* | 262 | thioesterase | thioesterase, *Streptomyces armeniacus* (AXK32421.1); 59/69 | - | - | - |
| *BZB76_RS31645* | 266 | AfsR/SARP family transcriptional regulator | activator protein, *Streptomyces armeniacus* (AXK32420.1); 71/79 | - | - | - |
| *BZB76_RS31650* | 754 | MMPL family transporter | MMPL family transporter, *Aeromicrobium* sp. Root236 (WP_056402437.1); 56/72 | - | - | - |
| *BZB76_RS31655* | 368 | alpha/beta hydrolase | alpha/beta hydrolase, *Actinomadura chibensis* (WP_067904591.1); 67/77 | - | - | - |
| *BZB76_RS31660* | 266 | acyltransferase | acyltransferase, *Actinocrispum wychmicini* (WP_132116034.1); 71/81 | - | - | - |
| *BZB76_RS31665* | 74 | acyl carrier protein | acyl carrier protein, *Actinocrispum wychmicini* (WP_132116036.1); 60/73 | - | - | - |
| *BZB76_RS31670* | 636 | HAD-IIIC family phosphatase | HAD-IIIC family phosphatase, *Micromonospora* sp. RP3T (WP_107154962.1); 65/76 | - | - | - |
| *BZB76_RS31675* | 343 | 3-oxoacyl-ACP synthase III family protein | 3-oxoacyl-ACP synthase III family protein, *Actinocrispum wychmicini* (WP_132116040.1); 76/87 | - | - | - |
| *BZB76_RS31680* | 501 | hypothetical protein | hypothetical protein DVA86_06945, *Streptomyces armeniacus* (AXK32428.1); 61/71 | - | - | - |
| *BZB76_RS31685* | 1507 | acyltransferase domain-containing protein | acyltransferase domain-containing protein, *Streptomyces armeniacus* (AXK32430.1); 57/67 | - | - | - |
| *BZB76_RS31690* | 3910 | PKS I | type I polyketide synthase, *Micromonospora* sp. Rc5 (WP_077939335.1); 61/71 | - | - | - |
| *BZB76_RS31695* | 589 | PKS I | SDR family NAD(P)-dependent oxidoreductase, partial, *Candidatus Streptomyces philanthi* (WP_114025722.1); 53/67 | - | - | - |
| *BZB76_RS31700* | 377 | PKS I | type I polyketide synthase, *Streptomyces eurocidicus* (WP_102919106.1); 48/61 | - | - | - |
| *BZB76_RS31705* | 777 | PKS I | ChlA4, *Streptomyces antibioticus* (AAZ77697.1); 65/77 | - | - | - |
| *BZB76_RS31710* | 5163 | PKS I | SDR family NAD(P)-dependent oxidoreductase, *Streptomyces alboflavus* (WP_125262906.1); 53/65 | - | - | - |
| *BZB76_RS31715* | 1542 | PKS I | SDR family NAD(P)-dependent oxidoreductase, *Actinocrispum wychmicini* (WP_132116050.1); 57/67 | - | - | - |
| *BZB76_RS31720* | 263 | PKS I | SDR family NAD(P)-dependent oxidoreductase, *Actinocrispum wychmicini* (WP_132116052.1); 69/78 | - | - | - |
| *BZB76_RS31725* | 112 | PKS I | type I polyketide synthase, *Actinomadura macra* (WP_067456430.1); 62/69 | - | - | - |
| *BZB76_RS31730* | 316 | PKS I | polyketide synthase subunit, partial, *Streptomyces* sp. RSD-27 (KIF04675.1); 68/76 | - | - | - |
| *BZB76_RS31735* | 67 | PKS I | Acyl transferase domain-containing protein, partial, *Actinoplanes regularis* (SNT04067.1); 70/79 | - | - | - |
| *BZB76_RS31740* | 881 | PKS I | type I polyketide synthase, *Streptomyces hygroscopicus* (WP_066029228.1); 64/75 | - | - | - |
| *BZB76_RS31745* | 967 | PKS I | SDR family NAD(P)-dependent oxidoreductase, *Actinomadura* sp. LHW52907 (WP_117405125.1); 64/73 | - | - | - |

| | | | | | | |
|---|---|---|---|---|---|---|
| *BZB76_RS31750* | 375 | PKS I | SDR family NAD(P)-dependent oxidoreductase, *Actinomadura* sp. LHW52907 (WP_117405125.1); 55/66 | - | - | - |
| *BZB76_RS31755* | 336 | PKS I | type I polyketide synthase, partial, *Streptomyces* sp. CNQ766 (WP_018840958.1); 61/69 | - | - | - |
| *BZB76_RS31760* | 251 | PKS I | KR domain-containing protein, partial, *Streptomyces* sp. AZ1-7 (WP_120745073.1); 46/55 | - | - | - |
| *BZB76_RS31765* | 1835 | PKS I | Acyl transferase domain-containing protein, *Streptomyces* sp. 2314.4 (SEE65811.1); 55/66 | - | - | - |
| *BZB76_RS31770* | 355 | glucose-1-phosphate thymidylyltransferase | ChlC1, *Streptomyces antibioticus* (AAZ77690.1); 64/75 | - | - | - |
| *BZB76_RS31775* | 579 | 3-hydroxyacyl-CoA dehydrogenase family protein | 3-hydroxyacyl-CoA dehydrogenase family protein, *Streptomyces sioyaensis* (WP_129246466.1); 63/73 | - | - | - |
| *BZB76_RS31780* | 347 | beta-ketoacyl-ACP synthase 3 | ketoacyl-ACP synthase III, *Streptomyces* sp. C (WP_007269134.1); 70/80 | - | - | - |
| *BZB76_RS31785* | 444 | crotonyl-CoA carboxylase/reductase | crotonyl-CoA carboxylase/reductase, *Streptoalloteichus hindustanus* (SHG03036.1); 81/91 | - | - | - |
| *BZB76_RS31790* | 89 | acyl carrier protein | ChlB2, *Streptomyces antibioticus* (AAZ77675.1); 51/67 | - | - | - |
| *BZB76_RS31795* | 453 | NAD(P)/FAD-dependent oxidoreductase | FAD-dependent oxidoreductase, *Actinocrispum wychmicini* (WP_132116064.1); 79/88 | - | - | - |
| *BZB76_RS31800* | 1785 | acyltransferase domain-containing protein | acyltransferase domain-containing protein, *Streptomyces armeniacus* (AXK33367.1); 66/74 | - | - | - |
| *BZB76_RS31805* | 347 | 3-oxoacyl-ACP synthase | 3-oxoacyl-ACP synthase, *Actinocrispum wychmicini* (WP_132116060.1); 74/84 | - | - | - |
| *BZB76_RS31810* | 67 | hypothetical protein | - | - | - | - |
| *BZB76_RS31815* | 186 | Diels-Alderase | hypothetical protein, *Actinocrispum wychmicini* (WP_132116074.1); 45/57 | AbyU | AbsU | AbmU |
| *BZB76_RS31820* | 384 | DegT/DnrJ/EryC1/StrS family aminotransferase | DegT/DnrJ/EryC1/StrS family aminotransferase, *Streptomyces exfoliatus* (WP_030554129.1); 69/81 | - | - | - |
| *BZB76_RS31825* | 348 | 3-oxoacyl-ACP synthase | 3-oxoacyl-ACP synthase, *Amycolatopsis palatopharyngis* (WP_116051422.1); 51/69 | - | - | - |
| *BZB76_RS31830* | 406 | DUF1205 domain-containing protein | DUF1205 domain-containing protein, *Actinocrispum wychmicini* (WP_132116056.1); 48/65 | - | - | - |
| *BZB76_RS31835* | 405 | cytochrome P450 | cytochrome P450, *Streptomyces* sp. LHW50302 (WP_114017402.1); 65/75 | - | - | - |

**Table S64.** Predicted functions of ORFs in potential BGC from *Candidatus Streptomyces philanthi* (NZ_QOIN01000036).

| ORF | Size (aa) | Proposed function | Closest homolog, host (protein ID); Identity/Similarity (%) | Aby homolog | Abs homolog | Abm homolog |
|---|---|---|---|---|---|---|
| *DTL70_RS08705* | - | PKS I | polyketide synthase, partial, *Streptomyces* sp. LHW50302 (RCG16060.1); 90/93 | - | - | - |
| *DTL70_RS08710* | 201 | Diels-Alderase | hypothetical protein, *Streptomyces* sp. LHW50302 (WP_114017401.1); 98/99 | AbyU | AbsU | AbmU |
| *DTL70_RS08715* | 403 | cytochrome P450 | cytochrome P450, *Streptomyces* sp. LHW50302 (WP_114017402.1); 99/99 | - | - | - |
| *DTL70_RS08720* | 253 | methyltransferase | methyltransferase, *Streptomyces* sp. LHW50302 (WP_114017403.1); 98/99 | - | - | - |
| *DTL70_RS08725* | 347 | 3-oxoacyl-ACP synthase | 3-oxoacyl-ACP synthase, *Streptomyces* sp. LHW50302 (WP_114017404.1); 99/99 | - | - | - |
| *DTL70_RS08730* | 1817 | PKS I | type I polyketide synthase, *Streptomyces* sp. LHW50302 (WP_114017405.1); 94/95 | - | - | - |
| *DTL70_RS08735* | 323 | dTDP-glucose 4,6-dehydratase | dTDP-glucose 4,6-dehydratase, *Streptomyces* sp. LHW50302 (WP_114017406.1); 98/99 | - | - | - |
| *DTL70_RS08740* | 974 | helix-turn-helix transcriptional regulator | helix-turn-helix transcriptional regulator, *Streptomyces* sp. LHW50302 (WP_114017407.1); 95/96 | - | - | - |
| *DTL70_RS08745* | 776 | DUF2075 domain-containing protein | DUF2075 domain-containing protein, *Streptomyces* sp. LHW50302 (WP_114017408.1); 97/98 | - | - | - |
| *DTL70_RS08750* | 202 | dTDP-4-keto-6-deoxy-D-glucose epimerase | dTDP-4-keto-6-deoxy-D-glucose epimerase, *Streptomyces* sp. LHW50302 (WP_114017409.1); 99/99 | - | - | - |
| *DTL70_RS08755* | 307 | putative sugar O-methyltransferase | NanM, *Streptomyces nanchangensis* (AAP42862.1); 61/77 | - | - | - |
| *DTL70_RS08760* | 403 | DUF1205 domain-containing protein | DUF1205 domain-containing protein, *Streptomyces* sp. LHW50302 (WP_114017411.1); 97/97 | - | - | - |
| *DTL70_RS08765* | 265 | class I SAM-dependent methyltransferase | class I SAM-dependent methyltransferase, *Streptomyces* sp. LHW50302 (WP_114017412.1); 98/100 | - | - | - |
| *DTL70_RS08770* | 415 | class I SAM-dependent methyltransferase | class I SAM-dependent methyltransferase, *Streptomyces* sp. LHW50302 (WP_114017413.1); 99/99 | - | - | - |

**Table S65.** Predicted functions of ORFs in potential BGC from *Streptomyces rimosus subsp. rimosus* NRRL B-16073 (NZ_JNWX01000004).

| ORF | Size (aa) | Proposed function | Closest homolog, host (protein ID); Identity/Similarity (%) | Aby homolog | Abs homolog | Abm homolog |
|---|---|---|---|---|---|---|
| *NH06_RS0106110* | 533 | amidase | amidase, *Streptomyces* sp. NRRL WC-3701 (KOT47522.1); 99/99 | - | - | - |
| *NH06_RS0106115* | 283 | oxidoreductase | SDR family NAD(P)-dependent oxidoreductase, *Streptomyces* sp. WAC 06783 (WP_125520864.1); 98/98 | - | - | - |
| *NH06_RS0106120* | 290 | transcriptional regulator | helix-turn-helix domain-containing protein, *Streptomyces* sp. WAC 06783 (WP_125520863.1); 100/100 | - | - | - |
| *NH06_RS0106125* | 421 | DegT/DnrJ/EryC1/StrS family aminotransferase | DegT/DnrJ/EryC1/StrS family aminotransferase, *Streptomyces albus* (WP_060732997.1); 99/99 | - | - | - |
| *NH06_RS0106130* | 335 | streptomycin biosynthesis protein | streptomycin biosynthesis protein, *Streptomyces* sp. WAC 06783 (WP_125520938.1); 99/99 | - | - | - |
| *NH06_RS0106135* | 79 | acyl carrier protein | acyl carrier protein, *Streptomyces* sp. WAC 06783 (RSO08986.1); 99/100 | - | - | - |
| *NH06_RS0106140* | 570 | (2,3-dihydroxybenzoyl)adenylate synthase | 2,3-dihydroxybenzoate--AMP ligase, *Kitasatospora aureofaciens* (KOG75552.1); 99/99 | - | - | - |
| *NH06_RS0106145* | 255 | thioesterase | alpha/beta fold hydrolase, *Streptomyces* sp. WAC 06725 (WP_125532514.1); 99/99 | - | - | - |
| *NH06_RS0106150* | 402 | FAD-dependent oxidoreductase | FAD-dependent oxidoreductase, *Streptomyces* sp. WAC 06725 (WP_125532515.1); 99/99 | - | - | - |
| *NH06_RS0106155* | 979 | PKS I | acyltransferase domain-containing protein, *Streptomyces* sp. WAC 06725 (WP_125532516.1); 99/98 | - | - | - |
| *NH06_RS0106160* | 173 | CGNR zinc finger domain-containing protein | CGNR zinc finger domain-containing protein, *Streptomyces* sp. WAC 06725 (RSO35445.1); 99/99 | - | - | - |
| *NH06_RS0106165* | 448 | MFS transporter | MFS transporter, *Streptomyces* sp. WAC 06783 (WP_125520937.1); 99/99 | - | - | - |
| *NH06_RS0106170* | 174 | Diels-Alderase | hypothetical protein DMH18_19415, *Streptomyces* sp. WAC 06783 (RSO08979.1); 99/100 | AbyU | AbsU | AbmU |
| *NH06_RS0106175* | 433 | MFS transporter | MFS transporter, *Kitasatospora aureofaciens* (KOG75546.1); 99/99 | - | - | - |
| *NH06_RS0106180* | 435 | hypothetical protein | hypothetical protein, *Streptomyces* sp. NRRL F-5755 (WP_053700239.1); 99/99 | - | - | - |
| *NH06_RS0106185* | 367 | UDP-N-acetylmuramate dehydrogenase | UDP-N-acetylmuramate dehydrogenase, *Streptomyces* sp. NRRL F-5755 (WP_053700238.1); 99/98 | - | - | - |
| *NH06_RS0106190* | 332 | iron ABC transporter | iron ABC transporter, *Streptomyces* sp. WAC 06783 (RSO09087.1); 99/99 | - | - | - |
| *NH06_RS0106195* | 376 | iron ABC transporter permease | iron ABC transporter permease, *Kitasatospora aureofaciens* (KOG75678.1); 99/100 | - | - | - |
| *NH06_RS0106200* | 331 | iron siderophore-binding protein | ABC transporter substrate-binding protein, *Streptomyces* sp. WAC 06783 (WP_125520857.1); 99/99 | - | - | - |

**Table S66.** Predicted functions of ORFs in abyssomicin BGC from *Streptomyces* sp. Amel2xE9 (NZ_KB912999 and NZ_KB912981).

| **ORF** | **Size (aa)** | **Proposed function** | **Closest homolog, host (protein ID); Identity/Similarity (%)** | **Aby homolog** | **Abs homolog** | **Abm homolog** |
|---|---|---|---|---|---|---|
| *B065_RS0132280* | 257 | AfsR/SARP family transcriptional regulator | SARP family transcriptional regulator, *Streptomyces* sp. E14 (WP_009191683.1); 99/99 | AbyI/AbyR | - | AbmI |
| *B065_RS0132285* | 945 | LuxR family transcriptional regulator | transcription regulator, *Streptomyces* sp. LC-6-2 (ARE67835.1); 99/99 | AbyH | - | AbmH |
| *B065_RS0132290* | 476 | MFS transporter | MFS transporter, *Streptomyces* sp. E14 (WP_043261567.1); 99/99 | AbyD | AbsD | AbmD |
| *B065_RS0132295* | 196 | TetR/AcrR family transcriptional regulator | AbsC2, *Streptomyces* sp. LC-6-2 (ARE67837.1); 99/100 | - | AbsC2 | - |
| *B065_RS0132300* | 128 | Diels-Alderase | conserved hypothetical protein, *Streptomyces* sp. E14 (EFF94138.1); 99/99 | AbyU | AbsU | AbmU |
| *B065_RS0132305* | 344 | aldo/keto reductase | aldo/keto reductase, *Streptomyces* sp. E14 (WP_050790870.1); 99/99 | - | AbsJ | AbmJ |
| *B065_RS0132310* | 64 | ferredoxin | AbsG2, *Streptomyces* sp. LC-6-2 (ARE67840.1); 100;100 | - | AbsG2 | - |
| *B065_RS0132315* | 403 | cytochrome P450 | cytochrome P450, *Streptomyces* sp. E14 (WP_009191676.1); 99/99 | AbyX | AbsX | - |
| *B065_RS0132320* | 387 | acyltransferase | acyltransferase, *Streptomyces* sp. E14 (WP_009191675.1); 99/99 | - | AbsI | - |
| *B065_RS0132325* | 68 | ferredoxin | AbsG1, *Streptomyces* sp. LC-6-2 (ARE67843.1); 100/100 | - | AbsG1 | AbmG |
| *B065_RS0132330* | 397 | cytochrome P450 | AbsV, *Streptomyces* sp. LC-6-2 (ARE67844.1); 99/100 | AbyV | AbsV | AbmV |
| *B065_RS0132335* | 555 | ABC transporter ATP-binding protein | AbsF4, *Streptomyces* sp. LC-6-2 (ARE67845.1); 98/98 | AbyF4 | AbsF4 | AbmF4 |
| *B065_RS0132340* | 249 | ABC transporter permease | AbsF3, *Streptomyces* sp. LC-6-2 (ARE67846.1); 99/99 | AbyF3 | AbsF3 | AbmF3 |
| *B065_RS0132345* | 333 | ABC transporter permease | AbsF2, *Streptomyces* sp. LC-6-2 (ARE67847.1); 99/99 | AbyF2 | AbsF2 | AbmF2 |
| *B065_RS0132350* | 554 | ABC transporter substrate-binding protein | AbsF1, *Streptomyces* sp. LC-6-2 (ARE67848.1); 99/99 | AbyF1 | AbsF1 | AbmF1 |
| *B065_RS0132355* | 328 | LLM class flavin-dependent oxidoreductase | AbsE, *Streptomyces* sp. LC-6-2 (ARE67849.1); 99/99 | AbyE | AbsE | AbmE1 |
| *B065_RS0132360* | 649 | HAD-IIIC family phosphatase | AbsA2, *Streptomyces* sp. LC-6-2 (ARE67850.1); 99/99 | AbyA2 | AbsA2 | AbmA2 |
| *B065_RS0132365* | 1049 | PKS I | AbsB3, *Streptomyces* sp. LC-6-2 (ARE67851.1); 98/98 | AbyB3 | AbsB3 | AbmB3 |
| *B065_RS0132370* | 3662 | PKS I | AbsB2, *Streptomyces* sp. LC-6-2 (ARE67852.1); 96/96 | AbyB2 | AbsB2 | AbmB2 |
| *B065_RS39245* | - | PKS I | AbsB1, *Streptomyces* sp. LC-6-2 (ARE67853.1); 97/97 | AbyB1 | AbsB1 | AbmB1 |
| //////////////////////////////////////////////////////////////////////// | | | | | | |
| *B065_RS38160* | - | PKS I | AbsB1, *Streptomyces* sp. LC-6-2 (ARE67853.1); 98/98 | AbyB1 | AbsB1 | AbmB1 |
| *B065_RS41620* | 69 | hypothetical protein | - | - | - | - |
| *B065_RS0128140* | 275 | thioesterase | thioesterase, *Streptomyces* sp. E14 (WP_009191660.1); 99/99 | AbyT | AbsN | AbmT |
| *B065_RS0128145* | 381 | alpha/beta hydrolase | AbsA5, *Streptomyces* sp. LC-6-2 (ARE67854.1); 98/98 | AbyA5 | AbsA5 | AbmA5 |

| | | | | | | |
|---|---|---|---|---|---|---|
| *B065_RS0128150* | 251 | acyltransferase | AbsA4, *Streptomyces* sp. LC-6-2 (ARE67855.1); 100/100 | AbyA4 | AbsA4 | AbmA4 |
| *B065_RS0128155* | 77 | acyl carrier protein | AbsA3, *Streptomyces* sp. LC-6-2 (ARE67856.1); 97/97 | AbyA3 | AbsA3 | AbmA3 |
| *B065_RS0128160* | 351 | 3-oxoacyl-ACP synthase III family protein | 3-oxoacyl-ACP synthase III family protein, *Streptomyces* sp. E14 (WP_063821841.1); 99/99 | AbyA1 | AbsA1 | AbmA1 |
| *B065_RS0128165* | 190 | flavin reductase family protein | flavin reductase domain-containing protein, *Streptomyces* sp. E14 (EFF94116.1); 98/97 | AbyZ | AbsH1 | AbmZ |
| *B065_RS0128170* | 117 | oxidoreductase | AbsH2, *Streptomyces* sp. LC-6-2 (ARE67859.1); 100/100 | - | AbsH2 | - |

**Table S67.** Predicted functions of ORFs in abyssomicin BGC from *Streptomyces* sp. e14 (NZ_GG753626.1).

| ORF | Size (aa) | Proposed function | Closest homolog, host (protein ID); Identity/Similarity (%) | Aby homolog | Abs homolog | Abm homolog |
|---|---|---|---|---|---|---|
| *SSTG_RS22930* | 1190 | RHS repeat protein | type IV secretion protein Rhs, *Streptomyces* sp. Amel2xE9 (WP_019984805.1); 99/99 | AbyK | - | - |
| *SSTG_RS22935* | 191 | short-chain dehydrogenase | short-chain dehydrogenase, *Rhodococcus* sp. 06-156-4C (OZD08776.1); 80/86 | - | - | - |
| *SSTG_RS22940* | 301 | alpha/beta fold hydrolase | AbsP, *Streptomyces* sp. LC-6-2 (ARE67863.1); 99/99 | - | AbsP | - |
| *SSTG_RS22945* | 209 | histidine phosphatase family protein | AbsK, *Streptomyces* sp. LC-6-2 (ARE67862.1); 97/98 | - | AbsK | - |
| *SSTG_RS22950* | 232 | TetR family transcriptional regulator | AbsC1, *Streptomyces* sp. LC-6-2 (ARE67861.1); 99/100 | - | AbsC1 | - |
| *SSTG_RS22955* | 387 | oxidoreductase | AbsH3, *Streptomyces* sp. LC-6-2 (ARE67860.1); 98/98 | - | AbsH3 | - |
| *SSTG_RS22960* | 117 | oxidoreductase | AbsH2, *Streptomyces* sp. LC-6-2 (ARE67859.1); 100/100 | AbyZ | AbsH1 | AbmZ |
| *SSTG_RS22965* | 163 | flavin reductase family protein | AbsH1, *Streptomyces* sp. LC-6-2 (ARE67858.1); 99/100 | - | AbsH2 | - |
| *SSTG_RS22970* | 351 | 3-oxoacyl-ACP synthase III family protein | 3-oxoacyl-ACP synthase III family protein, *Streptomyces* sp. Amel2xE9 (WP_019984792.1); 99/99 | AbyA1 | AbsA1 | AbmA1 |
| *SSTG_RS22975* | 77 | acyl carrier protein | acyl carrier protein, *Streptomyces* sp. Amel2xE9 (WP_019984791.1); 97/97 | AbyA3 | AbsA3 | AbmA3 |
| *SSTG_RS22980* | 251 | acyltransferase | AbsA4, *Streptomyces* sp. LC-6-2 (ARE67855.1); 100/100 | AbyA4 | AbsA4 | AbmA4 |
| *SSTG_RS22985* | 89 | alpha/beta hydrolase | AbsA5, *Streptomyces* sp. LC-6-2 (ARE67854.1); 98/98 | AbyA5 | AbsA5 | AbmA5 |
| *SSTG_RS22990* | 275 | thioesterase | AbsN, *Streptomyces* sp. LC-6-2 (ARE67865.1); 99/99 | AbyT | AbsN | AbmT |
| *SSTG_RS34725* | 87 | hypothetical protein | - | - | - | - |
| *SSTG_RS33420* | 561 | PKS I | AbsB1, *Streptomyces* sp. LC-6-2 (ARE67853.1); 96/96 | PKS I | PKS I | PKS I |
| *SSTG_RS34730* | - | PKS I | AbsB1, *Streptomyces* sp. LC-6-2 (ARE67853.1); 99/99 | PKS I | PKS I | PKS I |
| *SSTG_RS34735* | - | PKS I | AbsB1, *Streptomyces* sp. LC-6-2 (ARE67853.1); 91/92 | PKS I | PKS I | PKS I |
| *SSTG_RS34740* | - | PKS I | AbsB1, *Streptomyces* sp. LC-6-2 (ARE67853.1); 98/99 | PKS I | PKS I | PKS I |
| *SSTG_RS34745* | - | PKS I | AbsB1, *Streptomyces* sp. LC-6-2 (ARE67853.1); 98/98 | PKS I | PKS I | PKS I |
| *SSTG_RS23005* | - | PKS I | - | PKS I | PKS I | PKS I |
| *SSTG_RS23010* | - | PKS I | - | PKS I | PKS I | PKS I |
| *SSTG_RS33430* | - | PKS I | - | PKS I | PKS I | PKS I |
| *SSTG_RS23020* | - | PKS I | AbsB2, *Streptomyces* sp. LC-6-2 (ARE67852.1); 100/100 | PKS I | PKS I | PKS I |
| *SSTG_RS23025* | - | PKS I | AbsB2, *Streptomyces* sp. LC-6-2 (ARE67852.1); 96/96 | PKS I | PKS I | PKS I |
| *SSTG_RS23030* | - | PKS I | - | PKS I | PKS I | PKS I |

| | | | | | | |
|---|---|---|---|---|---|---|
| *SSTG_RS23035* | - | PKS I | - | PKS I | PKS I | PKS I |
| *SSTG_RS23040* | - | PKS I | AbsB3, *Streptomyces* sp. LC-6-2 (ARE67851.1); 99/100 | PKS I | PKS I | PKS I |
| *SSTG_RS23045* | - | PKS I | - | PKS I | PKS I | PKS I |
| *SSTG_RS23050* | 588 | methoxymalonyl-ACP biosynthesis protein FkbH | AbsA2, Streptomyces sp. LC-6-2 (ARE67850.1); 98/97 | AbyA2 | AbsA2 | AbmA2 |
| *SSTG_RS23055* | 353 | cytochrome P450 | AbsV, *Streptomyces* sp. LC-6-2 (ARE67844.1); 100/100 | AbyV | AbsV | AbmV |
| *SSTG_RS23060* | 68 | ferredoxin | AbsG1, *Streptomyces* sp. LC-6-2 (ARE67843.1); 100/100 | - | AbsG1 | AbmG |
| *SSTG_RS23065* | 387 | acyltransferase | acyltransferase, *Streptomyces* sp. Amel2xE9 (WP_019985553.1); 99/99 | - | AbsI | - |
| *SSTG_RS23070* | 403 | cytochrome P450 | cytochrome P450, *Streptomyces* sp. Amel2xE9 (WP_019985552.1); 99/99 | AbyX | AbsX | - |
| *SSTG_RS23075* | 64 | ferredoxin | AbsG2, *Streptomyces* sp. LC-6-2 (ARE67840.1); 100/100 | - | AbsG2 | - |
| *SSTG_RS23080* | 344 | aldo/keto reductase | AbsJ, *Streptomyces* sp. LC-6-2 (ARE67839.1); 100/100 | - | AbsJ | AbmJ |
| *SSTG_RS23085* | 171 | Diels-Alderase | AbsU, *Streptomyces* sp. LC-6-2 (ARE67838.1); 100/100 | AbyU | AbsU | AbmU |
| *SSTG_RS23090* | 196 | TetR/AcrR family transcriptional regulato | AbsC2, *Streptomyces* sp. LC-6-2 (ARE67837.1); 99/99 | - | AbsC2 | - |
| *SSTG_RS23095* | 476 | MFS transporter | AbsD, *Streptomyces* sp. LC-6-2 (ARE67836.1); 100/100 | AbyD | AbsD | AbmD |
| *SSTG_RS23100* | 688 | ATP-binding protein | LuxR family transcriptional regulator, *Streptomyces* sp. Amel2xE9 (WP_019985546.1); 99/99 | AbyH | - | AbmH |
| *SSTG_RS23105* | 257 | AfsR/SARP family transcriptional regulator | SARP family transcriptional regulator, *Streptomyces* sp. Amel2xE9 (WP_019985545.1); 99/99 | AbyI | - | AbmI |

**Table S68.** Predicted functions of ORFs in abyssomicin BGC from *Streptomyces fragilis* NBRC 12862 (NZ_BEVZ01000002.1).

| **ORF** | **Size (aa)** | **Proposed function** | **Closest homolog, host (protein ID); Identity/Similarity (%)** | **Aby homolog** | **Abs homolog** | **Abm homolog** |
|---|---|---|---|---|---|---|
| *Sfr03f_RS07880* | 859 | RHS repeat protein | hypothetical protein ACZ91_47855, *Streptomyces regensis* (KMS84453.1); 76/81 | AbyK | - | - |
| *Sfr03f_RS07885* | 134 | Diels-Alderase | YD repeat-containing protein, *Streptomyces regensis* (KMS84434.1); 95/95 | AbyU | AbsU | AbmU |
| *Sfr03f_RS07890* | 344 | 3-oxoacyl-ACP synthase III family protein | 3-oxoacyl-ACP synthase III family protein, *Streptomyces regalis* (WP_062712132.1); 87/93 | AbyA1 | AbsA1 | AbmA1 |
| *Sfr03f_RS07895* | 626 | HAD-IIIC family phosphatase | HAD-IIIC family phosphatase, *Streptomyces* (WP_078865191.1); 84/88 | AbyA2 | AbsA2 | AbmA2 |
| *Sfr03f_RS07900* | 75 | acyl carrier protein | acyl carrier protein, *Streptomyces* sp. NRRL WC-3725 (WP_031029037.1); 92/95 | AbyA3 | AbsA3 | AbmA3 |
| *Sfr03f_RS07905* | 258 | acyltransferase | Acyltransferase, *Streptomyces* (WP_051818896.1); 84/90 | AbyA4 | AbsA4 | AbmA4 |
| *Sfr03f_RS07910* | 360 | alpha/beta hydrolase | alpha/beta hydrolase, *Streptomyces* (WP_030991268.1); 88/92 | AbyA5 | AbsA5 | AbmA5 |
| *Sfr03f_RS07915* | 166 | flavin reductase | flavin oxidoreductase, *Streptomyces regensis* (KMS84438.1); 84/87 | AbyZ | AbsH1 | AbmZ |
| *Sfr03f_RS07920* | 229 | TetR/AcrR family transcriptional regulator | TetR/AcrR family transcriptional regulator, *Streptomyces* (WP_031100136.1); 90/95 | AbyC | - | AbmC |
| *Sfr03f_RS07925* | 461 | DHA2 family efflux MFS transporter permease subunit | DHA2 family efflux MFS transporter permease subunit, *Streptomyces* sp. NRRL WC-3744 (WP_030991264.1); 90/93 | AbyD | AbsD | AbmD |
| *Sfr03f_RS07930* | 345 | LLM class flavin-dependent oxidoreductase | LLM class flavin-dependent oxidoreductase, *Streptomyces regalis* (WP_062712142.1); 87/91 | AbyE | AbsE | AbmE1 |
| *Sfr03f_RS07935* | 546 | ABC transporter substrate-binding protein | ABC transporter substrate-binding protein, *Streptomyces* sp. NRRL WC-3744 (WP_030991261.1); 80/86 | AbyF1 | AbsF1 | AbmF1 |
| *Sfr03f_RS07940* | 315 | ABC transporter permease | ABC transporter permease, *Streptomyces regensis* (KMS84443.1); 84/90 | AbyF2 | AbsF2 | AbmF2 |
| *Sfr03f_RS07945* | 297 | ABC transporter permease | ABC transporter permease, *Streptomyces regalis* (WP_062712151.1); 80/86 | AbyF3 | AbsF3 | AbmF3 |
| *Sfr03f_RS07950* | 543 | ABC transporter ATP-binding protein | ABC transporter ATP-binding protein, *Streptomyces* (WP_031029029.1); 84/87 | AbyF4 | AbsF4 | AbmF4 |
| *Sfr03f_RS07955* | 434 | acyltransferase | Acyltransferase, *Streptomyces regalis* (WP_062712154.1); 79/85 | - | AbsI | - |
| *Sfr03f_RS07960* | 406 | cytochrome P450 | cytochrome P450, *Streptomyces* (WP_030654493.1); 92/95 | AbyV/AbyX | AbsV/AbsX | AbmV |
| *Sfr03f_RS07965* | 64 | ferredoxin | Ferredoxin, *Streptomyces* (WP_030654496.1); 92/95 | - | AbsG1 | AbmG |
| *Sfr03f_RS07970* | 307 | alpha/beta hydrolase | alpha/beta hydrolase, *Streptomyces* (WP_030654499.1); 90/92 | - | - | - |
| *Sfr03f_RS07975* | 6216 | PKS I | type I polyketide synthase, *Streptomyces* sp. 2131.1 (WP_093709985.1); 64/72 | AbyB1 | AbsB1 | AbmB1 |
| *Sfr03f_RS07980* | 3846 | PKS I | type I polyketide synthase, *Streptomyces* sp. 2131.1 (WP_093709984.1); 68/76 | AbyB2 | AbsB2 | AbmB2 |
| *Sfr03f_RS07985* | 960 | PKS I | type I polyketide synthase, *Streptomyces* sp. 2131.1 (WP_093709983.1); 73/80 | AbyB3 | AbsB3 | AbmB3 |
| *Sfr03f_RS07990* | 77 | PKS I | type I polyketide synthase, *Streptomyces* sp. NRRL WC-3725 (WP_043195775.1); 89/97 | | | |
| *Sfr03f_RS07995* | 393 | cytochrome P450 | cytochrome P450, *Streptomyces* (WP_031101897.1); 85/90 | AbyX/AbyV | AbsV/AbsX | AbmV |

**Table S69.** Predicted functions of ORFs in potential abyssomicin BGC from *Streptomyces incarnatus* NRRL 8089 (CP011497).

| ORF | Size (aa) | Proposed function | Closest homolog, host (protein ID); Identity/Similarity (%) | Aby homolog | Abs homolog | Abm homolog |
|---|---|---|---|---|---|---|
| *ABB07_01775* | 339 | luciferase | LLM class flavin-dependent oxidoreductase, *Streptomyces* sp. Amel2xE9 (WP_106962144.1); 91/94 | - | - | AbmE2 |
| *ABB07_01780* | 399 | cytochrome P450 | cytochrome P450, *Streptomyces* sp. NRRL WC-3742 (WP_031075102.1); 72/81 | AbyV | AbsV | AbmV |
| *ABB07_01785* | 71 | hypothetical protein | ferredoxin, *Streptomyces* sp. KE1 (WP_047140955.1); 55/71 | AbyW | - | - |
| *ABB07_01790* | 347 | luciferase | LLM class flavin-dependent oxidoreductase, *Frankia alni* (WP_011601492.1); 57/69 | AbyE | AbsE | AbmE1 |
| *ABB07_01795* | 497 | MFS transporter | MFS transporter, *Streptomyces* sp. Amel2xE9 (WP_019985547.1); 64/76 | AbyD | AbsD | AbmD |
| *ABB07_01800* | 201 | TetR family transcriptional regulator | TetR/AcrR family transcriptional regulator, *Microbispora rosea* (WP_076442332.1); 775/82 | - | AbsC2 | - |
| *ABB07_01805* | 447 | nitrilotriacetate monooxygenase | LLM class flavin-dependent oxidoreductase, *Frankia symbiont of Coriaria ruscifolia* (WP_131785890.1); 78/87 | - | - | - |
| *ABB07_01810* | 514 | hypothetical protein | hypothetical protein, *Frankia* sp. BMG5.30 (WP_047223156.1); 55/72 | AbyF1 | AbsF1 | AbmF1 |
| *ABB07_01815* | 270 | hypothetical protein | ABC transporter ATP-binding protein, *Frankia* sp. BMG5.30 (WP_083731087.1); 62/74 | AbyF4 | AbsF4 | AbmF4 |
| *ABB07_01820* | 269 | hypothetical protein | ATP-binding cassette domain-containing protein, *Frankia symbiont of Datisca glomerata* (WP_013873338.1); 65/75 | AbyF4 | AbsF4 | AbmF4 |
| *ABB07_01825* | 289 | hypothetical protein | ABC transporter permease subunit, *Frankia symbiont of Coriaria nepalensis* (WP_131772430.1); 57/71 | AbyF3 | AbsF3 | AbmF3 |
| *ABB07_01830* | 342 | ABC transporter permease | ABC-type transporter, integral membrane subunit, *Frankia symbiont of Datisca glomerata* (AEH09395.1); 62/75 | AbyF2 | AbsF2 | AbmF2 |
| *ABB07_01835* | 332 | aldo/keto reductase | aldo/keto reductase, *Streptacidiphilus* sp. DSM 106435 (WP_111492774.1); 76/84 | - | AbsJ | AbmJ |
| *ABB07_01840* | 1069 | hypothetical protein | acyltransferase domain-containing protein, *Streptacidiphilus* sp. DSM 106435 (WP_111490411.1); 60/68 | AbyB3 | AbsB3 | AbmB3 |
| *ABB07_01855* | 343 | 3-oxoacyl-ACP synthase | 3-oxoacyl-ACP synthase III family protein, *Streptomyces* sp. NRRL WC-3742 (WP_031075097.1); 71/81 | AbyA1 | AbsA1 | AbmA1 |
| *ABB07_01860* | 135 | Diels-Alderase | hypothetical protein, Streptomyces sp. CB03911 (WP_073928710.1); 71/87 | AbyU | AbsU | AbmU |
| *ABB07_01865* | 6174 | PKS I | type I polyketide synthase, *Streptomyces fragilis* (WP_108952947.1); 55/63 | AbyB1 | AbsB1 | AbmB1 |
| *ABB07_01880* | 365 | hydrolase superfamily dihydrolipoamide acyltransferase-like protein | alpha/beta hydrolase, *Streptomyces fragilis* (WP_108952935.1); 62/72 | AbyA5 | AbsA5 | AbmA5 |
| *ABB07_01885* | 69 | hypothetical protein | hypothetical protein, *Streptomyces hokutonensis* (WP_019071716.1); 51/61 | - | - | |
| *ABB07_01890* | 256 | hypothetical protein | activator protein, *Actinocrispum wychmicini* (WP_132114020.1); 62/75 | AbyI | - | AbmI |
| *ABB07_01895* | 260 | hypothetical protein | thioesterase, *Streptomyces paucisporeus* (WP_073501301.1); 67/75 | AbyT | AbsN | AbmT |
| *ABB07_01900* | 619 | methoxymalonyl-ACP biosynthesis protein FkbH | HAD-IIIC family phosphatase, *Streptomyces formicae* (WP_098241224.1); 64/73 | AbyA2 | AbsA2 | AbmA2 |
| *ABB07_01905* | 88 | hypothetical protein | acyl carrier protein, *Streptomyces* sp. CNQ329 (WP_027774411.1); 58/75 | AbyA3 | AbsA3 | AbmA3 |
| *ABB07_01910* | 172 | hypothetical protein | flavin reductase, *Nocardiopsis potens* (WP_017592460.1); 60/72 | AbyZ | AbsH1 | AbmZ |

**Table S70.** Predicted functions of ORFs in abyssomicin BGC from *Streptomyces* sp. NRRL F-6491 (NZ_LGEE01000251 and NZ_LGEE01000286).

| ORF | Size (aa) | Proposed function | Closest homolog, host (protein ID); Identity/Similarity (%) | Aby homolog | Abs homolog | Abm homolog |
|---|---|---|---|---|---|---|
| *ADL06_RS28345* | 175 | hypothetical protein | DsbA family oxidoreductase, *Streptomyces* sp. CNH287 (WP_027750659.1); 82/88 | - | - | - |
| *ADL06_RS28350* | 94 | hypothetical protein | - | - | - | - |
| *ADL06_RS28355* | 63 | hypothetical protein | MFS transporter, *Streptomyces* sp. Or20 (WP_097967670.1); 71/74 | AbyD | AbsD | AbmD |
| *ADL06_RS28360* | 212 | 4'-phosphopantetheinyl transferase superfamily protein | 4'-phosphopantetheinyl transferase superfamily protein, *Streptomyces* sp. S1 (WP_121827868.1); 93/95 | - | - | - |
| *ADL06_RS28365* | 349 | 3-oxoacyl-ACP synthase III family protein | AbsA1, *Streptomyces* sp. LC-6-2 (ARE67857.1); 87/91 | AbyA1 | AbsA1 | AbmA1 |
| *ADL06_RS28370* | 81 | acyl carrier protein | AbsA3, *Streptomyces* sp. LC-6-2 (ARE67856.1); 75/84 | AbyA3 | AbsA3 | AbmA3 |
| *ADL06_RS28375* | 251 | acyltransferase | AbsA4, *Streptomyces* sp. LC-6-2 (ARE67855.1); 90/94 | AbyA4 | AbsA4 | AbmA4 |
| *ADL06_RS28380* | 370 | alpha/beta hydrolase | AbsA5, *Streptomyces* sp. LC-6-2 (ARE67854.1); 83/86 | AbyA5 | AbsA5 | AbmA5 |
| *ADL06_RS28385* | 274 | thioesterase | thioesterase, *Streptomyces* sp. E14 (WP_009191660.1); 80/83 | AbyT | AbsN | AbmT |
| *ADL06_RS28390* | - | PKS I | AbsB1, *Streptomyces* sp. LC-6-2 (ARE67853.1); 80/84 | AbyB1 | AbsB1 | AbmB1 |
| //////////////////////////////////////////////////////////////////////////////////////////////////// | | | | | | |
| *ADL06_RS34415* | - | PKS I | AbsB1, *Streptomyces* sp. LC-6-2 (ARE67853.1); 74/78 | AbyB1 | AbsB1 | AbmB1 |
| *ADL06_RS36865* | - | PKS I | - | PKS I | PKS I | PKS I |
| *ADL06_RS34420* | - | PKS I | SDR family NAD(P)-dependent oxidoreductase, partial, *Microbispora triticiradicis* (WP_117408275.1); 64/71 | PKS I | PKS I | PKS I |
| *ADL06_RS34425* | - | PKS I | AbsB3, *Streptomyces* sp. LC-6-2 (ARE67851.1); 78/84 | AbyB3 | AbsB3 | AbmB3 |
| *ADL06_RS34430* | 652 | HAD-IIIC family phosphatase | AbsA2, *Streptomyces* sp. LC-6-2 (ARE67850.1); 81/86 | AbyA2 | AbsA2 | AbmA2 |
| *ADL06_RS34435* | 320 | LLM class flavin-dependent oxidoreductase | AbsE, *Streptomyces* sp. LC-6-2 (ARE67849.1); 90/94 | AbyE | AbsE | AbmE1 |
| *ADL06_RS34440* | 551 | ABC transporter substrate-binding protein | AbsF1, *Streptomyces* sp. LC-6-2 (ARE67848.1); 84/90 | AbyF1 | AbsF1 | AbmF1 |
| *ADL06_RS34445* | 334 | ABC transporter permease | ABC transporter permease, *Streptomyces* sp. Amel2xE9 (WP_019985558.1); 82/90 | AbyF2 | AbsF2 | AbmF2 |
| *ADL06_RS34450* | 269 | ABC transporter permease | AbsF3, *Streptomyces* sp. LC-6-2 (ARE67846.1); 90/92 | AbyF3 | AbsF3 | AbmF3 |
| *ADL06_RS34455* | 551 | ABC transporter ATP-binding protein | AbsF4, *Streptomyces* sp. LC-6-2 (ARE67845.1); 86/89 | AbyF4 | AbsF4 | AbmF4 |
| *ADL06_RS34460* | 395 | cytochrome P450 | cytochrome P450, *Streptomyces* sp. Amel2xE9 (WP_027758724.1); 93/96 | AbyV | AbsV | AbmV |
| *ADL06_RS34465* | 68 | ferredoxin | AbsG1, *Streptomyces* sp. LC-6-2 (ARE67843.1); 84/94 | - | AbsG1 | AbmG |
| *ADL06_RS34470* | 395 | acyltransferase | AbsI, *Streptomyces* sp. LC-6-2 (ARE67842.1); 84/88 | - | AbsI | - |

| | | | | | | |
|---|---|---|---|---|---|---|
| *ADL06_RS34475* | 403 | cytochrome P450 | AbsX, *Streptomyces* sp. LC-6-2 (ARE67841.1); 91/94 | AbyX | AbsX | - |
| *ADL06_RS34480* | 69 | ferredoxin | AbsG2, *Streptomyces* sp. LC-6-2 (ARE67840.1); 89/95 | - | AbsG2 | - |
| *ADL06_RS34485* | 346 | aldo/keto reductase | aldo/keto reductase, *Streptomyces* sp. Amel2xE9 (WP_020657195.1); 84/88 | - | AbsJ | AbmJ |
| *ADL06_RS34490* | 171 | Diels-Alderase | AbsU, *Streptomyces* sp. LC-6-2 (ARE67838.1); 92/97 | AbyU | AbsU | AbmU |
| *ADL06_RS34495* | 199 | TetR/AcrR family transcriptional regulator | AbsC2, *Streptomyces* sp. LC-6-2 (ARE67837.1); 90/94 | - | AbsC2 | - |
| *ADL06_RS34500* | 476 | MFS transporter | AbsD, *Streptomyces* sp. LC-6-2 (ARE67836.1); 88/92 | AbyD | AbsD | AbmD |

**Table S71.** Predicted functions of ORFs in potential BGC from *Streptomyces olivaceus* KLBMP 5084 (NZ_CP016795.1).

| ORF | Size (aa) | Proposed function | Closest homolog, host (protein ID); Identity/Similarity (%) | Aby homolog | Abs homolog | Abm homolog |
|---|---|---|---|---|---|---|
| *BC342_RS34475* | 260 | thioesterase | thioesterase, *Streptomyces qinglanensis* (WP_069991949.1); 81/87 | - | - | - |
| *BC342_RS34480* | 367 | serine hydrolase | serine hydrolase, *Streptomyces longwoodensis* (WP_067241675.1); 83/90 | - | - | - |
| *BC342_RS34485* | 382 | homoserine O-acetyltransferase | homoserine O-acetyltransferase, *Streptomyces* sp. E14 (EFF89200.1); 86/90 | - | - | - |
| *BC342_RS34490* | 453 | bifunctional o-acetylhomoserine/o-acetylserine sulfhydrylase | bifunctional o-acetylhomoserine/o-acetylserine sulfhydrylase, *Streptomyces* sp. XY006 (WP_094053066.1); 91/96 | - | - | - |
| *BC342_RS34495* | 201 | dihydrofolate reductase | dihydrofolate reductase, *Streptomyces* sp. CB01635 (WP_100599268.1); 87/95 | - | - | - |
| *BC342_RS34500* | 269 | response regulator transcription factor | response regulator transcription factor, *Streptomyces* sp. NRRL F-5639 (WP_051705713.1); 53/65 | - | - | - |
| *BC342_RS34505* | 290 | aldo/keto reductase family oxidoreductase | aldo/keto reductase family oxidoreductase, *Streptomyces* sp. FXJ7.023 (WP_037772516.1); 99/99 | - | - | - |
| *BC342_RS34510* | 148 | helix-turn-helix transcriptional regulator | transcriptional regulator, *Streptomyces* sp. FXJ7.023 (WP_037772514.1); 99/100 | - | - | - |
| *BC342_RS34515* | 94 | helix-turn-helix transcriptional regulator | transcriptional regulator, *Brevibacterium aurantiacum* (WP_096147035.1); 72/90 | - | - | - |
| *BC342_RS34520* | 363 | NADH:flavin oxidoreductase/NADH oxidase | NADH:flavin oxidoreductase/NADH oxidase, *Corynebacterium sputi* (WP_027019135.1); 69/80 | - | - | - |
| *BC342_RS34525* | 745 | transcriptional regulator | XRE family transcriptional regulator, *Streptomyces viridosporus* (WP_081238662.1); 83/88 | - | - | - |
| *BC342_RS34530* | 185 | hypothetical protein | hypothetical protein, *Streptomyces* sp. Root369 (WP_057616167.1); 91/96 | - | - | - |
| *BC342_RS34535* | 191 | hypothetical protein | hypothetical protein, *Streptomyces* sp. FXJ7.023 (WP_037772510.1); 99/99 | - | - | - |
| *BC342_RS34540* | 276 | PKS I | methyltransferase domain-containing protein, *Streptomyces* sp. FXJ7.023 (WP_037772509.1); 99/99 | - | - | - |
| *BC342_RS34545* | 1364 | PKS I | acyltransferase domain-containing protein, *Streptomyces* sp. E5N91 SAI-083 (WP_123627584.1); 95/96 | - | - | - |
| *BC342_RS34550* | 308 | PKS I | acyl transferase, partial, *Streptomyces varsoviensis* (KOG86991.1); 90/91 | - | - | - |
| *BC342_RS34555* | 173 | pyridoxamine 5'-phosphate oxidase family protein | pyridoxamine 5'-phosphate oxidase family protein, *Streptomyces* sp. FXJ7.023 (WP_037772507.1); 99/100 | - | - | - |
| *BC342_RS34560* | 2799 | PKS I | type I polyketide synthase, *Streptomyces cattleya* (WP_014140914.1); 87/90 | - | - | - |
| *BC342_RS34565* | 6128 | PKS I | SDR family NAD(P)-dependent oxidoreductase, *Streptomyces* sp. E5N91 SAI-083 (WP_123627588.1); 96/97 | - | - | - |
| *BC342_RS34570* | 140 | Diels-Alderase | hypothetical protein, *Streptomyces* sp. E5N91 SAI-083 (WP_123627591.1); 98/99 | AbyU | AbsU | AbmU |
| *BC342_RS34575* | 133 | nuclear transport factor 2 family protein | nuclear transport factor 2 family protein, *Streptomyces* sp. E5N91 SAI-083 (WP_123627590.1); 95/97 | - | - | - |
| *BC342_RS34580* | 416 | hypothetical protein | cytochrome P450, *Streptomyces* sp. E5N91 SAI-083 (WP_123627589.1); 97/98 | - | - | - |
| *BC342_RS34585* | 280 | NAD(P)-dependent oxidoreductase | NAD(P)-dependent oxidoreductase, *Streptomyces* sp. FXJ7.023 (WP_037772598.1); 99/100 | - | - | - |

| | | | | | | |
|---|---|---|---|---|---|---|
| *BC342_RS34590* | 157 | nuclear transport factor 2 family protein | nuclear transport factor 2 family protein, *Streptomyces* sp. E5N91 SAI-083 (WP_123627593.1); 98/99 | - | - | - |
| *BC342_RS34595* | 273 | thioesterase | thioesterase, *Streptomyces cattleya* (WP_014140907.1); 85/90 | - | - | - |
| *BC342_RS34600* | 408 | cytochrome P450 | cytochrome P450, *Streptomyces* sp. E5N91 SAI-083 (WP_123627594.1); 98/98 | - | - | - |
| *BC342_RS36425* | 99 | hypothetical protein | hypothetical protein EDC84_6803, *Streptomyces* sp. E5N91 SAI-083 (ROO97925.1); 94/94 | - | - | - |
| *BC342_RS34605* | 393 | LLM class flavin-dependent oxidoreductase | LLM class flavin-dependent oxidoreductase, *Streptomyces* sp. E5N91 SAI-083 (WP_123627595.1); 98/99 | - | - | - |
| *BC342_RS34610* | 277 | SDR family oxidoreductase | SDR family oxidoreductase, *Streptomyces* sp. E5N91 SAI-083 (WP_123627596.1); 98/98 | - | - | - |

**Table S72.** Predicted functions of ORFs in potential abyssomicin BGC from *Streptomyces regalis* NRRL 3151 (NZ_LLZG01000385).

| ORF | Size (aa) | Proposed function | Closest homolog, host (protein ID); Identity/Similarity (%) | Aby homolog | Abs homolog | Abm homolog |
|---|---|---|---|---|---|---|
| *ADL12_RS39130* | 257 | AfsR/SARP family transcriptional regulator | activator protein, *Streptomyces regensis* (KMS84432.1); 89/94 | AbyI/AbyR | - | AbmI |
| *ADL12_RS39135* | 257 | thioesterase | thioesterase, *Streptomyces* sp. NRRL WC-3744 (WP_063743390.1); 67/75 | AbyT | AbsN | AbmT |
| *ADL12_RS39140* | 909 | LuxR family transcriptional regulator | helix-turn-helix transcriptional regulator, *Streptomyces* sp. NRRL WC-3744 (WP_106978307.1); 74/82 | AbyH | - | AbmH |
| *ADL12_RS39145* | 860 | RHS repeat protein | RHS repeat protein, *Streptomyces fragilis* (WP_108953524.1); 72/79 | AbyK | - | - |
| *ADL12_RS39150* | 134 | Diels-Alderase | hypothetical protein, *Streptomyces fragilis* (WP_108952931.1); 91/94 | AbyU | AbsU | AbmU |
| *ADL12_RS39155* | 344 | 3-oxoacyl-ACP synthase III family protein | 3-oxoacyl-ACP synthase, *Streptomyces regensis* (KMS84435.1); 89/93 | AbyA1 | AbsA1 | AbmA1 |
| *ADL12_RS39160* | 589 | HAD-IIIC family phosphatase | HAD-IIIC family phosphatase, *Streptomyces* sp. NRRL WC-3744 (WP_078630008.1); 83/88 | AbyA2 | AbsA2 | AbmA2 |
| *ADL12_RS39165* | 75 | hypothetical protein | acyl carrier protein, *Streptomyces* sp. NRRL WC-3725 (WP_031029037.1); 92/94 | AbyA3 | AbsA3 | AbmA3 |
| *ADL12_RS39170* | 255 | acyltransferase | acyltransferase, *Streptomyces* sp. NRRL WC-3744 (WP_051816343.1); 84/91 | AbyA4 | AbsA4 | AbmA4 |
| *ADL12_RS39175* | 360 | alpha/beta hydrolase | hydrolase superfamily dihydrolipoamide acyltransferase-like protein, *Streptomyces regensis* (KMS84437.1); 87/89 | AbyA5 | AbsA5 | AbmA5 |
| *ADL12_RS39180* | 166 | flavin reductase | flavin oxidoreductase, *Streptomyces regensis* (KMS84438.1); 84/90 | AbyZ | AbsH1 | AbmZ |
| *ADL12_RS39185* | 227 | TetR/AcrR family transcriptional regulator | TetR family transcriptional regulator, *Streptomyces* sp. NRRL WC-3723 (KOV74725.1); 89/95 | AbyC | - | AbmC |
| *ADL12_RS39190* | 483 | DHA2 family efflux MFS transporter permease subunit | DHA2 family efflux MFS transporter permease subunit, *Streptomyces* sp. NRRL WC-3744 (WP_030991264.1); 89/93 | AbyD | AbsD | AbmD |
| *ADL12_RS39195* | 345 | LLM class flavin-dependent oxidoreductase | FMN-linked alkanal monooxygenase, *Streptomyces regensis* (KMS84441.1); 89/94 | AbyE | AbsE | AbmE1 |
| *ADL12_RS39200* | 543 | ABC transporter substrate-binding protein | ABC transporter substrate-binding protein, *Streptomyces fragilis* (WP_108952939.1); 78/86 | AbyF1 | AbsF1 | AbmF1 |
| *ADL12_RS39205* | 316 | ABC transporter permease | ABC transporter permease, *Streptomyces regensis* (KMS84443.1); 87/92 | AbyF2 | AbsF2 | AbmF2 |
| *ADL12_RS39210* | 291 | ABC transporter permease | ABC transporter permease, *Streptomyces fragilis* (WP_108952941.1); 81/88 | AbyF3 | AbsF3 | AbmF3 |
| *ADL12_RS39215* | 548 | ABC transporter ATP-binding protein | ABC transporter ATP-binding protein, *Streptomyces fragilis* (WP_108952942.1); 82/87 | AbyF4 | AbsF4 | AbmF4 |
| *ADL12_RS39220* | 389 | acyltransferase | acyltransferase, *Streptomyces* sp. NRRL WC-3725 (WP_031029026.1); 87/90 | - | AbsI | - |
| *ADL12_RS39225* | 406 | cytochrome P450 | cytochrome P450, *Streptomyces regensis* (KMS84447.1); 90/93 | AbyV/AbyX | AbsV/AbsX | AbmV |
| *ADL12_RS39230* | 64 | ferredoxin | ferredoxin, *Streptomyces regensis* (KMS84448.1); 92/95 | - | AbsG2/AbsG1 | AbmG |
| *ADL12_RS39235* | 305 | alpha/beta hydrolase | alpha/beta hydrolase, *Streptomyces regensis* (KMS84449.1); 89/91 | - | - | - |

| | | | | | | |
|---|---|---|---|---|---|---|
| *ADL12_RS39240* | - | PKS I | type I polyketide synthase, *Streptomyces* sp. 2131.1 (WP_093709985.1); 68/76 | PKS I | PKS I | PKS I |

**Table S73.** Predicted functions of ORFs in potential abyssomicin BGC from *Streptomyces regensis* NRRL B-11479 (LFVR01000395.1).

| ORF | Size (aa) | Proposed function | Closest homolog, host (protein ID); Identity/Similarity (%) | Aby homolog | Abs homolog | Abm homolog |
|---|---|---|---|---|---|---|
| *ACZ91_47835* | 69 | hypothetical protein | hypothetical protein, *Streptomyces* sp. NRRL WC-3744 (WP_030991280.1); 100/100 | - | - | - |
| *ACZ91_47840* | 257 | activator protein | activator protein, *Streptomyces* sp. NRRL WC-3725 (WP_031029046.1); 100/100 | AbyI | - | AbmI |
| *ACZ91_47850* | 914 | hypothetical protein | helix-turn-helix transcriptional regulator, *Streptomyces* sp. NRRL WC-3725 (WP_107054736.1); 99/99 | AbyH | - | AbmH |
| *ACZ91_47855* | 882 | hypothetical protein | RHS repeat protein, *Streptomyces fragilis* (WP_108953524.1); 76/81 | AbyK | - | - |
| *ACZ91_47860* | 134 | Diels-Alderase | hypothetical protein, *Streptomyces fragilis* (WP_108952931.1); 95/95 | AbyU | AbsU | AbmU |
| *ACZ91_47865* | 344 | 3-oxoacyl-ACP synthase | 3-oxoacyl-ACP synthase, *Streptomyces antibioticus* (KOG62350.1); 100/100 | AbyA1 | AbsA1 | AbmA1 |
| *ACZ91_47870* | 630 | methoxymalonyl-ACP biosynthesis protein FkbH | HAD-IIIC family phosphatase, *Streptomyces* sp. NRRL WC-3725 (WP_078918059.1); 99/99 | AbyA2 | AbsA2 | AbmA2 |
| *ACZ91_47875* | 75 | acyl carrier protein | acyl carrier protein, *Streptomyces antibioticus* (KOG62349.1); 100/100 | AbyA3 | AbsA3 | AbmA3 |
| *ACZ91_47880* | 246 | acyltransferase | acyltransferase, *Streptomyces* sp. NRRL WC-3744 (WP_051816343.1); 99/99 | AbyA4 | AbsA4 | AbmA4 |
| *ACZ91_47885* | 360 | hydrolase superfamily dihydrolipoamide acyltransferase-like protein | hydrolase superfamily dihydrolipoamide acyltransferase-like protein, *Streptomyces antibioticus* (KOG62348.1); 100/100 | AbyA5 | AbsA5 | AbmA5 |
| *ACZ91_47890* | 172 | flavin oxidoreductase | flavin oxidoreductase, *Streptomyces* sp. NRRL WC-3723 (KOV74748.1); 99/100 | AbyZ | AbsH1 | AbmZ |
| *ACZ91_47895* | 227 | TetR family transcriptional regulator | TetR/AcrR family transcriptional regulator, *Streptomyces* sp. NRRL WC-3744 (WP_030991266.1); 99/100 | AbyC | - | AbmC |
| *ACZ91_47900* | 483 | EmrB/QacA family drug resistance transporter | EmrB/QacA family drug resistance transporter, *Streptomyces antibioticus* (KOG62345.1); 100/100 | AbyD | AbsD | AbmD |
| *ACZ91_47905* | 345 | FMN-linked alkanal monooxygenase | FMN-linked alkanal monooxygenase, *Streptomyces antibioticus* (KOG62344.1); 100/100 | AbyE | AbsE | AbmE1 |
| *ACZ91_47910* | 544 | ABC transporter substrate-binding protein | ABC transporter substrate-binding protein, *Streptomyces antibioticus* (KOG62343.1); 100/100 | AbyF1 | AbsF1 | AbmF1 |
| *ACZ91_47915* | 315 | ABC transporter permease | ABC transporter permease, *Streptomyces* sp. NRRL WC-3744 (WP_030991259.1); 99/100 | AbyF2 | AbsF2 | AbmF2 |
| *ACZ91_47920* | 291 | peptide ABC transporter permease | peptide ABC transporter permease, *Streptomyces antibioticus* (KOG62341.1); 100/100 | AbyF3 | AbsF3 | AbmF3 |
| *ACZ91_47925* | 544 | ABC transporter ATP-binding protein | ABC transporter ATP-binding protein, *Streptomyces antibioticus* (KOG62340.1); 100/100 | AbyF4 | AbsF4 | AbmF4 |
| *ACZ91_47930* | 388 | hypothetical protein | acyltransferase, *Streptomyces antibioticus* (WP_053212164.1); 100/100 | - | AbsI | - |
| *ACZ91_47935* | 406 | cytochrome P450 | cytochrome P450, *Streptomyces antibioticus* (KOG62338.1); 100/100 | AbyV/AbyX | AbsV/AbsX | AbmV |
| *ACZ91_47940* | 64 | ferredoxin | ferredoxin, *Streptomyces antibioticus* (KOG62337.1); 100/100 | - | AbsG2/AbsG1 | AbmG |

| | | | | | | |
|---|---|---|---|---|---|---|
| *ACZ91_47945* | 294 | alpha/beta hydrolase | alpha/beta hydrolase, *Streptomyces antibioticus* (KOG62336.1); 100/100 | - | - | - |

**Table S74.** Predicted functions of ORFs in potential abyssomicin BGC from *Saccharothrix syringae* NRRL B-16468 (NZ_JNYO01000044.1).

| **ORF** | **Size (aa)** | **Proposed function** | **Closest homolog, host (protein ID); Identity/Similarity (%)** | **Aby homolog** | **Abs homolog** | **Abm homolog** |
|---|---|---|---|---|---|---|
| *OQ01_RS37195* | 262 | hypothetical protein | hypothetical protein, *Actinoplanes regularis* (WP_089299276.1); 32/46 | - | - | - |
| *OQ01_RS48955* | 223 | TetR/AcrR family transcriptional regulator | TetR/AcrR family transcriptional regulator, *Nocardia* sp. NRRL S-836 (WP_053731530.1); 82/88 | - | AbsC1 | - |
| *OQ01_RS37205* | 295 | alpha/beta hydrolase | alpha/beta hydrolase, *Nocardia* sp. NRRL S-836 (WP_053731531.1); 65/83 | - | - | - |
| *OQ01_RS37210* | 251 | SDR family oxidoreductase | 3-oxoacyl-ACP reductase, *Frankia coriariae* (KLL10339.1); 61/73 | - | AbsM | - |
| *OQ01_RS37215* | 200 | TetR/AcrR family transcriptional regulator | TetR/AcrR family transcriptional regulator, *Microbispora* sp. GKU 823 (WP_079317081.1); 62/73 | - | AbsC2 | - |
| *OQ01_RS37220* | 470 | MFS transporter | MFS transporter, *Microbispora* sp. GKU 823 (WP_079317079.1); 65/80 | AbyD | AbsD | AbmD |
| *OQ01_RS37225* | 438 | LLM class flavin-dependent oxidoreductase | Nitrilotriacetate monooxygenase, *Frankia alni* ACN14a (CAJ58912.1); 74/83 | - | - | - |
| *OQ01_RS37230* | 341 | LLM class flavin-dependent oxidoreductase | LLM class flavin-dependent oxidoreductase, *Frankia alni* (WP_011601492.1); 74/82 | AbyE | AbsE | AbmE1 |
| *OQ01_RS37235* | 551 | ABC transporter substrate-binding protein | ABC-type transporter, periplasmic subunit, *Frankia symbiont of Datisca glomerata* (AEH09361.1); 66/80 | AbyF1 | AbsF1 | AbmF1 |
| *OQ01_RS37240* | 305 | ABC transporter permease | ABC transporter permease, *Frankia symbiont of Coriaria ruscifolia* (WP_131785888.1); 72/83 | AbyF2 | AbsF2 | AbmF2 |
| *OQ01_RS37245* | 278 | ABC transporter permease | ABC transporter permease subunit, *Frankia symbiont of Coriaria ruscifolia* (WP_131785887.1); 74/86 | AbyF3 | AbsF3 | AbmF3 |
| *OQ01_RS37250* | 516 | ABC transporter ATP-binding protein | ABC transporter ATP-binding protein, *Frankia* sp. BMG5.30 (WP_076844589.1); 67/77 | AbyF4 | AbsF4 | AbmF4 |
| *OQ01_RS37255* | 242 | hypothetical protein | hypothetical protein, *Saccharothrix australiensis* (WP_121006568.1); 61/70 | - | - | - |
| *OQ01_RS37260* | 168 | flavin reductase | flavin reductase, *Actinomadura* sp. NEAU-Ht49 (WP_122194805.1); 62/73 | AbyZ | AbsH1 | AbmZ |
| *OQ01_RS37265* | 79 | acyl carrier protein | acyl carrier protein, *Streptomyces malaysiense* (WP_071387400.1); 53/76 | AbyA3 | AbsA3 | AbmA3 |
| *OQ01_RS37270* | 618 | HAD-IIIC family phosphatase | HAD-IIIC family phosphatase, *Kutzneria buriramensis* (WP_116181645.1); 66/76 | AbyA2 | AbsA2 | AbmA2 |
| *OQ01_RS37275* | 253 | hypothetical protein | activator protein, *Kutzneria buriramensis* (WP_116181646.1); 68/84 | AbyI | - | AbmI |
| *OQ01_RS37280* | 913 | hypothetical protein | LuxR family transcriptional regulator, *Rhodococcus yunnanensis* (WP_072806082.1); 40/57 | AbyH | - | AbmH |
| *OQ01_RS37285* | 460 | propionyl-CoA carboxylase subunit beta | propionyl-CoA carboxylase subunit beta, *Streptomyces cellostaticus* (WP_079058127.1); 70/78 | - | - | - |
| *OQ01_RS37290* | 330 | aldo/keto reductase | aldo/keto reductase, *Frankia coriariae* (WP_047223142.1); 75/84 | - | AbsJ | AbmJ |

| | | | | | | |
|---|---|---|---|---|---|---|
| *OQ01_RS37295* | 125 | Diels-Alderase | hypothetical protein FrCorBMG51_12000, *Frankia coriariae* (KLL11361.1); 80/85 | AbyU | AbsU | AbmU |
| *OQ01_RS37300* | 1323 | PKS I | pimaricinolide synthase PimS1, *Kutzneria buriramensis* (REH39151.1); 67/74 | AbyB3 | AbsB3 | AbmB3 |
| *OQ01_RS37305* | 3369 | PKS I | SDR family NAD(P)-dependent oxidoreductase, *Kutzneria buriramensis* (WP_116178516.1); 59/69 | AbyB2 | AbsB2 | AbmB2 |
| *OQ01_RS47295* | 5489 | PKS I | type I polyketide synthase, *Streptomyces fragilis* (WP_108952947.1); 55/64 | AbyB1 | AbsB1 | AbmB1 |
| *OQ01_RS50975* | 256 | PKS I | type I polyketide synthase, partial, *Streptomyces* sp. 4R-3d (WP_135068668.1); 72/81 | | | |
| *OQ01_RS37315* | 339 | alpha/beta hydrolase | alpha/beta hydrolase, *Kutzneria buriramensis* (WP_116181688.1); 66/74 | AbyA5 | AbsA5 | AbmA5 |
| *OQ01_RS37320* | 215 | hypothetical protein | acyltransferase*, Frankia* sp. BMG5.30 (WP_076843553.1); 73/79 | AbyA4 | AbsA4 | AbmA4 |
| *OQ01_RS37325* | 345 | 3-oxoacyl-ACP synthase III family protein | 3-oxoacyl-ACP synthase, *Frankia coriariae* (KLL11317.1); 78/85 | AbyA1 | AbsA1 | AbmA1 |
| *OQ01_RS37330* | 491 | hypothetical protein | monooxygenase FAD-binding protein, *Frankia symbiont of Datisca glomerata* (AEH09375.1); 57/64 | - | - | - |
| *OQ01_RS37335* | 353 | LLM class flavin-dependent oxidoreductase | LLM class flavin-dependent oxidoreductase, *Kutzneria buriramensis* (WP_116181640.1); 70/81 | - | - | AbmE2 |
| *OQ01_RS37340* | 399 | cytochrome P450 | cytochrome P450, *Frankia coriariae* (KLL11355.1); 73/84 | AbyV | AbsV | AbmV |
| *OQ01_RS37345* | 68 | ferredoxin | ferredoxin, *Streptomyces* sp. NRRL WC-3742 (WP_078911468.1); 68/82 | AbyW | AbsG2/AbsG1 | AbmG |
| *OQ01_RS37350* | 313 | alpha/beta hydrolase | alpha/beta hydrolase, *Saccharothrix texasensis* (WP_123742887.1); 67/75 | - | - | - |

**Table S75.** Predicted functions of ORFs in potential BGC from *Saccharothrix syringae* NRRL B-16468 (NZ_JNYO01000017.1).

| ORF | Size (aa) | Proposed function | Closest homolog, host (protein ID); Identity/Similarity (%) | Aby homolog | Abs homolog | Abm homolog |
|---|---|---|---|---|---|---|
| *OQ01_RS21070* | 406 | riboflavin synthase | riboflavin synthase, *Nonomuraea polychroma* (WP_127932388.1); 81/89 | - | - | - |
| *OQ01_RS21075* | 282 | class I SAM-dependent methyltransferase | methyltransferase domain-containing protein, *Nonomuraea polychroma* (WP_127932389.1); 76/83 | - | - | - |
| *OQ01_RS21080* | 131 | hypothetical protein | hypothetical protein, *Saccharothrix* sp. NRRL B-16314 (WP_033438186.1); 73/79 | - | - | - |
| *OQ01_RS21085* | 184 | sigma-70 family RNA polymerase sigma factor | sigma-70 family RNA polymerase sigma factor, *Lentzea jiangxiensis* (WP_090097117.1); 94/96 | - | - | - |
| *OQ01_RS21090* | 344 | hypothetical protein | hypothetical protein, *Lechevalieria atacamensis* (WP_112227485.1); 82/90 | - | - | - |
| *OQ01_RS21095* | 378 | serine/threonine protein kinase | serine/threonine protein kinase, *Saccharothrix carnea* (WP_106615887.1); 70/75 | - | - | - |
| *OQ01_RS21100* | 224 | ArsR family transcriptional regulator | MarR family transcriptional regulator, *Actinokineospora inagensis* (WP_026421213.1); 81/86 | - | - | - |
| *OQ01_RS21105* | 157 | arsenate reductase ArsC | arsenate reductase ArsC, *Streptomyces megasporus* (WP_031511409.1); 88/94 | - | - | - |
| *OQ01_RS21110* | 446 | FAD-dependent oxidoreductase | FAD-dependent oxidoreductase, *Saccharothrix* sp. ALI-22-I (WP_077008908.1); 82/86 | - | - | - |
| *OQ01_RS21115* | 294 | metalloregulator ArsR/SmtB family transcription factor | L-amino acid N-acyltransferase YncA, *Saccharothrix variisporea* (RKT67768.1); 86/91 | - | - | - |
| *OQ01_RS21120* | 153 | nuclear transport factor 2 family protein | nuclear transport factor 2 family protein, *Streptomyces iranensis* (WP_078957437.1); 87/92 | - | - | - |
| *OQ01_RS21125* | 281 | NAD(P)-dependent oxidoreductase | NAD(P)-dependent oxidoreductase, *Streptomyces iranensis* (WP_078957138.1); 87/93 | - | - | - |
| *OQ01_RS21130* | 140 | Diels-Alderase | hypothetical protein, *Streptomyces iranensis* (WP_078957139.1); 88/97 | AbyU | AbsU | AbmU |
| *OQ01_RS21135* | 119 | nuclear transport factor 2 family protein | nuclear transport factor 2 family protein, *Streptomyces cattleya* (WP_014627233.1); 92/98 | - | - | - |
| *OQ01_RS21140* | 417 | cytochrome P450 | cytochrome P450, *Streptomyces iranensis* (WP_044580004.1); 93/95 | - | - | - |
| *OQ01_RS21145* | 6097 | PKS I | type I polyketide synthase, *Streptomyces iranensis* (WP_044580005.1); 89/92 | - | - | - |
| *OQ01_RS21150* | 4278 | PKS I | type I polyketide synthase, *Streptomyces cattleya* (WP_014140914.1); 83/88 | - | - | - |
| *OQ01_RS21155* | 175 | hypothetical protein | pyridoxamine 5'-phosphate oxidase family protein, *Streptomyces iranensis* (WP_044580008.1); 88/94 | - | - | - |
| *OQ01_RS21160* | 3933 | PKS I | type I polyketide synthase, *Streptomyces iranensis* (WP_044580009.1); 89/93 | - | - | - |
| *OQ01_RS21165* | 1385 | PKS I | type I polyketide synthase, *Streptomyces iranensis* (WP_044580010.1); 89/92 | - | - | - |
| *OQ01_RS21170* | 448 | crotonyl-CoA carboxylase/reductase | crotonyl-CoA carboxylase/reductase, *Streptomyces xiamenensis* (WP_046722665.1); 79/89 | - | - | - |
| *OQ01_RS21175* | 336 | ketoacyl-ACP synthase III | ketoacyl-ACP synthase III, *Saccharothrix* sp. NRRL B-16314 (WP_081915703.1); 72/82 | - | - | - |

| | | | | | | |
|---|---|---|---|---|---|---|
| *OQ01_RS21180* | 292 | 3-hydroxybutyryl-CoA dehydrogenase | 3-hydroxybutyryl-CoA dehydrogenase*, Amycolatopsis* sp. 8-3EHSu (WP_130478881.1); 63/78 | - | - | - |
| *OQ01_RS21185* | 408 | cytochrome P450 | cytochrome P450, *Streptomyces iranensis* (WP_078957143.1); 83/90 | - | - | - |
| *OQ01_RS21190* | 68 | thioesterase | thioesterase, *Streptomyces cattleya* (WP_014140907.1); 87/91 | - | - | - |
| *OQ01_RS21195* | 393 | LLM class flavin-dependent oxidoreductase | LLM class flavin-dependent oxidoreductase, *Streptomyces iranensis* (WP_044580012.1); 93/96 | - | - | - |
| *OQ01_RS21200* | 277 | SDR family oxidoreductase | SDR family oxidoreductase, *Streptomyces cattleya* (WP_014140897.1); 91/94 | - | - | - |
| *OQ01_RS50340* | 128 | hypothetical protein | IS5 family transposase, *Actinosynnema* sp. ALI-1.44 (WP_076986720.1); 38/41 | - | - | - |
| *OQ01_RS21205* | 186 | barstar family protein | barnase inhibitor, *Micromonospora saelicesensis* (WP_112675927.1); 74/81 | - | - | - |
| *OQ01_RS50345* | 40 | hypothetical protein | - | - | - | - |
| *OQ01_RS21210* | 390 | glutathione-dependent formaldehyde dehydrogenase | glutathione-dependent formaldehyde dehydrogenase, *Actinomadura fibrosa* (WP_131759867.1); 78/87 | - | - | - |
| *OQ01_RS21215* | 158 | hypothetical protein | hypothetical protein, *Streptosporangium* sp. 'caverna' (WP_110699844.1); 52/65 | - | - | - |

**Table S76.** Predicted functions of ORFs in potential abyssomicin BGC from *Streptomyces* sp. SCA2-2 (NZ_PKMX01000004 and NZ_PKMX01000005).

| ORF | Size (aa) | Proposed function | Closest homolog, host (protein ID); Identity/Similarity (%) | Aby homolog | Abs homolog | Abm homolog |
|---|---|---|---|---|---|---|
| *C0L86_RS05520* | 981 | hypothetical protein | AbmH, *Streptomyces koyangensis* (AVI57436.1); 99/99 | AbyH | - | AbmH |
| *C0L86_RS05525* | 1040 | PKS I | AbmB3, *Streptomyces koyangensis* (AVI57435.1); 99/99 | AbyB3 | AbsB3 | AbmB3 |
| *C0L86_RS05530* | - | PKS I | AbmB2, *Streptomyces koyangensis* (AVI57434.1); 99/99 | AbyB2 | AbsB2 | AbmB2 |
| //////////////////////////////////////////////////////////////////////////////////////////////////////////////////////////////////////////////////////////////////////////////////////////////////// | | | | | | |
| *C0L86_RS05535* | - | PKS I | AbmB1, *Streptomyces koyangensis* (AVI57433.1); 99/99 | AbyB1 | AbsB1 | AbmB1 |
| *C0L86_RS05540* | 178 | flavin reductase family protein | flavin reductase, *Streptomyces* sp. Ru62 (WP_103810817.1); 58/71 | AbyZ | AbsH1 | AbmZ |
| *C0L86_RS05545* | 274 | alpha/beta fold hydrolase | AbmT, *Streptomyces koyangensis* (AVI57431.1); 99/99 | AbyT | AbsN | AbmT |
| *C0L86_RS05550* | 373 | alpha/beta hydrolase | AbmA5, *Streptomyces koyangensis* (AVI57430.1); 99/99 | AbyA5 | AbsA5 | AbmA5 |
| *C0L86_RS05555* | 280 | acyltransferase | AbmA4, *Streptomyces koyangensis* (AVI57429.1); 99/100 | AbyA4 | AbsA4 | AbmA4 |
| *C0L86_RS05560* | 75 | acyl carrier protein | acyl carrier protein, *Streptomyces malaysiense* (WP_071387400.1); 63/78 | AbyA3 | AbsA3 | AbmA3 |
| *C0L86_RS05565* | 628 | HAD-IIIC family phosphatase | AbmA2, *Streptomyces koyangensis* (AVI57427.1); 99/99 | AbyA2 | AbsA2 | AbmA2 |
| *C0L86_RS05570* | 343 | 3-oxoacyl-ACP synthase III family protein | AbmA1, *Streptomyces koyangensis* (AVI57426.1); 99/99 | AbyA1 | AbsA1 | AbmA1 |
| *C0L86_RS05575* | 482 | DHA2 family efflux MFS transporter permease subunit | AbmD, *Streptomyces koyangensis* (AVI57425.1); 99/99 | AbyD | AbsD | AbmD |
| *C0L86_RS05580* | 353 | MsnO8 family LLM class oxidoreductase | AbmE1, *Streptomyces koyangensis* (AVI57424.1); 99/99 | AbyE | AbsE | AbmE1 |
| *C0L86_RS05585* | 257 | TetR/AcrR family transcriptional regulator | TetR/AcrR family transcriptional regulator, *Streptomyces regalis* (WP_062712137.1); 62/75 | AbyC | - | AbmC |
| *C0L86_RS05590* | 405 | cytochrome P450 | cytochrome P450, *Streptomyces* sp. 4R-3d (WP_135068654.1); 69/80 | AbyV | AbsV | AbmV |
| *C0L86_RS05595* | 70 | ferredoxin | AbmG, *Streptomyces koyangensis* (AVI57421.1); 99/100 | - | AbsG1 | AbmG |
| *C0L86_RS05600* | 308 | aldo/keto reductase | AbmJ, *Streptomyces koyangensis* (AVI57420.1); 99/99 | - | AbsJ | AbmJ |
| *C0L86_RS05605* | 546 | ABC transporter substrate-binding protein | AbmF1, *Streptomyces koyangensis* (AVI57419.1); 99/99 | AbyF1 | AbsF1 | AbmF1 |
| *C0L86_RS05610* | 313 | ABC transporter permease | ABC transporter permease, *Streptomyces* sp. 4R-3d (TFI25402.1); 69/82 | AbyF2 | AbsF2 | AbmF2 |
| *C0L86_RS05615* | 413 | amidohydrolase family protein | AbmM, *Streptomyces koyangensis* (AVI57417.1); 99/99 | - | - | AbmM |
| *C0L86_RS0562* | 298 | ABC transporter permease | ABC transporter permease, *Actinomadura macra* (WP_067456459.1); | AbyF3 | AbsF3 | AbmF3 |

| | | | | | | |
|---|---|---|---|---|---|---|
| *0* | | | 63/80 | | | |
| *C0L86_RS05625* | 561 | dipeptide ABC transporter ATP-binding protein | AbmF4, *Streptomyces koyangensis* (AVI57415.1); 99/98 | AbyF4 | AbsF4 | AbmF4 |
| *C0L86_RS05630* | 281 | metallophosphoesterase | AbmL, *Streptomyces koyangensis* (AVI57414.1); 99/99 | - | - | AbmL |
| *C0L86_RS05635* | 253 | 4'-phosphopantetheinyl transferase superfamily protein | AbmK, *Streptomyces koyangensis* (AVI57413.1); 97/97 | - | | AbmK |
| *C0L86_RS05640* | 219 | Diels-Alderase | AbmU, *Streptomyces koyangensis* (AVI57412.1); 99/99 | AbyU | AbsU | AbmU |
| *C0L86_RS05645* | 356 | LLM class flavin-dependent oxidoreductase | LLM class flavin-dependent oxidoreductase, *Saccharothrix syringae* (WP_033434377.1); 57/67 | - | - | AbmE2 |
| *C0L86_RS05650* | 256 | activator protein | AbmI, *Streptomyces koyangensis* (AVI57410.1); 99/100 | AbyI | - | AbmI |

**Table S77.** Predicted functions of ORFs surrounding AbyU homolog from *Streptomyces* sp. SolWspMP-5a-2 (NZ_FMCI01000204).

| ORF | Size (aa) | Proposed function | Closest homolog, host (protein ID); Identity/Similarity (%) | Aby homolog | Abs homolog | Abm homolog |
|---|---|---|---|---|---|---|
| *GA0115242_RS20650* | 510 | MFS transporter | DHA2 family efflux MFS transporter permease subunit, *Streptomyces* sp. NRRL F-5126 (WP_078849779.1); 91/95 | - | - | - |
| *GA0115242_RS20655* | 196 | TetR/AcrR family transcriptional regulator | TetR/AcrR family transcriptional regulator, *Streptomyces* sp. NRRL F-5126 (WP_078849780.1); 90/95 | - | - | - |
| *GA0115242_RS20660* | 493 | hypothetical protein | hypothetical protein, *Streptomyces* sp. NRRL F-5126 (WP_030904229.1); 90/93 | - | - | - |
| *GA0115242_RS20665* | 131 | Diels-Alderase | hypothetical protein, *Streptomyces* sp. NRRL F-5126 (WP_030904231.1); 91/94 | AbyU | AbsU | AbmU |
| *GA0115242_RS20670* | 77 | hypothetical protein | - | - | - | - |
| *GA0115242_RS20675* | 343 | 3-oxoacyl-ACP synthase III family protein | 3-oxoacyl-ACP synthase III family protein, *Streptomyces* sp. NRRL F-5126 (WP_030904235.1); 97/99 | - | - | - |
| *GA0115242_RS20680* | 265 | AfsR/SARP family transcriptional regulator | hypothetical protein, *Streptomyces* sp. NRRL F-5126 (WP_030904237.1); 93/96 | - | - | - |
| *GA0115242_RS20685* | 373 | alpha/beta hydrolase | alpha/beta hydrolase, *Streptomyces* sp. NRRL F-5126 (WP_107059313.1); 90/94 | - | - | - |
| *GA0115242_RS20690* | - | PKS I | type I polyketide synthase, *Streptomyces* sp. NRRL F-5126 (WP_051839911.1); 91/94 | - | - | - |
| *GA0115242_RS20695* | - | PKS I | type I polyketide synthase, *Streptomyces* sp. NRRL F-5126 (WP_051839911.1); 85/87 | - | - | - |
| *GA0115242_RS20700* | 65 | ferredoxin | ferredoxin, *Streptomyces* sp. NRRL F-5126 (WP_030904243.1); 89/96 | - | - | - |
| *GA0115242_RS20705* | 410 | cytochrome P450 | cytochrome P450, *Streptomyces* sp. NRRL F-5126 (WP_030904245.1); 92/95 | - | - | - |
| *GA0115242_RS20710* | 269 | thioesterase | thioesterase, *Streptomyces* sp. NRRL F-5126 (WP_107059319.1); 90/93 | - | - | - |

**Table S78.** Predicted functions of ORFs in potential abyssomicin BGC from *Streptacidiphilus* sp. DSM 106435 (NZ_CP031264.1).

| ORF | Size (aa) | Proposed function | Closest homolog, host (protein ID); Identity/Similarity (%) | Aby homolog | Abs homolog | Abm homolog |
|---|---|---|---|---|---|---|
| *C7M71_RS25195* | 179 | NADPH-dependent FMN reductase | NADPH-dependent FMN reductase, *Streptomyces* sp. CB03911 (WP_073928518.1); 94/96 | AbyZ | AbsH1 | AbmZ |
| *C7M71_RS25200* | 81 | acyl carrier protein | acyl carrier protein, *Streptomyces* sp. CB03911 (WP_073928517.1); 93/95 | AbyA3 | AbsA3 | AbmA3 |
| *C7M71_RS25205* | 638 | HAD-IIIC family phosphatase | HAD-IIIC family phosphatase, *Streptomyces* sp. CB03911 (WP_073928516.1); 94/96 | AbyA2 | AbsA2 | AbmA2 |
| *C7M71_RS25210* | 67 | hypothetical protein | hypothetical protein, *Streptomyces* sp. CB03911 (WP_073928515.1); 68/73 | - | - | - |
| *C7M71_RS25215* | 697 | PKS I | type I polyketide synthase, *Streptomyces* sp. CB03911 (WP_079198462.1); 92/92 | AbyB1 | AbsB1 | AbyB1 |
| *C7M71_RS25220* | 4494 | PKS I | type I polyketide synthase, *Streptomyces* sp. CB03911 (WP_079198462.1); 91/93 | | | |
| *C7M71_RS25225* | 1275 | PKS I | type I polyketide synthase, *Streptomyces* sp. CB03911 (WP_079198462.1); 91/93 | | | |
| *C7M71_RS25230* | 269 | thioesterase | thioesterase, *Streptomyces* sp. CB03911 (WP_079198461.1); 89/91 | AbyT | AbsN | AbmT |
| *C7M71_RS25235* | 295 | AfsR/SARP family transcriptional regulator | AfsR/SARP family transcriptional regulator, *Streptomyces* sp. CB03911 (WP_073928514.1); 96/97 | AbyI | - | AbmI |
| *C7M71_RS25240* | 369 | alpha/beta hydrolase | alpha/beta hydrolase, *Streptomyces* sp. CB03911 (WP_073928513.1); 96/97 | AbyA5 | AbsA5 | AbmA5 |
| *C7M71_RS25245* | 275 | acyltransferase | acyltransferase, *Streptomyces* sp. CB03911 (WP_073928512.1); 95/97 | AbyA4 | AbsA4 | AbmA4 |
| *C7M71_RS25250* | 609 | LuxR family transcriptional regulator | LuxR family transcriptional regulator, *Streptomyces* sp. CB03911 (WP_079198460.1); 91/93 | AbyH | - | AbmH |
| *C7M71_RS25255* | 225 | hypothetical protein | LLM class flavin-dependent oxidoreductase, *Streptomyces* sp. CB03911 (WP_073928509.1); 90/92 | - | - | - |
| *C7M71_RS25260* | 475 | MFS transporter | MFS transporter, *Streptomyces* sp. CB03911 (WP_073928508.1); 94/95 | AbyD | AbsD | AbmD |
| *C7M71_RS25265* | 281 | AfsR/SARP family transcriptional regulator | AfsR/SARP family transcriptional regulator, *Streptomyces* sp. NRRL WC-3742 (WP_063763291.1); 89/93 | AbyI | - | AbmI |
| *C7M71_RS25270* | 330 | aldo/keto reductase | aldo/keto reductase, *Streptomyces* sp. CB03911 (WP_073928507.1); 98/98 | - | AbsJ | AbmJ |
| *C7M71_RS25275* | 388 | acyltransferase | acyltransferase, *Streptomyces* sp. CB03911 (WP_073928506.1); 91/94 | - | AbsI | - |
| *C7M71_RS25280* | 483 | DHA2 family efflux MFS transporter permease subunit | DHA2 family efflux MFS transporter permease subunit, *Streptomyces* sp. NRRL WC-3742 (WP_037973487.1); 87/93 | AbyD | AbsD | AbmD |
| *C7M71_RS25285* | 69 | ferredoxin | ferredoxin, *Streptomyces* sp. CB03911 (WP_073928505.1); 88/94 | - | AbsG2/AbsG1 | AbmG |
| *C7M71_RS25290* | 436 | cytochrome P450 | cytochrome P450, *Streptomyces* sp. CB03911 (WP_079198459.1); 98/99 | AbyV | AbsV | AbmV |
| *C7M71_RS25295* | 125 | Diels-Alderase | hypothetical protein, *Streptomyces* sp. CB03911 (WP_073928710.1); 92/96 | AbyU | AbsU | AbmU |
| *C7M71_RS25300* | 343 | 3-oxoacyl-ACP synthase III family protein | 3-oxoacyl-ACP synthase III family protein, *Streptomyces* sp. CB03911 (WP_073928503.1); 97/97 | AbyA1 | AbsA1 | AbmA1 |

| | | | | | | |
|---|---|---|---|---|---|---|
| *C7M71_RS25305* | 558 | PKS I | type I polyketide synthase, *Streptomyces* sp. CB03911 (WP_073928502.1); 91/93 | AbyB2 | AbsB2 | AbmB2 |
| *C7M71_RS25310* | 91 | PKS I | type I polyketide synthase, *Streptomyces* sp. CB03911 (WP_073928502.1); 89/89 | | | |
| *C7M71_RS25315* | 197 | PKS I | type I polyketide synthase, *Streptomyces* sp. CB03911 (WP_073928502.1); 88/89 | | | |
| *C7M71_RS25320* | 2670 | PKS I | type I polyketide synthase, *Streptomyces* sp. CB03911 (WP_073928502.1); 91/93 | | | |
| *C7M71_RS25325* | 1078 | PKS I | type I polyketide synthase, *Streptomyces* sp. CB03911 (WP_079198458.1); 93/95 | AbyB3 | AbsB3 | AbmB3 |
| *C7M71_RS25330* | 347 | LLM class flavin-dependent oxidoreductase | LLM class flavin-dependent oxidoreductase, *Streptomyces* sp. CB03911 (WP_073928501.1); 95/97 | - | - | AbmE2 |
| *C7M71_RS25335* | 341 | LLM class flavin-dependent oxidoreductase | LLM class flavin-dependent oxidoreductase, *Streptomyces* sp. CB03911 (WP_073928467.1); 95/96 | AbyE | AbsE | AbmE1 |
| *C7M71_RS25340* | 385 | NtaA/DmoA family FMN-dependent monooxygenase | LLM class flavin-dependent oxidoreductase, *Streptomyces* sp. CB03911 (WP_073928466.1); 96/98 | - | - | - |
| *C7M71_RS25345* | 628 | ABC transporter ATP-binding protein | ABC transporter ATP-binding protein, *Streptomyces* sp. CB03911 (WP_079198451.1); 95/96 | AbyF4 | AbsF4 | AbmF4 |
| *C7M71_RS25350* | 272 | ABC transporter permease | ABC transporter permease, *Streptomyces* sp. CB03911 (WP_073928702.1); 93/97 | AbyF3 | AbsF3 | AbmF3 |
| *C7M71_RS25355* | 308 | ABC transporter permease | ABC transporter permease, *Streptomyces* sp. CB03911 (WP_073928465.1); 97/99 | AbyF2 | AbsF2 | AbmF2 |
| *C7M71_RS25360* | 584 | ABC transporter substrate-binding protein | ABC transporter substrate-binding protein, *Streptomyces* sp. CB03911 (WP_079198450.1); 93/95 | AbyF1 | AbsF1 | AbmF1 |
| *C7M71_RS25365* | 316 | alpha/beta hydrolase | alpha/beta hydrolase, *Streptomyces* sp. CB03911 (WP_079198449.1); 94/95 | - | - | - |
| *C7M71_RS25370* | 220 | TetR/AcrR family transcriptional regulator | TetR/AcrR family transcriptional regulator, *Streptomyces* sp. CB03911 (WP_073928463.1); 99/99 | AbyC | - | AbmC |

**Table S79.** Predicted functions of ORFs surrounding AbyU homolog from *Streptomyces hoynatensis* KCTC 29097 (NZ_RBAL01000026.1).

| ORF | Size (aa) | Proposed function | Closest homolog, host (protein ID); Identity/Similarity (%) | Aby homolog | Abs homolog | Abm homolog |
|---|---|---|---|---|---|---|
| *D7294_RS28425* | 233 | TetR/AcrR family transcriptional regulator | TetR family transcriptional regulator, *Actinomadura* sp. KC345 (WP_131884222.1); 77/86 | - | - | - |
| *D7294_RS28430* | 740 | MMPL family transporter | membrane protein, *Streptomyces olindensis* (KDN75573.1); 60/75 | - | - | - |
| *D7294_RS28435* | 254 | AfsR/SARP family transcriptional regulator | activator protein, *Actinomadura* sp. 7K534 (WP_132046165.1); 56/69 | - | - | - |
| *D7294_RS28440* | 181 | Diels-Alderase | hypothetical protein, *Streptomyces* sp. NRRL S-350 (WP_030245418.1); 34/55 | AbyU | AbsU | AbmU |
| *D7294_RS28445* | 261 | thioesterase | thioesterase, *Streptomyces* sp. AZ1-7 (RKN06823.1); 57/68 | - | - | - |
| *D7294_RS28450* | 448 | hypothetical protein | hypothetical protein, *Streptomyces* sp. MP131-18 (WP_079251913.1); 72/84 | - | - | - |
| *D7294_RS28455* | 441 | glycosyl transferase | hypothetical protein, *Catenuloplanes japonicus* (WP_033345696.1); 50/65 | - | - | - |
| *D7294_RS28460* | 503 | MFS transporter | DHA2 family efflux MFS transporter permease subunit, *Frankia* sp. BMG5.36 (WP_071049484.1); 53/68 | - | - | - |
| *D7294_RS28465* | 290 | LLM class F420-dependent oxidoreductase | LLM class F420-dependent oxidoreductase, *Frankia* sp. BMG5.36 (WP_071051364.1); 68/78 | - | - | - |
| *D7294_RS28470* | 1550 | PKS I | polyketide synthase, *Streptomyces* sp. 211726 (ARM20279.1); 51/62 | - | - | - |
| *D7294_RS28475* | 4001 | PKS I | VerV, *Actinomadura* sp. XM-4-3 (AYW35158.1); 60/70 | - | - | - |
| *D7294_RS28480* | 2194 | PKS I | type I polyketide synthase, *Actinomadura macra* (WP_067456433.1); 59/69 | - | - | - |
| *D7294_RS28485* | 584 | PKS I | SDR family NAD(P)-dependent oxidoreductase, partial, *Streptomyces* sp. WAC 01420 (WP_126896698.1); 57/66 | - | - | - |
| *D7294_RS28490* | 373 | PKS I | Polyketide synthase dehydratase, partial, *Streptomyces* sp. MnatMP-M27 (SCG12060.1); 51/62 | - | - | - |
| *D7294_RS28495* | 830 | PKS I | type I polyketide synthase, *Streptomyces* sp. SBT349 (WP_053171085.1); 69/79 | - | - | - |
| *D7294_RS28500* | 296 | PKS I | SDR family NAD(P)-dependent oxidoreductase, *Streptacidiphilus neutrinimicus* (WP_084729882.1); 68/76 | - | - | - |

**Table S80.** Predicted functions of ORFs in abyssomicin BGC from *Streptomyces* sp. 57 (NZ_RCCZ01000005.1).

| ORF | Size (aa) | Proposed function | Closest homolog, host (protein ID); Identity/Similarity (%) | Aby homolog | Abs homolog | Abm homolog |
|---|---|---|---|---|---|---|
| *CLZ79_6704* | 902 | regulatory LuxR family protein | LuxR family transcriptional regulator, *Streptomyces regalis* (WP_062712128.1); 52/63 | AbyH | - | AbmH |
| *CLZ79_6705* | 861 | RHS repeat-associated protein | RHS repeat protein, *Streptomyces fragilis* (WP_108953524.1); 59/67 | AbyK | - | - |
| *CLZ79_6706* | 135 | Diels-Alderase | hypothetical protein, *Streptomyces regalis* (WP_062712130.1); 87/93 | AbyU | AbsU | AbmU |
| *CLZ79_6707* | 345 | 3-oxoacyl-ACP synthase III family protein | 3-oxoacyl-ACP synthase III family protein, *Frankia* sp. EAN1pec (WP_020461027.1); 78/88 | AbyA1 | AbsA1 | AbmA1 |
| *CLZ79_6708* | 633 | HAD-IIIC family phosphatase | methoxymalonyl-ACP biosynthesis protein FkbH, *Streptomyces regalis* (KUL23675.1); 70/77 | AbyA2 | AbsA2 | AbmA2 |
| *CLZ79_6709* | 75 | acyl carrier protein | acyl carrier protein, *Streptomyces* sp. NRRL WC-3725 (WP_031029037.1); 75/85 | AbyA3 | AbsA3 | AbmA3 |
| *CLZ79_6710* | 255 | acyltransferase | 2-oxoacid dehydrogenase/acyltransferase catalytic subunit, *Streptomyces* sp. BK438 (TCP45291.1); 76/84 | AbyA4 | AbsA4 | AbmA4 |
| *CLZ79_6711* | 362 | alpha/beta hydrolase | alpha/beta hydrolase, *Streptomyces* sp. 2131.1 (WP_093709996.1); 81/87 | AbyA5 | AbsA5 | AbmA5 |
| *CLZ79_6712* | 165 | flavin reductase | flavin reductase, *Streptomyces fragilis* (WP_108952936.1); 78/85 | AbyZ | AbsH1 | AbmZ |
| *CLZ79_6713* | 223 | TetR/AcrR family transcriptional regulator | TetR/AcrR family transcriptional regulator, *Streptomyces* sp. KhCrAH-43 (WP_018522887.1); 86/93 | AbyC | - | AbmC |
| *CLZ79_6714* | 482 | DHA2 family efflux MFS transporter permease subunit | DHA2 family efflux MFS transporter permease subunit, *Streptomyces* sp. NRRL WC-3744 (WP_030991264.1); 85/91 | AbyD | AbsD | AbmD |
| *CLZ79_6715* | 348 | LLM class flavin-dependent oxidoreductase | LLM class flavin-dependent oxidoreductase, *Streptomyces* sp. KhCrAH-43 (WP_018522885.1); 77/85 | AbyE | AbsE | AbmE1 |
| *CLZ79_6716* | 544 | ABC transporter substrate-binding protein | ABC transporter substrate-binding protein, *Streptomyces* sp. KhCrAH-43 (WP_018522884.1); 76/84 | AbyF1 | AbsF1 | AbmF1 |
| *CLZ79_6717* | 319 | ABC transporter permease | ABC transporter permease, *Streptomyces* sp. CB01249 (WP_073865209.1); 76/87 | AbyF2 | AbsF2 | AbmF2 |
| *CLZ79_6718* | 286 | ABC transporter permease | ABC transporter permease, *Herbidospora cretacea* (WP_034385023.1); 77/83 | AbyF3 | AbsF3 | AbmF3 |
| *CLZ79_6719* | 539 | ABC transporter ATP-binding protein | ABC transporter ATP-binding protein, *Streptomyces* sp. 2131.1 (WP_093709989.1); 75/83 | AbyF4 | AbsF4 | AbmF4 |
| *CLZ79_6720* | 392 | acyltransferase | acyltransferase ,*Streptomyces* sp. KhCrAH-43 (WP_018522880.1); 63/71 | - | AbsI | - |
| *CLZ79_6721* | 407 | cytochrome P450 | cytochrome P450, *Streptomyces fragilis* (WP_108952944.1); 84/89 | AbyV | AbsV | AbmV |
| *CLZ79_6722* | 82 | ferredoxin | ferredoxin, *Streptomyces regensis* (KMS84448.1); 84/87 | - | AbmG1 | AbmG |
| *CLZ79_6723* | 294 | ferredoxin | alpha/beta hydrolase, *Streptomyces fragilis* (WP_108952946.1); 72/80 | - | - | - |
| *CLZ79_6724* | 357 | LLM class flavin-dependent oxidoreductase | LLM class flavin-dependent oxidoreductase, *Frankia* sp. EAN1pec (WP_020461009.1); 56/67 | - | - | AbmE2 |
| *CLZ79_6725* | 251 | SDR family oxidoreductase | SDR family oxidoreductase, *Streptoalloteichus hindustanus* (WP_073481867.1); 55/66 | - | - | - |
| *CLZ79_6726* | 6183 | PKS I | ype I polyketide synthase, *Streptomyces* sp. KhCrAH-43 (WP_018522876.1); 64/71 | AbyB1 | AbsB1 | AbmB1 |

| | | | | | | |
|---|---|---|---|---|---|---|
| *CLZ79_RS33565* | 3864 | PKS I | SDR family NAD(P)-dependent oxidoreductase, *Streptomyces* sp. BK438 (WP_132903672.1); 68/76 | AbyB2 | AbsB2 | AbmB2 |
| *CLZ79_RS33570* | 1017 | PKS I | type I polyketide synthase, *Streptomyces* sp. KhCrAH-43 (WP_018522873.1); 72/78 | AbyB3 | AbsB3 | AbmB3 |
| *CLZ79_RS33575* | 406 | cytochrome P450 | cytochrome P450, *Streptomyces* sp. BK438 (WP_132903670.1); 80/87 | AbyX/AbyV | AbsV/AbsX | AbmV |

**Table S81.** Predicted functions of ORFs in potential abyssomicin BGC from *Streptomyces subroseum* CGMCC 4.2132 (NZ_FZOD01000006.1).

| ORF | Size (aa) | Proposed function | Closest homolog, host (protein ID); Identity/Similarity (%) | Aby homolog | Abs homolog | Abm homolog |
|---|---|---|---|---|---|---|
| *CHC08_RS08495* | 320 | LLM class flavin-dependent oxidoreductase | MsnO8 family LLM class oxidoreductase, *Microbispora triticiradicis* (WP_117408853.1); 85/90 | AbyE | AbsE | AbmE1 |
| *CHC08_RS08500* | 559 | ABC transporter substrate-binding protein | ABC transporter substrate-binding protein, *Microbispora triticiradicis* (WP_117408852.1); 77/84 | AbyF1 | AbsF1 | AbmF1 |
| *CHC08_RS08505* | 331 | ABC transporter permease | ABC transporter permease, *Microbispora triticiradicis* (WP_117408849.1); 77/85 | AbyF2 | AbsF2 | AbmF2 |
| *CHC08_RS08510* | 268 | ABC transporter permease | AbsF3, *Streptomyces* sp. LC-6-2 (ARE67846.1); 80/88 | AbyF3 | AbsF3 | AbmF3 |
| *CHC08_RS08515* | 555 | ABC transporter ATP-binding protein | AbsF4, *Streptomyces* sp. LC-6-2 (ARE67845.1); 73/81 | AbyF4 | AbsF4 | AbmF4 |
| *CHC08_RS08525* | 273 | acyltransferase | AbsI, *Streptomyces* sp. LC-6-2 (ARE67842.1); 66/76 | - | AbsI | - |
| *CHC08_RS08530* | 408 | cytochrome P450 | cytochrome P450, *Microbispora triticiradicis* (WP_117409466.1); 85/91 | AbyX | AbsX | AbmV |
| *CHC08_RS08535* | 72 | ferredoxin | ferredoxin, *Microbispora triticiradicis* (WP_117409458.1); 87/91 | - | AbsG2/AbsG1 | AbmG |
| *CHC08_RS08540* | 332 | aldo/keto reductase | aldo/keto reductase, *Microbispora triticiradicis* (WP_117409459.1); 82/89 | - | AbsJ | AbmJ |
| *CHC08_RS08545* | 128 | Diels-Alderase | hypothetical protein, *Microbispora triticiradicis* (WP_117409467.1); 90/96 | AbyU | AbsU | AbmU |
| *CHC08_RS08550* | 928 | hypothetical protein | LuxR family transcriptional regulator, *Microbispora triticiradicis* (WP_133306130.1); 71/80 | AbyH | - | AbmH |
| *CHC08_RS08555* | 198 | TetR/AcrR family transcriptional regulator | TetR/AcrR family transcriptional regulator, *Microbispora* sp. GKU 823 (WP_079317081.1); 82/87 | - | AbsC2 | - |
| *CHC08_RS08560* | 479 | MFS transporter | MFS transporter, *Microbispora* sp. GKU 823 (WP_079317079.1); 79/87 | AbyD | AbsD | AbmD |
| *CHC08_RS08565* | 252 | AfsR/SARP family transcriptional regulator | AfsR/SARP family transcriptional regulator, *Microbispora triticiradicis* (WP_117409463.1); 83/90 | AbyR/AbyI | - | AbmI |

**Table S82.** Predicted functions of ORFs in potential BGC from *Streptomyces varsoviensis* NRRL B-3589 (NZ_JOFN01000010.1).

| ORF | Size (aa) | Proposed function | Closest homolog, host (protein ID); Identity/Similarity (%) | Aby homolog | Abs homolog | Abm homolog |
|---|---|---|---|---|---|---|
| *IF95_RS0115745* | 409 | MFS transporter | MFS transporter, *Streptacidiphilus jiangxiensis* (WP_042456014.1); 33/51 | - | - | - |
| *IF95_RS0115750* | 1347 | PKS I | type I polyketide synthase, *Streptomyces iranensis* (WP_044580010.1); 86/91 | - | - | - |
| *IF95_RS0115755* | 3948 | PKS I | type I polyketide synthase, *Streptomyces cattleya* (WP_014140917.1); 90/93 | - | - | - |
| *IF95_RS0115760* | 172 | pyridoxamine 5'-phosphate oxidase family protein | pyridoxamine 5'-phosphate oxidase family protein, *Streptomyces cattleya* (WP_014140915.1); 87/93 | - | - | - |
| *IF95_RS39245* | 1594 | PKS I | type I polyketide synthase, *Streptomyces cattleya* (WP_014140914.1); 87/90 | - | - | - |
| *IF95_RS39250* | 2101 | PKS I | type I polyketide synthase, *Streptomyces cattleya* (WP_014140914.1); 88/91 | - | - | - |
| *IF95_RS37565* | 5303 | PKS I | type I polyketide synthase, *Streptomyces cattleya* (WP_014140913.1); 90/92 | - | - | - |
| *IF95_RS0115780* | 416 | cytochrome P450 | cytochrome P450, *Streptomyces* sp. E5N91 SAI-083 (WP_123627589.1); 91/95 | - | - | - |
| *IF95_RS0115785* | 119 | nuclear transport factor 2 family protein | nuclear transport factor 2 family protein, *Streptomyces cattleya* (WP_014627233.1); 94/96 | - | - | - |
| *IF95_RS0115790* | 140 | Diels-Alderase | hypothetical protein, *Streptomyces cattleya* (WP_014140910.1); 96/98 | AbyU | AbsU | AbmU |
| *IF95_RS0115795* | 280 | NAD(P)-dependent oxidoreductase | NAD(P)-dependent oxidoreductase, *Streptomyces cattleya* (WP_014140909.1); 93/97 | - | - | - |
| *IF95_RS0115800* | 149 | nuclear transport factor 2 family protein | nuclear transport factor 2 family protein, *Streptomyces cattleya* (WP_014627231.1); 90/94 | - | - | - |
| *IF95_RS0115805* | 265 | thioesterase | thioesterase, *Streptomyces cattleya* (WP_014140907.1); 90/94 | - | - | - |
| *IF95_RS0115810* | 408 | cytochrome P450 | cytochrome P450, *Streptomyces* sp. E5N91 SAI-083 (WP_123627594.1); 90/94 | - | - | - |
| *IF95_RS0115815* | 924 | LuxR family transcriptional regulator | helix-turn-helix transcriptional regulator, *Streptomyces cattleya* (WP_014140903.1); 89/93 | - | - | - |
| *IF95_RS0115820* | 454 | crotonyl-CoA carboxylase/reductase | crotonyl-CoA carboxylase/reductase, *Streptomyces olivaceus* (WP_070390081.1); 96/97 | - | - | - |
| *IF95_RS0115825* | 145 | hypothetical protein | hypothetical protein EDC84_6806, *Streptomyces* sp. E5N91 SAI-083 (ROO97928.1); 89/91 | - | - | - |
| *IF95_RS0115830* | 277 | SDR family NAD(P)-dependent oxidoreductase | SDR family oxidoreductase, *Streptomyces cattleya* (WP_014140897.1);95/97 | - | - | - |
| *IF95_RS0115835* | 393 | LLM class flavin-dependent oxidoreductase | LLM class flavin-dependent oxidoreductase, *Streptomyces cattleya* (WP_014140896.1); 94/96 | - | - | - |

**Table S83.** Predicted functions of ORFs in potential abyssomicin BGC from *Actinocrispum wychmicini* DSM 45934 (NZ_SLWS01000002).

| ORF | Size (aa) | Proposed function | Closest homolog, host (protein ID); Identity/Similarity (%) | Aby homolog | Abs homolog | Abm homolog |
|---|---|---|---|---|---|---|
| *EV192_RS10885* | 196 | TetR family transcriptional regulator | TetR/AcrR family transcriptional regulator, *Corallococcus* sp. H22C18031201 (WP_120202919.1); 55/70 | - | AbsC2 | - |
| *EV192_RS10890* | 492 | DHA2 family efflux MFS transporter permease subunit | DHA2 family efflux MFS transporter permease subunit, *Streptomyces formicae* (WP_098241238.1); 54/72 | AbyD | AbsD | AbmD |
| *EV192_RS10895* | 444 | FAD-dependent oxidoreductase | monooxygenase, *Frankia canadensis* (WP_101832044.1); 54/65 | - | - | - |
| *EV192_RS10900* | 396 | cytochrome P450 | cytochrome P450, *Actinoplanes* sp. N902-109 (WP_015620517.1); 51/70 | AbyX/AbyV | AbsV/AbsX | AbmV |
| *EV192_RS10905* | 346 | MsnO8 family LLM class oxidoreductase | luciferase family oxidoreductase, group 1, *Kibdelosporangium aridum* (SMD20744.1); 64/76 | AbyE | AbsE | AbmE1 |
| *EV192_RS10910* | 534 | dipeptide ABC transporter ATP-binding protein | AbmF4, *Streptomyces koyangensis* (AVI57415.1); 63/77 | AbyF4 | AbsF4 | AbmF4 |
| *EV192_RS10915* | 282 | ABC transporter permease subunit | ABC transporter permease, *Actinomadura macra* (WP_067456459.1); 64/76 | AbyF3 | AbsF3 | AbmF3 |
| *EV192_RS10920* | 325 | ABC transporter permease subunit | ABC transporter permease, *Actinomadura* sp. 5-2 (WP_103566287.1); 66/79 | AbyF2 | AbsF2 | AbmF2 |
| *EV192_RS10925* | 587 | ABC transporter substrate-binding protein | ABC transporter substrate-binding protein, *Streptomyces formicae* (WP_098241233.1); 56/69 | AbyF1 | AbsF1 | AbmF1 |
| *EV192_RS10930* | 264 | SDR family oxidoreductase | SDR family oxidoreductase, *Actinomadura macra* (WP_084265034.1); 66/76 | - | - | - |
| *EV192_RS10935* | 138 | hypothetical protein | hypothetical protein, *Actinomadura meyerae* (WP_089329757.1); 53/70 | - | - | - |
| *EV192_RS10940* | 312 | 3-oxoacyl-ACP synthase III family protein | 3-oxoacyl-ACP synthase III family protein, *Streptomyces* sp. E5N91 SAI-083 (WP_123627122.1); 66/81 | AbyA1 | AbsA1 | AbmA1 |
| *EV192_RS10945* | 75 | acyl carrier protein | acyl carrier protein, *Frankia* sp. ACN1ag (KQC35070.1); 66/82 | AbyA3 | AbsA3 | AbmA3 |
| *EV192_RS10950* | 216 | acyltransferase | acyltransferase, *Streptomyces armeniacus* (AXK32424.1); 69/81 | AbyA4 | AbsA4 | AbmA4 |
| *EV192_RS10955* | 356 | alpha/beta hydrolase | alpha/beta hydrolase, *Streptomyces* sp. NRRL F-525 (WP_033287161.1); 62/74 | AbyA5 | AbsA5 | AbmA5 |
| *EV192_RS10960* | 4517 | PKS I | QmnA1, *Amycolatopsis orientalis* (AFI57005.1); 56/66 | PKS I | PKS I | PKS I |
| *EV192_RS10965* | 2460 | PKS I | type-I PKS, *Streptomyces noursei* (PNE39797.1); 53/65 | PKS I | PKS I | PKS I |
| *EV192_RS10970* | 861 | PKS I | SDR family NAD(P)-dependent oxidoreductase, *Streptomyces lydicus* (WP_129293092.1); 57/69 | PKS I | PKS I | PKS I |
| *EV192_RS10975* | 240 | PKS I | type I polyketide synthase, *Allokutzneria* sp. NRRL B-24872 (WP_086824400.1); 60/68 | PKS I | PKS I | PKS I |
| *EV192_RS10980* | 292 | PKS I | polyketide synthase, partial, *Streptomyces rubellomurinus subsp. indigoferus* (KJS52288.1); 65/74 | PKS I | PKS I | PKS I |
| *EV192_RS10985* | 132 | PKS I | polyketide synthase, partial, *Streptomyces hygroscopicus subsp. hygroscopicus* (BAH67173.1); 65/74 | PKS I | PKS I | PKS I |
| *EV192_RS10990* | 325 | PKS I | ChlA1, *Streptomyces antibioticus* (AAZ77693.1); 60/74 | PKS I | PKS I | PKS I |
| *EV192_RS10995* | 1266 | PKS I | type I polyketide synthase, *Streptomyces* sp. NBRC 109436 (WP_079150001.1); 57/67 | PKS I | PKS I | PKS I |

| | | | | | | |
|---|---|---|---|---|---|---|
| *EV192_RS11000* | 314 | activator protein | activator protein, *Kutzneria buriramensis* (WP_116181646.1); 69/78 | AbyI/AbyR | - | AbmI |

**Table S84.** Predicted functions of ORFs in potential chlorothricin BGC from *Actinocrispum wychmicini* DSM 45934 (NZ_SLWS01000003).

| ORF | Size (aa) | Proposed function | Closest homolog, host (protein ID); Identity/Similarity (%) | Aby homolog | Abs homolog | Abm homolog |
|---|---|---|---|---|---|---|
| *EV192_RS16140* | 338 | beta-ketoacyl-ACP synthase 3 | ketoacyl-ACP synthase III, *Streptomyces diastatochromogenes* (WP_094222628.1); 71/82 | - | - | - |
| *EV192_RS16145* | 291 | 3-hydroxybutyryl-CoA dehydrogenase | 3-hydroxybutyryl-CoA dehydrogenase, *Streptomyces* sp. NRRL B-3648 (WP_053711641.1); 68/79 | - | - | - |
| *EV192_RS16150* | 520 | acyl-CoA carboxylase subunit beta | acyl-CoA carboxylase subunit beta, *Amycolatopsis orientalis* (WP_043836795.1); 83/90 | - | - | - |
| *EV192_RS16155* | 345 | aldo/keto reductase | aldo/keto reductase, *Streptomyces* sp. DSM 15324 (WP_079079511.1); 76/84 | - | - | - |
| *EV192_RS16160* | 462 | NDP-hexose 2,3-dehydratase | NDP-hexose 2,3-dehydratase, *Streptomyces* sp. Ru73 (WP_103830972.1); 70/81 | - | - | - |
| *EV192_RS16165* | 404 | cytochrome P450 | cytochrome P450, *Actinomadura pelletieri* (WP_121438133.1); 61/71 | - | - | - |
| *EV192_RS16170* | 79 | hypothetical protein | hypothetical protein, *Stackebrandtia nassauensis* (WP_013019899.1); 69/80 | - | - | - |
| *EV192_RS16175* | 158 | hypothetical protein | Clp protease, *Nakamurella* sp. 12Sc4-1 (WP_111766871.1); 45/60 | - | - | - |
| *EV192_RS16180* | 490 | DHA2 family efflux MFS transporter permease subunit | DHA2 family efflux MFS transporter permease subunit, *Plantactinospora* sp. KBS50 (WP_095565847.1); 52/66 | - | - | - |
| *EV192_RS16185* | 927 | LuxR family transcriptional regulator | LuxR family transcriptional regulator, *Streptomyces armeniacus* (AXK33357.1); 45/58 | - | - | - |
| *EV192_RS16190* | 354 | alpha/beta hydrolase | alpha/beta hydrolase, *Actinomadura pelletieri* (WP_121438111.1); 65/77 | - | - | - |
| *EV192_RS16195* | 256 | acyltransferase | acyltransferase, *Actinomadura pelletieri* (WP_121438112.1); 71/81 | - | - | - |
| *EV192_RS16200* | 76 | acyl carrier protein | acyl carrier protein, *Streptomyces armeniacus* (AXK32426.1); 60/80 | - | - | - |
| *EV192_RS16205* | 634 | HAD-IIIC family phosphatase | HAD-IIIC family phosphatase, *Actinomadura pelletieri* (WP_121438114.1); 63/76 | - | - | - |
| *EV192_RS16210* | 343 | 3-oxoacyl-ACP synthase III family protein | 3-oxoacyl-ACP synthase III family protein, *Actinomadura pelletieri* (WP_121438115.1); 76/87 | - | - | - |
| *EV192_RS16215* | 474 | monooxygenase | hypothetical protein, *Actinomadura pelletieri* (WP_121438116.1); 62/71 | - | - | - |
| *EV192_RS16220* | 1355 | PKS I | acyltransferase domain-containing protein, *Actinomadura pelletieri* (WP_121438117.1); 55/64 | - | - | - |
| *EV192_RS16225* | 2904 | PKS I | SDR family NAD(P)-dependent oxidoreductase, *Actinomadura pelletieri* (WP_121438118.1); 63/74 | - | - | - |
| *EV192_RS16230* | 1074 | PKS I | SDR family NAD(P)-dependent oxidoreductase, *Actinomadura pelletieri* (WP_121438118.1); 70/79 | - | - | - |
| *EV192_RS16235* | 239 | PKS I | Phosphopantetheine attachment site, partial, *Micromonospora matsumotoense* (SCF50095.1); 53/65 | - | - | - |
| *EV192_RS16240* | 355 | PKS I | SDR family NAD(P)-dependent oxidoreductase, *Actinomadura pelletieri* (WP_121438352.1); 60/70 | - | - | - |

| | | | | | | |
|---|---|---|---|---|---|---|
| *EV192_RS16245* | 759 | PKS I | SDR family NAD(P)-dependent oxidoreductase, *Streptomyces* sp. 11-1-2 (WP_119988543.1); 65/76 | - | - | - |
| *EV192_RS16250* | 1261 | PKS I | modular polyketide synthase, *Streptomyces* sp. RK95-74 (BAW35613.1); 52/64 | - | - | - |
| *EV192_RS16255* | 680 | PKS I | ChlA3, *Streptomyces antibioticus* (AAZ77696.1); 65/76 | - | - | - |
| *EV192_RS16260* | 301 | PKS I | SDR family NAD(P)-dependent oxidoreductase, *Streptomyces alboflavus* (WP_125262907.1); 65/73 | - | - | - |
| *EV192_RS16265* | 340 | PKS I | SDR family NAD(P)-dependent oxidoreductase, *Actinomadura pelletieri* (WP_121438119.1); 57/69 | - | - | - |
| *EV192_RS16270* | 130 | PKS I | QmnA3, *Amycolatopsis orientalis* (AFI57007.1); 63/75 | - | - | - |
| *EV192_RS16275* | 245 | PKS I | SDR family NAD(P)-dependent oxidoreductase, *Actinomadura pelletieri* (WP_121438119.1); 83/89 | - | - | - |
| *EV192_RS16280* | 159 | PKS I | SDR family NAD(P)-dependent oxidoreductase, partial, *Streptomyces* sp. AM-2504 (WP_131124269.1); 58/68 | - | - | - |
| *EV192_RS16285* | 314 | PKS I | KR domain-containing protein, partial, *Streptomyces* sp. MnatMP-M27 (SCG13790.1); 65/74 | - | - | - |
| *EV192_RS16290* | 899 | PKS I | SDR family NAD(P)-dependent oxidoreductase, *Actinomadura pelletieri* (WP_121438119.1); 62/73 | - | - | - |
| *EV192_RS16295* | 1566 | PKS I | SDR family NAD(P)-dependent oxidoreductase, *Actinomadura pelletieri* (WP_121438120.1); 57/67 | - | - | - |
| *EV192_RS16300* | 4361 | PKS I | Ann4, *Streptomyces calvus* (AGY30676.1); 52/63 | - | - | - |
| *EV192_RS16305* | 386 | acyl-CoA dehydrogenase | acyl-CoA dehydrogenase, *Saccharomonospora saliphila* (WP_019815635.1); 69/80 | - | - | - |
| *EV192_RS16310* | 401 | DUF1205 domain-containing protein | DUF1205 domain-containing protein, *Actinomadura pelletieri* (WP_121438132.1); 48/65 | - | - | - |
| *EV192_RS16315* | 384 | aminotransferase class I/II-fold pyridoxal phosphate-dependent enzyme | aminotransferase class I/II-fold pyridoxal phosphate-dependent enzyme, *Amycolatopsis japonica* (WP_051972467.1); 62/75 | - | - | - |
| *EV192_RS16320* | 347 | 3-oxoacyl-ACP synthase | 3-oxoacyl-ACP synthase, *Actinomadura pelletieri* (WP_121438128.1); 74/84 | - | - | - |
| *EV192_RS16325* | 1724 | acyltransferase domain-containing protein | acyltransferase domain-containing protein, *Actinomadura pelletieri* (WP_121438127.1); 61/72 | - | - | - |
| *EV192_RS16330* | 448 | FAD-dependent oxidoreductase | NAD(P)/FAD-dependent oxidoreductase, *Actinomadura pelletieri* (WP_121438126.1); 79/88 | - | - | - |
| *EV192_RS16335* | 88 | acyl carrier protein | ChlB2, *Streptomyces antibioticus* (AAZ77675.1); 55/72 | - | - | - |
| *EV192_RS16340* | 349 | 3-oxoacyl-ACP synthase | 3-oxoacyl-ACP synthase, *Micromonospora haikouensis* (WP_091284724.1); 58/71 | - | - | - |
| *EV192_RS16345* | 261 | alpha/beta fold hydrolase | thioesterase, *Streptomyces aurantiacus* (WP_055507818.1); 59/72 | - | - | - |
| *EV192_RS16350* | 256 | activator protein | activator protein, *Actinomadura* sp. LMG 30035 (WP_131738462.1); 67/80 | - | - | - |
| *EV192_RS16355* | 182 | Diels-Alderase | hypothetical protein, *Actinomadura pelletieri* (WP_121438130.1); 50/62 | AbyU | AbsU | AbmU |

| | | | | | | |
|---|---|---|---|---|---|---|
| *EV192_RS1636 0* | 344 | methyltransferase | methyltransferase family protein, *Actinocrispum wychmicini* (TCO60814.1); 99/100 | - | - | - |

**Table S85.** Predicted genomic islands nearby the abyssomicin BGC from *Actinokineospora auranticolor* YU 961-1 (PTIX01000011.1).

| Island number | Island start | Island end | Length | Method | Gene name | Locus | Gene start | Gene end | Strand | Product |
|---|---|---|---|---|---|---|---|---|---|---|
| 1 | 183112 | 188952 | 5840 | IslandPath-DIMOB | PPK66181.1 | CLV40_111145 | 182417 | 183133 | 1 | hypothetical protein |
| | 183112 | 188952 | 5840 | IslandPath-DIMOB | PPK66182.1 | CLV40_111146 | 183112 | 183891 | -1 | DDE superfamily endonuclease |
| | 183112 | 188952 | 5840 | IslandPath-DIMOB | PPK66183.1 | CLV40_111147 | 184081 | 184374 | 1 | transposase |
| | 183112 | 188952 | 5840 | IslandPath-DIMOB | PPK66184.1 | CLV40_111148 | 184371 | 185255 | 1 | transposase InsO family protein |
| | 183112 | 188952 | 5840 | IslandPath-DIMOB | PPK66185.1 | CLV40_111149 | 185345 | 185479 | -1 | hypothetical protein |
| | 183112 | 188952 | 5840 | IslandPath-DIMOB | PPK66186.1 | CLV40_111150 | 186133 | 187383 | -1 | deoxyribonuclease NucA/NucB |
| | 183112 | 188952 | 5840 | IslandPath-DIMOB | PPK66187.1 | CLV40_111151 | 187781 | 188074 | 1 | transposase-like protein |
| | 183112 | 188952 | 5840 | IslandPath-DIMOB | PPK66188.1 | CLV40_111152 | 188134 | 188952 | 1 | putative transposase |

**Table S86.** Predicted genomic islands in the abyssomicin BGC from *Frankia sp.* EAN1pec (CP000820.1).

| Island number | Island start | Island end | Length | Method | Gene name | Locus | Gene start | Gene end | Strand | Product |
|---|---|---|---|---|---|---|---|---|---|---|
| 1 | 4037626 | 4083572 | 45946 | IslandPath-DIMOB | WP_020460948.1 | FRANEAN1_RS16505 | 4036943 | 4037629 | 1 | hydrogenase |
| | 4037626 | 4083572 | 45946 | IslandPath-DIMOB | WP_020460949.1 | FRANEAN1_RS16510 | 4037626 | 4038201 | 1 | HybD peptidase |
| | 4037626 | 4083572 | 45946 | IslandPath-DIMOB | WP_020460950.1 | FRANEAN1_RS16515 | 4038194 | 4038514 | 1 | hypothetical protein |
| | 4037626 | 4083572 | 45946 | IslandPath-DIMOB | WP_041254273.1 | FRANEAN1_RS16520 | 4038597 | 4039307 | 1 | transposase |
| | 4037626 | 4083572 | 45946 | IslandPath-DIMOB | | FRANEAN1_RS16525 | 4039538 | 4040605 | -1 | integrase |
| | 4037626 | 4083572 | 45946 | IslandPath-DIMOB | | FRANEAN1_RS16530 | 4041090 | 4042106 | 1 | transposase |
| | 4037626 | 4083572 | 45946 | IslandPath-DIMOB | WP_041254274.1 | FRANEAN1_RS16535 | 4042771 | 4044030 | 1 | group II intron reverse transcriptase/maturase |
| | 4037626 | 4083572 | 45946 | IslandPath-DIMOB | WP_041254276.1 | FRANEAN1_RS16540 | 4044349 | 4045413 | -1 | integrase |
| | 4037626 | 4083572 | 45946 | IslandPath-DIMOB | WP_020460958.1 | FRANEAN1_RS16545 | 4045559 | 4046086 | 1 | transposase |
| | 4037626 | 4083572 | 45946 | IslandPath-DIMOB | WP_020460959.1 | FRANEAN1_RS16550 | 4046250 | 4047026 | 1 | endonuclease |
| | 4037626 | 4083572 | 45946 | IslandPath-DIMOB | WP_020460962.1 | FRANEAN1_RS16555 | 4049067 | 4049702 | 1 | transposase |
| | 4037626 | 4083572 | 45946 | IslandPath-DIMOB and SIGI-HMM | WP_020460963.1 | FRANEAN1_RS16560 | 4049877 | 4050305 | 1 | hypothetical protein |
| | 4037626 | 4083572 | 45946 | IslandPath-DIMOB and SIGI-HMM | WP_020460964.1 | FRANEAN1_RS16565 | 4050568 | 4051017 | -1 | hypothetical protein |
| | 4037626 | 4083572 | 45946 | IslandPath-DIMOB and SIGI-HMM | WP_020460965.1 | FRANEAN1_RS16570 | 4051014 | 4052702 | -1 | long-chain-fatty-acid--CoA ligase |
| | 4037626 | 4083572 | 45946 | IslandPath-DIMOB and SIGI-HMM | WP_020460966.1 | FRANEAN1_RS16575 | 4053138 | 4054232 | 1 | transposase |
| | 4037626 | 4083572 | 45946 | IslandPath-DIMOB and SIGI-HMM | WP_041254279.1 | FRANEAN1_RS16580 | 4056659 | 4056949 | -1 | hypothetical protein |
| | 4037626 | 4083572 | 45946 | IslandPath-DIMOB and SIGI-HMM | WP_041254281.1 | FRANEAN1_RS16585 | 4057354 | 4057620 | -1 | hypothetical protein |
| | 4037626 | 4083572 | 45946 | IslandPath-DIMOB | WP_020460972.1 | FRANEAN1_RS16590 | 4059514 | 4060779 | 1 | transposase |
| | 4037626 | 4083572 | 45946 | IslandPath- | WP_049795687.1 | FRANEAN1_RS16595 | 4060884 | 4061411 | -1 | hypothetical protein |

| | | | | | | | | | | |
|---|---|---|---|---|---|---|---|---|---|---|
| | | | | DIMOB | | | | | | |
| | 4037626 | 4083572 | 45946 | IslandPath-DIMOB | WP_020460974.1 | FRANEAN1_RS16600 | 4061557 | 4061805 | 1 | transposase |
| | 4037626 | 4083572 | 45946 | IslandPath-DIMOB | WP_020460977.1 | FRANEAN1_RS16610 | 4063942 | 4065843 | 1 | hypothetical protein |
| | 4037626 | 4083572 | 45946 | IslandPath-DIMOB | WP_020460978.1 | FRANEAN1_RS16615 | 4065997 | 4066383 | 1 | peptidase C15 pyroglutamyl peptidase I |
| | 4037626 | 4083572 | 45946 | IslandPath-DIMOB | WP_020460979.1 | FRANEAN1_RS16620 | 4066403 | 4068013 | 1 | hypothetical protein |
| | 4037626 | 4083572 | 45946 | IslandPath-DIMOB | WP_020458361.1 | FRANEAN1_RS16625 | 4068230 | 4069618 | -1 | transposase |
| | 4037626 | 4083572 | 45946 | IslandPath-DIMOB | | FRANEAN1_RS16630 | 4069765 | 4070127 | -1 | transposase |
| | 4037626 | 4083572 | 45946 | IslandPath-DIMOB | WP_020460981.1 | FRANEAN1_RS16635 | 4070259 | 4070747 | -1 | transposase |
| | 4037626 | 4083572 | 45946 | IslandPath-DIMOB | WP_049795688.1 | FRANEAN1_RS16640 | 4070863 | 4071195 | 1 | hypothetical protein |
| | 4037626 | 4083572 | 45946 | IslandPath-DIMOB | WP_020460983.1 | FRANEAN1_RS16645 | 4071324 | 4073876 | -1 | hypothetical protein |
| | 4037626 | 4083572 | 45946 | IslandPath-DIMOB | | FRANEAN1_RS16650 | 4074150 | 4075388 | 1 | hypothetical protein |
| | 4037626 | 4083572 | 45946 | IslandPath-DIMOB | WP_041254282.1 | FRANEAN1_RS16655 | 4076180 | 4076416 | 1 | hypothetical protein |
| | 4037626 | 4083572 | 45946 | IslandPath-DIMOB | WP_020460987.1 | FRANEAN1_RS16660 | 4076633 | 4077709 | 1 | recombinase |
| | 4037626 | 4083572 | 45946 | IslandPath-DIMOB | WP_020460988.1 | FRANEAN1_RS16665 | 4077805 | 4078125 | 1 | XRE family transcriptional regulator |
| | 4037626 | 4083572 | 45946 | IslandPath-DIMOB | WP_020460989.1 | FRANEAN1_RS16670 | 4078128 | 4079762 | 1 | hypothetical protein |
| | 4037626 | 4083572 | 45946 | IslandPath-DIMOB | WP_049795689.1 | FRANEAN1_RS16675 | 4080154 | 4080387 | -1 | hypothetical protein |
| | 4037626 | 4083572 | 45946 | IslandPath-DIMOB | WP_020460992.1 | FRANEAN1_RS37180 | 4081077 | 4081268 | 1 | hypothetical protein |
| | 4037626 | 4083572 | 45946 | IslandPath-DIMOB | WP_020460993.1 | FRANEAN1_RS16685 | 4081417 | 4081926 | 1 | molecular chaperone DnaJ |
| | 4037626 | 4083572 | 45946 | IslandPath-DIMOB | | FRANEAN1_RS16690 | 4082160 | 4083245 | 1 | HNH endonuclease |
| | 4037626 | 4083572 | 45946 | IslandPath-DIMOB | WP_020460995.1 | FRANEAN1_RS16695 | 4083267 | 4083572 | -1 | hypothetical protein |
| | 4049877 | 4057620 | 7743 | IslandPath-DIMOB | WP_020460963.1 | FRANEAN1_RS16560 | 4049877 | 4050305 | 1 | hypothetical protein |
| | 4049877 | 4057620 | 7743 | IslandPath-DIMOB | WP_020460964.1 | FRANEAN1_RS16565 | 4050568 | 4051017 | -1 | hypothetical protein |
| | 4049877 | 4057620 | 7743 | IslandPath-DIMOB | WP_020460965.1 | FRANEAN1_RS16570 | 4051014 | 4052702 | -1 | long-chain-fatty-acid--CoA ligase |
| | 4049877 | 4057620 | 7743 | IslandPath-DIMOB | WP_020460966.1 | FRANEAN1_RS16575 | 4053138 | 4054232 | 1 | transposase |

| | | | | | | | | | | |
|---|---|---|---|---|---|---|---|---|---|---|
| | 4049877 | 4057620 | 7743 | IslandPath-DIMOB | WP_041254279.1 | FRANEAN1_RS16580 | 4056659 | 4056949 | -1 | hypothetical protein |
| | 4049877 | 4057620 | 7743 | IslandPath-DIMOB | WP_041254281.1 | FRANEAN1_RS16585 | 4057354 | 4057620 | -1 | hypothetical protein |
| 2 | 4162485 | 4178361 | 15876 | IslandPath-DIMOB | WP_020461039.1 | FRANEAN1_RS16930 | 4162485 | 4162649 | 1 | 5-methyltetrahydropteroyltriglutamate--homocysteine methyltransferase |
| | 4162485 | 4178361 | 15876 | IslandPath-DIMOB | WP_020461040.1 | FRANEAN1_RS16935 | 4162734 | 4163462 | -1 | hypothetical protein |
| | 4162485 | 4178361 | 15876 | IslandPath-DIMOB | WP_020461041.1 | FRANEAN1_RS16940 | 4163485 | 4163964 | 1 | biotin carboxylase |
| | 4162485 | 4178361 | 15876 | IslandPath-DIMOB | WP_049795955.1 | FRANEAN1_RS16945 | 4164164 | 4165327 | -1 | transposase |
| | 4162485 | 4178361 | 15876 | IslandPath-DIMOB | WP_020461044.1 | FRANEAN1_RS16950 | 4165962 | 4166978 | 1 | hypothetical protein |
| | 4162485 | 4178361 | 15876 | IslandPath-DIMOB | WP_020461045.1 | FRANEAN1_RS16955 | 4166975 | 4167850 | 1 | N-acetyltransferase GCN5 |
| | 4162485 | 4178361 | 15876 | IslandPath-DIMOB | WP_020461046.1 | FRANEAN1_RS37185 | 4167903 | 4168769 | -1 | restriction endonuclease |
| | 4162485 | 4178361 | 15876 | IslandPath-DIMOB | WP_020461047.1 | FRANEAN1_RS16965 | 4169026 | 4169463 | 1 | hypothetical protein |
| | 4162485 | 4178361 | 15876 | IslandPath-DIMOB | WP_049795690.1 | FRANEAN1_RS37190 | 4169505 | 4169693 | -1 | hypothetical protein |
| | 4162485 | 4178361 | 15876 | IslandPath-DIMOB | WP_049795691.1 | FRANEAN1_RS37195 | 4169904 | 4170146 | 1 | hypothetical protein |
| | 4162485 | 4178361 | 15876 | IslandPath-DIMOB | WP_020461048.1 | FRANEAN1_RS16975 | 4170166 | 4171362 | -1 | transposase |
| | 4162485 | 4178361 | 15876 | IslandPath-DIMOB | WP_020461049.1 | FRANEAN1_RS16980 | 4171738 | 4172307 | 1 | ATPase AAA |
| | 4162485 | 4178361 | 15876 | IslandPath-DIMOB | WP_020461050.1 | FRANEAN1_RS16985 | 4172548 | 4173339 | 1 | alpha-hydroxy acid dehydrogenase |
| | 4162485 | 4178361 | 15876 | IslandPath-DIMOB | WP_041254292.1 | FRANEAN1_RS16990 | 4173460 | 4173927 | -1 | polyketide cyclase |
| | 4162485 | 4178361 | 15876 | IslandPath-DIMOB | WP_020461052.1 | FRANEAN1_RS16995 | 4174246 | 4175001 | 1 | hypothetical protein |
| | 4162485 | 4178361 | 15876 | IslandPath-DIMOB | WP_020461053.1 | FRANEAN1_RS17000 | 4175403 | 4175837 | -1 | N-acetyltransferase GCN5 |
| | 4162485 | 4178361 | 15876 | IslandPath-DIMOB | WP_020461054.1 | FRANEAN1_RS17005 | 4176055 | 4177173 | -1 | type 11 methyltransferase |
| | 4162485 | 4178361 | 15876 | IslandPath-DIMOB | WP_041254293.1 | FRANEAN1_RS17010 | 4177531 | 4178361 | -1 | aminoglycoside phosphotransferase |

**Table S87.** Predicted genomic islands nearby the abyssomicin BGC from *Herbidospora sakaeratensis* NBRC 102641 (NZ_BBXC01000032).

| **Island number** | **Island start** | **Island end** | **Length** | **Method** | **Gene name** | **Locus** | **Gene start** | **Gene end** | **Strand** | **Product** |
|---|---|---|---|---|---|---|---|---|---|---|
| 1 | 56040 | 60649 | 4609 | SIGI-HMM | WP_062343049.1 | AW271_RS37545 | 56040 | 56351 | 1 | hypothetical protein |
| | 56040 | 60649 | 4609 | SIGI-HMM | WP_062343051.1 | AW271_RS37550 | 56430 | 57275 | -1 | helix-turn-helix domain-containing protein |
| | 56040 | 60649 | 4609 | SIGI-HMM | WP_062343072.1 | AW271_RS37555 | 57419 | 58267 | 1 | SDR family NAD(P)-dependent oxidoreductase |
| | 56040 | 60649 | 4609 | SIGI-HMM | WP_062343053.1 | AW271_RS37560 | 58577 | 59458 | -1 | haloalkane dehalogenase |
| | 56040 | 60649 | 4609 | SIGI-HMM | WP_062343055.1 | AW271_RS37565 | 59571 | 59795 | -1 | 4-oxalocrotonate tautomerase family protein |
| | 56040 | 60649 | 4609 | SIGI-HMM | WP_062343058.1 | AW271_RS37570 | 60152 | 60649 | 1 | helix-turn-helix transcriptional regulator |

**Table S88.** Predicted genomic islands in the potential BGC from *Streptomyces cattleya* DSM 46488 (NC_017586.1).

| Island number | Island start | Island end | Length | Method | Gene name | Locus | Gene start | Gene end | Strand | Product |
|---|---|---|---|---|---|---|---|---|---|---|
| 1 | 118712 | 149510 | 30798 | IslandPath-DIMOB | WP_014140884.1 | SCATT_RS00500 | 118712 | 119890 | 1 | hypothetical protein |
| | 118712 | 149510 | 30798 | IslandPath-DIMOB | WP_014627221.1 | SCATT_RS00505 | 120052 | 121530 | 1 | hypothetical protein |
| | 118712 | 149510 | 30798 | IslandPath-DIMOB | WP_014140886.1 | SCATT_RS00510 | 121668 | 122774 | 1 | hypothetical protein |
| | 118712 | 149510 | 30798 | IslandPath-DIMOB | WP_014140887.1 | SCATT_RS00515 | 122863 | 123426 | 1 | GNAT family N-acetyltransferase |
| | 118712 | 149510 | 30798 | IslandPath-DIMOB | WP_014140888.1 | SCATT_RS00520 | 123516 | 124982 | 1 | phosphohydrolase |
| | 118712 | 149510 | 30798 | IslandPath-DIMOB | WP_014140889.1 | SCATT_RS00525 | 125072 | 125857 | 1 | DUF72 domain-containing protein |
| | 118712 | 149510 | 30798 | IslandPath-DIMOB | WP_014140890.1 | SCATT_RS00530 | 125974 | 126939 | 1 | D-alanyl-D-alanine carboxypeptidase |
| | 118712 | 149510 | 30798 | IslandPath-DIMOB | WP_014140891.1 | SCATT_RS00535 | 127454 | 127849 | 1 | lytic transglycosylase domain-containing protein |
| | 118712 | 149510 | 30798 | IslandPath-DIMOB | WP_014140892.1 | SCATT_RS00540 | 128044 | 130809 | 1 | SpoIIE family protein phosphatase |
| | 118712 | 149510 | 30798 | IslandPath-DIMOB | WP_086010104.1 | SCATT_RS00545 | 130879 | 131373 | -1 | DUF1360 domain-containing protein |
| | 118712 | 149510 | 30798 | IslandPath-DIMOB | WP_014140894.1 | SCATT_RS00550 | 131632 | 133032 | 1 | dihydrolipoyl dehydrogenase |
| | 118712 | 149510 | 30798 | IslandPath-DIMOB | WP_014140895.1 | SCATT_RS00555 | 134176 | 134448 | 1 | hypothetical protein |
| | 118712 | 149510 | 30798 | IslandPath-DIMOB | WP_014140896.1 | SCATT_RS00560 | 134445 | 135626 | 1 | LLM class flavin-dependent oxidoreductase |
| | 118712 | 149510 | 30798 | IslandPath-DIMOB | WP_014140897.1 | SCATT_RS00565 | 135869 | 136702 | 1 | SDR family oxidoreductase |
| | 118712 | 149510 | 30798 | IslandPath-DIMOB | WP_014140898.1 | SCATT_RS35820 | 136793 | 137098 | 1 | hypothetical protein |
| | 118712 | 149510 | 30798 | IslandPath-DIMOB | WP_014140899.1 | SCATT_RS00570 | 137239 | 138603 | 1 | crotonyl-CoA carboxylase/reductase |
| | 118712 | 149510 | 30798 | IslandPath-DIMOB | WP_014140901.1 | SCATT_RS35620 | 139045 | 139374 | 1 | transposase |
| | 118712 | 149510 | 30798 | IslandPath-DIMOB | WP_014140903.1 | SCATT_RS00580 | 139609 | 142386 | 1 | helix-turn-helix transcriptional regulator |
| | 118712 | 149510 | 30798 | IslandPath-DIMOB | WP_014627227.1 | SCATT_RS00585 | 142847 | 143074 | 1 | hypothetical protein |
| | 118712 | 149510 | 30798 | IslandPath-DIMOB | WP_014627229.1 | SCATT_RS00590 | 143554 | 144780 | -1 | cytochrome P450 |
| | 118712 | 149510 | 30798 | IslandPath-DIMOB | WP_014140907.1 | SCATT_RS00595 | 144777 | 145796 | -1 | thioesterase |
| | 118712 | 149510 | 30798 | IslandPath-DIMOB | WP_014627231.1 | SCATT_RS00600 | 145951 | 146427 | -1 | nuclear transport factor 2 family protein |
| | 118712 | 149510 | 30798 | IslandPath-DIMOB | WP_014140909.1 | SCATT_RS00605 | 146521 | 147363 | -1 | NAD(P)-dependent oxidoreductase |
| | 118712 | 149510 | 30798 | IslandPath-DIMOB | WP_014140910.1 | SCATT_RS00610 | 147412 | 147834 | -1 | hypothetical protein |
| | 118712 | 149510 | 30798 | IslandPath-DIMOB | WP_014627233.1 | SCATT_RS00615 | 147862 | 148263 | -1 | nuclear transport factor 2 family protein |

| | | | | | | | | | | |
|---|---|---|---|---|---|---|---|---|---|---|
| | 118712 | 149510 | 30798 | IslandPath-DIMOB | WP_014140912.1 | SCATT_RS00620 | 148260 | 149510 | -1 | cytochrome P450 |

**Table S89.** Predicted genomic islands nearby AbyU homolog from *Streptomyces armeniacus* ATCC 15676 (CP031320.1).

| Island number | Island start | Island end | Length | Method | Gene name | Locus | Gene start | Gene end | Strand | Product |
|---|---|---|---|---|---|---|---|---|---|---|
| 1 | 7286945 | 7297167 | 10222 | IslandPath-DIMOB | AXK36495.1 | DVA86_31895 | 7285777 | 7286970 | 1 | DUF1205 domain-containing protein |
| | 7286945 | 7297167 | 10222 | IslandPath-DIMOB | AXK36496.1 | DVA86_31900 | 7286945 | 7287814 | 1 | hypothetical protein |
| | 7286945 | 7297167 | 10222 | IslandPath-DIMOB | AXK36497.1 | DVA86_31905 | 7288161 | 7288919 | 1 | phage Gp37/Gp68 family protein |
| | 7286945 | 7297167 | 10222 | IslandPath-DIMOB | AXK36498.1 | DVA86_31910 | 7288959 | 7290155 | -1 | hypothetical protein |
| | 7286945 | 7297167 | 10222 | IslandPath-DIMOB | AXK36499.1 | DVA86_31915 | 7290475 | 7290762 | 1 | hypothetical protein |
| | 7286945 | 7297167 | 10222 | IslandPath-DIMOB | AXK36500.1 | DVA86_31920 | 7290944 | 7293595 | -1 | ATP/GTP-binding protein |
| | 7286945 | 7297167 | 10222 | IslandPath-DIMOB | AXK37815.1 | DVA86_31925 | 7293886 | 7294101 | 1 | hypothetical protein |
| | 7286945 | 7297167 | 10222 | IslandPath-DIMOB | AXK37814.1 | DVA86_31930 | 7294077 | 7295885 | -1 | ATP/GTP-binding protein |
| | 7286945 | 7297167 | 10222 | IslandPath-DIMOB | AXK36501.1 | DVA86_31935 | 7296034 | 7297167 | 1 | Fic family protein |

**Table S90.** Predicted genomic islands in potential BGC from *Streptomyces iranensis* DSM 41954 (NZ_LK022848).

| Island number | Island start | Island end | Length | Method | Gene name | Locus | Gene start | Gene end | Strand | Product |
|---|---|---|---|---|---|---|---|---|---|---|
| 1 | 11040571 | 11048595 | 8024 | IslandPick | WP_044580002.1 | SIRAN_RS44125 | 11037896 | 11040580 | 1 | glucan biosynthesis protein |
| | 11040571 | 11048595 | 8024 | IslandPick | WP_078957437.1 | SIRAN_RS51845 | 11040877 | 11041341 | -1 | nuclear transport factor 2 family protein |
| | 11040571 | 11048595 | 8024 | IslandPick | WP_078957138.1 | SIRAN_RS51850 | 11041468 | 11042301 | -1 | NAD(P)-dependent oxidoreductase |
| | 11040571 | 11048595 | 8024 | IslandPick | WP_078957139.1 | SIRAN_RS51855 | 11042451 | 11042873 | -1 | hypothetical protein |
| | 11040571 | 11048595 | 8024 | IslandPick | WP_078957140.1 | SIRAN_RS44130 | 11042902 | 11043261 | -1 | nuclear transport factor 2 family protein |
| | 11040571 | 11048595 | 8024 | IslandPick | WP_044580004.1 | SIRAN_RS44135 | 11043300 | 11044553 | -1 | cytochrome P450 |
| | 11040571 | 11048595 | 8024 | IslandPick | WP_044580005.1 | SIRAN_RS44140 | 11044641 | 11063105 | -1 | type I polyketide synthase |
| 2 | 11062737 | 11066838 | 4101 | IslandPick | WP_044580005.1 | SIRAN_RS44140 | 11044641 | 11063105 | -1 | type I polyketide synthase |
| | 11062737 | 11066838 | 4101 | IslandPick | | SIRAN_RS44145 | 11063102 | 11074674 | -1 | 3-ketoacyl-ACP synthase |
| 3 | 11072390 | 11078862 | 6472 | IslandPick | | SIRAN_RS44145 | 11063102 | 11074674 | -1 | 3-ketoacyl-ACP synthase |
| | 11072390 | 11078862 | 6472 | IslandPick | WP_078957141.1 | SIRAN_RS51860 | 11074773 | 11076665 | -1 | type I polyketide synthase |
| | 11072390 | 11078862 | 6472 | IslandPick | WP_044580008.1 | SIRAN_RS44165 | 11076907 | 11077464 | -1 | pyridoxamine 5'-phosphate oxidase family protein |
| | 11072390 | 11078862 | 6472 | IslandPick | WP_044580009.1 | SIRAN_RS44170 | 11077848 | 11089649 | 1 | type I polyketide synthase |
| 4 | 11085080 | 11090496 | 5416 | IslandPick | WP_044580009.1 | SIRAN_RS44170 | 11077848 | 11089649 | 1 | type I polyketide synthase |
| | 11085080 | 11090496 | 5416 | IslandPick | WP_044580010.1 | SIRAN_RS44175 | 11089703 | 11093776 | 1 | type I polyketide synthase |
| 5 | 11091079 | 11102579 | 11500 | IslandPick | WP_044580010.1 | SIRAN_RS44175 | 11089703 | 11093776 | 1 | type I polyketide synthase |
| | 11091079 | 11102579 | 11500 | IslandPick | WP_078957142.1 | SIRAN_RS51865 | 11093717 | 11094226 | -1 | hypothetical protein |
| | 11091079 | 11102579 | 11500 | IslandPick | WP_044580011.1 | SIRAN_RS44180 | 11094468 | 11095301 | -1 | SDR family oxidoreductase |
| | 11091079 | 11102579 | 11500 | IslandPick | WP_044580012.1 | SIRAN_RS44185 | 11095683 | 11096864 | 1 | LLM class flavin-dependent oxidoreductase |
| | 11091079 | 11102579 | 11500 | IslandPick | | SIRAN_RS44190 | 11096920 | 11097126 | 1 | thioesterase |
| | 11091079 | 11102579 | 11500 | IslandPick | WP_078957143.1 | SIRAN_RS44195 | 11097123 | 11098349 | 1 | cytochrome P450 |
| | 11091079 | 11102579 | 11500 | IslandPick | | SIRAN_RS51870 | 11098657 | 11099193 | -1 | hypothetical protein |
| | 11091079 | 11102579 | 11500 | IslandPick | WP_044580016.1 | SIRAN_RS44205 | 11099197 | 11101998 | -1 | helix-turn-helix transcriptional regulator |
| | 11091079 | 11102579 | 11500 | IslandPick | WP_107073432.1 | SIRAN_RS53370 | 11102469 | 11102720 | 1 | hypothetical protein |

**Table S91.** Predicted genomic islands nearby AbyU homolog from *Streptomyces caatingaensis* CMAA 1322 (NZ_LFXA01000017).

| Island number | Island start | Island end | Length | Method | Gene name | Locus | Gene start | Gene end | Strand | Product |
|---|---|---|---|---|---|---|---|---|---|---|
| 1 | 223990 | 272192 | 48202 | IslandPath-DIMOB | WP_049718312.1 | AC230_RS23380 | 223990 | 225192 | -1 | beta-ketoacyl-ACP synthase II |
| | 223990 | 272192 | 48202 | IslandPath-DIMOB | WP_049718313.1 | AC230_RS23385 | 225404 | 226003 | 1 | TetR/AcrR family transcriptional regulator |
| | 223990 | 272192 | 48202 | IslandPath-DIMOB | WP_049718314.1 | AC230_RS23390 | 225972 | 226367 | -1 | hypothetical protein |
| | 223990 | 272192 | 48202 | IslandPath-DIMOB | WP_049718315.1 | AC230_RS23395 | 226575 | 227609 | 1 | zinc-dependent alcohol dehydrogenase family protein |
| | 223990 | 272192 | 48202 | IslandPath-DIMOB | WP_078871521.1 | AC230_RS31430 | 227616 | 228092 | -1 | DUF4188 domain-containing protein |
| | 223990 | 272192 | 48202 | IslandPath-DIMOB | WP_049718316.1 | AC230_RS23410 | 228394 | 229050 | -1 | DUF1211 domain-containing protein |
| | 223990 | 272192 | 48202 | IslandPath-DIMOB | WP_049718317.1 | AC230_RS23415 | 229187 | 230215 | 1 | N-acetyl-gamma-glutamyl-phosphate reductase |
| | 223990 | 272192 | 48202 | IslandPath-DIMOB | WP_049718318.1 | AC230_RS23420 | 230212 | 231363 | 1 | bifunctional glutamate N-acetyltransferase/amino-acid acetyltransferase ArgJ |
| | 223990 | 272192 | 48202 | IslandPath-DIMOB | WP_049718319.1 | AC230_RS23425 | 231360 | 232268 | 1 | acetylglutamate kinase |
| | 223990 | 272192 | 48202 | IslandPath-DIMOB | WP_049718320.1 | AC230_RS23430 | 232265 | 233452 | 1 | acetylornithine transaminase |
| | 223990 | 272192 | 48202 | IslandPath-DIMOB | WP_049718321.1 | AC230_RS23435 | 233512 | 234057 | 1 | arginine repressor |
| | 223990 | 272192 | 48202 | IslandPath-DIMOB | WP_049718322.1 | AC230_RS23440 | 234137 | 235183 | 1 | hypothetical protein |
| | 223990 | 272192 | 48202 | IslandPath-DIMOB | WP_078871522.1 | AC230_RS31435 | 235122 | 235313 | -1 | DUF397 domain-containing protein |
| | 223990 | 272192 | 48202 | IslandPath-DIMOB | WP_078871523.1 | AC230_RS23445 | 235310 | 236563 | -1 | helix-turn-helix domain-containing protein |
| | 223990 | 272192 | 48202 | IslandPath-DIMOB | WP_078871524.1 | AC230_RS23455 | 236681 | 237448 | 1 | signal peptidase I |
| | 223990 | 272192 | 48202 | IslandPath-DIMOB | WP_049718325.1 | AC230_RS23460 | 238453 | 238845 | -1 | hypothetical protein |
| | 223990 | 272192 | 48202 | IslandPath-DIMOB | WP_049718326.1 | AC230_RS23465 | 238903 | 239382 | -1 | IS5/IS1182 family transposase |
| | 223990 | 272192 | 48202 | IslandPath-DIMOB | WP_049718327.1 | AC230_RS23470 | 239688 | 240203 | -1 | DUF2247 family protein |
| | 223990 | 272192 | 48202 | IslandPath-DIMOB | WP_078871525.1 | AC230_RS23475 | 240220 | 243255 | -1 | hypothetical protein |
| | 223990 | 272192 | 48202 | IslandPath-DIMOB | WP_049718328.1 | AC230_RS23480 | 243221 | 243604 | 1 | hypothetical protein |
| | 223990 | 272192 | 48202 | IslandPath-DIMOB | | AC230_RS23485 | 243611 | 243889 | -1 | GNAT family N-acetyltransferase |
| | 223990 | 272192 | 48202 | IslandPath-DIMOB | WP_049718329.1 | AC230_RS23490 | 244234 | 244626 | -1 | hypothetical protein |
| | 223990 | 272192 | 48202 | IslandPath-DIMOB | WP_078871526.1 | AC230_RS23495 | 244648 | 251268 | -1 | sugar-binding protein |
| | 223990 | 272192 | 48202 | IslandPath-DIMOB | WP_078871587.1 | AC230_RS23500 | 251663 | 255541 | -1 | LamG domain-containing protein |

| | | | | | | | | | | |
|---|---|---|---|---|---|---|---|---|---|---|
| | 223990 | 272192 | 48202 | IslandPath-DIMOB | WP_049718331.1 | AC230_RS23505 | 256202 | 256687 | 1 | MarR family transcriptional regulator |
| | 223990 | 272192 | 48202 | IslandPath-DIMOB | WP_049718332.1 | AC230_RS23510 | 256705 | 257391 | -1 | respiratory nitrate reductase subunit gamma |
| | 223990 | 272192 | 48202 | IslandPath-DIMOB | WP_049718333.1 | AC230_RS23515 | 257408 | 257986 | -1 | nitrate reductase molybdenum cofactor assembly chaperone |
| | 223990 | 272192 | 48202 | IslandPath-DIMOB | WP_078871588.1 | AC230_RS23520 | 257983 | 259566 | -1 | nitrate reductase subunit beta |
| | 223990 | 272192 | 48202 | IslandPath-DIMOB | | AC230_RS23525 | 259577 | 263262 | -1 | nitrate reductase subunit alpha |
| | 223990 | 272192 | 48202 | IslandPath-DIMOB | WP_049718334.1 | AC230_RS23530 | 264241 | 266856 | 1 | M4 family peptidase |
| | 223990 | 272192 | 48202 | IslandPath-DIMOB | WP_049718335.1 | AC230_RS23535 | 267031 | 268311 | 1 | DegT/DnrJ/EryC1/StrS family aminotransferase |
| | 223990 | 272192 | 48202 | IslandPath-DIMOB | WP_053161355.1 | AC230_RS30460 | 268308 | 269315 | 1 | hypothetical protein |
| | 223990 | 272192 | 48202 | IslandPath-DIMOB | WP_049718851.1 | AC230_RS23545 | 269404 | 269637 | 1 | hypothetical protein |
| | 223990 | 272192 | 48202 | IslandPath-DIMOB | WP_049718336.1 | AC230_RS23550 | 269634 | 271289 | 1 | (2,3-dihydroxybenzoyl)adenylate synthase |
| | 223990 | 272192 | 48202 | IslandPath-DIMOB | WP_049718337.1 | AC230_RS23555 | 271401 | 272192 | 1 | thioesterase |
| | 223990 | 272192 | 48202 | IslandPath-DIMOB | WP_049718338.1 | AC230_RS23560 | 272189 | 273382 | 1 | FAD-dependent oxidoreductase |

**Table S92.** Predicted genomic islands in potential abyssomicin BGC from *Streptomyces* sp. SCA2-2 (NZ_PKMX01000004 and NZ_PKMX01000005).

| Island number | Island start | Island end | Length | Method | Gene name | Locus | Gene start | Gene end | Strand | Product |
|---|---|---|---|---|---|---|---|---|---|---|
| 1 | 3631 | 8735 | 5104 | SIGI-HMM | WP_129847664.1 | C0L86_RS05545 | 2810 | 3634 | -1 | thioesterase |
| | 3631 | 8735 | 5104 | SIGI-HMM | WP_129847665.1 | C0L86_RS05550 | 3631 | 4752 | -1 | alpha/beta hydrolase |
| | 3631 | 8735 | 5104 | SIGI-HMM | WP_129847666.1 | C0L86_RS05555 | 4749 | 5591 | -1 | acyltransferase |
| | 3631 | 8735 | 5104 | SIGI-HMM | WP_129847667.1 | C0L86_RS05560 | 5588 | 5815 | -1 | acyl carrier protein |
| | 3631 | 8735 | 5104 | SIGI-HMM | WP_129847668.1 | C0L86_RS05565 | 5812 | 7698 | -1 | HAD-IIIC family phosphatase |
| | 3631 | 8735 | 5104 | SIGI-HMM | WP_129847669.1 | C0L86_RS05570 | 7704 | 8735 | -1 | 3-oxoacyl-ACP synthase III family protein |

**Table S93.** Predicted genomic islands in abyssomicin BGC from *Streptomyces koyangensis* SCSIO 5802 (MG243704).

| Island number | Island start | Island end | Length | Method | Gene name | Gene start | Gene end | Strand | Product |
|---|---|---|---|---|---|---|---|---|---|
| 1 | 20783 | 25887 | 5104 | SIGI-HMM | AVI57426.1 | 20783 | 21814 | 1 | AbmA1 |
| | 20783 | 25887 | 5104 | SIGI-HMM | AVI57427.1 | 21820 | 23706 | 1 | AbmA2 |
| | 20783 | 25887 | 5104 | SIGI-HMM | AVI57428.1 | 23703 | 23930 | 1 | AbmA3 |
| | 20783 | 25887 | 5104 | SIGI-HMM | AVI57429.1 | 23927 | 24769 | 1 | AbmA4 |
| | 20783 | 25887 | 5104 | SIGI-HMM | AVI57430.1 | 24766 | 25887 | 1 | AbmA5 |
| | 20783 | 25887 | 5104 | SIGI-HMM | AVI57431.1 | 25884 | 26708 | 1 | AbmT |

**Table S94.** Predicted genomic islands in potential abyssomicin BGC from *Streptomyces griseorubiginosus* SAI-142 (NZ_RJKZ01000001.1).

| Island number | Island start | Island end | Length | Method | Gene name | Locus | Gene start | Gene end | Strand | Product |
|---|---|---|---|---|---|---|---|---|---|---|
| 1 | 6603102 | 6609081 | 5979 | IslandPick | WP_123763198.1 | EDC83_RS30455 | 6598078 | 6604542 | 1 | hypothetical protein |
| | 6603102 | 6609081 | 5979 | IslandPick | WP_123763199.1 | EDC83_RS30460 | 6604545 | 6604937 | 1 | hypothetical protein |
| | 6603102 | 6609081 | 5979 | IslandPick | WP_123765188.1 | EDC83_RS30465 | 6604975 | 6605424 | 1 | hypothetical protein |
| | 6603102 | 6609081 | 5979 | IslandPick | WP_123763200.1 | EDC83_RS30470 | 6605491 | 6607155 | 1 | hypothetical protein |
| | 6603102 | 6609081 | 5979 | IslandPick | WP_123763201.1 | EDC83_RS30475 | 6607568 | 6608059 | 1 | hypothetical protein |
| | 6603102 | 6609081 | 5979 | IslandPick | WP_123763202.1 | EDC83_RS30480 | 6608210 | 6608725 | 1 | hypothetical protein |
| 2 | 6609385 | 6614640 | 5255 | IslandPick | WP_123763203.1 | EDC83_RS30485 | 6609521 | 6612259 | 1 | AAA family ATPase |
| | 6609385 | 6614640 | 5255 | IslandPick | WP_123763204.1 | EDC83_RS30490 | 6612705 | 6613478 | 1 | AfsR/SARP family transcriptional regulator |
| | 6609385 | 6614640 | 5255 | IslandPick | WP_123763205.1 | EDC83_RS30495 | 6613527 | 6614363 | -1 | thioesterase |
| | 6609385 | 6614640 | 5255 | IslandPick | WP_123763206.1 | EDC83_RS30500 | 6614604 | 6615635 | 1 | 3-oxoacyl-ACP synthase III family protein |
| 3 | 6650159 | 6655065 | 4906 | IslandPick | WP_123763213.1 | EDC83_RS30540 | 6646381 | 6651045 | 1 | acyltransferase domain-containing protein |
| | 6650159 | 6655065 | 4906 | IslandPick | WP_123763214.1 | EDC83_RS30545 | 6650954 | 6652699 | 1 | hypothetical protein |
| | 6650159 | 6655065 | 4906 | IslandPick | WP_123763215.1 | EDC83_RS30550 | 6652748 | 6652933 | 1 | hypothetical protein |
| | 6650159 | 6655065 | 4906 | IslandPick | WP_123763216.1 | EDC83_RS30555 | 6653032 | 6654078 | 1 | methyltransferase |
| | 6650159 | 6655065 | 4906 | IslandPick | WP_123763217.1 | EDC83_RS30560 | 6654139 | 6654552 | -1 | hypothetical protein |
| | 6650159 | 6655065 | 4906 | IslandPick | WP_123763218.1 | EDC83_RS30565 | 6654713 | 6654949 | -1 | ferredoxin |
| | 6650159 | 6655065 | 4906 | IslandPick | WP_123765190.1 | EDC83_RS30570 | 6654943 | 6656052 | -1 | cytochrome P450 |

**Table S95.** Predicted genomic islands in potential tetronomycin BGC from *Streptomyces olindensis* DAUFPE 5622 (JJOH01000019.1).

| Island number | Island start | Island end | Length | Method | Gene name | Locus | Gene start | Gene end | Strand | Product |
|---|---|---|---|---|---|---|---|---|---|---|
| 1 | 797257 | 801732 | 4475 | SIGI-HMM | KDN76185.1 | DF19_21910 | 797257 | 798648 | -1 | enterotoxin |
| | 797257 | 801732 | 4475 | SIGI-HMM | KDN76186.1 | DF19_21915 | 798882 | 799103 | 1 | hypothetical protein |
| | 797257 | 801732 | 4475 | SIGI-HMM | KDN76187.1 | DF19_21920 | 799389 | 800591 | 1 | cytochrome P450 |
| | 797257 | 801732 | 4475 | SIGI-HMM | KDN76188.1 | DF19_21925 | 800704 | 801732 | 1 | 3-oxoacyl-ACP synthase |

**Table S96.** Predicted genomic islands in potential BGC from *Streptomyces sp.* E5N91 SAI-083 (NZ_RJKF01000001.1).

| Island number | Island start | Island end | Length | Method | Gene name | Locus | Gene start | Gene end | Strand | Product |
|---|---|---|---|---|---|---|---|---|---|---|
| 1 | 7571262 | 7575300 | 4038 | IslandPick | WP_123627566.1 | EDC84_RS34290 | 7570858 | 7571283 | 1 | CBS domain-containing protein |
| | 7571262 | 7575300 | 4038 | IslandPick | WP_123627567.1 | EDC84_RS34295 | 7571341 | 7572360 | 1 | sigma-70 family RNA polymerase sigma factor |
| | 7571262 | 7575300 | 4038 | IslandPick | WP_123627568.1 | EDC84_RS34300 | 7572389 | 7573828 | -1 | FAD-dependent oxidoreductase |
| | 7571262 | 7575300 | 4038 | IslandPick | WP_123627569.1 | EDC84_RS34305 | 7573843 | 7575081 | -1 | polysaccharide pyruvyl transferase |
| | 7571262 | 7575300 | 4038 | IslandPick | WP_123627570.1 | EDC84_RS34310 | 7575238 | 7576188 | 1 | TerC family protein |
| 2 | 7585854 | 7591889 | 6035 | IslandPick | WP_123627580.1 | EDC84_RS34370 | 7584925 | 7585890 | 1 | pirin family protein |
| | 7585854 | 7591889 | 6035 | IslandPick | WP_123627581.1 | EDC84_RS34375 | 7586121 | 7586444 | 1 | transposase |
| | 7585854 | 7591889 | 6035 | IslandPick | WP_123627582.1 | EDC84_RS34380 | 7586562 | 7588097 | -1 | alpha/beta hydrolase |
| | 7585854 | 7591889 | 6035 | IslandPick | | EDC84_RS34385 | 7588336 | 7589144 | 1 | MerR family transcriptional regulator |
| | 7585854 | 7591889 | 6035 | IslandPick | | EDC84_RS34390 | 7589215 | 7589752 | -1 | IS701 family transposase |
| | 7585854 | 7591889 | 6035 | IslandPick | WP_123627583.1 | EDC84_RS34395 | 7589925 | 7590770 | -1 | methyltransferase domain-containing protein |
| | 7585854 | 7591889 | 6035 | IslandPick | WP_123627584.1 | EDC84_RS34400 | 7590827 | 7594921 | -1 | acyltransferase domain-containing protein |
| 3 | 7607165 | 7620916 | 13751 | SIGI-HMM | WP_123627586.1 | EDC84_RS34410 | 7607165 | 7607695 | 1 | pyridoxamine 5'-phosphate oxidase family protein |
| | 7607165 | 7620916 | 13751 | SIGI-HMM | WP_123627587.1 | EDC84_RS34415 | 7607786 | 7620916 | 1 | SDR family NAD(P)-dependent oxidoreductase |
| | 7607165 | 7620916 | 13751 | SIGI-HMM | WP_123627588.1 | EDC84_RS34420 | 7620913 | 7639290 | 1 | SDR family NAD(P)-dependent oxidoreductase |
| 4 | 7638150 | 7646034 | 7884 | SIGI-HMM and IslandPick | WP_123627588.1 | EDC84_RS34420 | 7620913 | 7639290 | 1 | SDR family NAD(P)-dependent oxidoreductase |
| | 7638150 | 7646034 | 7884 | SIGI-HMM and IslandPick | WP_123627589.1 | EDC84_RS34425 | 7639359 | 7640618 | 1 | cytochrome P450 |
| | 7638150 | 7646034 | 7884 | SIGI-HMM and IslandPick | WP_123627590.1 | EDC84_RS34430 | 7640615 | 7641016 | 1 | nuclear transport factor 2 family protein |
| | 7638150 | 7646034 | 7884 | SIGI-HMM and IslandPick | WP_123627591.1 | EDC84_RS34435 | 7641045 | 7641467 | 1 | hypothetical protein |
| | 7638150 | 7646034 | 7884 | SIGI-HMM and IslandPick | WP_123627592.1 | EDC84_RS34440 | 7641572 | 7642414 | 1 | NAD(P)-dependent oxidoreductase |
| | 7638150 | 7646034 | 7884 | SIGI-HMM and IslandPick | WP_123627593.1 | EDC84_RS34445 | 7642507 | 7642980 | 1 | nuclear transport factor 2 family protein |
| | 7638150 | 7646034 | 7884 | SIGI-HMM and IslandPick | | EDC84_RS34450 | 7643041 | 7644037 | 1 | thioesterase |
| | 7638150 | 7646034 | 7884 | SIGI-HMM and IslandPick | WP_123627594.1 | EDC84_RS34455 | 7644034 | 7645260 | 1 | cytochrome P450 |

| | | | | | | | | | | |
|---|---|---|---|---|---|---|---|---|---|---|
| | 7638150 | 7646034 | 7884 | IslandPick | WP_123627595.1 | EDC84_RS34460 | 7645913 | 7647094 | 1 | LLM class flavin-dependent oxidoreductase |
| 5 | 7640615 | 7645260 | 4645 | IslandPick | WP_123627589.1 | EDC84_RS34425 | 7639359 | 7640618 | 1 | cytochrome P450 |
| | 7640615 | 7645260 | 4645 | IslandPick | WP_123627590.1 | EDC84_RS34430 | 7640615 | 7641016 | 1 | nuclear transport factor 2 family protein |
| | 7640615 | 7645260 | 4645 | IslandPick | WP_123627591.1 | EDC84_RS34435 | 7641045 | 7641467 | 1 | hypothetical protein |
| | 7640615 | 7645260 | 4645 | IslandPick | WP_123627592.1 | EDC84_RS34440 | 7641572 | 7642414 | 1 | NAD(P)-dependent oxidoreductase |
| | 7640615 | 7645260 | 4645 | IslandPick | WP_123627593.1 | EDC84_RS34445 | 7642507 | 7642980 | 1 | nuclear transport factor 2 family protein |
| | 7640615 | 7645260 | 4645 | IslandPick | | EDC84_RS34450 | 7643041 | 7644037 | 1 | thioesterase |
| | 7640615 | 7645260 | 4645 | IslandPick | WP_123627594.1 | EDC84_RS34455 | 7644034 | 7645260 | 1 | cytochrome P450 |
| 6 | 7649811 | 7657319 | 7508 | IslandPick | WP_123627597.1 | EDC84_RS34475 | 7649079 | 7650443 | 1 | crotonyl-CoA carboxylase/reductase |
| | 7649811 | 7657319 | 7508 | IslandPick | WP_123628731.1 | EDC84_RS34480 | 7651139 | 7653916 | 1 | helix-turn-helix transcriptional regulator |
| | 7649811 | 7657319 | 7508 | IslandPick | | EDC84_RS34485 | 7653975 | 7654518 | -1 | IS5/IS1182 family transposase |
| | 7649811 | 7657319 | 7508 | IslandPick | | EDC84_RS34490 | 7654686 | 7655182 | -1 | ISAzo13 family transposase |
| | 7649811 | 7657319 | 7508 | IslandPick | WP_123627598.1 | EDC84_RS34495 | 7655281 | 7656108 | -1 | endo alpha-1,4 polygalactosaminidase |
| | 7649811 | 7657319 | 7508 | IslandPick | WP_123627599.1 | EDC84_RS34500 | 7656863 | 7657768 | 1 | class A beta-lactamase |
| 7 | 7657633 | 7661732 | 4099 | IslandPick | WP_123627599.1 | EDC84_RS34500 | 7656863 | 7657768 | 1 | class A beta-lactamase |
| | 7657633 | 7661732 | 4099 | IslandPick | WP_123627600.1 | EDC84_RS34505 | 7657765 | 7658109 | 1 | transposase |
| | 7657633 | 7661732 | 4099 | IslandPick | WP_123627601.1 | EDC84_RS34510 | 7658288 | 7658944 | 1 | transposase |
| | 7657633 | 7661732 | 4099 | IslandPick | | EDC84_RS34515 | 7659020 | 7659535 | 1 | NAD(P)-dependent alcohol dehydrogenase |
| | 7657633 | 7661732 | 4099 | IslandPick | WP_123627602.1 | EDC84_RS34520 | 7659701 | 7660423 | 1 | alpha/beta hydrolase |
| | 7657633 | 7661732 | 4099 | IslandPick | WP_123627603.1 | EDC84_RS34525 | 7660579 | 7661721 | 1 | Gfo/Idh/MocA family oxidoreductase |
| | 7657633 | 7661732 | 4099 | IslandPick | | EDC84_RS34530 | 7661725 | 7662434 | -1 | IS5/IS1182 family transposase |

**Table S97.** Predicted genomic islands in potential BGC from *Streptomyces olivaceus* KLBMP 5084 (NZ_CP016795.1).

| Island number | Island start | Island end | Length | Method | Gene name | Locus | Gene start | Gene end | Strand | Product |
|---|---|---|---|---|---|---|---|---|---|---|
| 1 | 7944586 | 7951229 | 6643 | Island Pick | WP_070390064.1 | BC342_RS34420 | 7943086 | 7945233 | -1 | MMPL family transporter |
| | 7944586 | 7951229 | 6643 | Island Pick | WP_037769115.1 | BC342_RS34425 | 7945383 | 7945961 | 1 | TetR/AcrR family transcriptional regulator |
| | 7944586 | 7951229 | 6643 | Island Pick | WP_079155146.1 | BC342_RS34430 | 7946091 | 7949228 | 1 | AAA family ATPase |
| | 7944586 | 7951229 | 6643 | Island Pick | WP_070390547.1 | BC342_RS34435 | 7949283 | 7950356 | -1 | alcohol dehydrogenase |
| | 7944586 | 7951229 | 6643 | Island Pick | | BC342_RS36420 | 7950486 | 7950671 | 1 | siderophore-interacting protein |
| | 7944586 | 7951229 | 6643 | Island Pick | WP_079155147.1 | BC342_RS34440 | 7950767 | 7952413 | 1 | serine/threonine protein kinase |
| 2 | 7979896 | 8005952 | 26056 | SIGI-HMM | | BC342_RS34550 | 7979896 | 7991707 | -1 | SDR family NAD(P)-dependent oxidoreductase |
| | 7979896 | 8005952 | 26056 | SIGI-HMM | WP_070390076.1 | BC342_RS34555 | 7992082 | 7992603 | 1 | pyridoxamine 5'-phosphate oxidase family protein |
| | 7979896 | 8005952 | 26056 | SIGI-HMM | | BC342_RS34560 | 7992716 | 8005952 | 1 | SDR family NAD(P)-dependent oxidoreductase |
| | 7979896 | 8005952 | 26056 | SIGI-HMM | WP_070390077.1 | BC342_RS34565 | 8005949 | 8024335 | 1 | type I polyketide synthase |
| 3 | 8023417 | 8032406 | 8989 | Island Pick, SIGI-HMM and IslandPath-DIMOB | WP_070390077.1 | BC342_RS34565 | 8005949 | 8024335 | 1 | type I polyketide synthase |
| | 8023417 | 8032406 | 8989 | Island Pick, SIGI-HMM and IslandPath-DIMOB | WP_037772595.1 | BC342_RS34570 | 8024403 | 8025653 | 1 | cytochrome P450 |
| | 8023417 | 8032406 | 8989 | Island Pick, SIGI-HMM and IslandPath-DIMOB | WP_070390078.1 | BC342_RS34575 | 8025650 | 8026051 | 1 | nuclear transport factor 2 family protein |
| | 8023417 | 8032406 | 8989 | Island Pick, SIGI-HMM and IslandPath-DIMOB | WP_037772597.1 | BC342_RS34580 | 8026080 | 8026502 | 1 | hypothetical protein |
| | 8023417 | 8032406 | 8989 | Island Pick, SIGI-HMM and IslandPath-DIMOB | WP_070390079.1 | BC342_RS34585 | 8026607 | 8027449 | 1 | NAD(P)-dependent oxidoreductase |
| | 8023417 | 8032406 | 8989 | Island Pick, SIGI-HMM and IslandPath-DIMOB | WP_037772601.1 | BC342_RS34590 | 8027542 | 8028015 | 1 | nuclear transport factor 2 family protein |
| | 8023417 | 8032406 | 8989 | Island Pick, SIGI-HMM and IslandPath-DIMOB | | BC342_RS34595 | 8028096 | 8029092 | 1 | thioesterase |
| | 8023417 | 8032406 | 8989 | Island Pick, SIGI- | WP_078536178.1 | BC342_RS34600 | 8029089 | 8030315 | 1 | cytochrome P450 |

| | | | | | | | | | | |
|---|---|---|---|---|---|---|---|---|---|---|
| | | | | HMM and IslandPath-DIMOB | | | | | | |
| | 8023417 | 8032406 | 8989 | Island Pick, SIGI-HMM and IslandPath-DIMOB | WP_123937984.1 | BC342_RS36425 | 8030672 | 8030971 | 1 | hypothetical protein |
| | 8023417 | 8032406 | 8989 | Island Pick, SIGI-HMM and IslandPath-DIMOB | WP_031047091.1 | BC342_RS34605 | 8030968 | 8032149 | 1 | LLM class flavin-dependent oxidoreductase |
| 4 | 8025650 | 8172811 | 147161 | Island Pick and IslandPath-DIMOB | WP_037772595.1 | BC342_RS34570 | 8024403 | 8025653 | 1 | cytochrome P450 |
| | 8025650 | 8172811 | 147161 | Island Pick and IslandPath-DIMOB | WP_070390078.1 | BC342_RS34575 | 8025650 | 8026051 | 1 | nuclear transport factor 2 family protein |
| | 8025650 | 8172811 | 147161 | Island Pick and IslandPath-DIMOB | WP_037772597.1 | BC342_RS34580 | 8026080 | 8026502 | 1 | hypothetical protein |
| | 8025650 | 8172811 | 147161 | Island Pick and IslandPath-DIMOB | WP_070390079.1 | BC342_RS34585 | 8026607 | 8027449 | 1 | NAD(P)-dependent oxidoreductase |
| | 8025650 | 8172811 | 147161 | Island Pick and IslandPath-DIMOB | WP_037772601.1 | BC342_RS34590 | 8027542 | 8028015 | 1 | nuclear transport factor 2 family protein |
| | 8025650 | 8172811 | 147161 | Island Pick and IslandPath-DIMOB | | BC342_RS34595 | 8028096 | 8029092 | 1 | thioesterase |
| | 8025650 | 8172811 | 147161 | Island Pick and IslandPath-DIMOB | WP_078536178.1 | BC342_RS34600 | 8029089 | 8030315 | 1 | cytochrome P450 |
| | 8025650 | 8172811 | 147161 | Island Pick and IslandPath-DIMOB | WP_123937984.1 | BC342_RS36425 | 8030672 | 8030971 | 1 | hypothetical protein |
| | 8025650 | 8172811 | 147161 | Island Pick and IslandPath-DIMOB | WP_031047091.1 | BC342_RS34605 | 8030968 | 8032149 | 1 | LLM class flavin-dependent oxidoreductase |
| | 8025650 | 8172811 | 147161 | Island Pick and IslandPath-DIMOB | WP_031047088.1 | BC342_RS34610 | 8032407 | 8033240 | 1 | SDR family oxidoreductase |
| | 8025650 | 8172811 | 147161 | Island Pick and IslandPath-DIMOB | | BC342_RS34615 | 8033502 | 8033938 | 1 | hypothetical protein |
| | 8025650 | 8172811 | 147161 | Island Pick and IslandPath-DIMOB | WP_070390081.1 | BC342_RS34620 | 8034080 | 8035444 | 1 | crotonyl-CoA carboxylase/reductase |
| | 8025650 | 8172811 | 147161 | Island Pick and | | BC342_RS34625 | 8036200 | 8038975 | 1 | helix-turn-helix transcriptional |

| | | | | | | | | | | |
|---|---|---|---|---|---|---|---|---|---|---|
| | | | | IslandPath-DIMOB | | | | | | regulator |
| | 8025650 | 8172811 | 147161 | Island Pick and IslandPath-DIMOB | WP_070390082.1 | BC342_RS34630 | 8039390 | 8040859 | 1 | SAVED domain-containing protein |
| | 8025650 | 8172811 | 147161 | Island Pick and IslandPath-DIMOB | WP_070390083.1 | BC342_RS34635 | 8041289 | 8042275 | -1 | hypothetical protein |
| | 8025650 | 8172811 | 147161 | Island Pick and IslandPath-DIMOB | WP_070387189.1 | BC342_RS34640 | 8042389 | 8043714 | 1 | ISL3 family transposase |
| | 8025650 | 8172811 | 147161 | Island Pick and IslandPath-DIMOB | WP_070387188.1 | BC342_RS34645 | 8045212 | 8046216 | 1 | nucleotidyltransferase |
| | 8025650 | 8172811 | 147161 | Island Pick and IslandPath-DIMOB | WP_070387187.1 | BC342_RS34650 | 8046213 | 8047679 | 1 | ThiF family adenylyltransferase |
| | 8025650 | 8172811 | 147161 | Island Pick and IslandPath-DIMOB | | BC342_RS36430 | 8047676 | 8048203 | 1 | hypothetical protein |
| | 8025650 | 8172811 | 147161 | Island Pick and IslandPath-DIMOB | | BC342_RS36435 | 8048298 | 8048636 | -1 | hypothetical protein |
| | 8025650 | 8172811 | 147161 | Island Pick and IslandPath-DIMOB | WP_070387186.1 | BC342_RS34655 | 8048719 | 8049045 | -1 | hypothetical protein |
| | 8025650 | 8172811 | 147161 | Island Pick and IslandPath-DIMOB | WP_070390084.1 | BC342_RS34660 | 8049152 | 8050225 | -1 | ATP/GTP-binding protein |
| | 8025650 | 8172811 | 147161 | Island Pick and IslandPath-DIMOB | WP_107405374.1 | BC342_RS34665 | 8050225 | 8052336 | -1 | DDE-type integrase/transposase/recombinase |
| | 8025650 | 8172811 | 147161 | Island Pick and IslandPath-DIMOB | WP_070390551.1 | BC342_RS34670 | 8052618 | 8053325 | -1 | hypothetical protein |
| | 8025650 | 8172811 | 147161 | Island Pick and IslandPath-DIMOB | | BC342_RS34675 | 8053536 | 8054456 | -1 | hypothetical protein |
| | 8025650 | 8172811 | 147161 | Island Pick and IslandPath-DIMOB | WP_123937986.1 | BC342_RS34680 | 8054469 | 8058035 | -1 | DNA-binding protein |
| | 8025650 | 8172811 | 147161 | Island Pick and IslandPath-DIMOB | WP_070390552.1 | BC342_RS34685 | 8058051 | 8059061 | -1 | AAA family ATPase |
| | 8025650 | 8172811 | 147161 | Island Pick and IslandPath-DIMOB | | BC342_RS36440 | 8059103 | 8061270 | -1 | transposase |
| | 8025650 | 8172811 | 147161 | Island Pick and | WP_107405229.1 | BC342_RS34700 | 8061267 | 8062250 | -1 | TnsA-like heteromeric transposase |

| | | | | | | | | | | |
|---|---|---|---|---|---|---|---|---|---|---|
| | | | | IslandPath-DIMOB | | | | | | endonuclease subunit |
| | 8025650 | 8172811 | 147161 | Island Pick and IslandPath-DIMOB | WP_079155150.1 | BC342_RS34705 | 8063902 | 8064456 | -1 | GNAT family N-acetyltransferase |
| | 8025650 | 8172811 | 147161 | Island Pick and IslandPath-DIMOB | WP_070390090.1 | BC342_RS34710 | 8064890 | 8065825 | 1 | hypothetical protein |
| | 8025650 | 8172811 | 147161 | Island Pick and IslandPath-DIMOB | WP_070390554.1 | BC342_RS34715 | 8066765 | 8067721 | 1 | IS481 family transposase |
| | 8025650 | 8172811 | 147161 | Island Pick and IslandPath-DIMOB | WP_079155151.1 | BC342_RS34720 | 8067652 | 8070510 | 1 | serine/threonine protein kinase |
| | 8025650 | 8172811 | 147161 | Island Pick and IslandPath-DIMOB | WP_070390092.1 | BC342_RS34725 | 8070565 | 8071245 | 1 | hypothetical protein |
| | 8025650 | 8172811 | 147161 | Island Pick and IslandPath-DIMOB | WP_070390093.1 | BC342_RS34730 | 8071242 | 8073329 | 1 | hypothetical protein |
| | 8025650 | 8172811 | 147161 | Island Pick and IslandPath-DIMOB | WP_070390094.1 | BC342_RS34735 | 8073379 | 8075418 | 1 | hypothetical protein |
| | 8025650 | 8172811 | 147161 | Island Pick and IslandPath-DIMOB | WP_079155152.1 | BC342_RS34740 | 8075506 | 8077569 | -1 | helicase |
| | 8025650 | 8172811 | 147161 | Island Pick and IslandPath-DIMOB | WP_070390555.1 | BC342_RS34745 | 8078860 | 8081727 | 1 | CRISPR-associated endonuclease Cas3" |
| | 8025650 | 8172811 | 147161 | Island Pick and IslandPath-DIMOB | WP_070389195.1 | BC342_RS34750 | 8082043 | 8083308 | -1 | IS701 family transposase |
| | 8025650 | 8172811 | 147161 | Island Pick and IslandPath-DIMOB | WP_079155248.1 | BC342_RS34755 | 8083539 | 8085026 | 1 | type I-E CRISPR-associated protein Cse1/CasA |
| | 8025650 | 8172811 | 147161 | Island Pick and IslandPath-DIMOB | WP_051159998.1 | BC342_RS34760 | 8085026 | 8085670 | 1 | hypothetical protein |
| | 8025650 | 8172811 | 147161 | Island Pick and IslandPath-DIMOB | WP_037772151.1 | BC342_RS34765 | 8085888 | 8087051 | 1 | type I-E CRISPR-associated protein Cas7/Cse4/CasC |
| | 8025650 | 8172811 | 147161 | Island Pick and IslandPath-DIMOB | WP_037772153.1 | BC342_RS34770 | 8087048 | 8087857 | 1 | type I-E CRISPR-associated protein Cas5/CasD |
| | 8025650 | 8172811 | 147161 | Island Pick and IslandPath-DIMOB | WP_070390095.1 | BC342_RS34775 | 8087854 | 8088498 | 1 | type I-E CRISPR-associated protein Cas6/Cse3/CasE |
| | 8025650 | 8172811 | 147161 | Island Pick and | WP_063741599.1 | BC342_RS34780 | 8088524 | 8090083 | 1 | DDE-type |

| | | | | | | | | | | |
|---|---|---|---|---|---|---|---|---|---|---|
| | | | | IslandPath-DIMOB | | | | | | integrase/transposase/recombinase |
| | 8025650 | 8172811 | 147161 | Island Pick and IslandPath-DIMOB | WP_123937988.1 | BC342_RS34785 | 8090080 | 8090889 | 1 | ATP-binding protein |
| | 8025650 | 8172811 | 147161 | Island Pick and IslandPath-DIMOB | WP_037772156.1 | BC342_RS34790 | 8091277 | 8092737 | 1 | glycosyl hydrolase |
| | 8025650 | 8172811 | 147161 | Island Pick and IslandPath-DIMOB | WP_037772159.1 | BC342_RS34795 | 8092741 | 8093640 | -1 | inorganic polyphosphate kinase |
| | 8025650 | 8172811 | 147161 | Island Pick and IslandPath-DIMOB | WP_037772162.1 | BC342_RS34800 | 8093637 | 8094647 | -1 | SPFH/Band 7/PHB domain protein |
| | 8025650 | 8172811 | 147161 | Island Pick and IslandPath-DIMOB | WP_070390096.1 | BC342_RS34805 | 8095178 | 8096605 | -1 | hypothetical protein |
| | 8025650 | 8172811 | 147161 | Island Pick and IslandPath-DIMOB | WP_070390097.1 | BC342_RS34810 | 8097039 | 8098148 | -1 | winged helix-turn-helix domain-containing protein |
| | 8025650 | 8172811 | 147161 | Island Pick and IslandPath-DIMOB | WP_070390098.1 | BC342_RS34815 | 8098145 | 8099434 | -1 | MFS transporter |
| | 8025650 | 8172811 | 147161 | Island Pick and IslandPath-DIMOB | WP_070390099.1 | BC342_RS34820 | 8099431 | 8100117 | -1 | hypothetical protein |
| | 8025650 | 8172811 | 147161 | Island Pick and IslandPath-DIMOB | | BC342_RS34825 | 8100225 | 8101275 | -1 | YncE family protein |
| | 8025650 | 8172811 | 147161 | Island Pick and IslandPath-DIMOB | WP_079155153.1 | BC342_RS34830 | 8101345 | 8102037 | -1 | hypothetical protein |
| | 8025650 | 8172811 | 147161 | Island Pick and IslandPath-DIMOB | WP_070390101.1 | BC342_RS34835 | 8102475 | 8102912 | 1 | glyoxalase |
| | 8025650 | 8172811 | 147161 | Island Pick and IslandPath-DIMOB | WP_070390102.1 | BC342_RS34840 | 8103305 | 8104564 | -1 | hypothetical protein |
| | 8025650 | 8172811 | 147161 | Island Pick and IslandPath-DIMOB | WP_079155154.1 | BC342_RS34845 | 8104920 | 8108648 | 1 | LamG domain-containing protein |
| | 8025650 | 8172811 | 147161 | Island Pick and IslandPath-DIMOB | | BC342_RS34850 | 8108777 | 8114473 | 1 | RHS repeat-associated core domain-containing protein |
| | 8025650 | 8172811 | 147161 | Island Pick and IslandPath-DIMOB | WP_079155155.1 | BC342_RS36455 | 8116473 | 8116682 | 1 | DUF4291 family protein |
| | 8025650 | 8172811 | 147161 | Island Pick and | WP_070390104.1 | BC342_RS37465 | 8116689 | 8117315 | -1 | hypothetical protein |

| | | | | | | | | | | |
|---|---|---|---|---|---|---|---|---|---|---|
| | | | | IslandPath-DIMOB | | | | | | |
| | 8025650 | 8172811 | 147161 | Island Pick and IslandPath-DIMOB | | BC342_RS36470 | 8117378 | 8118030 | 1 | hypothetical protein |
| | 8025650 | 8172811 | 147161 | Island Pick and IslandPath-DIMOB | | BC342_RS34865 | 8118066 | 8118745 | 1 | hypothetical protein |
| | 8025650 | 8172811 | 147161 | Island Pick and IslandPath-DIMOB | | BC342_RS36475 | 8118783 | 8119242 | 1 | DUF4291 family protein |
| | 8025650 | 8172811 | 147161 | Island Pick and IslandPath-DIMOB | WP_070390106.1 | BC342_RS34870 | 8119191 | 8119823 | -1 | hypothetical protein |
| | 8025650 | 8172811 | 147161 | Island Pick and IslandPath-DIMOB | WP_070390107.1 | BC342_RS34875 | 8120019 | 8121062 | -1 | hypothetical protein |
| | 8025650 | 8172811 | 147161 | Island Pick and IslandPath-DIMOB | WP_107405230.1 | BC342_RS37255 | 8121154 | 8121495 | 1 | helix-turn-helix transcriptional regulator |
| | 8025650 | 8172811 | 147161 | Island Pick and IslandPath-DIMOB | WP_079155158.1 | BC342_RS36480 | 8122467 | 8122946 | 1 | hypothetical protein |
| | 8025650 | 8172811 | 147161 | Island Pick and IslandPath-DIMOB | WP_107405375.1 | BC342_RS34900 | 8122943 | 8124439 | 1 | ATP-dependent helicase |
| | 8025650 | 8172811 | 147161 | Island Pick and IslandPath-DIMOB | WP_070390113.1 | BC342_RS34905 | 8125175 | 8128555 | 1 | RHS repeat-associated core domain-containing protein |
| | 8025650 | 8172811 | 147161 | Island Pick and IslandPath-DIMOB | | BC342_RS34910 | 8129626 | 8129910 | -1 | IS630 family transposase |
| | 8025650 | 8172811 | 147161 | Island Pick and IslandPath-DIMOB | WP_070390114.1 | BC342_RS34915 | 8130188 | 8130613 | 1 | SRPBCC family protein |
| | 8025650 | 8172811 | 147161 | Island Pick and IslandPath-DIMOB | | BC342_RS36485 | 8130942 | 8131115 | -1 | VOC family protein |
| | 8025650 | 8172811 | 147161 | Island Pick and IslandPath-DIMOB | WP_123937990.1 | BC342_RS34925 | 8131592 | 8131951 | -1 | transposase |
| | 8025650 | 8172811 | 147161 | Island Pick and IslandPath-DIMOB | WP_070390556.1 | BC342_RS34930 | 8132352 | 8133011 | 1 | HNH endonuclease |
| | 8025650 | 8172811 | 147161 | Island Pick and IslandPath-DIMOB | | BC342_RS34935 | 8133117 | 8133317 | 1 | hypothetical protein |
| | 8025650 | 8172811 | 147161 | Island Pick and | WP_070390116.1 | BC342_RS34940 | 8133499 | 8134689 | -1 | helix-turn-helix domain-containing |

| | | | | | | | | | | |
|---|---|---|---|---|---|---|---|---|---|---|
| | | | | IslandPath-DIMOB | | | | | | protein |
| | 8025650 | 8172811 | 147161 | Island Pick and IslandPath-DIMOB | WP_037772213.1 | BC342_RS34945 | 8135099 | 8135374 | -1 | hypothetical protein |
| | 8025650 | 8172811 | 147161 | Island Pick and IslandPath-DIMOB | WP_123937992.1 | BC342_RS37260 | 8135776 | 8136273 | -1 | CHAT domain-containing protein |
| | 8025650 | 8172811 | 147161 | Island Pick and IslandPath-DIMOB | WP_079155249.1 | BC342_RS34950 | 8136327 | 8136536 | -1 | hypothetical protein |
| | 8025650 | 8172811 | 147161 | Island Pick and IslandPath-DIMOB | WP_123937994.1 | BC342_RS34955 | 8136840 | 8137049 | 1 | hypothetical protein |
| | 8025650 | 8172811 | 147161 | Island Pick and IslandPath-DIMOB | WP_107405232.1 | BC342_RS34960 | 8137262 | 8137801 | 1 | hypothetical protein |
| | 8025650 | 8172811 | 147161 | Island Pick and IslandPath-DIMOB | WP_070390119.1 | BC342_RS34965 | 8138057 | 8138302 | 1 | DUF3761 domain-containing protein |
| | 8025650 | 8172811 | 147161 | Island Pick and IslandPath-DIMOB | WP_070390120.1 | BC342_RS34970 | 8138561 | 8139271 | 1 | HNH endonuclease |
| | 8025650 | 8172811 | 147161 | Island Pick and IslandPath-DIMOB | WP_070390121.1 | BC342_RS34975 | 8139268 | 8139558 | 1 | hypothetical protein |
| | 8025650 | 8172811 | 147161 | Island Pick and IslandPath-DIMOB | WP_070390122.1 | BC342_RS34980 | 8140230 | 8140733 | 1 | lamin tail domain-containing protein |
| | 8025650 | 8172811 | 147161 | Island Pick and IslandPath-DIMOB | | BC342_RS34985 | 8140831 | 8141085 | -1 | HNH endonuclease |
| | 8025650 | 8172811 | 147161 | Island Pick and IslandPath-DIMOB | | BC342_RS34990 | 8141088 | 8141768 | 1 | IS110 family transposase |
| | 8025650 | 8172811 | 147161 | Island Pick and IslandPath-DIMOB | | BC342_RS34995 | 8142090 | 8142375 | -1 | hypothetical protein |
| | 8025650 | 8172811 | 147161 | Island Pick and IslandPath-DIMOB | WP_107405233.1 | BC342_RS35000 | 8142490 | 8145183 | 1 | hypothetical protein |
| | 8025650 | 8172811 | 147161 | Island Pick and IslandPath-DIMOB | | BC342_RS37270 | 8145315 | 8145419 | -1 | IS5/IS1182 family transposase |
| | 8025650 | 8172811 | 147161 | Island Pick and IslandPath-DIMOB | WP_037771683.1 | BC342_RS35005 | 8145582 | 8146313 | -1 | MBL fold metallo-hydrolase |
| | 8025650 | 8172811 | 147161 | Island Pick and | WP_031048538.1 | BC342_RS35010 | 8146976 | 8147158 | 1 | toxin-antitoxin system HicB family |

| | | | | | | | | | | |
|---|---|---|---|---|---|---|---|---|---|---|
| | | | | IslandPath-DIMOB | | | | | | antitoxin |
| | 8025650 | 8172811 | 147161 | Island Pick and IslandPath-DIMOB | WP_070390126.1 | BC342_RS35015 | 8147155 | 8147451 | 1 | hypothetical protein |
| | 8025650 | 8172811 | 147161 | Island Pick and IslandPath-DIMOB | WP_079155163.1 | BC342_RS35020 | 8147526 | 8148077 | 1 | kinase |
| | 8025650 | 8172811 | 147161 | Island Pick and IslandPath-DIMOB | WP_070390128.1 | BC342_RS35030 | 8148466 | 8148660 | -1 | hypothetical protein |
| | 8025650 | 8172811 | 147161 | Island Pick and IslandPath-DIMOB | WP_123937996.1 | BC342_RS35035 | 8148684 | 8149055 | -1 | hypothetical protein |
| | 8025650 | 8172811 | 147161 | Island Pick and IslandPath-DIMOB | | BC342_RS36490 | 8149212 | 8149489 | 1 | hypothetical protein |
| | 8025650 | 8172811 | 147161 | Island Pick and IslandPath-DIMOB | WP_070390130.1 | BC342_RS35040 | 8149778 | 8149999 | 1 | hypothetical protein |
| | 8025650 | 8172811 | 147161 | Island Pick and IslandPath-DIMOB | WP_070390131.1 | BC342_RS35045 | 8150033 | 8150884 | -1 | helix-turn-helix domain-containing protein |
| | 8025650 | 8172811 | 147161 | Island Pick and IslandPath-DIMOB | WP_070390132.1 | BC342_RS35050 | 8151064 | 8151864 | 1 | oxidoreductase |
| | 8025650 | 8172811 | 147161 | Island Pick and IslandPath-DIMOB | WP_078536146.1 | BC342_RS36495 | 8151914 | 8152225 | 1 | hypothetical protein |
| | 8025650 | 8172811 | 147161 | Island Pick and IslandPath-DIMOB | | BC342_RS35055 | 8152393 | 8153387 | 1 | RacO protein |
| | 8025650 | 8172811 | 147161 | Island Pick and IslandPath-DIMOB | WP_031031886.1 | BC342_RS35060 | 8153390 | 8153809 | 1 | hypothetical protein |
| | 8025650 | 8172811 | 147161 | Island Pick and IslandPath-DIMOB | WP_037772147.1 | BC342_RS35065 | 8153876 | 8154718 | 1 | IS5 family transposase |
| | 8025650 | 8172811 | 147161 | Island Pick and IslandPath-DIMOB | WP_031031882.1 | BC342_RS35070 | 8154769 | 8154987 | -1 | hypothetical protein |
| | 8025650 | 8172811 | 147161 | Island Pick and IslandPath-DIMOB | WP_070390559.1 | BC342_RS35075 | 8155244 | 8156233 | -1 | AraC family transcriptional regulator |
| | 8025650 | 8172811 | 147161 | Island Pick and IslandPath-DIMOB | WP_070390133.1 | BC342_RS35080 | 8156265 | 8157179 | 1 | alpha/beta hydrolase |
| | 8025650 | 8172811 | 147161 | Island Pick and | | BC342_RS36500 | 8157529 | 8157847 | 1 | transposase |

| | | | | | | | | | | |
|---|---|---|---|---|---|---|---|---|---|---|
| | | | | IslandPath-DIMOB | | | | | | |
| | 8025650 | 8172811 | 147161 | Island Pick and IslandPath-DIMOB | WP_070390134.1 | BC342_RS35085 | 8157992 | 8158960 | 1 | SDR family NAD(P)-dependent oxidoreductase |
| | 8025650 | 8172811 | 147161 | Island Pick and IslandPath-DIMOB | | BC342_RS37275 | 8158957 | 8159121 | 1 | TetR/AcrR family transcriptional regulator |
| | 8025650 | 8172811 | 147161 | Island Pick and IslandPath-DIMOB | WP_070390136.1 | BC342_RS35095 | 8159586 | 8160857 | 1 | M24 family metallopeptidase |
| | 8025650 | 8172811 | 147161 | Island Pick and IslandPath-DIMOB | WP_070390137.1 | BC342_RS35100 | 8160978 | 8161292 | -1 | transcriptional regulator |
| | 8025650 | 8172811 | 147161 | Island Pick and IslandPath-DIMOB | WP_123937998.1 | BC342_RS37470 | 8161546 | 8161731 | -1 | hypothetical protein |
| | 8025650 | 8172811 | 147161 | Island Pick and IslandPath-DIMOB | | BC342_RS37280 | 8161733 | 8161858 | -1 | IS5/IS1182 family transposase |
| | 8025650 | 8172811 | 147161 | Island Pick and IslandPath-DIMOB | WP_070390139.1 | BC342_RS35110 | 8162200 | 8162796 | 1 | helix-turn-helix domain-containing protein |
| | 8025650 | 8172811 | 147161 | Island Pick and IslandPath-DIMOB | WP_070390140.1 | BC342_RS35115 | 8162823 | 8164934 | 1 | hypothetical protein |
| | 8025650 | 8172811 | 147161 | Island Pick and IslandPath-DIMOB | WP_070390141.1 | BC342_RS37285 | 8164934 | 8165653 | 1 | helix-turn-helix domain containing protein |
| | 8025650 | 8172811 | 147161 | Island Pick and IslandPath-DIMOB | WP_070390142.1 | BC342_RS35125 | 8165937 | 8166293 | -1 | hypothetical protein |
| | 8025650 | 8172811 | 147161 | Island Pick and IslandPath-DIMOB | WP_079155166.1 | BC342_RS36510 | 8166379 | 8166582 | -1 | hypothetical protein |
| | 8025650 | 8172811 | 147161 | Island Pick and IslandPath-DIMOB | WP_123938000.1 | BC342_RS37475 | 8166679 | 8166867 | -1 | hypothetical protein |
| | 8025650 | 8172811 | 147161 | Island Pick and IslandPath-DIMOB | WP_070390144.1 | BC342_RS35135 | 8167664 | 8170015 | -1 | XRE family transcriptional regulator |
| | 8025650 | 8172811 | 147161 | Island Pick and IslandPath-DIMOB | WP_070390146.1 | BC342_RS35145 | 8171127 | 8172332 | 1 | hypothetical protein |
| | 8025650 | 8172811 | 147161 | Island Pick and IslandPath-DIMOB | WP_070390147.1 | BC342_RS35150 | 8172329 | 8172811 | 1 | hypothetical protein |
| 5 | 8026607 | 8030971 | 4364 | SIGI-HMM | WP_070390079.1 | BC342_RS34585 | 8026607 | 8027449 | 1 | NAD(P)-dependent oxidoreductase |

| | | | | | | | | | | |
|---|---|---|---|---|---|---|---|---|---|---|
| | 8026607 | 8030971 | 4364 | SIGI-HMM | WP_037772601.1 | BC342_RS34590 | 8027542 | 8028015 | 1 | nuclear transport factor 2 family protein |
| | 8026607 | 8030971 | 4364 | SIGI-HMM | | BC342_RS34595 | 8028096 | 8029092 | 1 | thioesterase |
| | 8026607 | 8030971 | 4364 | SIGI-HMM | WP_078536178.1 | BC342_RS34600 | 8029089 | 8030315 | 1 | cytochrome P450 |
| | 8026607 | 8030971 | 4364 | SIGI-HMM | WP_123937984.1 | BC342_RS36425 | 8030672 | 8030971 | 1 | hypothetical protein |
| | 8026607 | 8030971 | 4364 | SIGI-HMM | WP_031047091.1 | BC342_RS34605 | 8030968 | 8032149 | 1 | LLM class flavin-dependent oxidoreductase |
| 6 | 8033093 | 8051282 | 18189 | Island Pick and IslandPath-DIMOB | WP_031047088.1 | BC342_RS34610 | 8032407 | 8033240 | 1 | SDR family oxidoreductase |
| | 8033093 | 8051282 | 18189 | Island Pick and IslandPath-DIMOB | | BC342_RS34615 | 8033502 | 8033938 | 1 | hypothetical protein |
| | 8033093 | 8051282 | 18189 | Island Pick and IslandPath-DIMOB | WP_070390081.1 | BC342_RS34620 | 8034080 | 8035444 | 1 | crotonyl-CoA carboxylase/reductase |
| | 8033093 | 8051282 | 18189 | Island Pick and IslandPath-DIMOB | | BC342_RS34625 | 8036200 | 8038975 | 1 | helix-turn-helix transcriptional regulator |
| | 8033093 | 8051282 | 18189 | Island Pick and IslandPath-DIMOB | WP_070390082.1 | BC342_RS34630 | 8039390 | 8040859 | 1 | SAVED domain-containing protein |
| | 8033093 | 8051282 | 18189 | Island Pick and IslandPath-DIMOB | WP_070390083.1 | BC342_RS34635 | 8041289 | 8042275 | -1 | hypothetical protein |
| | 8033093 | 8051282 | 18189 | Island Pick and IslandPath-DIMOB | WP_070387189.1 | BC342_RS34640 | 8042389 | 8043714 | 1 | ISL3 family transposase |
| | 8033093 | 8051282 | 18189 | Island Pick and IslandPath-DIMOB | WP_070387188.1 | BC342_RS34645 | 8045212 | 8046216 | 1 | nucleotidyltransferase |
| | 8033093 | 8051282 | 18189 | Island Pick and IslandPath-DIMOB | WP_070387187.1 | BC342_RS34650 | 8046213 | 8047679 | 1 | ThiF family adenylyltransferase |
| | 8033093 | 8051282 | 18189 | Island Pick and IslandPath-DIMOB | | BC342_RS36430 | 8047676 | 8048203 | 1 | hypothetical protein |
| | 8033093 | 8051282 | 18189 | Island Pick and IslandPath-DIMOB | | BC342_RS36435 | 8048298 | 8048636 | -1 | hypothetical protein |
| | 8033093 | 8051282 | 18189 | Island Pick and IslandPath-DIMOB | WP_070387186.1 | BC342_RS34655 | 8048719 | 8049045 | -1 | hypothetical protein |
| | 8033093 | 8051282 | 18189 | Island Pick and IslandPath-DIMOB | WP_070390084.1 | BC342_RS34660 | 8049152 | 8050225 | -1 | ATP/GTP-binding protein |
| | 8033093 | 8051282 | 18189 | Island Pick and | WP_107405374.1 | BC342_RS34665 | 8050225 | 8052336 | -1 | DDE-type |

| | | | | | | | | | | |
|---|---|---|---|---|---|---|---|---|---|---|
| | | | | IslandPath-DIMOB | | | | | | integrase/transposase/recombinase |

**Table S98.** Predicted genomic islands in abyssomicin BGC from *Streptomyces* sp. Amel2xE9 (NZ_KB912999 and NZ_KB912981).

| Island number | Island start | Island end | Length | Method | Gene name | Locus | Gene start | Gene end | Strand | Product |
|---|---|---|---|---|---|---|---|---|---|---|
| 1 | 252366 | 258784 | 6418 | SIGI-HMM | WP_019985544.1 | B065_RS38560 | 252366 | 252560 | -1 | hypothetical protein |
| | 252366 | 258784 | 6418 | SIGI-HMM | WP_019985545.1 | B065_RS0132280 | 252764 | 253537 | 1 | AfsR/SARP family transcriptional regulator |
| | 252366 | 258784 | 6418 | SIGI-HMM | WP_019985546.1 | B065_RS0132285 | 254243 | 257080 | -1 | LuxR family transcriptional regulator |
| | 252366 | 258784 | 6418 | SIGI-HMM | WP_019985547.1 | B065_RS0132290 | 257354 | 258784 | -1 | MFS transporter |
| 2 | 264101 | 271884 | 7783 | SIGI-HMM | WP_019985554.1 | B065_RS0132325 | 264101 | 264307 | -1 | ferredoxin |
| | 264101 | 271884 | 7783 | SIGI-HMM | WP_027758724.1 | B065_RS0132330 | 264313 | 265506 | -1 | cytochrome P450 |
| | 264101 | 271884 | 7783 | SIGI-HMM | WP_019985556.1 | B065_RS0132335 | 266764 | 268431 | -1 | ABC transporter ATP-binding protein |
| | 264101 | 271884 | 7783 | SIGI-HMM | WP_106962190.1 | B065_RS0132340 | 268428 | 269177 | -1 | ABC transporter permease |
| | 264101 | 271884 | 7783 | SIGI-HMM | WP_019985558.1 | B065_RS0132345 | 269234 | 270235 | -1 | ABC transporter permease |
| | 264101 | 271884 | 7783 | SIGI-HMM | WP_027758725.1 | B065_RS0132350 | 270220 | 271884 | -1 | ABC transporter substrate-binding protein |

**Table S99.** Predicted genomic islands in potential abyssomicin BGC from *Streptomyces incarnatus* NRRL 8089 (CP011497).

| Island number | Island start | Island end | Length | Method | Gene name | Locus | Gene start | Gene end | Strand | Product |
|---|---|---|---|---|---|---|---|---|---|---|
| 1 | 395452 | 405212 | 9760 | IslandPick | AKJ08807.1 | ABB07_01775 | 395560 | 396579 | -1 | luciferase |
| | 395452 | 405212 | 9760 | IslandPick | AKJ08808.1 | ABB07_01780 | 396806 | 398005 | 1 | cytochrome P450 |
| | 395452 | 405212 | 9760 | IslandPick | AKJ08809.1 | ABB07_01785 | 398038 | 398253 | 1 | hypothetical protein |
| | 395452 | 405212 | 9760 | IslandPick | AKJ08810.1 | ABB07_01790 | 398213 | 399256 | -1 | luciferase |
| | 395452 | 405212 | 9760 | IslandPick | AKJ08811.1 | ABB07_01795 | 399560 | 401053 | -1 | MFS transporter |
| | 395452 | 405212 | 9760 | IslandPick | AKJ08812.1 | ABB07_01800 | 401255 | 401860 | 1 | TetR family transcriptional regulator |
| | 395452 | 405212 | 9760 | IslandPick | AKJ08813.1 | ABB07_01805 | 401950 | 403293 | 1 | nitrilotriacetate monooxygenase |
| | 395452 | 405212 | 9760 | IslandPick | AKJ08814.1 | ABB07_01810 | 403277 | 404821 | -1 | hypothetical protein |
| | 395452 | 405212 | 9760 | IslandPick | AKJ08815.1 | ABB07_01815 | 404898 | 405710 | -1 | hypothetical protein |

**Table S100.** Predicted genomic islands in potential BGC from *Micromonospora eburnea* DSM 44814 (NZ_FMHY01000002.1).

| Island number | Island start | Island end | Length | Method | Gene name | Locus | Gene start | Gene end | Strand | Product |
|---|---|---|---|---|---|---|---|---|---|---|
| 1 | 4701227 | 4709030 | 7803 | IslandPick | WP_091120850.1 | GA0070604_RS20495 | 4700484 | 4701254 | 1 | LLM class F420-dependent oxidoreductase |
| | 4701227 | 4709030 | 7803 | IslandPick | | GA0070604_RS20500 | 4702315 | 4704202 | 1 | hypothetical protein |
| | 4701227 | 4709030 | 7803 | IslandPick | | GA0070604_RS20505 | 4703926 | 4705122 | 1 | hypothetical protein |
| | 4701227 | 4709030 | 7803 | IslandPick | | GA0070604_RS20510 | 4706074 | 4706952 | 1 | hypothetical protein |
| | 4701227 | 4709030 | 7803 | IslandPick | | GA0070604_RS20515 | 4707523 | 4708200 | 1 | hypothetical protein |
| | 4701227 | 4709030 | 7803 | IslandPick | | GA0070604_RS20520 | 4708213 | 4708737 | 1 | hypothetical protein |
| | 4701227 | 4709030 | 7803 | IslandPick | | GA0070604_RS20525 | 4708825 | 4709556 | 1 | hypothetical protein |
| | 4709107 | 4714377 | 5270 | IslandPick | | GA0070604_RS20525 | 4708825 | 4709556 | 1 | hypothetical protein |
| | 4709107 | 4714377 | 5270 | IslandPick | | GA0070604_RS20530 | 4709884 | 4710294 | 1 | hypothetical protein |
| | 4709107 | 4714377 | 5270 | IslandPick | | GA0070604_RS20535 | 4711306 | 4712406 | 1 | hypothetical protein |
| | 4709107 | 4714377 | 5270 | IslandPick | | GA0070604_RS20540 | 4712743 | 4713576 | 1 | malonyl CoA-ACP transacylase |
| | 4709107 | 4714377 | 5270 | IslandPick | | GA0070604_RS20545 | 4714183 | 4715653 | 1 | hypothetical protein |
| 2 | 4716822 | 4723967 | 7145 | IslandPick | WP_091120853.1 | GA0070604_RS20550 | 4715710 | 4719729 | 1 | SDR family NAD(P)-dependent oxidoreductase |
| | 4716822 | 4723967 | 7145 | IslandPick | WP_091120857.1 | GA0070604_RS20555 | 4719968 | 4720201 | 1 | hypothetical protein |
| | 4716822 | 4723967 | 7145 | IslandPick | WP_091120861.1 | GA0070604_RS20560 | 4720423 | 4721184 | -1 | hypothetical protein |
| | 4716822 | 4723967 | 7145 | IslandPick | WP_091120868.1 | GA0070604_RS20565 | 4721598 | 4722908 | 1 | cytochrome P450 |
| | 4716822 | 4723967 | 7145 | IslandPick | WP_091120872.1 | GA0070604_RS20570 | 4723627 | 4725360 | 1 | ABC transporter ATP-binding protein |
| | 4735352 | 4739428 | 4076 | IslandPick | WP_091120887.1 | GA0070604_RS20590 | 4730526 | 4735802 | -1 | type I polyketide synthase |
| | 4735352 | 4739428 | 4076 | IslandPick | WP_091120890.1 | GA0070604_RS20595 | 4735878 | 4736921 | -1 | 3-oxoacyl-ACP synthase |
| | 4735352 | 4739428 | 4076 | IslandPick | WP_091120894.1 | GA0070604_RS20600 | 4736988 | 4738196 | -1 | cytochrome P450 |
| | 4735352 | 4739428 | 4076 | IslandPick | WP_091120898.1 | GA0070604_RS20605 | 4738235 | 4738834 | -1 | hypothetical protein |
| | 4735352 | 4739428 | 4076 | IslandPick | WP_091120901.1 | GA0070604_RS20610 | 4739148 | 4744745 | 1 | type I polyketide synthase |
| 3 | 4759083 | 4764629 | 5546 | IslandPick | | GA0070604_RS20625 | 4756912 | 4759682 | 1 | hypothetical protein |
| | 4759083 | 4764629 | 5546 | IslandPick | | GA0070604_RS20630 | 4759384 | 4760877 | 1 | hypothetical protein |
| | 4759083 | 4764629 | 5546 | IslandPick | WP_091127296.1 | GA0070604_RS20635 | 4760929 | 4761408 | 1 | hypothetical protein |
| | 4759083 | 4764629 | 5546 | IslandPick | WP_091120909.1 | GA0070604_RS20640 | 4761408 | 4762883 | 1 | hypothetical protein |
| | 4759083 | 4764629 | 5546 | IslandPick | WP_091120913.1 | GA0070604_RS20645 | 4762944 | 4763975 | 1 | 3-oxoacyl-ACP synthase III family protein |
| | 4759083 | 4764629 | 5546 | IslandPick | | GA0070604_RS20650 | 4764056 | 4765045 | 1 | alpha/beta hydrolase |
| 4 | 4765348 | 4770810 | 5462 | IslandPick | WP_091120916.1 | GA0070604_RS20655 | 4765165 | 4765938 | -1 | AfsR/SARP family transcriptional regulator |
| | 4765348 | 4770810 | 5462 | IslandPick | WP_091120921.1 | GA0070604_RS20660 | 4765935 | 4766723 | -1 | thioesterase |
| | 4765348 | 4770810 | 5462 | IslandPick | WP_091120924.1 | GA0070604_RS20665 | 4767358 | 4769694 | -1 | AAA family ATPase |
| | 4765348 | 4770810 | 5462 | IslandPick | WP_091120927.1 | GA0070604_RS20670 | 4770202 | 4770429 | 1 | hypothetical protein |
| | 4765348 | 4770810 | 5462 | IslandPick | WP_091120932.1 | GA0070604_RS20675 | 4770588 | 4771895 | 1 | lipopolysaccharide biosynthesis protein RfbH |
